# FIND-IT: Ultrafast mining of genome diversity

**DOI:** 10.1101/2021.05.20.444969

**Authors:** Søren Knudsen, Toni Wendt, Christoph Dockter, Hanne C. Thomsen, Magnus Rasmussen, Morten Egevang Jørgensen, Qiongxian Lu, Cynthia Voss, Emiko Murozuka, Jeppe Thulin Østerberg, Jesper Harholt, Ilka Braumann, Jose A. Cuesta-Seijo, Sabrina Bodevin, Lise T. Petersen, Massimiliano Carciofi, Pai Rosager Pedas, Jeppe Opstrup Husum, Martin Toft Simmelsgaard Nielsen, Kasper Nielsen, Mikkel K. Jensen, Lillian A. Møller, Zoran Gojkovic, Alexander Striebeck, Klaus Lengeler, Ross T. Fennessy, Michael Katz, Rosa Garcia Sanchez, Natalia Solodovnikova, Jochen Förster, Ole Olsen, Birger Lindberg Møller, Geoffrey B. Fincher, Birgitte Skadhauge

## Abstract

Novel crop improvement methodologies, including the exploitation of natural genetic variation, are urgently required to feed our rapidly growing human population in the context of global climate change. Here we describe a ‘Fast Identification of Nucleotide variants by DigITal PCR’ (FIND-IT) method for the rapid identification of pre-targeted genetic variants or rare alleles in large genomic populations. Libraries of 500,000 individuals can be screened and desired variants isolated within two weeks. FIND-IT is widely applicable for mining valuable diversity in any genomic population, including elite breeding and wild germplasm collections. The method provides single nucleotide resolution that has been validated by identifying and isolating knockout lines, non-synonymous codon changes and variants of miRNA and transcription factor binding sites in the agronomically important crop barley. In contrast to existing methods, FIND-IT does not require transformation, cloning or enzymatic steps, and is exempt from GMO regulations. Thus, FIND-IT can be applied immediately to elite crop cultivars and can be tailored to minimize or eliminate time-consuming backcrossing requirements.

## Introduction

Domestication and subsequent improvement of cereal crops was initiated by human selection of traits for enhanced crop productivity and edibility some 10,000 years ago. Early Neolithic hunter-gatherers, peasants and later farmers, breeders, and scientists, using ever evolving technologies, have strived to improve annual average crop yields and quality. At a global scale, societies are now challenged by crop productivity stagnation across many environments, aggravated by climate change and soil nutrient depletion^1,2^. Current yield increase rates will not accommodate the projected food requirements of 9-10 billion people by 2050^3^, demanding a boost of sustainable crop improvement through new, targeted genome engineering technologies. This can be achieved through harnessing natural or induced genetic variation in crop species or their wild relatives and acceleration of the application of variants in elite breeding stock programmes^3^.

Despite advances in plant genome editing by CRISPR-Cas9 technologies, which hold great promise for the long term future of crop adaptation^4^, their use in commercial plant breeding is not yet widely applied. This is because *i*) they require elaborate transformation protocols and tissue culture steps that can adversely affect genome integrity^5,6^, *ii*) they are therefore non-compatible with most existing breeding pipelines, i.e. not directly applicable to elite, high-yielding, climate-resilient and regionally adapted breeding stock, and *iii*) they are considered GM-technologies in many countries.

Large, natural variant populations (e.g. crop fields and germplasm collections), with low spontaneous mutation densities have been the source of natural gene variation used in human endeavours to domesticate and adapt crops to superior performance, independent of changing environments and novel agricultural practices^7^. Induced mutagenesis increases variant densities in populations and has been investigated and utilized for crop development for nearly 100 years^8^, notably in the genetic improvements underlying the green revolution crop traits^9^. In particular, the more recently developed chemical treatments, which primarily induce single nucleotide substitutions^10^, can be used to obtain large variant populations and eventually single nucleotide resolution, where a substitution of any genome nucleotide is present with high probability.

Traditional, phenotype-based methods to screen natural or induced variant populations for desirable genetic variants are painstakingly slow and labour intensive. Targeted screening methods like TILLING (Targeting Induced Local Lesions IN Genomes)^11^ or EcoTILLING^12^, provide genome-wide scanning of induced and natural variant populations directly at the genotype level. However, technical hurdles and low detection sensitivity usually restrict population sizes to fewer than 10,000 individuals. This prevents the achievement of single nucleotide resolution or, alternatively, demands very high mutation rates per genome^13^, which result in high off-target background mutation loads that are undesirable in downstream breeding applications^14^.

Here, we present a simple, agile, and high-throughput strategy to screen extremely large, low mutation-density variant populations, scalable to single nucleotide resolution, for targeted identification of desired traits that are immediately applicable in crop or organism improvement strategies. In this ‘<u>F</u>ast <u>I</u>dentification of <u>N</u>ucleotide variants by <u>D</u>ig<u>IT</u>al PCR’ (FIND-IT) method, we induce, identify and isolate specific targeted nucleotide variants by combining systematic sample pooling-and-splitting with high-sensitivity genotyping. FIND-IT *i*) does not require transformation or tissue culture protocols, *ii*) is applicable to any living organism that can be grown in the field or in culture (i.e. elite plant crop varieties, bacteria, yeasts), and *iii*) is exempt from GMO regulation. Our pooling strategy reduces the time required for DNA isolation up to 300-fold and, using cultivated barley as an example, we show that it is feasible to screen, identify and isolate pre-targeted variants within two weeks, in large variant libraries (>500,000 individuals) with low individual mutation loads. This reduces or eliminates extensive backcrossing requirements and thereby greatly reduces the time from variant detection to field evaluation. By targeting specified single-nucleotide changes in genes that modify barley crop performance, grain morphology and grain quality properties, we demonstrate how FIND-IT circumvents limitations of comparable techniques and provides a novel methodology for ultrafast variant mining and improvement of today’s germplasm, through precision breeding.

## Results

### Designing libraries for targeted variant identification

The mutational load and the type of genetic variance introduced by mutagenesis form the basis for successful identification of the desired genetic variant. Chemical mutagenesis has a major advantage over radiation-induced mutagenesis by predominantly causing single nucleotide changes and, depending on the respective library size, facilitating single nucleotide resolution without additional changes associated with base/nucleotide excision repair^14^. We generated large variant libraries in barley (Fig. 1a), wheat, rapeseed, oat, yeast and bacteria, using either sodium azide (NaN_3_) or ethyl methanesulfonate (EMS) as the mutagen (Supplementary Notes 1, Section 1). The spring barley libraries used here to demonstrate and validate the FIND-IT methodology were designed with the aim of obtaining a large population of 500,000 individual plants with 1,000 – 5,000 induced nucleotide substitutions per genome. The recent publication of the barley pan genome identified large structural chromosome rearrangements and approximately 1.5 million presence/absence variations (PAV) in the range of 50 basepairs to 1 million bases (Mb) between 20 pan genome assemblies^7^. Our summary statistics of genome variation in whole genome sequencing data from the 200 domesticated and 100 wild barley lines published with the barley pan genome revealed more than 223 million naturally occurring nucleotide substitutions, with 299 of 300 lines having more than 5 million nucleotide substitutions when compared to the barley reference genome Morex_V2 (Supplementary notes 1, Section 2). Thus, our aim of increasing the natural variation by 1,000 – 5,000 induced nucleotide substitutions per plant is extremely low in the context of existing genome variation and novel genome variation created by crossing events in traditional breeding.

**Fig. 1.**
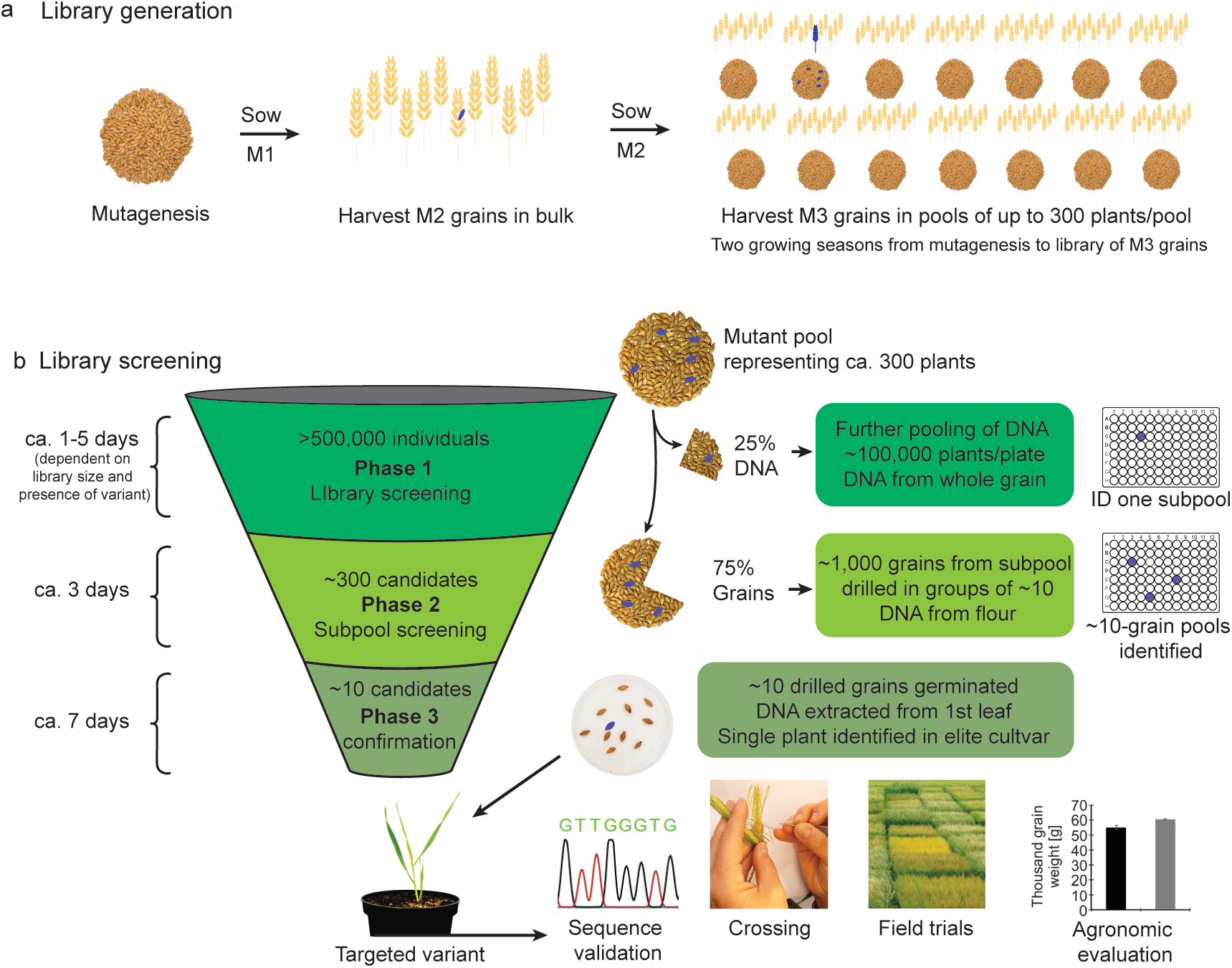
FIND-IT screening procedure. **a**, Barley library generation: Mutagenized barley plants are grown densely to maturity in the field. M3 grains are harvested in pools of approximately 300 plants. **b**, Library screening consists of three phases that narrow down the search of a target variant from >500,000 to eventually a single plant within days. See detailed description of the method in Main Text, Online Methods and Supplementary Notes 3, Section 1. Screening Phase 1: The harvested grain pools are each split into two fractions; one fraction (25%) is utilized for total DNA extraction, the other (75%) is stored for further analysis. DNA from the 25% pools is further pooled and one 96-well library plate contains DNA from >100,000 individuals. The following ddPCR screening (Fig. Supplementary Notes 3.2a), using competing fluorescently labelled probes to target the nucleotide variants of interest, identifies one well, representing ~300 candidates, for further analysis. Screening Phase 2: the corresponding 75% grain pool of the identified DNA pool in Phase 1 is selected for further analysis. DNA from approximately 1,000 grains is sampled non-destructively in pools of 10 and rescreened using ddPCR (Fig. Supplementary Notes 3.2b), and several wells are potentially identified to contain the targeted variant. Screening Phase 3: the grains from the identified wells in Phase 2 are germinated and total DNA is extracted from the first leaf of each individual plant and re-screened by ddPCR (Fig. Supplementary Notes 3.2c). From this, a single plant is identified containing the targeted variant which can be sequence-validated (Fig. Supplementary Notes 3.2d). Identified variants of interest are immediately available for field trials, agronomic analyses, and incorporation into routine breeding backcrossing programs.

### Determining mutation load of variant libraries

To define library mutation density, we applied two levels of characterization (Supplementary Notes 1, Section 3). We first Sanger sequenced four individual gene fragments in some 5,000 barley plants from the library treated with low NaN_3_ concentration (0.3 mM) (Table Supplementary Notes 1.3a). Here, mutation densities were in the range of 0.29 to 1.45 induced variants per Mb per plant. Given an average barley genome size of approx. 4.2 gigabases^7^, this equals approx. 1,200 to 6,000 induced variations per plant and is within the desired mutation density range.

In a complementary approach, Genotyping-By-Sequencing (GBS) of 40 barley individuals from low and high (0.3 and 1.67 mM NaN_3_) mutagen treated libraries (20 individuals/library) verified low mutation rates after 0.3 mM NaN_3_ treatment (0.46 to 0.9 induced variants/1Mb, accounting for approx. 2,000 to 3,750 induced variations per plant) and indicated a tendency for higher mutation rates after treatment with the higher NaN_3_ concentration (Table Supplementary Notes 1.3b). It should be noted that in 16 and 11 genotyped individuals chosen from both libraries no mutation was detected, respectively (Supplementary Notes 1, Section 3).

Thus, libraries can be designed to have the desired mutation load by controlling treatment intensity and population size.

### FIND-IT employs a split-and-pool strategy coupled with high-sensitivity droplet digital PCR (ddPCR)

To facilitate high-throughput screening of >500,000 individuals, we integrated a systematic and cost-effective split-and-pool strategy. The ddPCR technology is 1,000 fold more sensitive than conventional PCR^15^, and we confirmed its sensitivity by spiking pools of 1,000 wild-type barley grains with 4, 2, 1, or a half grain homozygous for a specific nucleotide exchange in the *LIPOXYGENASE-1* (*Lox-1*) gene. When targeted in total DNA of such dilution pools, the *Lox-1* variant signal achieved by ddPCR was clearly distinguishable from the background signal at all dilutions tested (Supplementary Notes 1, Section 4). We based the FIND-IT libraries on DNA pools representing grains from up to 1,200 individual M2 plants. For practical reasons, we harvested all material in pools of up to 300 plants, and only after DNA extraction we combined DNA samples to represent an estimate of 1,200 plants per pool (see Online Methods)^16^. The initial pooling strategy of up to 300 plants reduced the time required for DNA isolation by up to 300-fold, compared with screening strategies that use DNA from individual plants in the library. This removes a key technological bottleneck inherent in screening methods involving single plant sequencing or genotyping.

In each pool, multiple grains will originate from the same plant and hence a given induced variant will be represented several times in a pool. Thus, pools can be split into a 25% sample for initial large-scale genotyping while the remaining grain pool (75%) is stored for subsequent specific variant identification and isolation (Fig. 1b).

In Phase 1 of the FIND-IT procedure, DNA representing approximately 100,000 plants is screened on a single 96-well PCR plate. Here, 94 DNA pools (representing up to 1,200 plants each), plus two negative controls, are genotyped by ddPCR for an induced nucleotide target using a single set of primers and two competitive hydrolysis probes that detect either wild-type or variant alleles (Fig. 1b).

Positive pools containing the targeted variant are further analyzed in FIND-IT’s Phase 2 (Fig. 1b). Here, a new 96-well plate where each well contains DNA from 10 grains from the Phase 1 identified grain pool (75% of original grain pool, and likely to contain additional grains of the targeted variant) are ddPCR genotyped to identify a positive sub-pool with the targeted variant. The DNA is extracted from flour sampled by non-destructively drilling into the endosperm of individual seeds^17^. Thus, by combining a split-and-pool strategy with ddPCR genotyping, we can quickly narrow down the number of candidate grains carrying the targeted gene variant from 500,000 plants to 10 grains. Total DNA from drilled grains or drilled grains from sub-pools can be stored safely at −20°C. DNA from drilled grain sub-pools can be re-screened multiple times for other desired gene variants if identified in FIND-IT’s Phase 1, further reducing sub-pool generating workload. Finally (Phase 3; Fig. 1b), the 10 grains from the positive sub-pool are germinated and genotyped by ddPCR and Sanger sequenced for identification and isolation of the plant carrying the desired induced gene variant. If present in the library, the FIND-IT workflow enables the identification and isolation of a desired variant within days and weeks, respectively. During Phase 2 and 3, Phase 1 can be re-initiated for screening of new targets and Phase 3-targets can be accumulated for germination and validation (Supplementary Notes 1, Section 5), further accelerating the FIND-IT workflow.

### Modulating plant- and grain-morphology by targeting loss-of-function variants

To demonstrate FIND-IT’s high capacity to identify knockout variants, we identified 100 induced premature stop codons in specified barley genes (Fig. 2a, Supplementary Table 1 and Supplementary Notes 2). Of these, we isolated and field characterized five primary knockout variants of genes that mediate well-documented changes in phenotypic characteristics of the barley grain (Fig. 2b-h).

**Fig. 2.**
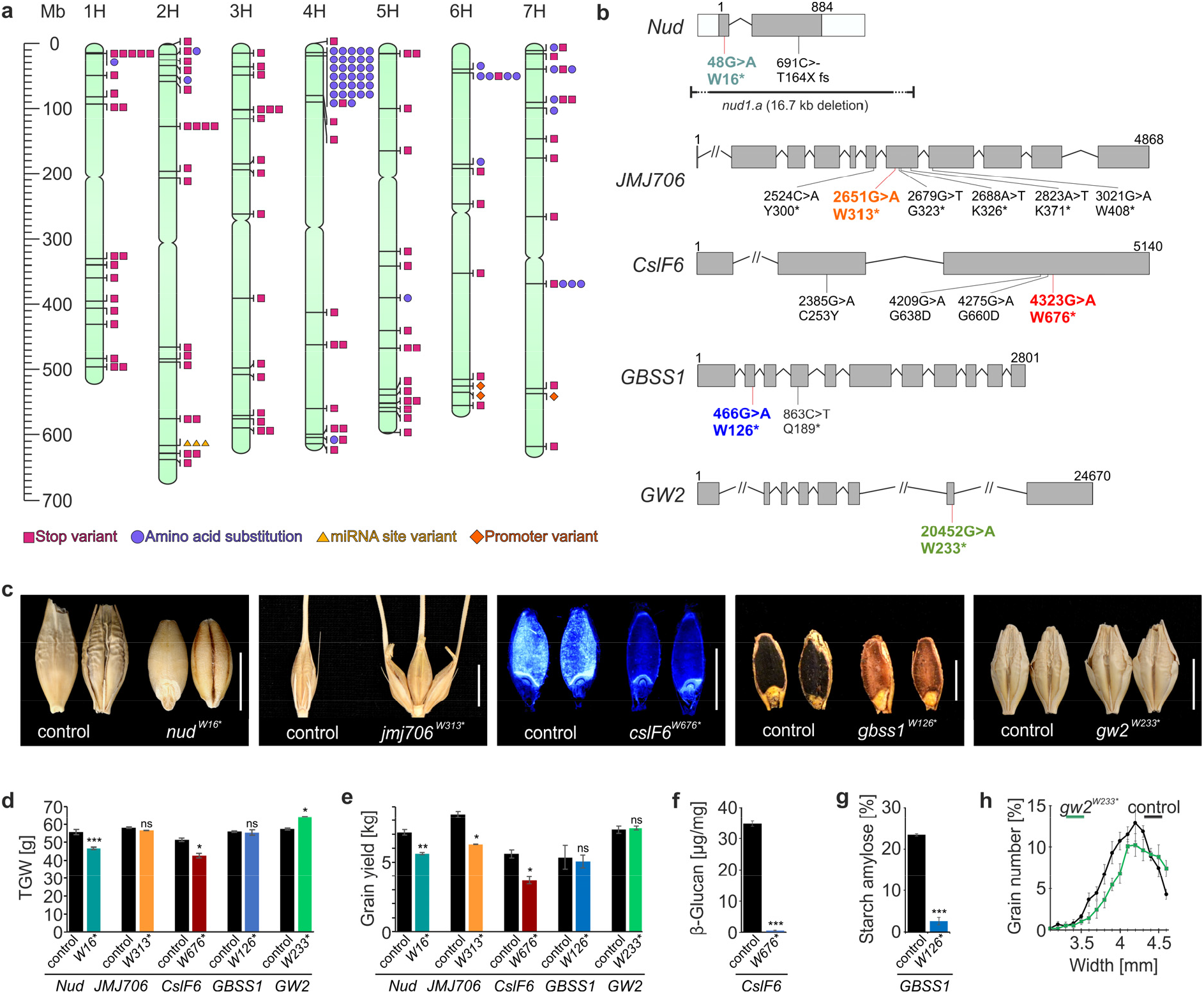
Barley knockout library with selected variants isolated and agronomically evaluated. **a**, Barley physical reference genome map with chromosomes 1H – 7H with more than 150 identified stop and other SNP mutations indicated by number (Supplementary Table 1). **b**, Gene models of selected barley reference targets with known loss-of-function mutations and novel, isolated stop variants (in colour). **c**, Loss-of-function phenotypes of novel variants (hulless grain, *nud^W16*^*; six-rowed spikelets, *jmj706^W313*^*; (1,3;1,4)-β-glucan Calcofluor staining, *cslF6^W676*^*; Starch amylose Lugol staining, *gbss1^W126*^*; grain width, *gw2^W233*^*). **d**, TGW and **e**, grain yield of field-grown novel barley variants. **f**, Grain (1,3;1,4)-β-glucan content of *cslF6^W676*^* grain. **g**, Starch amylose content of *gbss1^W126*^* grain. **h**, Grain width distribution of *gw2^W233*^* grain. Scale bars in c, 5 mm. Error bars are standard diviation, two-tailed t-test was performed to obtain P values (*P<0.05, **P<0.01, ***P<0.001, ns not statistically significant). See Supplementary Table 3 for statistical test details and number of replicates.

The *Nud* gene (Supplementary Notes 3, Section 1) encodes a transcription factor of the ethylene response factor (ERF) family that controls the covered/naked caryopsis trait in barley and knockout of this gene (Fig. 2b) converts hulled to hull-less grain^18^ (Fig. 2c). Using the FIND-IT technology, we isolated a novel premature stop allele of the *Nud* gene (*nud^W16*^*; Fig. 2b). Field trial evaluation of *nud^W16*^* in an elite spring barley cultivar (RGT Planet) background showed the expected loosely attached hull and accompanying reduced grain length, width, and thousand grain weight (TGW) after threshing and hull removal (Fig. 2b-e and Fig. Supplementary Notes 3.8a-e). This known domestication trait renders any wild, locally adapted or climate-resilient barley, or any elite hulled barley cultivar, suitable for human consumption and the ability to directly proceed to agronomic evaluation allows precise trait market value evaluation by comparison to elite hulled barley varieties.

The *JMJ706/VRS3* locus (Supplementary Notes 3, Section 1) encodes a putative Jumonji C-type H3K9me2/me3 histone demethylase that is involved in defining the barley inflorescence (spike) row-type, a yield-related domestication trait controlled by several genes^19^. We identified a novel homozygous loss-of-function variant in the *JMJ706* gene in cultivar Planet (*jmj706^W313*^*; Fig. 2b) that has the reported barley spikelet phenotype, important for grain size uniformity in six-rowed barley^19^ (Fig. 2c). Using FIND-IT, novel alleles of row-type genes, as shown here for *JMJ706/VRS3*, can be directly isolated from elite two- and six-rowed barley libraries to quickly optimize agronomic performance^19^.

The *Cellulose synthase like* gene *CslF6* (Supplementary Notes 3, Section 2) is a key determinant controlling the biosynthesis and structure of (1,3;1,4)-β-glucan in the barley grain, making it a prime target for lowering (1,3;1,4)-β-glucan^20^, a preferred grain trait for the distilling and brewing industries. Using FIND-IT we identified and isolated a novel *CslF6* knockout variant (*cslF6^W676*^*; Fig. 2b) in cultivar Planet with undetectable levels of (1,3;1,4)-β-glucan content in the grain (Fig. 2c,f). Field-testing for agronomic performance of *cslF6^W676*^* confirmed previously published greenhouse experiments on *CslF6* knockouts, namely that TGW and grain width were reduced and changed, respectively^21^ (Fig. 2d-f; Fig. Supplementary Notes 3.17a-f). Here, we show for the first time that total grain yield of *CslF6* knockout variants grown in the field is reduced by more than 30% (Fig. 2e), limiting the applicability of *CslF6* knockout varieties.

Granule-Bound Starch Synthase I (GBSS1) (Supplementary Notes 3, Section 1) is the enzyme for synthesizing amylose in barley^22^. We identified a variant carrying a novel loss-of-function allele (*gbss1^W126*^*; Fig. 2b) that features the characteristic waxy barley grain phenotype and the underlying reduction of amylose content in the grain starch (Fig. 2c,g), with no significant yield and TGW change (Fig 2d,e). Waxy starches are of high value in many processed foods as the amylose content directly influences the texture of cooked starches^23^. Now waxy variants of common cereals can be isolated from elite, high yielding varieties directly for market potential evaluation.

To illustrate the translation of agronomic traits from one cereal to another we used the FIND-IT methodology to induce a yield related loss-of-function trait *Grain Width 2* (*GW2)* (Supplementary Notes 3, Section 1), a modulator of grain morphology known in rice, into barley^24^ by isolating a novel loss-of-function allele in the elite malting barley Paustian (*gw2^W233*^*; Fig. 2b). To date, no knockout of barley *GW2* has been identified or studied^25^. Initial field yield trials (Denmark 2020) showed that *gw2^W233*^* has increased TGW, caused by wider grain size as shown in rice, but that total yield per area was not increased (Fig. 2c-e, h). Knockout variants of *OsGW2* improve grain yield in rice via acceleration of the grain milk filling rate, a trait of potential importance for cereals grown in suboptimal growth conditions such as drought and heat stress associated with shortened grain filling periods. The novel barley *gw2^W233*^*variant can now be field-tested in diverse climatic environments to identify its potential for yield improvement and climate adaptation in barley and other cereals.

### Targeting non-synonymous single-nucleotide variants and variants that modulate transcript abundance

The ability to finely regulate gene expression, transcript abundance and protein function is a highly desirable objective of genome engineering tools for precision breeding. FIND-IT enables the identification and isolation of such finely tuned gene variants. Here, 55 genetic variants with single, non-synonymous base-changes were targeted, either for changing transcript levels or by altering the amino acid sequences of encoded proteins (Fig. 2a, Supplementary Table 1, Supplementary Notes 2).

In one example we targeted the GROWTH-REGULATING FACTOR 4 (GRF4) transcription factor and the DELLA protein (SLENDER1/SLN1) that promote and repress growth, respectively, as well as influencing nitrogen assimilation and carbon fixation^26^ (Supplementary Notes 3, Section 2). In rice, enhanced nitrogen-use efficiency and grain yield are achieved by increasing *GRF4* transcript abundance and by reducing DELLA activity^26^. We identified 31 variants of *SLN1/DELLA* with specific amino acid exchanges (Fig. 3a, Supplementary Table 1) and characterized one variant *sln1^A277T^* (Fig. 3b-d), showing normal plant height, increased seed length and increased TGW but no change in yield (Fig. Supplementary Notes 3.4). Next, we identified three variants targeting the *miRNA396*-binding site of *GRF4* (Fig. 3e) with the aim of increasing GRF4 activity by reducing miRNA-mediated transcript repression. The genetic variants of *GRF4* and *SLN1/DELLA* identified will enable field trials aimed at generating new elite barley varieties with enhanced nitrogen-use efficiency^26^.

**Fig. 3.**
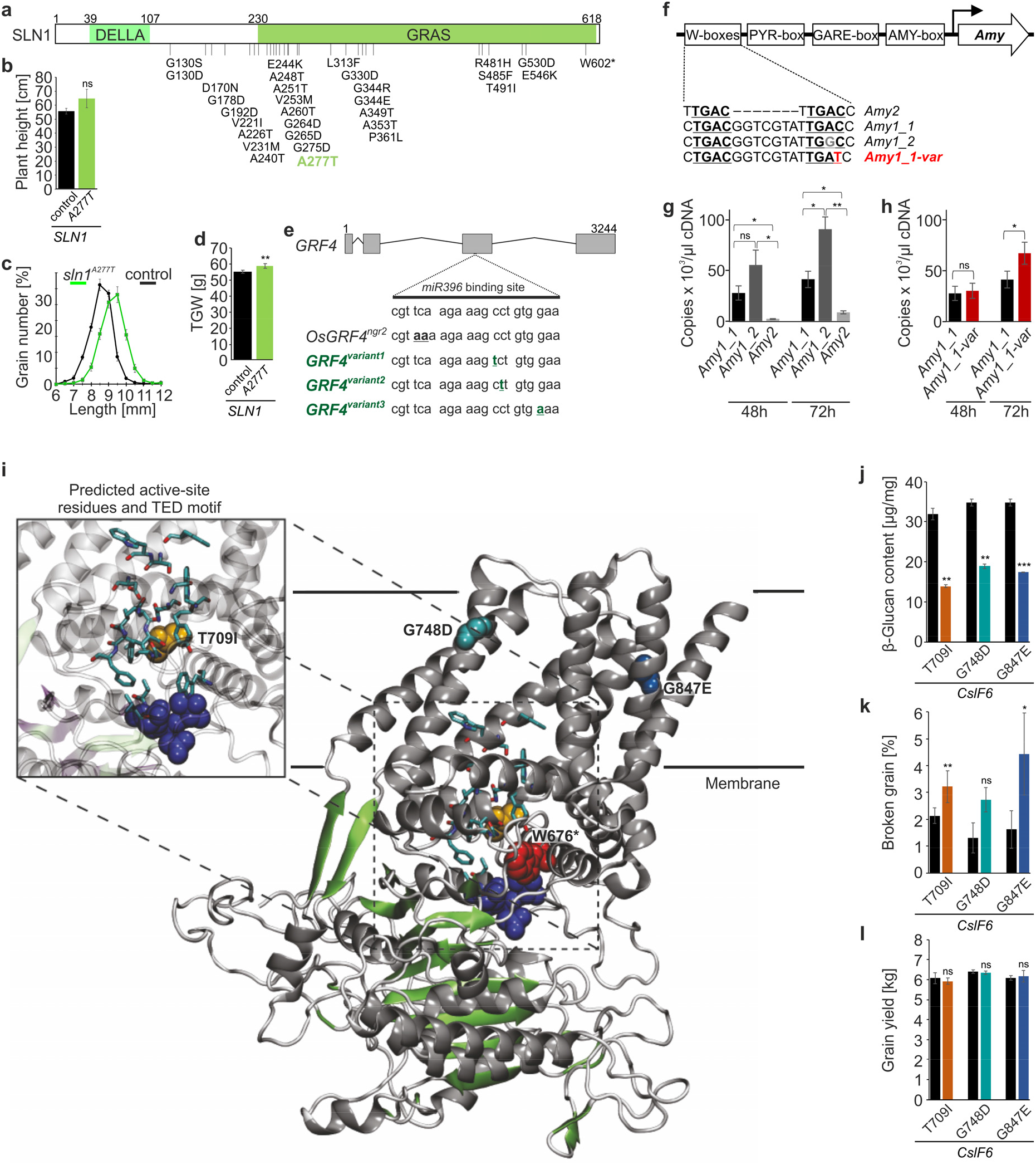
FIND-IT libraries with distinct non-synonymous nucleotide variants and variants to modulate transcript abundance for precision breeding. **a**, Identified and isolated variants (in black and green, respectively) of the SLN1/DELLA protein. **b**, Plant height, **c**, grain length distribution and **d**, TGW of *sln1^A277T^*. **e**, Gene model of barley *GRF4* showing the 21-bp *miR396* binding site on exon 3 including a known double substitution in rice (*OsGRF4^ngr2^*) and three novel barley variants (in green). **f**, Known cis-acting elements in α-amylase promoters with natural and induced W-box sequence variation. TGAC core motifs in W-boxes are underlined and in bold, natural variation in the *Amy1_2* gene in grey and a novel, induced variation in an *Amy1_1* gene in red. **g**, Expression of α-amylase genes in Planet grain 48h and 72h after initiation of germination. **h**, Expression of *Amy1_1* genes in Planet grain (black) and Planet variant *Amy1_1-var* (red) 48h and 72h after initiation of germination. **i**, Protein homology model of CslF6 based on poplar cellulose synthase isoform 8 (PDB# 6wlb.1.A). Amino acid positions at residue W676, T709, G748 and G847 are shown as spheres. Insert shows UDP-glucose binding sites – predicted by alignment of CslF6 and poplar cellulose synthase isoform 8 – as light blue sticks and the TED motif is shown as dark blue sticks. The insert is shown at a slightly tilted angle to visually optimize the position of the residues discussed. **j**, Grain (1,3;1,4)-β-glucan content of controls (black bars) and *CslF6* variants. **k**, Percentage of broken grain after threshing in controls (black bars) and *CslF6* variants. **l**, Field grain yield of controls (black bars) and *CslF6* variants. Error bars are standard diviation, two-tailed t-test was performed to obtain P values (*P<0.05, **P<0.01, ***P<0.001, ns not statistically significant). See Supplementary Table 3 for statistical test details and number of replicates.

In a second example, we targeted a promoter variant of a barley α-amylase gene to enhance gene expression and seed germination vigour in Planet barley (Fig. 3f and Supplementary Notes 3, Section 2). Expression of the *Amy1_2* gene in the germinated Planet control grain exceeded the expression of genes in the *Amy1_1* or *Amy2* clusters at 72 hours after germination^27^ (Fig. 3g). Inspired by natural variation in the tandem repeat W-box/O2S binding domain in α-amylase promoters, to relieve the repression by WRKY38^28^, we identified and isolated a novel allele of *Amy1_1* (*Amy1_1-var*) with a single nucleotide substitution in its promoter that changes one TGAC core motif to TGAT (Fig. 3f). We then demonstrated that the overall *Amy1_1* transcript level in the novel Planet variant *Amy1_1-var* increased 72h after initiation of germination by close to 100% relative to control (Fig. 3h), indicating that changing the TGAC to TGAT released repression of expression. This result shows the function of the tandem repeat W-box/O2S binding domain in α-amylase promoters *in vivo* and highlights the possibilities of modifying gene transcript levels by targeting promoter domains using FIND-IT, both for gene function analyses and for specific breeding purposes.

To isolate variants with an intermediate (1,3;1,4)-β-glucan content phenotype, to overcome the adverse yield and grain morphology phenotypes of the knockout line (Fig.2 c-e), we identified and isolated three additional *CslF6* variants with amino acid exchanges in close proximity to the TED motif that controls (1,3;1,4)-β-glucan fine structure (T709I)^29^, or within CslF6 transmembrane helices (G748D and G847E)^20^ (Fig. 3i and Supplementary Notes 3, Section 2). These novel *CslF6* variants have a 50% reduction in grain (1,3;1,4)-β-glucan levels but maintain elite barley yields and for G748D and G847E also TGW (Fig. 3j,l and Supplementary Notes 3, Section 2). While grains of variants *cslF6^G847E^* and *cslF6^T709I^* are susceptible to breakage in hard mechanical threshing (Fig. 3k), variant *cslF6^G748D^* is proving valuable in malting and brewing processes, where high (1,3;1,4)-β-glucan causes filtration difficulties and undesirable hazes in the final product^30^.

### FIND-IT beyond barley

The broad applicability of FIND-IT makes it possible to screen large variant libraries of any living organism that can be grown in the field or in culture, as highlighted here by additional examples in wheat, yeast (*Saccharomyces cerevisiae)* and bacteria (*Lactobacillus pasteurii)* (Supplementary Notes 3, Section 3). Diverse libraries of rapeseed and oat are also ready for variant targeting (Supplementary Notes 1, Section 1).

## Discussion

Here we have described and validated the FIND-IT method as a new species- and variety-agnostic method for untargeted generation and targeted isolation of single nucleotide variants in genome pools of plants and other biological systems. FIND-IT relies on three crucial components, *i*) the century old method of induced mutagenesis, especially through chemical treatment, *ii*) a sample pool-and-split strategy, and *iii)* high-sensitivity ddPCR technology^15^. Based on the integrated use of these three components, FIND-IT allows the size of routinely screen-able variant libraries to be expanded beyond 500,000 individuals and enables variant pool generation towards single nucleotide resolution. The size of the library and its mutation load are inversely correlated with respect to the probability of finding the variant of interest. FIND-IT allows each of these two parameters to be independently adjusted to meet the intended application of the library (e.g. for direct incorporation into a breeding pipeline or for functional analysis of individual genes).

In the present study, we chose low or high mutagen concentrations in combination with large library sizes to demonstrate the versatility of the method and to optimize the fast inclusion of the identified variant of interest into breeding programs with no or little backcrossing required. We show that very large libraries can be efficiently handled using the demonstrated protocol and that the low overall library mutation load achieved here nevertheless provides a high probability of finding *i*) a knockout of a specified target gene, *ii*) a non-synonymous variant in a single target codon and *iii*) a single nucleotide variant in a promoter element or a miRNA binding site. The total number of more than 150 targeted and successfully identified single nucleotide variants show the high-capacity of the FIND-IT method. We demonstrate the strength of FIND-IT to quickly evolve elite germplasms in accelerated commercial precision breeding through isolation, detailed analysis, and direct field assessment of multiple variants. These are either novel or mimicked from existing natural cereal variants and improve important agronomical or grain quality traits.

Precision breeding by CRISPR-Cas9 mediated genome editing of crop plants is rapidly developing despite unwanted off-target edits^4,31^, which are challenging to identify^32^. However, full applicability to commercial breeding pipelines awaits further improvements, such as solving the need for time-consuming and potentially chromosome rearranging transformation and/or tissue culture steps that limit the technology to the relatively small number of plant varieties that are transformable and will grow in culture^33^, or where *de novo* meristem induction can be used^34^. Across multiple jurisdictions, lines generated by CRISPR-Cas9 currently fall under GMO guidelines^35^, and are subject to the regulatory barriers associated with GMO methodologies. These intrinsic drawbacks and uncertainties in the accompanying legislation of CRISPR-Cas9 driven gene editing currently precludes its widespread utilization for accelerated development of crops with desired traits. Other traditional breeding methods dependent on phenotypic or targeted selection from natural or induced genome pools are either slow and imprecise or limited in population size and therefore restricted in their resolution. FIND-IT releases the bottlenecks of phenotypic selection and TILLING, and stands together with evolving CRISPR-Cas9 technologies for the efficient improvement of today’s germplasms.

We have successfully initiated FIND-IT based targeting of single nucleotide variants in elite-varieties of barley, wheat, and in industrial production strains of yeast and bacteria. This demonstrates the species-independent potential of FIND-IT for precision breeding.

FIND-IT will prove invaluable in new approaches to crop improvement by providing a direct means to select and implement: *i*) domestication traits into undomesticated or semi-domesticated plants such as perennials or wild crop relatives (e.g. quinoa^36^), *ii*) potentially valuable gene alleles retained in wild relatives that were replaced during domestication^37^, *iii*) gene alleles found in variomes (e.g. exome collections and pan genomes^7,38–40^) and *iv*) novel improved variants identified *in vitro*.

In conclusion, FIND-IT constitutes a powerful and ready-to-use, technological innovation-platform for trait development that allows fast and simple precision breeding of the high-yielding, nutrient-use-efficient and stress-tolerant crops that have become essential for food security in a world faced with a changing climate and increasing population.

## Acknowledgements

We thank the Carlsberg Foundation for funding support (grants CF14-0461 to B.L.M. and CF15-0236 to B.S.).

## Author Contributions

S.K., T.W., C.D. and H.C.T. contributed equally. S.K., T.W. and B.S. conceived the project, S.K., H.C.T., C.V., T.W., M.R., C.D. and P.R.P. were responsible for the mutagenesis and variant propagation, L.T.P., C.V., S.K., H.C.T., T.W., M.C., M.R., A.S., L.M., M.K.J. and J.A.C.S. generated plant libraries, Q.L., C.D., M.E.J., J.T.Ø., K.N., J.O.H. validated library mutation load, H.C.T., L.T.P., C.V., M.C., M.R. and T.W. performed the ddPCR experiments, C.D., J.A.C.S., I.B., S.B. and S.K. designed barley variant targets, C.D., S.B., J.A.C.S., P.R.P., E. M., J.H., H.C.T., M.T.S.N. and S.K. analysed barley variants, K.L., M.K., R.G.S. and N.S. prepared and screened the yeast library, R.T.F. and T.W. prepared and screened the bacterial library, Z.G., J.F., O.O. and B.L.M. assisted with experimental design, and C.D., M.E.J., H.C.T., M. R., B.L.M., G.B.F. and B.S. were responsible for the preparation of the manuscript, with assistance from the other authors.

## Competing interests

S.K., T.W., H.C.T, M.R., M.C., A.S., O.O. and B.S. are listed as inventors in a patent application WO2018/001884, and S.K., T.W., M.R., J.Ø and B.S. are listed as inventors in patent application PCT/EP2020/078321, both relating to the methodology and filed by the Carlsberg A/S. Furthermore, S.K., S.B, O.O., H.T., T.W. and J.H. are listed as inventors in a patent application WO2019129736A1, and C.D., P.R.P., S.K., O.O., T.W., M.C., H.T. and M.R. are listed as inventors in a patent application WO2019129739A1, both relating to traits and filed by the Carlsberg A/S. All authors are previous or current Carlsberg A/S employees. B.L.M. is a Distinguished Professor at the Carlsberg Research Laboratory and serves as a consultant for the laboratory. G.B.F. is a Senior Scientific Consultant and Distinguished Professor of Raw Materials at the Carlsberg Research Laboratory. The methodology presented in the manuscript is applied commercially by Traitomic, a business unit at Carlsberg A/S.

## Notes

### Competing Interest Statement

S.K., T.W., H.C.T, M.R., M.C., A.S., O.O. and B.S. are listed as inventors in a patent application WO2018/001884, and S.K., T.W., M.R., J.O and B.S. are listed as inventors in patent application PCT/EP2020/078321, both relating to the methodology and filed by the Carlsberg A/S. Furthermore, S.K., S.B, O.O., H.T., T.W. and J.H. are listed as inventors in a patent application WO2019129736A1, and C.D., P.R.P., S.K., O.O., T.W., M.C., H.T. and M.R. are listed as inventors in a patent application WO2019129739A1, both relating to traits and filed by the Carlsberg A/S. All authors are previous or current Carlsberg A/S employees. B.L.M. is a Distinguished Professor at the Carlsberg Research Laboratory and serves as a consultant for the laboratory. G.B.F. is a Senior Scientific Consultant and Distinguished Professor of Raw Materials at the Carlsberg Research Laboratory. The methodology presented in the manuscript is applied commercially by Traitomic, a business unit at Carlsberg A/S.

