## Supplementary material for "FIND-IT: Ultrafast mining of genome diversity": Methods

### Online Methods

**Method sensitivity validation.** The FIND-IT method is based on droplet digital polymerase chain reaction (ddPCR) technology, which is a highly sensitive technique for identifying rare variants in high background samples[1]. To validate the sensitivity of ddPCR for identification of single nucleotide variation in grain samples, 1,000 wild-type grains (cv. Barke) were milled with 0.5, 1, 2, 3 or 4 grains (cv. China) containing a known single nucleotide mutation in the *Lipoxygenase 1* gene (*Lox1*, HORVU.MOREX. r2.4HG0280010) followed by extraction of total DNA to be analysed using ddPCR (Supplementary Notes 1, Section 4).

The ddPCR assay reactants include competing fluorescently labelled TaqMan probes. The wild-type probe is labelled with the hexachlorofluorescein (HEX; green) fluorophore and the 'variant' probe is labelled with 6-carboxyfluorescein (FAM; blue) fluorophore. The TaqMan PCR probes and primers were designed using the BioRad design tool (www.bio-rad.com) with the two probes distinguishing between two different nucleotides at the same location in a targeted gene of interest. The reaction mixture for the ddPCR assay consisted of 2x QX200™ ddPCR™ Supermix for probes (no dUTP) (Bio-Rad, Hercules, CA, USA), 900nM/250nM of 20x target primers/probe (FAM for mutant, HEX for wild-type), 5 µL of diluted DNA and molecular grade water added up to total reaction volume of 20 µL. *Lox1* probes were designed to specifically distinguish between the cv. Barke and cv. China alleles (lox1-fwd: GCCATTGAGCAGTACG; lox1_rev: CATGGGGCGTCCTTG; lox1_G_hex: AGGCGTGGTGGAAGG; lox1_A_fam: AGGCGTGATGGAAGGA). ddPCR was performed as described below (FIND-IT phase 1).

**Generation of FIND-IT libraries.** A range of FIND-IT libraries were generated by induced, chemical mutagenesis of elite varieties of barley, wheat, rapeseed, oat, as well as yeast (*Saccharomyces cerevisiae*) and a bacterium (*Lactobacillus pasteurii*; DSM 23907). Here we describe in detail the preparation of the barley libraries and while the methods for preparing libraries from other species were similar, key differences are presented in Supplementary Notes 1, Section 1.

Barley mutagenesis was performed by soaking 2-3 kg grain in 0.3 or 1.67 mM sodium azide at pH 3.0 for 2 h[2]. After thoroughly washing with water, the grains were air dried in a fume hood for 16 h. The mutagenized M1 grains were sown at high density (600 plants/$m^2$) to reduce tillering and grown to maturity under field conditions. At maturity, M1 plants were harvested in pools of approximately 300 plants (0.5 $m^2$/pool). Grains from each pool were threshed, cleaned, and used to generate a low zygosity M1 library (containing M2 grains). Bulk harvested grains from M1 plants were used to generate a high-zygosity library the following season. The M2 grains were again sown at high density (600 plants/$m^2$) and field grown to maturity. At maturity, M2 plants were harvested in pools of approximately 300 plants (0.5 $m^2$/pool). Grains from each pool were threshed and cleaned individually, avoiding carryover between pools, and used to generate an M2 library (containing M3 grains). We initially built M1 libraries for developing the FIND-IT method (containing M2 grains). Working with M1 plants significantly speeds up library construction, and a fully screen-able library can be available within one growing cycle. All cells are susceptible to mutagenesis in the seed embryo, but only a few cells in the embryo's shoot meristem will contribute to seed heredity. These "germline cells" are independently mutagenized, giving rise to a chimerical plant, descending into several independent mutant lines[3]. Therefore, when screening M2 grains, we harvested in pools of approximately 300 plants. We kept on harvesting in pools of 300 plants when building M2 and M3 libraries for practical reasons. Only after DNA extraction, we combined samples to represent ca. 1,200 plants per pool in the M2 and M3 libraries.

Grains from each pool of up to 300 plants (200-600 g grain) were split using a Perten Sequential Divider SPD4200 (Perkin Elmer, Waltham, MA USA) to obtain a representative 25% sample of grains from the pool for DNA extraction. The representative 25% sample was milled (Retsch GM200, Haan, Germany) and DNA was extracted from 50 g of the flour by LGC Genomics (LGC Genomics GmbH, Berlin, Germany). DNA concentrations were adjusted to 150 ng/µl for each sample extract and distributed in 96-well library plates that each represented approximately 25,000-30,000 individual mutant plants. All M2 and M3 libraries were further pooled after DNA extraction to represent 1,200 plants per sample and approximately 100,000 individual mutant plants per 96-well plate. The remaining 75% sample of each grain pool was stored at 4 °C until potentially identified in a DNA library screen for further screening. Currently, 20 plates in our library, representing a range of barley cultivars, generations and mutagenesis treatments, are available for screening (Supplementary Notes 1, Section 1).

**Screening of FIND-IT barley libraries for specific nucleotide substitutions.** The FIND-IT screening pipeline is a three-phase process (Fig. 1b) that includes a Phase 1, the initial library screening where the number of candidates is narrowed down from the 100,000s to 300 candidates, followed by a Phase 2, the sub-pool screen where the number of candidates is further narrowed down from 300 to approximately 10 candidate grains, and a Phase 3, where the presence of the targeted nucleotide polymorphism is confirmed in a single individual.

In Phase 1 of the FIND-IT procedure, 100,000s of plants were screened in 96-well library plates. Each well contained DNA extracted from 25% of a grain pool representing approximately 300 individual plants. Dependent on the library used (M2 vs M3 grains), it was further possible to pool four DNA library plates, thus screening DNA representing up to 1,200 individual plants in a single well. Library plates were screened until a certain well showed an increased abundance of the nucleotide of interest. This pool was then advanced to Phase 2 in the screening process (Fig. 1b).

For Phase 1 screening, a master mixture of the TaqMan probe solution, distinguishing between specific nucleotides, and the ddPCR reagents was prepared and mixed with diluted DNA from a library DNA plate in a 96-well PCR plate. The reaction mixture was then prepared on a QX200™ AutoDG Droplet Digital PCR system (BioRad, Hercules, CA, USA) where each 20 µL of reaction mixture is separated and encapsulated into approximately 20,000 droplets using an oil film. Each droplet encases, on average, one genome equivalent of DNA and droplets are dispensed into 96-well plates for ddPCR analysis. The PCR plate containing the droplets was heat-sealed at 180°C for 5 sec with pierceable foil using a PX1 PCR plate sealer (Bio-Rad, Hercules, CA, USA) followed by PCR amplification (Uno96, VWR, Radnor, PA, USA) as follows: enzyme activation at 95°C/10 min followed by 40 cycles of denaturation at 94°C/30 s followed by annealing/extension at 55°C/1 min ending with enzyme deactivation at 98°C/10 min, all steps with a ramp of 2°C/s. During the PCR reaction, the fluorophore is displaced from the oligonucleotide and quencher and its fluorescence is unmasked. The droplet PCR plate was placed in a QX200™ droplet reader (Bio-Rad, Hercules, CA, USA) to count positive and negative droplets for each fluorophore and data was subsequently analysed using Bio-Rad QuantaSoft™ software (Bio-Rad, Hercules, CA, USA), based on the amplitude and concentration of fluorophore signal for each probe.

In Phase 2, about 1,000 grains were randomly selected from the stored 75% sample (approximately 5,000 remaining grains) of the positively identified DNA pool to further identify and isolate the variant grain of interest. Flour from each endosperm was sampled non-destructively by drilling a 1 mm hole approximately 2-3 mm into the endosperm (Marathon-3 Champion, Saeyang Microtech, Daegu, Korea) and collecting the flour emerging from that hole on weighing paper (about 1.0-1.5 mg per grain; [4]). The flour from 1,000 grains, in pools of ~10 grains, was distributed into a single deep 96-well plate and DNA was extracted (NucleoSpin Plant II, Mini kit, Macherey Nagel, Düren, Germany). An identical plate containing the small sub-pools of ~10 drilled grains was stored at 4°C until further analysis. The ddPCR reaction was repeated, as described above, using the same TaqMan assay as in the Phase 1 library screen. From the 'flour' plate, positives for the nucleotide substitution of interest were detected with a fractional abundance of 5-10%, depending on zygosity. Thus, the Phase 2 screen narrowed down the potential candidates to ~10 grains (Fig. 1b).

In Phase 3, all ~10 grains from the corresponding positive well were germinated on filter paper on a petri dish at 15°C. Germinated grains were transferred to soil after 4 days and DNA was extracted (DNeasy, Plant Mini Kit, Qiagen, Hilden, Germany) from the first emerging leaf of individual plants after 5-7 days. DNA from the ~10 plants was then analysed by repeating the initial ddPCR procedure, using the same TaqMan assay as in the original Phase 1 library screen. This Phase 3 screen identified one variant plant that contained the nucleotide substitution of interest (Fig. 1b). The mutant carrying the nucleotide substitution of interest was transferred to a large pot and DNA was re-extracted from a random leaf for verification of the targeted nucleotide substitution, using both the original ddPCR procedure and nucleotide sequencing.

With established crop libraries, the FIND-IT pipeline can isolate a given variant within a few days or weeks dependent on the presence of a given variant in the library being screened (Supplementary Notes 1, Section 5).

**Library validation - Sequencing of target gene fragments and Genotyping-By-Sequencing/ddRAD sequencing for determination of mutation load.** Mutation load in the barley libraries was measured at two levels. First, 464-497 bp fragments of four selected genes from a barley mutant library were sequenced to identify polymorphisms, and then re-sequenced to confirm the recorded differences. Here, a barley mutant population of cv. Quench was prepared as described above with the conditions shown in Supplementary Notes 1, Section 1. The mutant population was field grown and 12,000 single spikes of the M2 generation were harvested. A single grain (M3) per spike was isolated for further genomic DNA extraction and single read Sanger sequencing of target gene sequences, using Eurofins Genomics services (Eurofins Genomics Germany GmbH, Ebersberg, Germany). The chosen sequencing targets were single fragments of the genes HORVU.MOREX.r2.7HG0553990 (*Amino Acid Permease3*, *APP3*) and HORVU.MOREX.r2.6HG0462530 (*No Apical Meristem1*, *NAM1*) as well as two individual fragments of HORVU.MOREX.r2.4HG0278150 (*Iron-Regulated Transporter 1*, *IRT1*) using the amplification primers listed below. In total, *APP3*, *NAM1* and two *IRT1* fragments were sequenced in 6,144, 6,048, 5,952 and 4,704 individual mutant grains, respectively. Here, 5,763 (*APP3*), 5,551 (*NAM1*), 5,446 (*IRT1*_Ex1) and 3,731 (*IRT1*_Ex2) sequences were of appropriate quality for detailed analysis (Supplementary Notes 1, Section 3).

A certain level of heterozygosity can be expected in genomic DNA derived from M3 generation barley seeds. Thus, for the sequence analysis of each fragment, the sangerseqR package[5] was used to capture heterozygous bases with a “makeBaseCalls” ratio set at 0.8. Consensus sequences from “Primary Basecalls” and “Secondary Basecalls” were built for each fragment at the target region using Biostrings[6], where potential heterozygous mutation bases were coded according to IUPAC codes. Consensus sequences shorter than median length were excluded. Multiple sequence alignment was performed using the MAFFT v.7 software[7]. Alignments were read with seqinr[8] and converted into a matrix from which 500 bp of sequence was extracted (300 bp from upstream and 200 bp from downstream of the mid base). Positions with non-unique bases were reported as mutation sites for further confirmation by manually checking chromatograms. Manually confirmed mutations are listed in Supplementary Notes 1, Section 3.

In each confirmed case, four additional grains from the respective candidate spike were collected and germinated on petri dishes. Genomic DNA from seedling leaves was extracted and the respective gene fragment was PCR-amplified using the REDExtract-N-AmpTM Plant PCR Kit (Sigma-Aldrich, St. Louis, MO, USA), according to the manufacturer’s instructions. PCRs were performed for 35 cycles (initial denaturation at 94°C/3 min followed by 35 cycles of 94°C/30 s, 59°C/30 s, and 72°C/60 s for extension, with a final extension step of 72°C/10 min), using the following primers: HvAAP3_Ex6_F1 (CGTGTACCTGGAAATGCAGG) and HvAAP3_Ex6_R1 (GGATTTCGGCCTTCCAAGTG), LU_HvNAM1_F2 (GTCTCATCGATCAGTTGGACG) HvNAM1_uhb7_F (CAACCCCGTTCAACTGGCT) and LU_HvNAM1_R2 HvNAM1_Ex3_R2 (GTGTCATTCGTTCAGG GATTCCTGCATGCAACTACTCAGTGCT), HvIRT1_Ex1_F (AATCGCAACCAAAAGATCGAGC) and HvIRT1_Ex1_R (GCAACACAAGATTACCTGAACG) or HvIRT1_Ex2_F (GTCTAGTTCCCGTGTTTCCATG) and HvIRT1_Ex2_R (AGACACACCCTCATCAACCATA). PCR products were purified using the NucleoSpin® Gel and PCR Clean-Up Kit (Macherey-Nagel GmbH & Co. AG, Düren, Germany) according to the manufacturer’s instructions for Sanger sequencing (Eurofins Genomics Germany GmbH, Ebersberg, Germany). Mutation rates for each fragment were estimated and are given in Supplementary Notes 1, Section 3.

**In a second approach,** mutation load was further evaluated at a sparse genome-wide level using Genotyping-By-Sequencing/ddRAD [(n)GBS/ddRAD]. This experiment was performed using 48 plants of RGT Planet, including 8 wild-type plants, 20 M3 mutant progeny plants derived from a 0.3 mM $NaN_3$ treatment and 20 M3 mutant progeny plants derived from a 1.67 mM $NaN_3$ treatment. The (n)GBS/ddRAD analysis, including gDNA extraction, library construction, sequencing and initial bioinformatic analysis were performed at LGC genomics (LGC Genomics GmbH, Germany). Briefly, genomic DNA was extracted from dried leaf material, the restriction enzymes PstI and ApekI were used for complexity reduction. Samples were sequenced on an Illumina sequencer (Illumina NextSeq 500/550 v2) and approximately 3 million single-end reads with a length of 75 bp per sample were obtained. Read pre-processing was performed using the Illumina bcl2fastq 2.20 software and demultiplexed, cleaned reads were aligned to the Morex_V2 reference genome using the BWA-MEM software version 0.7.12[9], followed by SNP calling using Freebayes v1.0.2-16[6]. To lower false positive rate, the following filters were applied: Removal of loci with coverage depth lower than 8 reads in any of the samples. Removal of samples which were outliers based on a Student's T-test ($P<0.01$) on genetic distance matrices[10] generated for all plants with similar $NaN_3$ treatment. Removal of loci which had any neighbouring loci within 50 bp. Disregarding loci whose alternate fraction lower than 25%. Disregarding loci which were detected in two or more samples. Mutation rates for each fragment were estimated and are given in Supplementary Notes 1, Section 3.

**Analysis of naturally occurring genome variation in domesticated and wild barley of the barley pan genome.** Data of RGT Planet presence/absence variations (PAVs) versus the Morex_V2 was extracted from the barley pan genome[11], and illustrated with Circos 0.69.8[12]. The SNP matrix from 200 domesticated and 100 wild barley lines published as part of the barley pan genome was retrieved from IPK (SNP_matrix_WGS_300_samples.vcf.gz, https://doi.ipk-gatersleben.de/DOI/c4d433dc-bf7c-4ad9-9368-69bb77837ca5/1b898e9a-11bf-4eb6-b165-e1fa2167f0ec/0 )[11]. Summary statistics for each sample was calculated by bcftools stats

(1.10.2+htslib-1.10.2-3)[13] with the command **bcftools stats -s - SNP_matrix_WGS_300_samples.vcf.gz**.

**Sequence validation of isolated barley variants.** A single grain from each identified and isolated variant was germinated and gDNA was extracted from the first emerging leaf (DNeasy, Plant Mini Kit, Qiagen, Hilden, Germany). PCRs were performed with the respective primers listed in Supplementary Table 2 for 35 cycles (initial denaturation at 94°C/3 min followed by 35 cycles of 94°C/30 s, 61°C/30 s, and 72°C/45 s for extension, with a final extension step of 72°C/10 min). Appropriate PCR products were purified using the NucleoSpin® Gel and PCR Clean-Up Kit (Macherey-Nagel GmbH & Co. AG, Düren, Germany) according to the manufacturer's instructions for Sanger sequencing (Eurofins Genomics Germany GmbH, Ebersberg, Germany).

**Background heritage validation by 50K SNP chip genotyping.** Isolated FIND-IT variants and their corresponding wild-types were germinated, leaf material was sampled, freeze-dried and sent for 50K SNP genotyping at TraitGenetics (SGS - TraitGenetics GmbH, Germany). The genetic distance between individuals was calculated as the average of their per-locus distances using R package stringdist[14]. Principal Coordinate Analysis (PCoA) was done with R base function cmdscale based on this genetic distance matrix. The first two PCs were illustrated by ggplot2.

**Plant material and growth conditions.** Barley cultivars and isolated FIND-IT variants were grown in a greenhouse at 18°C under 16 h/8 h light/dark cycles. At maturity, grains were harvested and further propagated, either in dozens or in hundreds in the greenhouse or in field row plots, respectively, followed by field trial plots for agronomic evaluation (7.5 $m^2$ plots) in Denmark. Grain from field plots was harvested and threshed using a Wintersteiger Elite plot combiner (Wintersteiger AG, Germany), and grains were sorted by size (threshold 2.5 mm) using a Pfeuffer SLN3 sample cleaner (Pfeuffer GmbH, Germany).

**Yield performance evaluation.** Thousand grain weight (TGW), grain width and grain length of mature dry grains (sample size approximately 500 grains per measurement) from field grown plants were determined using the digital seed analyzer MARVIN (GTA Sensorik GmbH, Neubrandenburg, Germany).

**Statistical analysis.** All field grown material used for agronomical, enzymatic or gene expression characterization were grown in field trials in either New Zealand (September-February) or Denmark (April-August) and in distinct replicates of n=2 to n=5. Microsoft Excel data analysis package was used for comparison of two groups by Paired Student's two-tailed T-test to test if the hypothesized mean difference is zero. Values of p were considered significant different if ***p ≤ 0,001, ** p ≤ 0,01 or * p ≤ 0,05. All statistical parameters and outputs are presented in Supplementary Table 3.

**Starch amylose Lugol staining.** Grains were embedded in a clay block and grinded with sandpaper. Remaining half-grains were dipped in diluted Lugol solution (0.1% $I_2$ and 0.2% KI) for 20 minutes, tapped dry with paper and exposed to air for approximately 10 min before photographing.

**Starch amylose assay.** Three replicates each of 50 grains of the variant *gbss1*$^{W126*}$ and of Planet control were ground to flour in an IKA Tube-mill 100 (IKA®-Werke GmbH & Co. KG, Germany) at 20,000 rpm and the amylose:starch ratio was determined using a miniaturized version of the K-AMYL kit (Megazyme, Wicklow, Ireland).

**Green revolution varieties genotyping - *Sdw1* sequencing.** Barley cvs. Paustian and Planet were grown in a greenhouse and genomic DNA was extracted from green leaf material using the NucleoSpin® 96 Plant II kit (Macherey-Nagel GmbH & Co. AG, Düren, Germany) according to the manufacturer's instructions. A fragment of exon 1, harboring the 7-bp deletion of the *sdw1.d* variant allele, was PCR-amplified using the TaKaRa LA Taq polymerase (0.25 µl) with 2 x GC buffer I (12.5 µl) (TaKaRa Bio Europe), mixed with MilliQ water (4.25 µl), 2.5mM dNTP (4 µl), 10 mM forward primer (1µl), reverse primer (1µl) and genomic DNA (10ng/µl, 2 µl). PCR with primers CD348_sdw1d_fw (5'-

GGTGCTCCAGACCGCTCAGC-3') and CD347_sdw1d_rc (5'- CCTCCGGAGGTCGTACACC-3') was performed for 35 cycles (initial denaturation at 94°C/3 min followed by 35 cycles of 94°C/30 s, 60°C/30 s, and 72°C/45 s for extension, with a final extension step of 72°C/10 min). PCR products were purified using the NucleoSpin Gel and PCR Clean-Up kit (Macherey-Nagel GmbH & Co. AG, Düren, Germany) according to the manufacturer's instructions. Purified PCR products were sequenced by Eurofins (Eurofins Genomics Germany GmbH, Ebersberg, Germany).

**Broken kernel test.** Field-propagated mature grain of *CslF6* variant plants $cslF6^{G748D}$, $cslF6^{G847E}$ and $cslF6^{T709I}$ as well as respective reference lines cvs. Paustian and Quench was harvested and threshed using a trial Wintersteiger Classic (Wintersteiger AG, Austria). Further cleaning of grains was performed on a Pfeuffer 20 sample Cleaner, model SLN4 (Pfeuffer GmbH, Germany) using a 2.5 mm screen. Broken grains were counted in four randomly selected 50 g samples of each line and respective wild-type, and the amount of broken grains for each line was calculated on weight basis.

**α-Amylase gene expression analysis.** Plant material and experimental setup: Triplicates of 100 grains of *Amy1_1-var* and wild-type cv. Planet were germinated on filter paper in a petri dish (ø 85mm, Frisenette, Knebel, Denmark) in 4 ml water at 16°C. At 48 or 72 h germination, the grains were snap frozen in liquid nitrogen and freeze-dried for 48 h before being finely ground (MM300, Retsch, Haan, Germany).

Primer and probe design: Primers and probes for *Amy1_1*, *Amy1_2* and *Amy2* were designed based on Morex_V2[15]. Primers and fluorescent probe for *Amy1_1* and *Amy2* were designed to specifically amplify and bind to the *Amy1_1* (*a,b,c,d*) and *Amy2* (*_1,_2,_3*) gene clusters, respectively. *Amy1_2* is present as a single gene. The amplicon location was specifically targeted to the 3'UTR or all targets and the probe was labelled with 6-carboxyfluorescein (FAM) for fluorescence detection of transcript.

Amy1_1: Fwd_GACTGGGGCCTGAAG, rev_ GTGCCGGGTCCTGAC, FAM_ AGATCGATCGCCTGGTGTC

Amy1_2: Fwd_ AGATCGATCGTCTGGTG, rev_ TCCATGATCTGCAGCTTG, FAM_ TCAATCAGGACCCGACAGG

Amy2: Fwd_ CGAGCTCAAGGAGTGG, rev_ CGTCGATGTACACCTTG, FAM_ AAGAGCGACCTCGGCTTC

RNA extraction and cDNA synthesis: Total RNA was extracted according to Betts et al. (2017)[16], from the equivalent of 2 grains (~100 mg freeze-dried and milled grain material) by a modified Spectrum Plant Total RNA Kit (Sigma-Aldrich) protocol. DNA was removed using DNase I (Sigma) and ZYMO Clean & Concentrator Kit (ZymoResearch) was used to clean and concentrate the RNA sample for further downstream applications. RNA quality was validated by gel electrophoresis (1% agarose) and quantified by Nanodrop (ThermoFischer, Waltham, MA, USA). cDNA was synthesized from 200 ng total RNA using BioRad iScript Select (oligodT) kit (Bio-Rad, Hercules, CA, USA) according to manufacturer's protocol and diluted 10x with milliQ water before ddPCR analysis.

ddPCR analysis: *Amy1_1*, *Amy1_2* and *Amy2* transcripts were quantified using the Biorad ddPCR system (Biorad, Hercules, CA, USA): A master mix of 2x QX200TM ddPCR™ Supermix for probes (no dUTP) (Bio-Rad, Hercules, CA, USA) and 900nM/250nM of 20x target primers and probe, 5 µL of diluted cDNA and molecular grade water was added up to total reaction volume of 20 µL and placed into the QX200™ Droplet Generator (BioRad, Hercules, CA, USA). Following droplet generation, the plate was heat sealed at 180°C for 6 s with pierceable foil using a PX1 PCR plate sealer (BioRad, Hercules, CA, USA). Subsequent PCR amplification (Uno96, VWR, Radnor, PA, USA) was as follows: enzyme activation at 95°C/10 min followed by 40 cycles of denaturation at 94°C/30 s followed by annealing/extension at 55°C/1 min ending with enzyme deactivation at 98°C/10 min, all steps with a ramp of 2°C/s. The PCR plate was then placed in QX200™ Droplet Reader (BioRad, Hercules, CA, USA) and positive and negative droplets were recorded and subsequently analysed using the Quantasoft™ software (BioRad, Hercules, Ca, USA) to evaluate amplitude and concentration (copies/µl cDNA) of the different transcripts.

**Analysis of grain (1,3;1,4)-β-glucan content.** Barley grains were milled on a Retch cyclone mill (Haan, Germany) and replicates of 20 mg flour were heated for 2 h at 100 °C.  After cooling to room temperature, 500 µl 50% (v/v) aqueous methanol was added, and the sample was shaken gently for 1 h. Following centrifugation at 16,000 g for 10 min, the supernatant was discarded, and the flour dried overnight. Sodium phosphate buffer (400 µl, 20 mM, pH 6.5) was added, together with 1 U/ml lichenase (Megazyme, Wicklow, Ireland) per 10 mg flour. After incubation at 50 °C for 2.5 h the sample was centrifuged at 16,000 g for 10 min and the supernatant filtered through 0.45 µm filters. Released oligosaccharides were quantified by HPAEC-PAD using a Dionex ICS 5000+ DC system (Thermo Fischer, Waltham MA USA) equipped with a 4 µm SA-10 column at 0.4 ml/min and a column temperature 40°C, with an isocratic 100 mM NaOH eluent for 15 min. Standards were produced by 1 U/ml lichenase (Megazyme, Wicklow, Ireland) digestion of known quantities of medium viscosity (1,3;1,4)-β-glucan (Megazyme, Wicklow, Ireland) in 50 mM MES buffer, pH 6.5. Visualization of (1,3;1,4)-β-glucan in longitudinal sections of *CslF6* knockout mutants and wild-type grains was carried out according to Aastrup et al. (1981)[17] using 0.05% Calcofluor for staining and 0.05% Fast Green for counter staining.

**Homology modelling of protein structures.** Homology models were built using the automated SWISS-MODEL homology modelling pipeline[18]. Briefly, this workflow searches for homology model templates using BLAST[19] (Camacho et al., 2009) and HHblits[20] that are subsequently analysed and ranked using QMQE (monomer) or QSQE (oligomer)[21]. SWISS-MODEL uses the modelling engine ProMod3 to extract structural information from the selected template and to perform the modelling. Non-conserved side chains are modelled using a back-bone dependent rotamer library and optimized using the Treepack algorithm[22]. Finally, energy minimization is performed by ProMod3 using the OpenMM library[23] and CHARM22/CAMP force field parameters[24].

The homology model of the *Hordeum vulgare* Cellulose Synthase–like F6 enzyme, encoded by *CslF6* (HORVU.MOREX.r2.7HG0579230), was based on the poplar cellulose synthase isoform 8 (PDBID: 6wlb, 43.18% identity). All models were visualized using VMD[25] and final figures arranged using Adobe Illustrator™ and CorelDRAW.

#### Yeast library generation.

A haploid *Saccharomyces cerevisiae* strain was treated with ethyl methanesulfonate (EMS) using a standard protocol [26]. In brief, the strain was first grown in standard yeast extract/peptone/dextrose (YPD) medium to saturation. The cells were spun down, washed once with sterile water and one more time with 0.1 M sodium phosphate buffer (pH 7). Finally, a cell suspension was generated in the same buffer with a cell titer of approx. $2 \times 10^8$ cells/ml. For each mutagenesis reaction 1 ml of this cell suspension was pipetted into a 2 ml safe-lock reaction tube and 30 µl EMS were added to the cells. The reactions were put on an Eppendorf Thermomixer comfort (1.5 ml) and incubated at 1000 rpm and 30 °C for 75 minutes. These conditions resulted in a survival rate of approximately 30 %. The mutagenesis was stopped by washing the cells three times with 1 ml of freshly prepared sterile sodium thiosulfate solution (5 %) and one time with sterile water. Cells were finally revived in 1 ml YPD for 1 hour at beforementioned conditions. Cells were either processed right away or were stored at 4 °C until use.

Literature suggested that growing yeast cells under selective pressure immediately after mutagenesis would select for cells with mutator-like phenotype accumulating significantly more mutations per single cell than under non-selective conditions [27,28]. Hence, one could in theory reduce the initial size of our mutant library significantly due to the cells' increased mutation density. We plated therefore the mutagenized yeast cells on solid synthetic dextrose (SD) medium containing 2 µg/ml of the herbicide metsulfuron methyl and incubated the plates at 30 °C until resistant colonies appeared. The cells of approx. 50,000 colonies were collected from the plates, washed with sterile phosphate buffer (see above) and used for another round of EMS mutagenesis and selection as described. The described mutagenesis/selection cycle was done four times overall to increase the mutation density in the yeast population even further.

After the final cycle, the cell titer of the yeast suspension was determined, and cells were spread onto solid YPD medium to create plates with approx. 1,200-1,500 colonies/plate. After three days at

30 °C, cells from 96 individual plates were washed off with phosphate buffer and collected separately to form 96 library pools. These cells were subsequently used for DNA isolation using the PureLink® Pro 96 genomic DNA Kit (ThermoFischer, Waltham, MA, USA) with an EveryPrep™ Universal Vacuum Manifold (ThermoFischer, Waltham, MA, USA), and on the other hand for generating glycerol stocks for long term storage. The final "total random library" created with the described procedure should contain approx. 120,000-150,000 mutant clones overall.

**Yeast library screening.** We used the FIND-IT approach to screen for mutations in the ferulic acid decarboxylase (*FDC1*) gene. This enzyme is responsible for converting aromatic carboxylic acids such as ferulic acid derived from malt to their corresponding vinyl derivatives, which often impart undesirable clove-like off-flavours in beer[29]. Two Trp residues in the enzyme (Trp159 and Trp171) were targeted for the formation of premature stop codons, using the S288C *ScFDC1* sequence as a reference (GenBank number NCBI: NM_001180847). The mutations were confirmed by sequence analysis. The assay of mutant lines was based on the inactivation of the *FDC1* gene, which resulted in increased sensitivity of the yeast to growth inhibition by cinnamic acid[30]; this effect was clearly visible in the mutant lines (Supplementary Notes 3, Section 3).

**Bacterial library generation and screening.** A *Lactobacillus pasteurii* (DSM 23907) library of about 960,000 genetic variants was generated by treating cells with 1.4% EMS for 30 min [31]. Post mutagenesis and recovery cells were counted using a NovoCyte flow cytometer (Acea Biosciences) and about 1,000 mutagenized bacterial cells were dispensed into each well of 10, 96-well plates. The library was screened for variants of the of the Galactose-1-phosphate uridylyltransferase gene (galT) which mediates the second step of the Leloir pathway in which D-galactose is metabolised [31,32]. Mutations in this gene should show no phenotype if the bacteria are grown on glucose. After screening about 10% of the library, two mutants with stop codons in this gene were identified. The mutants were Lac-GalMut_W89* and Lac-GalMut_Q148* (Supplementary Notes 2, Section 2).

**Nucleotide and amino acid numbering.** Throughout the manuscript, nucleotide and amino acid numbering are based on publicly available gene and protein sequences. The exact amino acid positions described are dependent on the template haplotype used for generating the probe. Where possible, we have aligned the numbering with the Morex_V2 genome sequence, but we also provide full-length Morex_V2 probe sequences that will guide readers to the correct nucleotide positions.
