## Supplemental notes 1 for "FIND-IT: Ultrafast mining of genome diversity"

### Supplementary Notes 1

#### Supplementary Notes 1, Section 1. Currently available FIND-IT libraries.

**Table Supplementary Notes 1.1a. Total available libraries in which variants have been identified, including mutagen treatment.**

| Type | Cultivar | Grains | Mutagen | Concentration of mutagen | Presoak 4°C, h | Treatment | Individuals for screening* |
|---|---|---|---|---|---|---|---|
| Spring barley | Planet, Paustian, Quench | M2, M3, M4 | $NaN_3$ | 0.3-1.67 mM | 16 | 2h, 20°C | ~500,000 |
| Spring wheat | Alondra, Harenda | M3 | EMS | 0.60% | 0 | 12h, 20°C | 60,000 |
| Winter wheat | Extase, Siskin | M3 | EMS | 0.60% | 0 | 12h, 20°C | 30,000 |
| Spring Rapeseed | Mosaik | M3 | EMS | 0.3% | 0 | 16h, 20°C | 25,000 |
| Winter Rapeseed | Epure | M2 | EMS | 0.25% | 0 | 16h, 20°C | 35,000 |
| Oats | Galant | M3 | EMS | 0.8% | 0 | 16h, 20°C | 40,000 |
| *Saccharomyces cerevisiae* | - | - | EMS | 3.0% | - | 75 min, 30°C | 120,000-150,000 |
| *Lactobacillus pasteurii* | - | - | EMS | 1.40% | - | 30 min, 30°C | 960,000 |

*the number of total individual variants is based on an estimation that each harvested pool contains grains from approximately 300 densely sown plants for barley and ~100 densely sown plants for wheat

**Table Supplementary Notes 1.1b. Spring barley cultivar libraries in which variants have been identified, including mutagen treatment.**

| Cultivar | Grains | Mutagen* | Total individuals available for screening** |
|---|---|---|---|
| Paustian | M2 | 0.3 mM $NaN_3$ | ~500,000 |
| Quench | M4 | 0.3 mM $NaN_3$ | |
| Planet | M2 | 0.3 mM $NaN_3$ | |
| Planet | M3 | 0.3 mM $NaN_3$ | |
| Planet | M3 | 1.67 mM $NaN_3$ | |

*barley grains were pre-soaked in water for 16 h at 4°C, drained and subjected to mutagenesis for 2 h at 20°C before air drying for 48 h in fume hood.
**number of individual mutants is based on an estimation that each harvested pool contains grains from approximately 300 densely sown plants

**Supplementary Notes 1, Section 2. Analysis of naturally occurring genome variation of domesticated and wild barley in the barley pan genome.**

The barley pan genome with 20 diverse genome assemblies identified approximately 1.5 million presence/absence variations (PAVs) ranging in size from 50 to 999,568 bp[1]. Comparison of the barley Morex_V2 and RGT Planet genomes alone identified 71,553 deletions and 56,063 insertions in the range from 50 - 987,841 bp (Fig. Supplementary Notes 1.1a; Online Methods). Whole genome sequencing data of 200 domesticated and 100 wild barley lines published with the barley pan genome[1] revealed that 299 out of 300 lines had between 5 million and 25 million naturally occurring nucleotide substitutions when compared to the Morex_V2 genome (Fig. Supplementary Notes 1.1b,c and Online Methods).

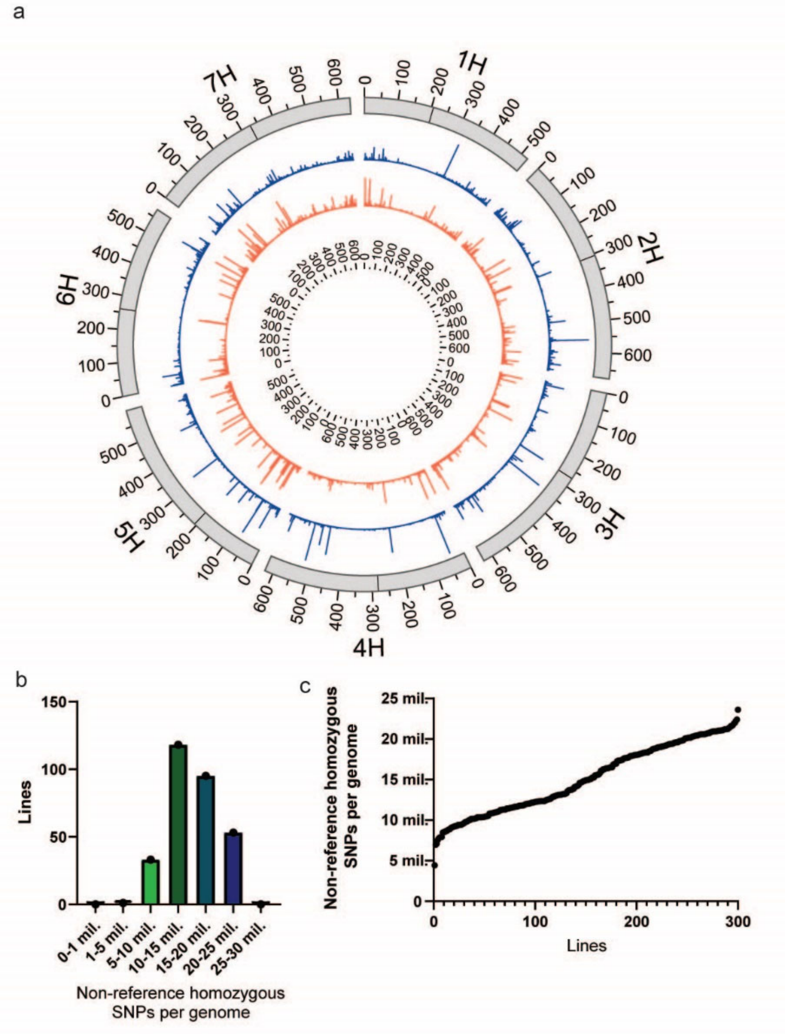

**Fig. Supplementary Notes 1.2. Naturally occurring genome variation of domesticated and wild barley in the pan genome. a**, PAVs (presence/absence variations) in the RGT Planet genome when compared to the Morex_V2 reference genome. Outmost track visualizes Morex_V2 chromosomes in Mb with approximate centromere highlighted as black bar. Inner tracks depict deletions (blue track) and insertion (orange track) found in RGT Planet with sizes indicated as bar height (ranging from 50bp to nearly 1Mb). **b**, Nucleotide substitution summary statistics of whole genome sequencing of 200 domesticated and 100 wild barley lines mapped to Morex_V2 reference genome. Non-reference homozygous SNPs in bins of 5 million SNPs per genome are shown. **c**, Non-reference homozygous SNPs in 200 domesticated and 100 wild barley lines.

#### Supplementary Notes 1, Section 3. Determination of mutation load of variant libraries.

**Table Supplementary Notes1.3a. Determination of mutation load of variant libraries by Sanger sequence analysis of gene fragments.**

| Barley gene | Fragment length (bp) | Mutants sequenced | Mutation type* | Sequence space (bp) | Mutation density (mutations /1Mb) | Estimated mutation rate |
|---|---|---|---|---|---|---|
| *APP3*<br>*(Amino Acid Permease3)*<br>HORVU.MOREX.r2.7HG053990<br>chr7H:98,808,452-98,810,950 | 464 | 5,763 | 820G>A, Hom<br>851C>T, Het<br>982C>T, Hom | 2,674,032 | 0.93 | 1.02E-6 |
| *NAM1*<br>*(No Apical Meristem1)*<br>HORVU.MOREX.r2.6HG0462530<br>chr6H:51,261,970-51,263,563 | 497 | 5,551 | 867C>T, Hom<br>976A>G, Hom<br>998G>A, Hom<br>1100C>T, Hom | 2,758,847 | 1.45 | |
| *IRT1*<br>*(Iron-Regulated Transporter 1)*<br>HORVU.MOREX.r2.4HG0278150<br>chr4H:6,593,933-6,596,483 | 480 | 5,446 | 193G>A, Hom<br>249G>A, Hom | 2,614,080 | 1.15 | |
| | 468 | 3,731 | 480C>T, Hom<br>795C>T, Het | 1,746,108 | 0.29 | |

*Zygosity status is denoted as Hom for homozygous and Het for heterozygous.

**Table Supplementary Notes1.3b Determination of mutation load of variant libraries by Genotyping-By-Sequencing (GBS).**

| Mutagen level and sample number* | Mutations | Mutation position | Mutation type | Mutation zygosity** | Mutation density (mutations /1Mb) | Estimated mutation rate*** |
|---|---|---|---|---|---|---|
| Low 6 | 2 | chr3H:506938316 | C>T | Het | 0.92 | 3.22E-07 |
| | | chr7H:534010302 | C>T | Het | | |
| Low 12 | 1 | chr2H:236578529 | G>A | Hom | 0.92 | |
| Low 17 | 1 | chr4H:457198053 | C>T | Hom | 0.92 | |
| Low 20 | 1 | chr7H:314434249 | G>A | Het | 0.46 | |
| High 3 | 1 | chr7H:141001621 | C>T | Het | 0.46 | 1.56E-6 |
| High 4 | 2 | chr1H:364657757 | C>T | Het | 0.92 | |
| | | chr7H:458672114 | C>T | Het | | |
| High 7 | 2 | chr1H:511484793 | G>A | Het | 0.92 | |
| | | chr5H:543755799 | C>T | Het | | |
| High 12 | 1 | chr1H:471596482 | C>T | Hom | 0.92 | |
| High 13 | 2 | chr3H:654773772 | G>A | Hom | 1.84 | |
| | | chr4H:609222511 | C>T | Hom | | |
| High 15 | 7 | chr1H:108247541 | G>A | Het | 5.06 | |
| | | chr1H:529698929 | G>A | Hom | | |
| | | chr3H:491484168 | G>T | Hom | | |
| | | chr3H:555808799 | G>A | Het | | |
| | | chr4H:130060307 | G>A | Hom | | |
| | | chr7H:37879999 | G>A | Hom | | |
| | | chr7H:450634289 | G>A | Het | | |
| High 17 | 1 | chr6H:518937333 | G>A | Hom | 0.92 | |
| High 18 | 2 | chr4H:544424142 | G>A | Het | 1.38 | |
| | | chr4H:560341634 | G>A | Hom | | |
| High 20 | 4 | chr2H:474460039 | C>T | Hom | 3.22 | |
| | | chr3H:556196067 | G>A | Het | | |
| | | chr3H:675147252 | C>A | Hom | | |
| | | chr4H:599974714 | G>A | Hom | | |

*Among 20 samples genotyped for each treatment, only samples with detected mutations are listed in the table.
**Zygosity status is denoted as Hom for homozygous and Het for heterozygous.
*** Mutation rate is based on a total sequence space of 21,757,380bp (1,087,869 bp per sample) from 20 samples with the same treatment.

**Supplementary Notes 1, Section 4. Sensitivity of ddPCR method for detection of rare events.**

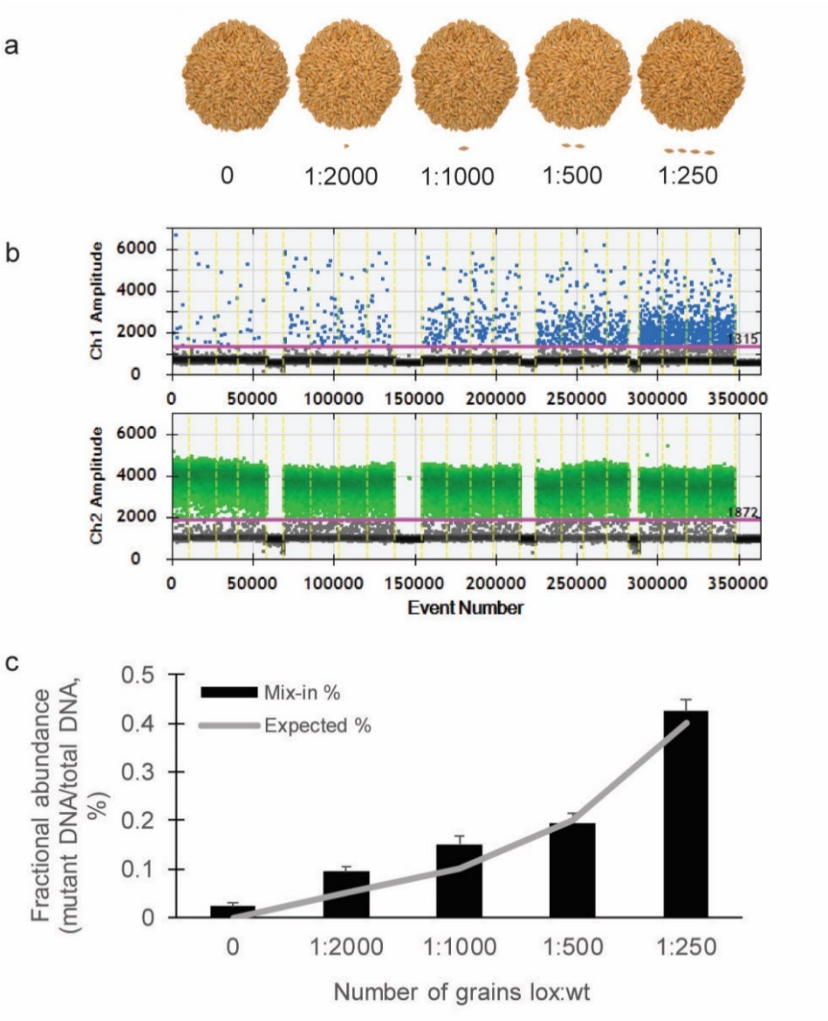

**Fig. Supplementary Notes 4.1. Validation of ddPCR sensitivity for detection of rare mutational events in grain pools. a**, 1,000 wild-type grains were milled with 0.5, 1, 2, 3 or 4 grains containing a known single nucleotide mutation in the lipoxygenase gene, *Lox1*, followed by total DNA extraction. **b**, DNA from each mix-in sample was subjected to ddPCR following standard procedure as described in Online Methods using hydrolysis probes specifically distinguishing between wild-type and variant allele. **c**, Fractional abundance of variant DNA in the total amount of DNA in each sample (black bars) compared to the expected abundance (gray line) was used to evaluate the sensitivity of the method.

**Supplementary Notes 1, Section 5. Estimated timeline for FIND-IT workflow following library generation.**

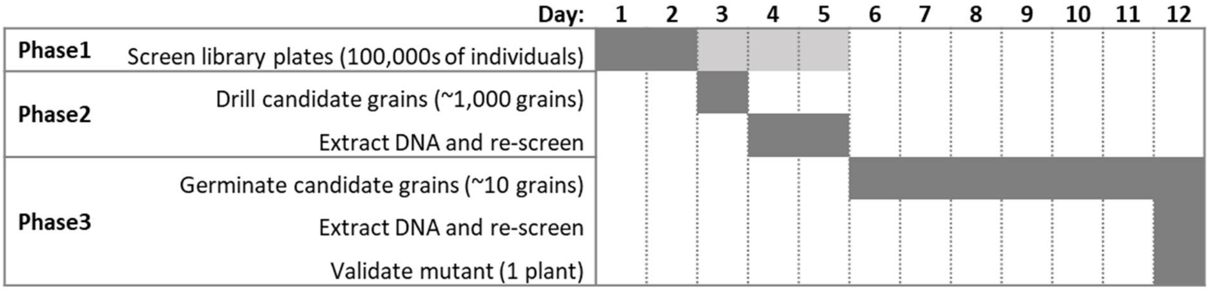
