## Supplemental notes 2 for "FIND-IT: Ultrafast mining of genome diversity"

### Supplementary Notes 2

#### Contents

Supplementary Notes 2, Section 1: Barley variants.................................................................................. 2

Supplementary Notes 2, Section 2: Beyond barley. ............................................................................. 55

**Supplementary Notes 2, Section 1: Barley variants. Fractional abundance values of variant DNA to total DNA following ddPCR in Phase 1 of the FINDIT screening process in each of 155 barley variants containing premature stop codons, nucleotide substitution leading to an amino acid substitution in gene of interest or nucleotide substitution in regulatory regions as presented in Supplementary Table 1.** Dots represent individual pools on a library plate, with the positive pool(s) identified for further downstream screening in FIND-IT phase 2 marked in gray open circles. The gene sequences and the targeted SNP are shown under the graphs.

1

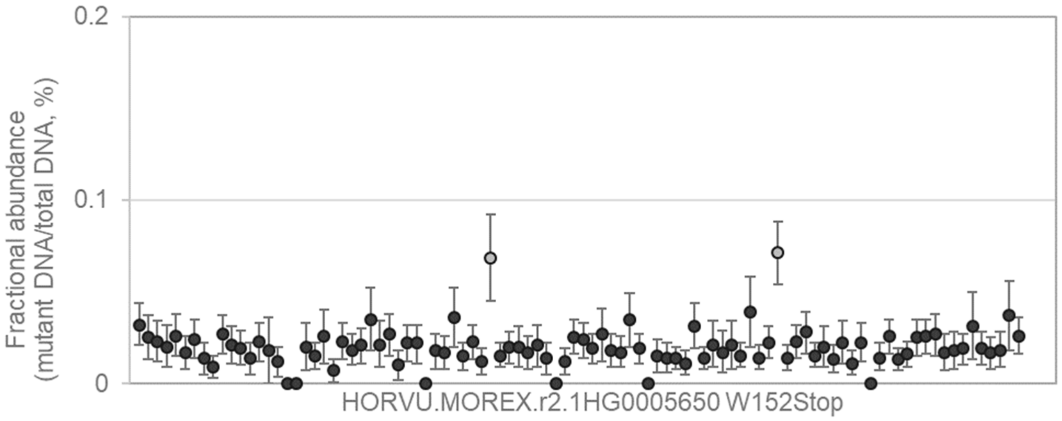


CCTTCGGTGAGCAGTCGACCAAGATGCGCCGGGTGCTCACCTCGCACATCGTCTCCCCGTCCCGCCA
CAAGTG[G/A]CTCCACGACAAGCGCGCCGAGGAGGCCGACAACATCACGTGGTACATGTACAACCTCA
CCGGCGAGGAGG

---

2

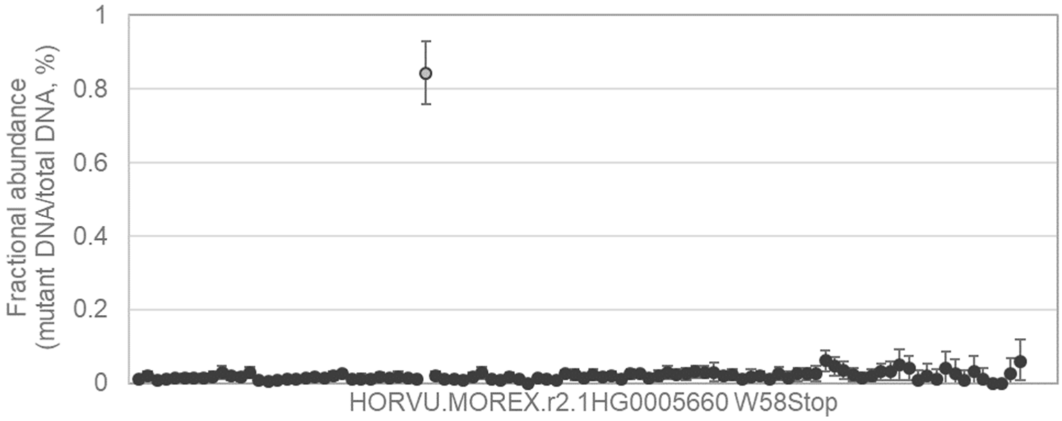


CGTGCGGCCCCACGATGGACATCGTCAGCTGCGGCGGCAAAGGCATCCTCTTGGCACCCTACGGCG
ACCACTG[G/A]CGGCAGATGCGTAAGGTTTGCGTCGTGGAGGTGCTCAGCGCAAGGCAGGTGCGGCG
TATCGAGTCCATCC

3

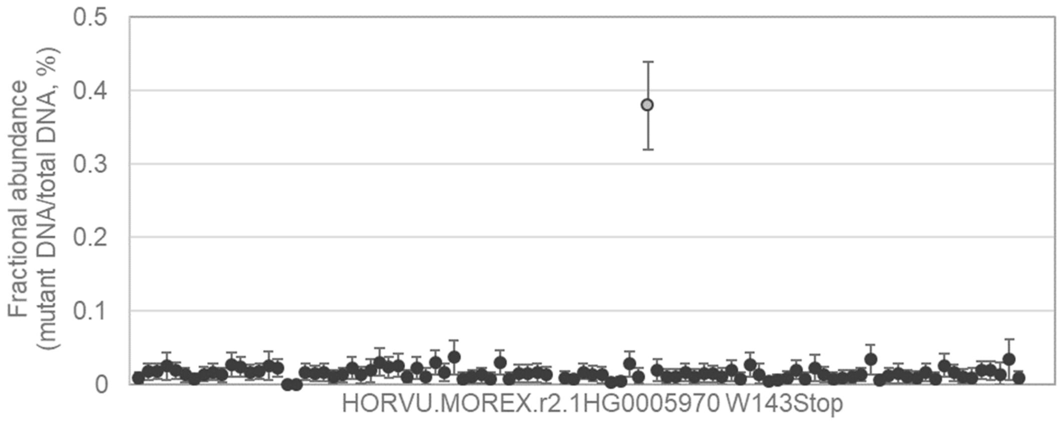


CGTGGCGGACAACGTGACGAGCTTCAGCGTCGACGCGGCGGCGGAGCTCGGGGTCCCCTGCGCGC
TCTTCTG[G/A]ACCGCCAGCGCCGCCGGTTACATGGGCTACTGCAACTTCCGGTTCCTCATGGACAAG
GGCATCGCTCCCC

4

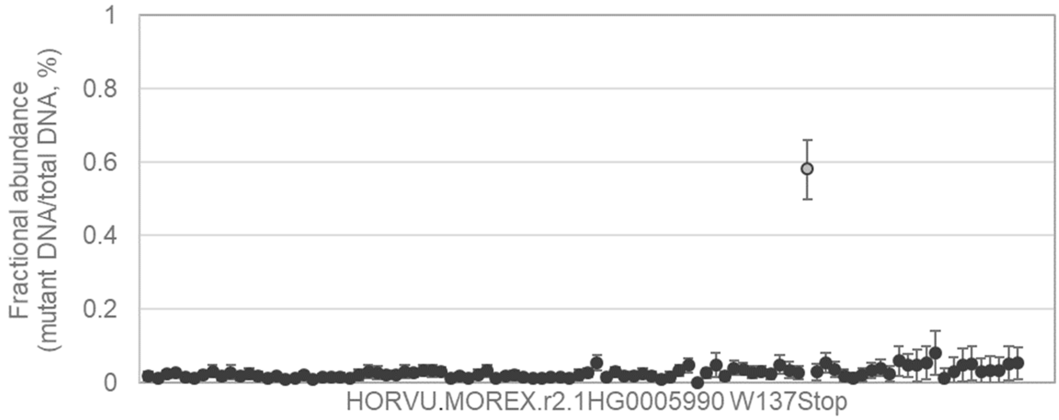


CCCCACAACCCCGCCACCGACATCATATTCTACGGCCCCAGTGACATCGCCTTCTGCCCCTACGGCG
ACCACTG[G/A]AGGCAGGTGAAGAAGATCGCCATGACGCACCTCCTCACCGCCAACAAGGTCCGCTC
CTACCGCCAGGCGC

5

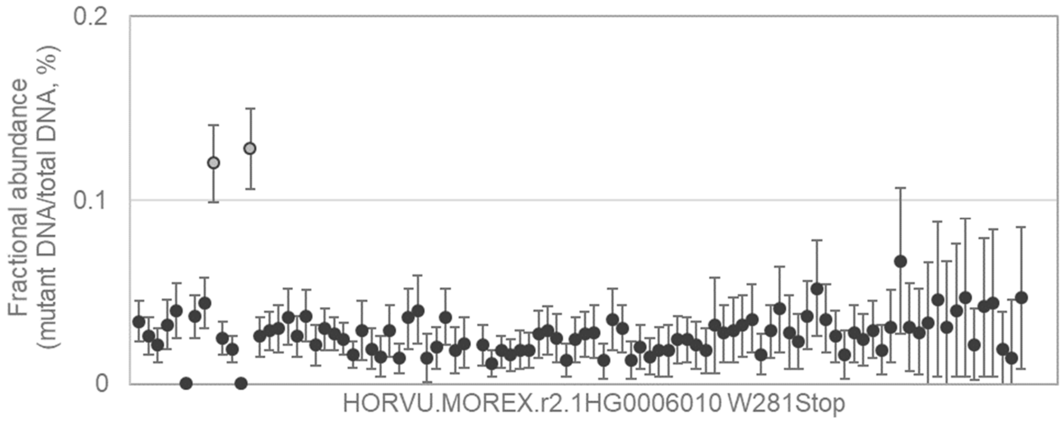


GCCCGCTCCACCTCCTCGCCCAAGGCCAAGGCCAAGGCCAAGTGCTGCTCCCCGAGATACCCGAGG
TGCTCTG[G/A]AAAGCAGACGGGTCCTGCCTCGAGTGGCTGGACGGAAGGGAGCCCGGGTCCGTGGT
GTACGTCAACTTTG

6

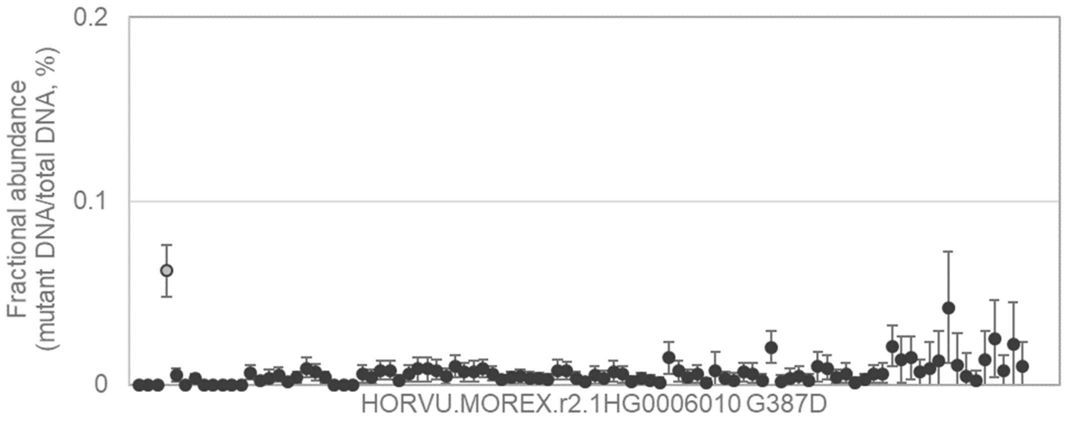


CTCCTGACGAGCTGGTGCGACCAGGAGGCGGTGCTGCAGCACCCGGCCCTCGGCGTCTTCCTCACG
CACT[G/A]CTGGAACTCCGCCTTGGTGGCCATCAGCGCCGGGGTGCCCATGCTGGGCTGGCCCTTCT
TCGCCGA

---

7

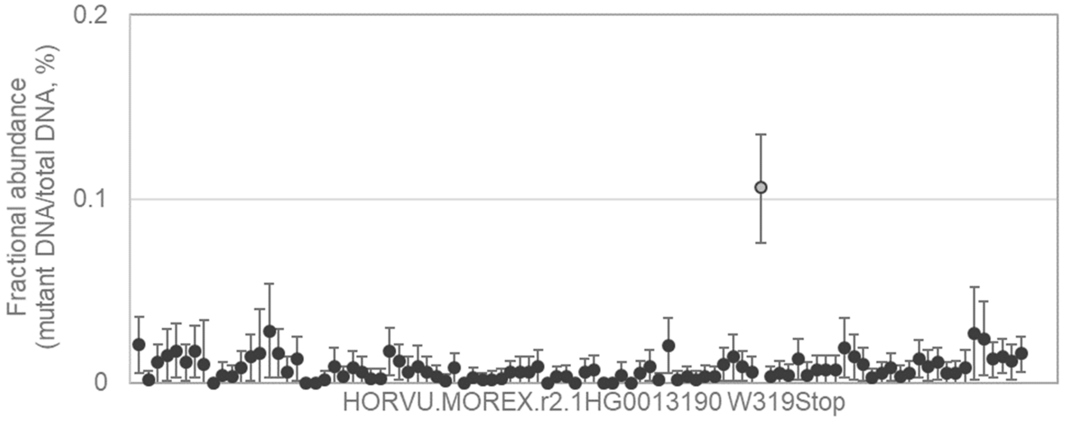


CTACACGAGAAGAAATTAAGTGCATGAATCCAAATTATACCGAGTTTAAATTCCCGCAAATCAAAGCTC
ACCCATG[G/A]CACAAGGTAGGAACCTTGTGTTAGCTGTTTTACTTGTCTATAAGGCTGCAGTATAACAT
TACTGTTTTTATCC

---

8

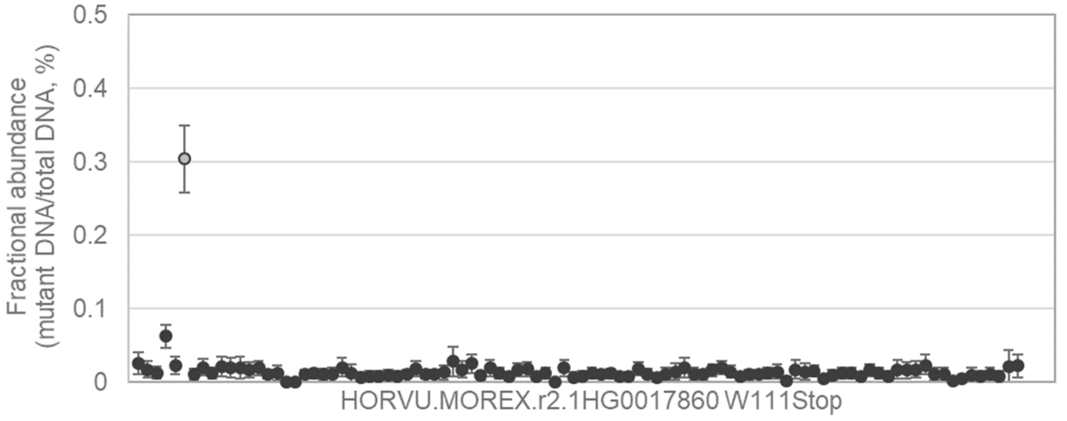


TACAGAACACAATAATGGGGATGAAGAAATTCTTGACACATACAATGCGCTATTACATACCGAGGCAG
CAGAATG[G/A]ACAGATACTAGAGAAGCTGAAACTGCTGAAGCAGACTCGTCACAAAATGCTTCAAGCA
GTTCTATAATCAGG

9

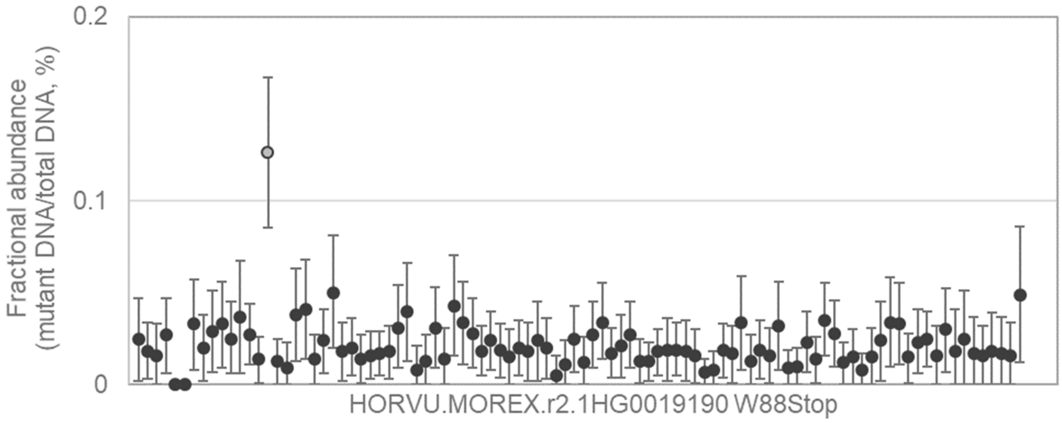


TCAAGGGCTTCTTCATCAACCTTTACAGGTTATTGACTCTGGTCAGAGTTATCGTGGTTATTCTATTCTT
CACGTG[G/A]CGCATGAGGCACCGGGACTCGGACGCGATGTGGCTGTGGTGGATCTCGGTCGTGGG
CGACCTCTGGTTC

10

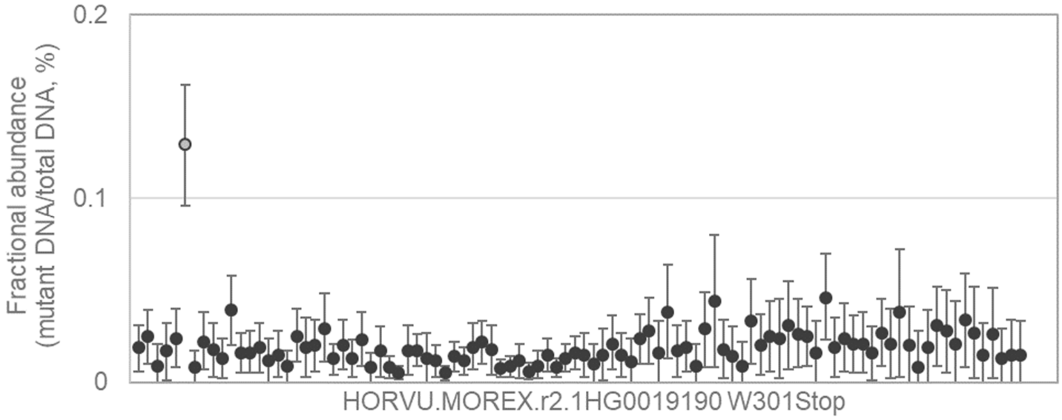


CGTGCAACCGCGCTGCAAACGAGAAAGAAGGGTGTGGGAACGCGACTTGGATGGCCGATGGGTCGA
CGCAATG[G/A]CAGGGGACGTGGATCAAGCCGGCCAAGGGCCACCGGAAAGGACACCATCCTGCAAT
TCTTCAGGTTATGC

11

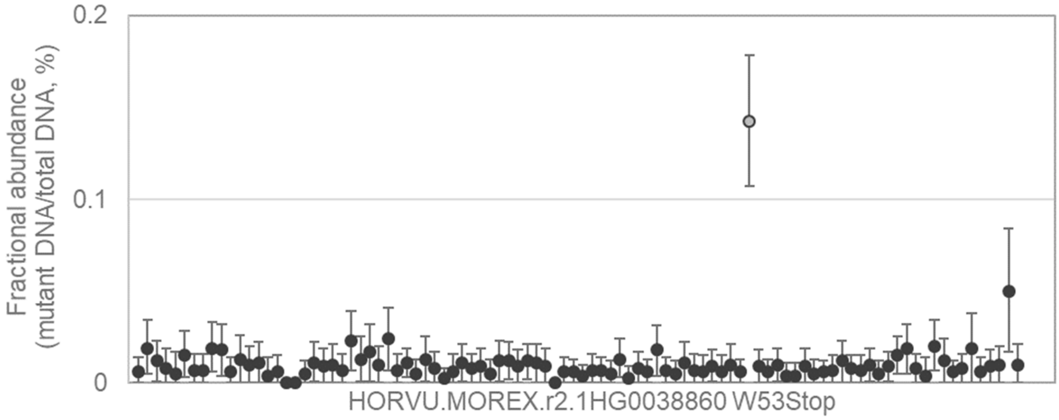


CACCGCCCGCGGACCGCCGGCGGCGAGCTCTCGGCCGCCCCGAGCAGATACCTGGCGCGGGAGGA
GCGCTG[G/A]ATGAGCCAACGCCTCGACCACTTCTCCTCCAGCGTAAGCCCTCTCCCCCTCCCAACCA
CCCGTGGATTTCG

12 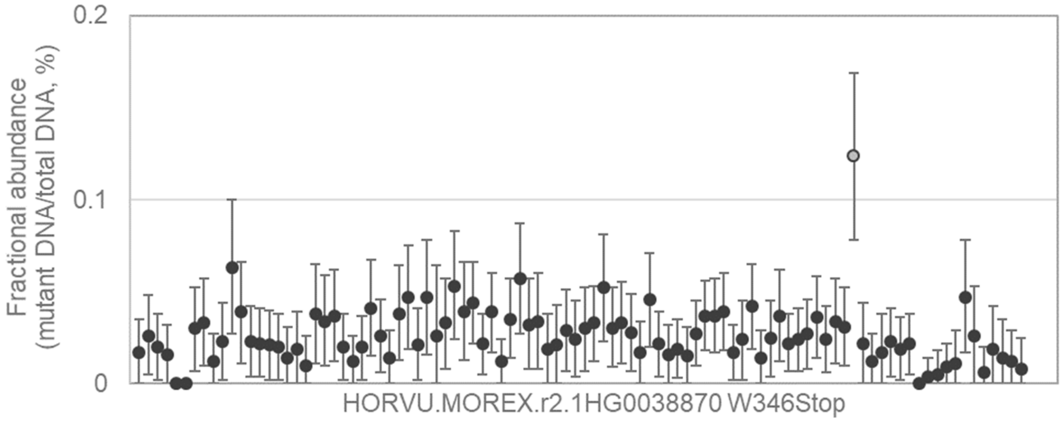


TTAATGCTGTCCGGATATTATATAATTGATCAGCATTTAACTGTTGAAATTTGATTTTGTGTAGCATATA
GATTGTG[G/A]TGGTATCAAGTTTGCAGTGAGGTTTCATATTTCCAAGTGGCACCAAAAAATGATAGCG
TCCGCTCTACAAAG

13 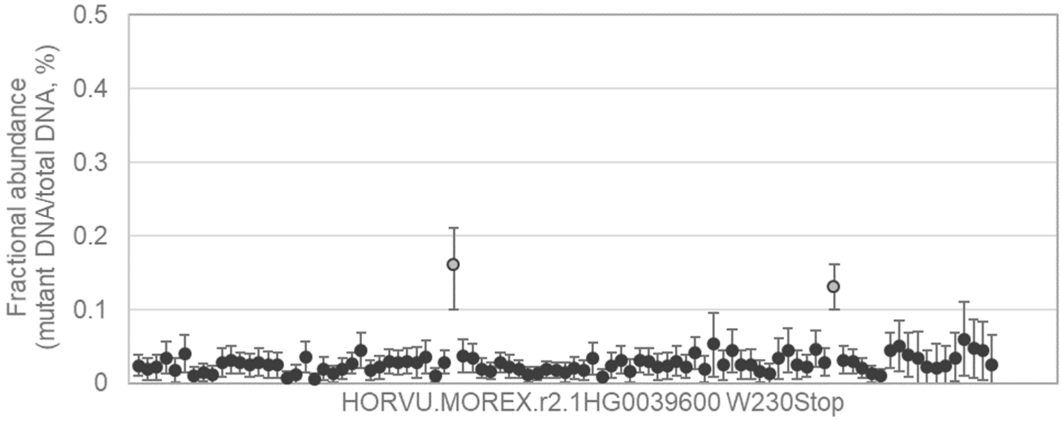


CCTGCTTGTTGTCACACTCAGGTGTATTATTACAGGAAATGGAGAAGCTGGATATATATTCCTGGGTCG
GCCATG[G/A]GGGCCCTTTGGGAGGGTTGTCTTTGCGCACACTTTTATGGACCGCTGTGTTAAGGCCA
CTGGGTGGCATAA

14 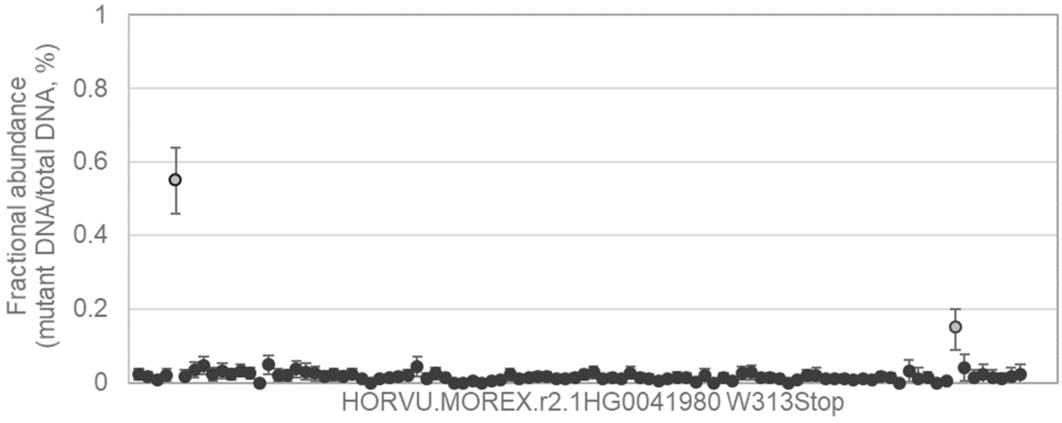


TTTTTTGCAGACAATTTTCTAACTTTTTCATCACATAACCAGCATCAATTATCATCATTGTGGGGCATTTA
AGACATG[G/A]TATGGGATACCAGGTGATGCTGCTCCTGGATTTGAAAAGGTGGCAAGCCAGTATGTG
TACAACAAGGATATT

15

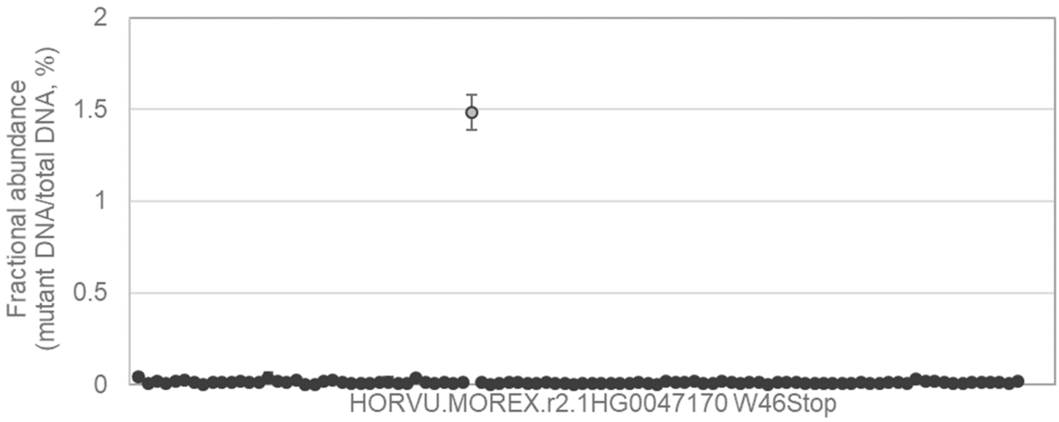


ATCCTATATAAGGGGCCTGCACCCCGTTGTGGCCTCACCAGAAAAGAAACAACAAGAGCTTTACAGAG
AGCCTTG[G/A]CATCACCCACCCACACCCTCACCCTCCAACGCAGGTAGAGAGAGAAAGAGAATGGCA
GGCCAAGGCGTTG

---

16

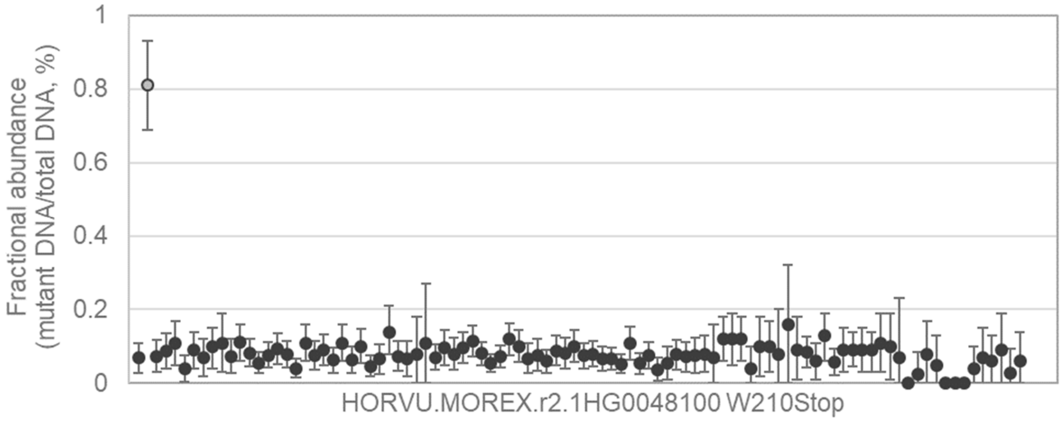


GCCAAGTTCCTCCTCACCAGCACTAGCGAGGTCTACGGTGATCCCCTCCAGCACCCTCAGGTGGAGA
CCTACTG[G/A]GGCAATGTCAACCCCATCGGTAAGAACTCTCCTCAATCCTCATACTCCCCCAACTCAA
TTTATTAAGTGGGAT

---

17

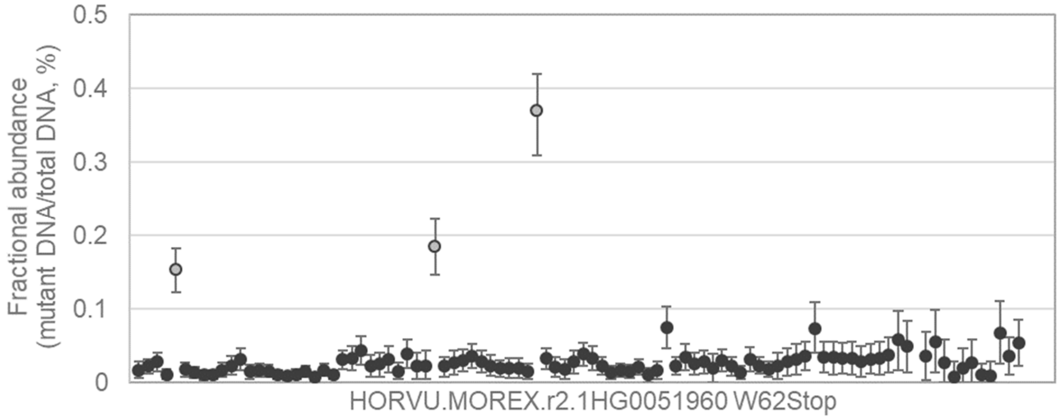


GCCGGCGGTGCGCCGGCCTGCCCATCTCCGTGCCCATCATCGCCGAGGTCGACCTCTACAAGTTCG
ACCCATG[G/A]CAGCTCCCAAGTACGTTCGCCTTGCCCTGATCATCGGTCCCCTCGATCGCTGCACCG
TTCGGCCAGGCCG

18

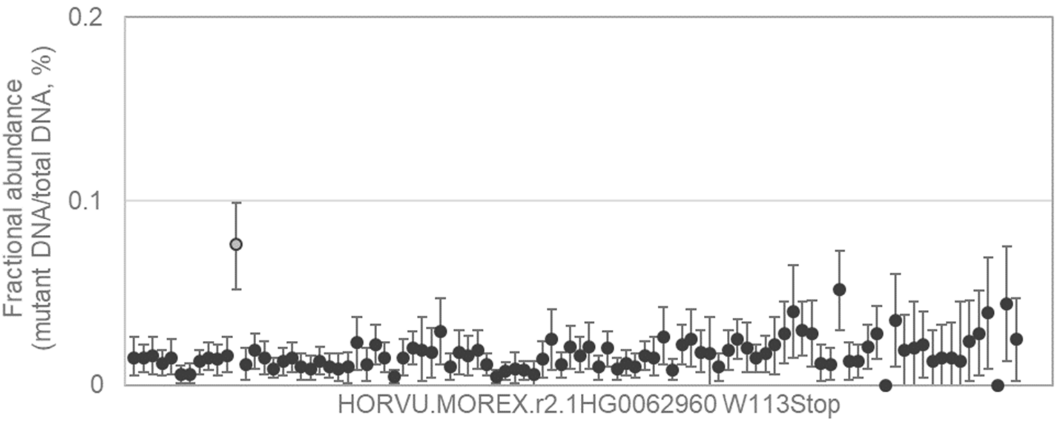


TCTTGGCAATGGTGGTCCTCGCGAAGCCGGCGGCGCTGGAGCAGATCGCGCTGATGAGGACTCCTG
AGCCGTG[G/A]GAGAGCTTCTCGGGCGTCCCGGCCGTGGACCTGTCCAGCCCCGGCGCCGCGGCGG
ACGTGGTGCGCGC

---

19

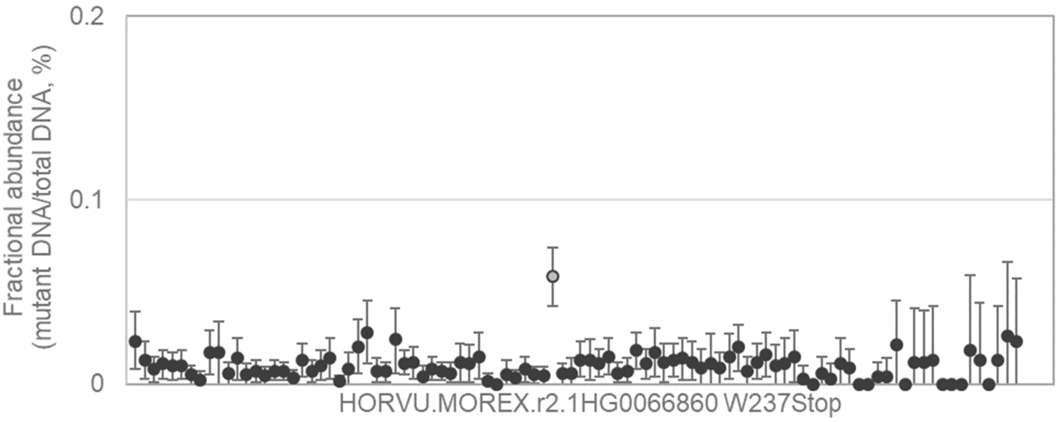


TTCCCTTAGGGAGCTGGAGTCCAGATTTATAGCTGCACAAGCAGATATGATGAAACTTGGTCCTAGGG
ATGCCTG[G/A]TGGGAGAAAGTAGAAAAATTGGAAGACTTGCTTGAGACCACAGCAAACCAAGTAGAG
CGTGCTGCTGTGAT

---

20

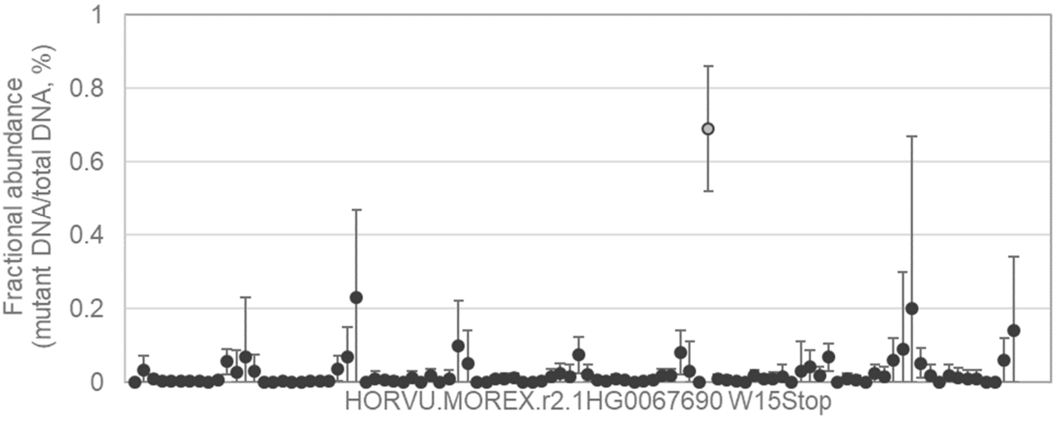


GACTGCTCTCCCGAACAAGGAGCCATCATGCGGGGGGCCGGCGGCGGCGGGACCCCGGGCCGGCC
GCGGTG[G/A]GGCGGCTCGGGCGCCACCACGCCGCGCTCGCTCAGCACCGGCTCCTCGCCGCGCG
GCTCCGACCGCAG

21 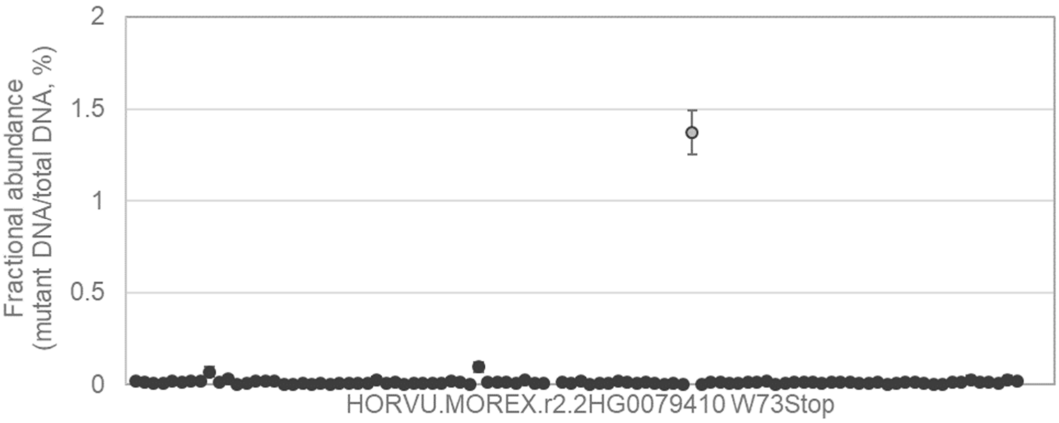


GACCTTTAAAGTTCAGTTCTTGTTTTGAGGTCGTAACTTCTGCCACACAGGGTATCAGTATGCATGAACATGGCTG[G/A]GTGCAGGGTCCGTCTGTAACTGGCAAGGAGAAAACGAGTTCACTTAACAATGTGAAATCACCGAGGTGCAA

---

22 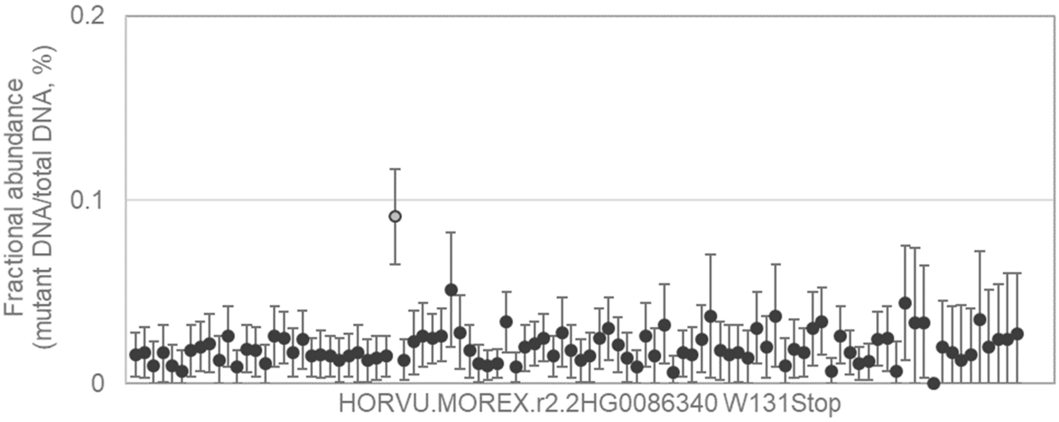


TAGCTTTCCTAGACTATCATGATATACCCTACAAAGTTGTGGAAGTTAATCCACTGAGCAAAAAGGAAATCAAGTG[G/A]TCTGAGTACAAGAAGGTCCCAATATTGACGGTAGATGGTGAACATCTGGTTGATTCTACAGGTTACGCTTGT

---

23 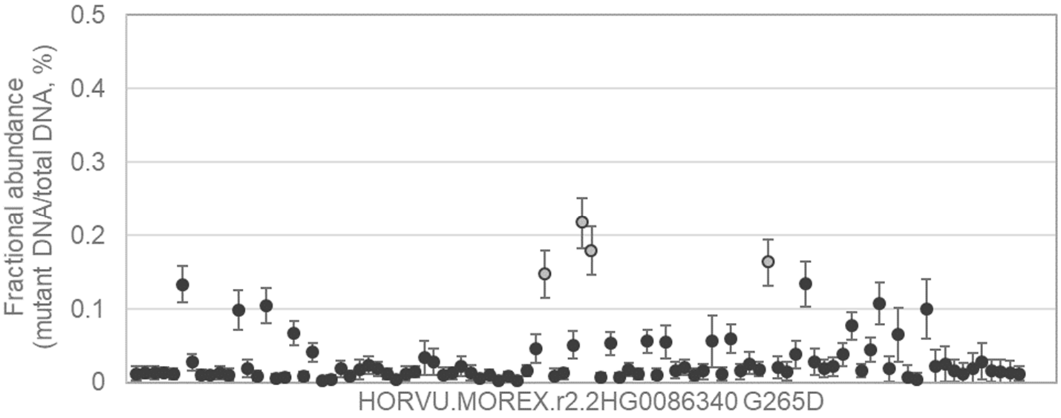


ACAGGAAAATTAGAATACAGCGAGTTTATTATATTCCTTAGCATTAATTTGGTTTTCACTTTCTTTGTCATCCA[G/A]TGGCTCCAAGCCTAACTTGGCAGATCTGGCAGCATTTGGTGTTCTGAGACCTATCAGATACCTGCAGT

24

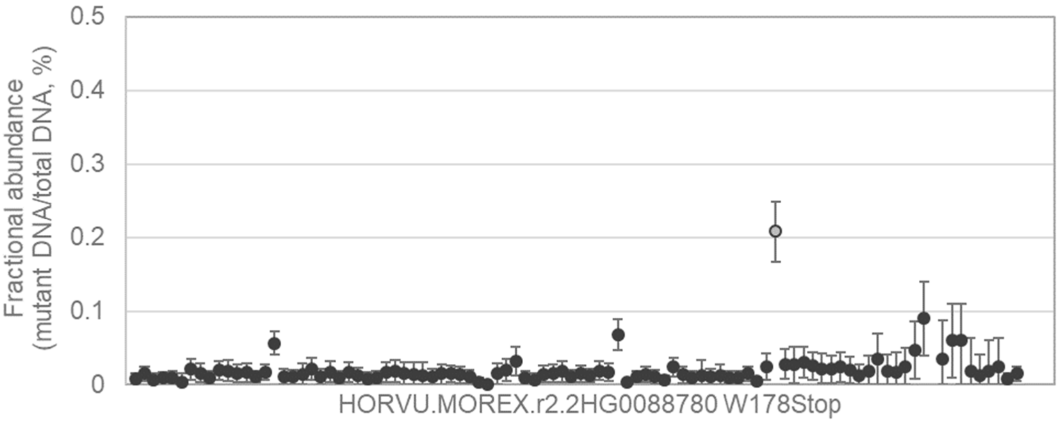


CAACACCAGCAGCAGCGGAGCAGCGGTGGTTGCGCAGCCGGAGCCGCCGGCGTGCAACCTGTACC
AGGGGTG[G/A]TGGACGTACGACGCCGAGGGGTCCCAGGTGCCGCTGTACCGGGAGCCGGAGTGCG
AGTTCCTGACGGAG

---

25

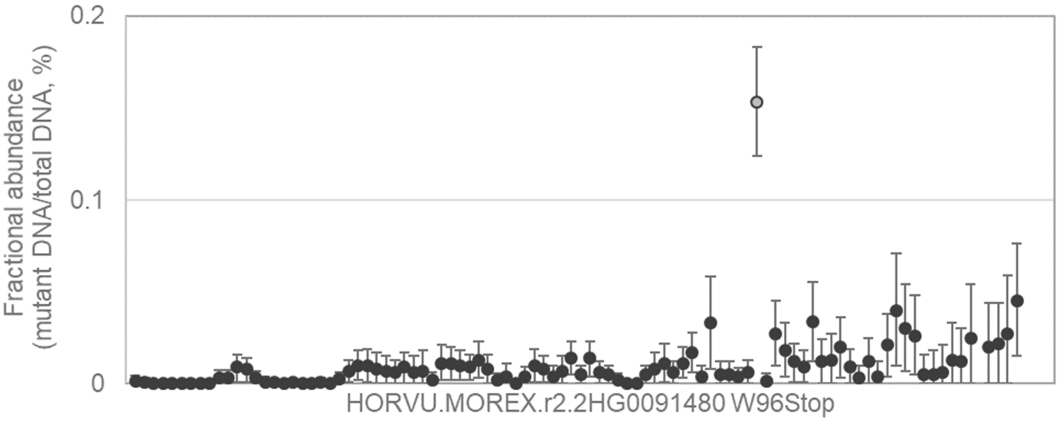


GCAACCGCGCCGCCGCCGCGCCGTGCCCGGTCCCCATCATCGCCGACGTCGACATCTACAAGTTCG
ACCCATG[G/A]GACCTCCCCTGTAAGTTCTCAACCAGTCAACCCATCACCGCTGTATGCAATCCCTGCT
TTGCTCTGCTTGCT

---

26

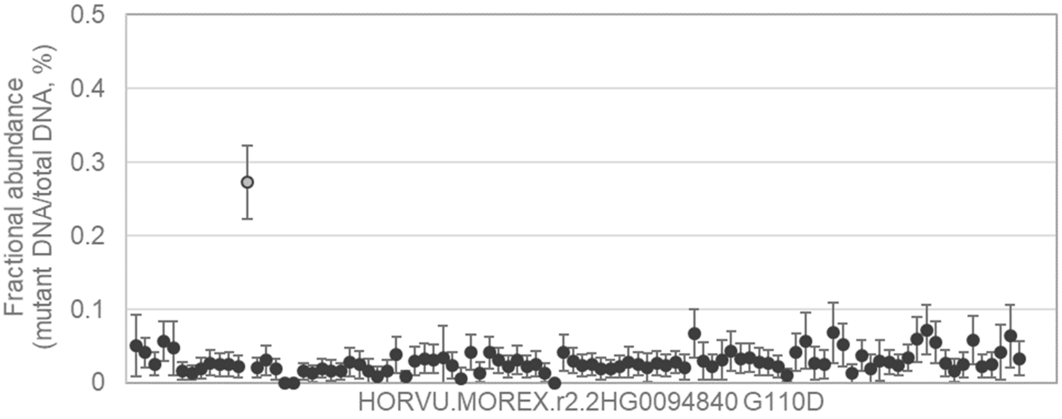


GCTACATCCGCTGCGACGTGACCGATGAGGCGCAGGTGGCCGCCGCCGTGGACCTCGCGGTGGCG
CGGC[G/A]CCGGCTCGACGTCATGTTCAACAACGCCGGGATCACGGGCGGCAACTACGCGGCCGCC
CCGATCGA

27

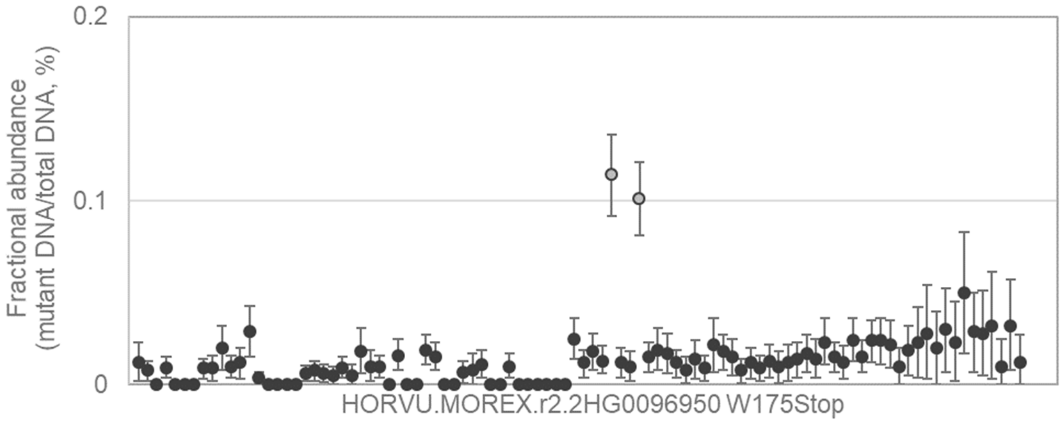


GCTACGGCAAGTCGATCTACGAGGACTGCACCCTCATGTCCAACATGCCGGCGTCGTCCCAGCAGCC
CGGGTG[G/A]GTGACCGCGCACGGCCGGGCCGCCGACAGGAGCGCCGCCTTCGTGTTCAAGGGCGG
CGTGATCACGGGC

---

28

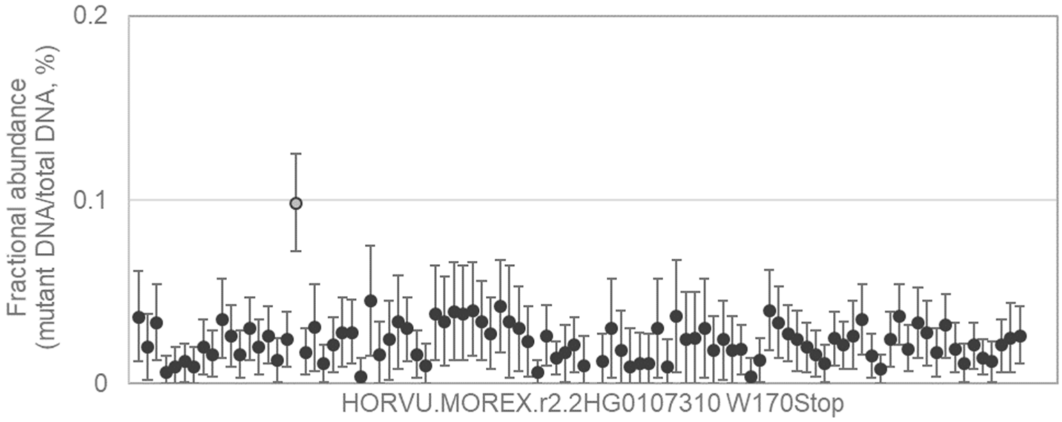


GCAAAAACAGATGGGTTCCAAAAAGGCAGCTGAGAAGACTGAAGGTCAGCTACTGGGAGTATTTGATT
ATGCATG[G/A]AATAAACTGCAGAATGGTGCGTGTCGTCGCGAAGAAATACTGCTCAAGTTAGAACAGG
GAAGCAATAGATTA

---

29

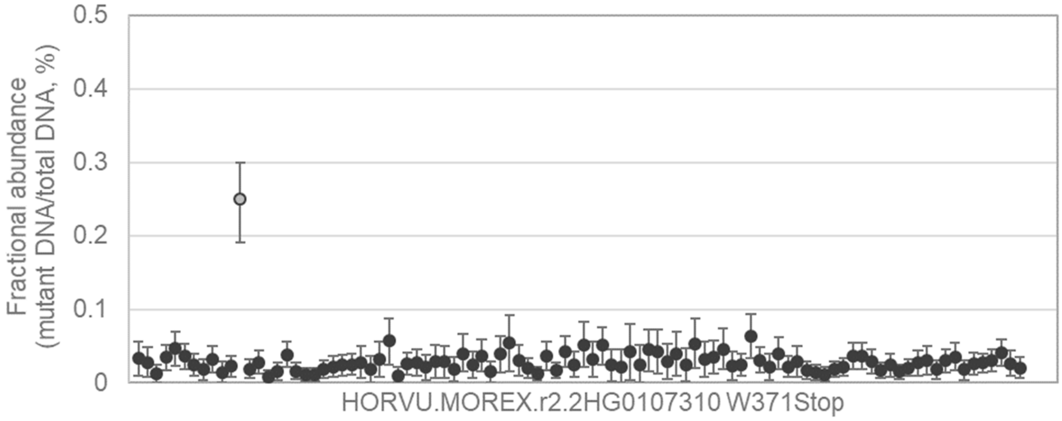


CTTGCCAAGTTGCTATTCCAACTGTTGAATCCATACCTCAACAAACTGCTAATCCAAGCAAACGAGATC
AAGCCTG[G/A]CAAACCAGTGCAGACCAATCCAAAAATCTGTCCACAAATGCAAAGGAGCCATCTTGGC
AAAGGAACAGCTTC

30 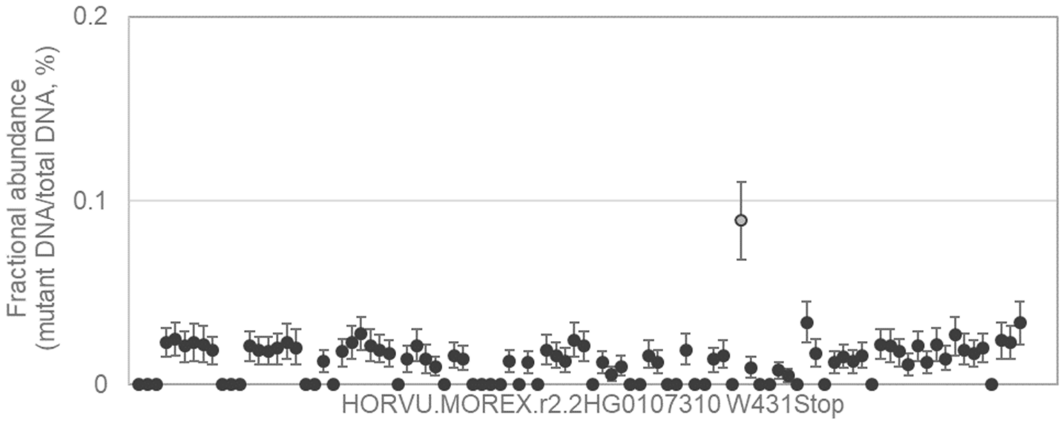


TACAGAAGATGAAGCATATGGAACCTCCCATGCAGAATCTCAGTTCCACGAAGATGAGCCTTGTGGAA GGAAGTG[G/A]TTTGAGCGGCTCAAAGCACAGAAATCAGCCAACGCACCATCGTCGAGAGGCCAAGG ATGTCAGCCAAGAGA

---

31 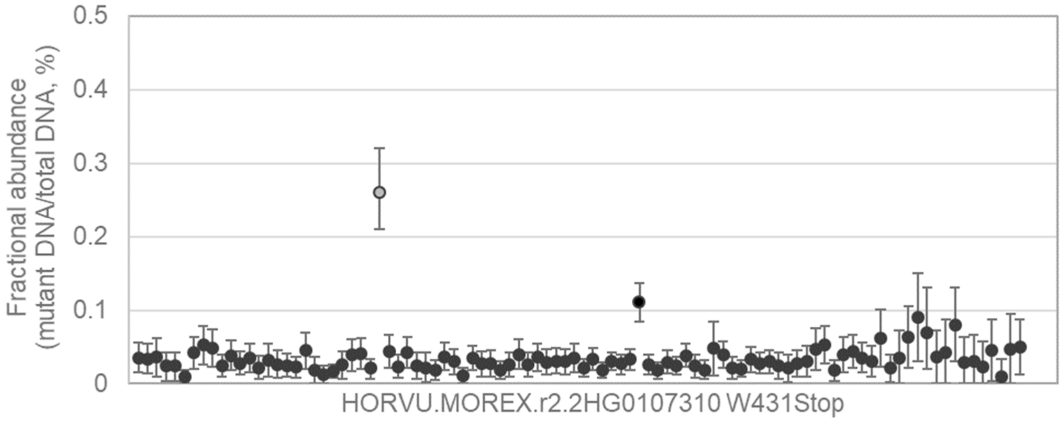


TACAGAAGATGAAGCATATGGAACCTCCCATGCAGAATCTCAGTTCCACGAAGATGAGCCTTGTGGAA GGAAGTG[G/A]TTTGAGCGGCTCAAAGCACAGAAATCAGCCAACGCACCATCGTCGAGAGGCCAAGG ATGTCAGCCAAGAGA

---

32 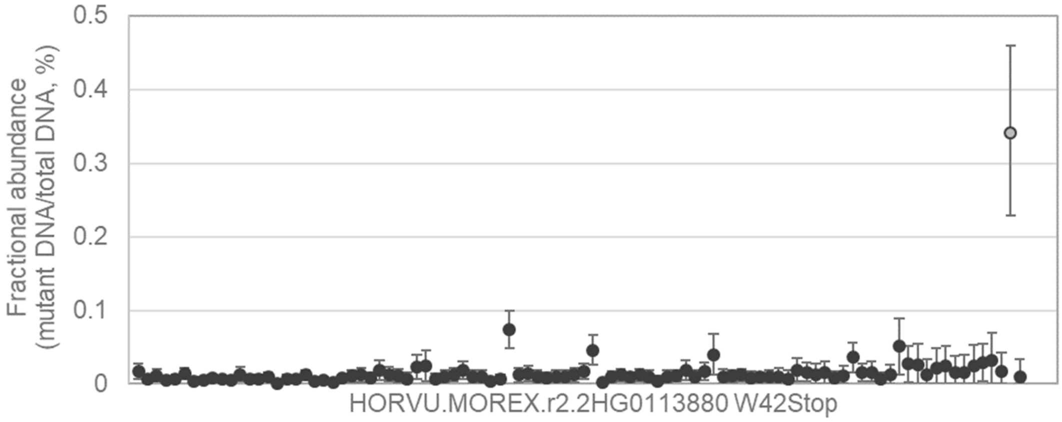


TCGTCGCTGCCTTATTCCTAACCCCGCCGGCCTTGGCGGCGGGGGGGAAGACCGGCCAGGTGACGG TGTTCTG[G/A]GGACGGAATAAGGCCGAGGGGTCCCTGCGTGAGGCCTGCGACTCCGGCATGTACAC CATGGTCACCATGT

33

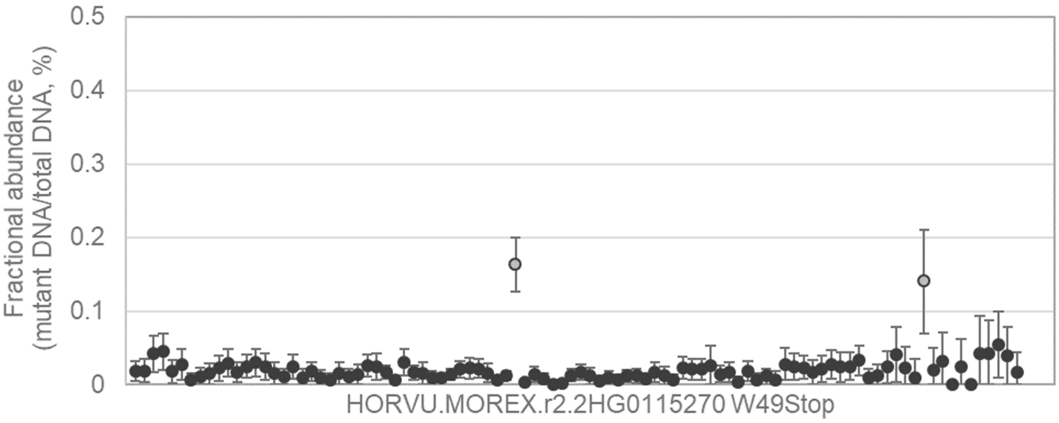


TGAAAAGGAAGATTATGGGGAAGAAACTTTTTGTTCAAGCTATCTCAGAGGTTGAGCTGTACAAATTTG
CTCCTTG[G/A]GACCTTCCTGGTAAATTTCTTCTATTGTTTGATTTTTTATGTAATATACTTACACCCAAAT
GAGCTTGAAATTAA

---

34

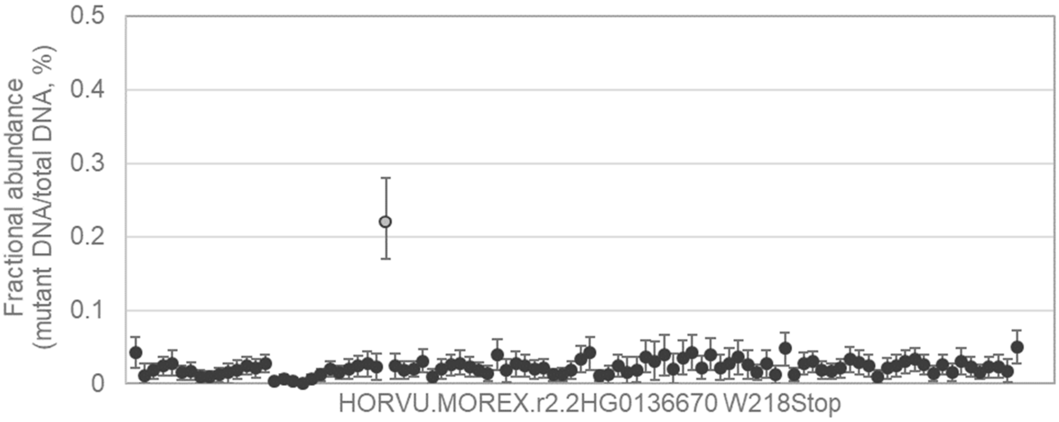


CAGTTACTCTAAATTTATGACCTGGCTTTTGCATCCTGTTACAGTCTGCAGCATTAGTAGGTGACTTCA
ACAATTG[G/A]AACCCAAATGCAGATACTATGACCAGAGTATGTCTACAGCTTGGCAATCTTCCACCTTT
GCTTCTTAACTATT

---

35

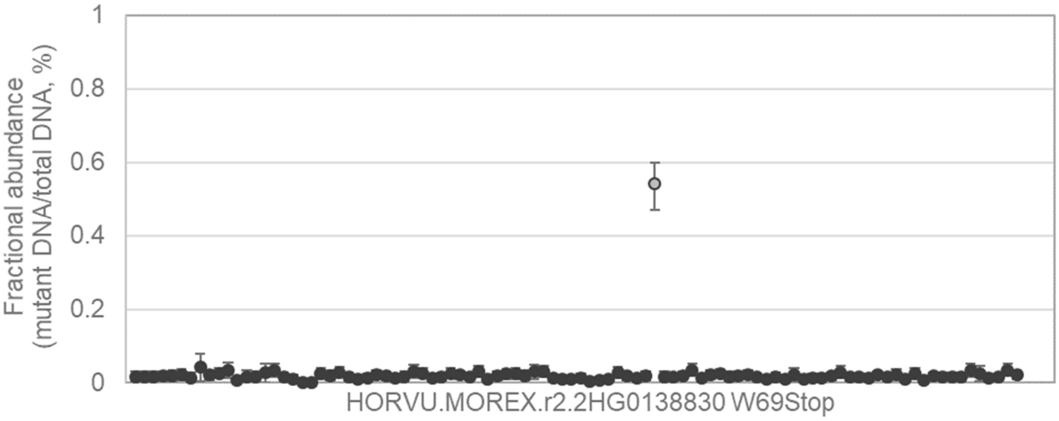


GGACGTGGTTGGCGGCGCTCGTCTGCGAGGCGTGGTTCGCCTTCGTGTGGATCCTCAACATGAACG
GCAAGTG[G/A]AGCCCCGTCCGGTTCGACACCTACCCCGACAACCTCGCCAACAGGTACTCTACGTAC
GTACCCACGGCGA

36 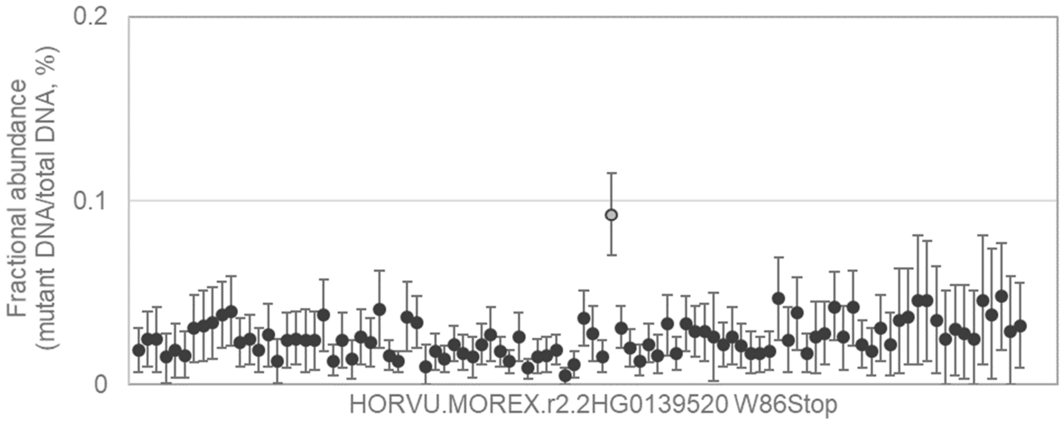


TACCTAAAATGTCCAGTGTGCTCTGTTCGTTTTTGTGTGGTTGTTAGTATTCTAATGAAACTTCTATGAC
AGGCCTG[G/A]CGTTCTGTGTGTAACACCCAGGTGACCTGTAAGAATTATTTAAGACCTGGTTTATCTG
CAAAAGTGAAAGATT

---

37 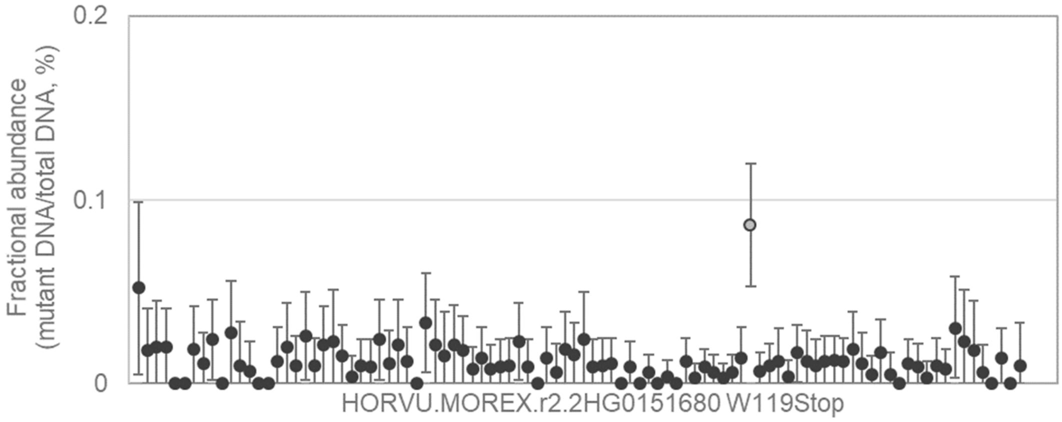


AAGATCATCCGGCTGTCGACCGACGTCCGCATCTCCTTCCGCGCCTACACGACGTGCCTGCAGTCCA
CTGAGTG[G/A]CACATCGACAGCGAGCTGGCGGCGGGCCGCCGGCACGTGATCACCGGCCCGGTCA
AGGACCCGAGCCCG

---

38 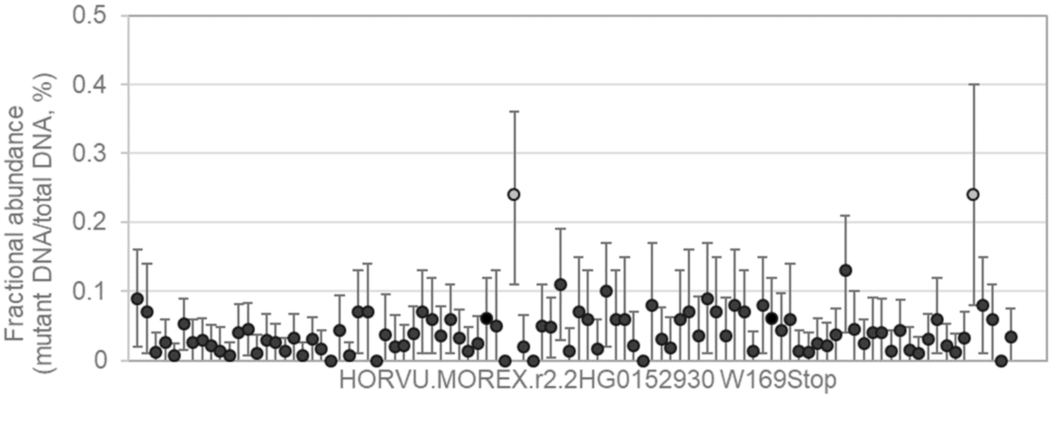


AGACTGGGAGCGACTGACAGGAGGTGCGGCGCCTCAGGTCGGCAGCTGGGAGCCACGGGGCATCT
GTG[G/A]ATGGCGGACACGCCGCTGGCGCCGTTGGCGTGTGCGGCGGGAGCCCGAGCTCGTCCTTC
TCGACGGGAACC

39 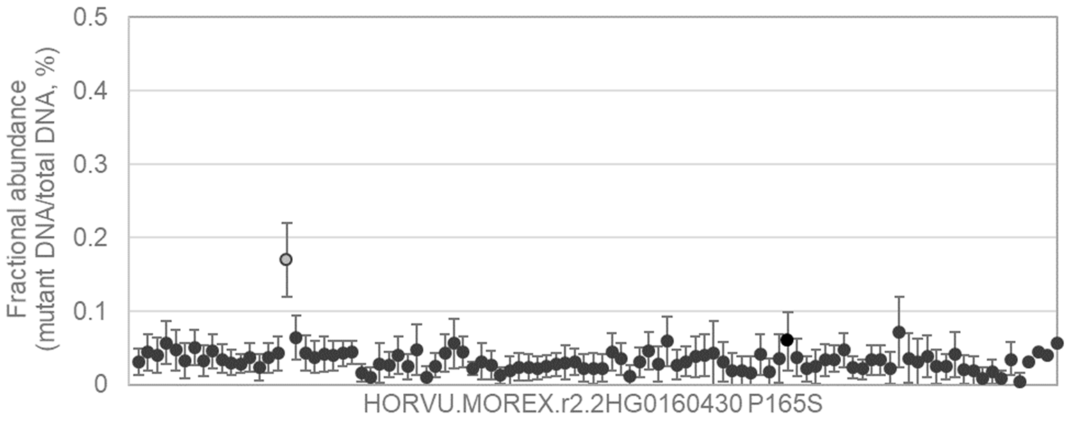


AGGCCGCTCAGGACTCCAAGTACTGCGAGCGCCACATGCACCGCGGCCGCAACCGTTCAAGAAAG
[C/T]CTGTGGAAACGCAGCTCGTCGCCAGCTCCCACTCCCAGTCCCAGCAGCACGCCACCGCCGCC

---

40 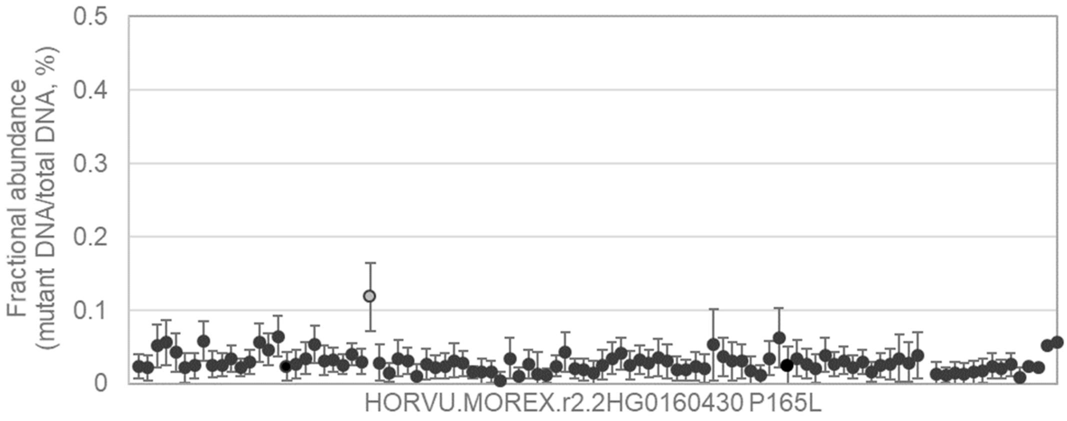


GGCCGCTCAGGACTCCAAGTACTGCGAGCGCCACATGCACCGCGGCCGCAACCGTTCAAGAAAGC
[C/T]TGTGGAAACGCAGCTCGTCGCCAGCTCCCACTCCCAGTCCCAGCAGCACGCCACCGCCGCCT

---

41 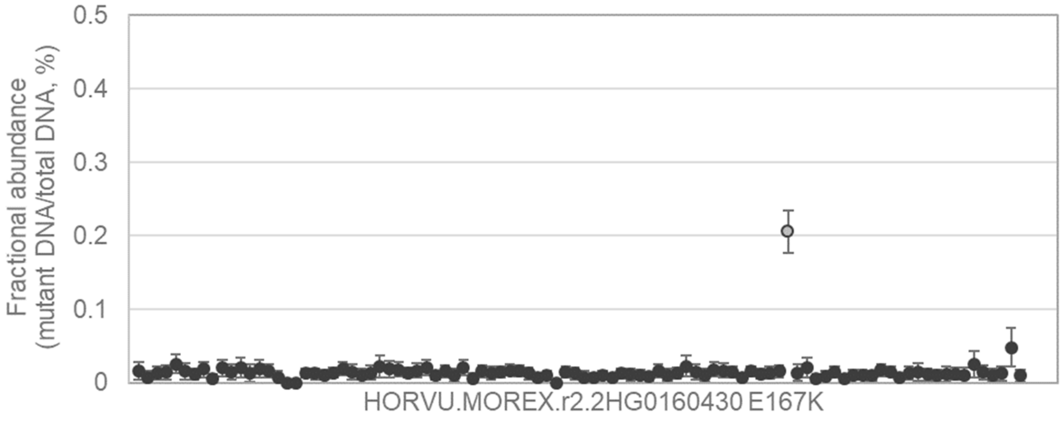


GAGGCCGCTCAGGACTCCAAGTACTGCGAGCGCCACATGCACCGCGGCCGCAACCGTTCAAGAAAG
CCTG[G/A]AAACGCAGCTCGTCGCCAGCTCCCACTCCCAGTCCCAGCAGCACGCCACCGCCGCCTTC
CACAACCA

42

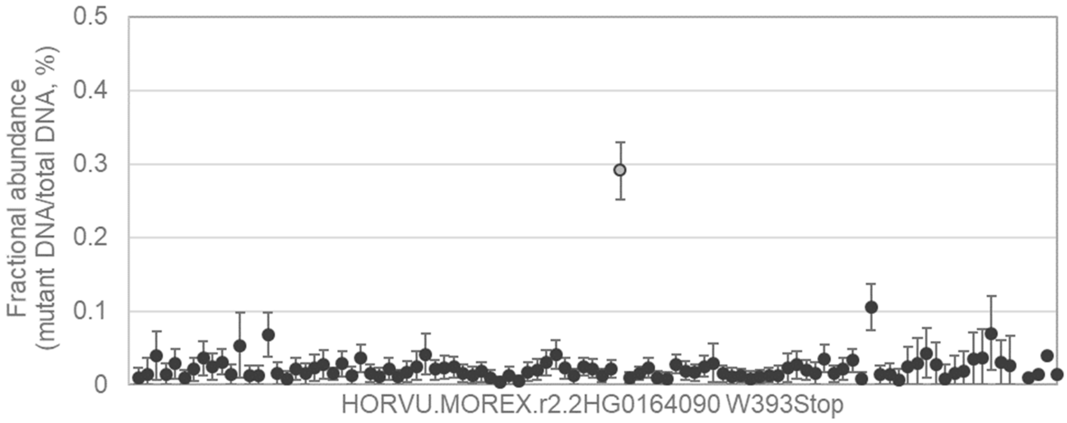


CAAAGGGGCATTCAATGGTTGGAGATGGAACCATTTCACTGAAAGATTGCATAAAAGTGAATTATCAG
GGGATTG[G/A]TGGTGTTGTAAGCTATACATACCCAAGCAGGCATACAGATTAGACTTTGTGTTCTTTAA
TGGTGACACCGTCT

---

43

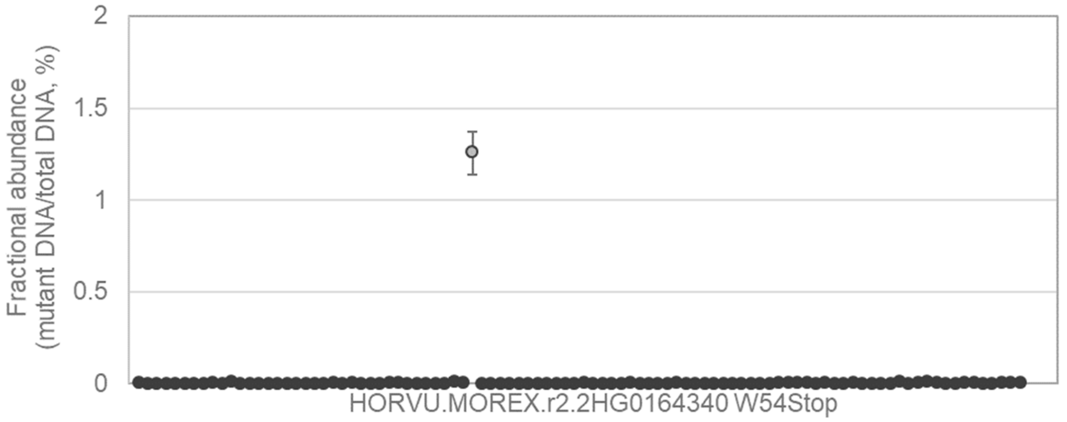


CTCTGACCATGGATGGGATTGGATGGGTGGCAGCGGTGGGGATCCGGTACGGCAAGTTCTGCGGCG
TGGGGTG[G/A]AGCGGGTGCGAGGGGGAAGAGCCCTGCGACGACCTCGACGCCTGCTGCCGCGACC
ACGACCACTGCGTC

---

44

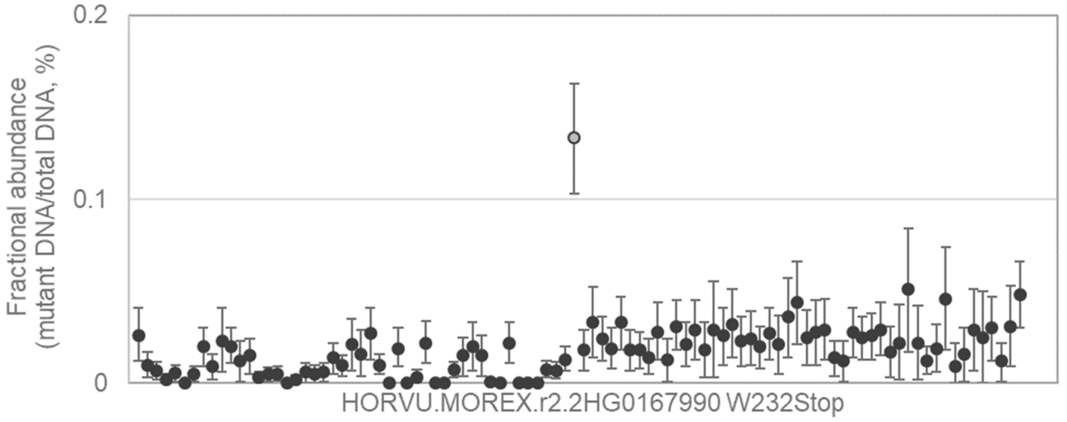


CATTCAACCGCGAGGTTCTGGGGCCCGTGATGTTCCCTGGCCTCCGGATTCGCAAGGAAGGGAAGCA
CAGGTG[G/A]GACACCCTTGAAGACATATGGAACGGCCTGTGTGCTAAGGTGATCTGCGACAGCTTGG
GCTACGGCGTGAA

45

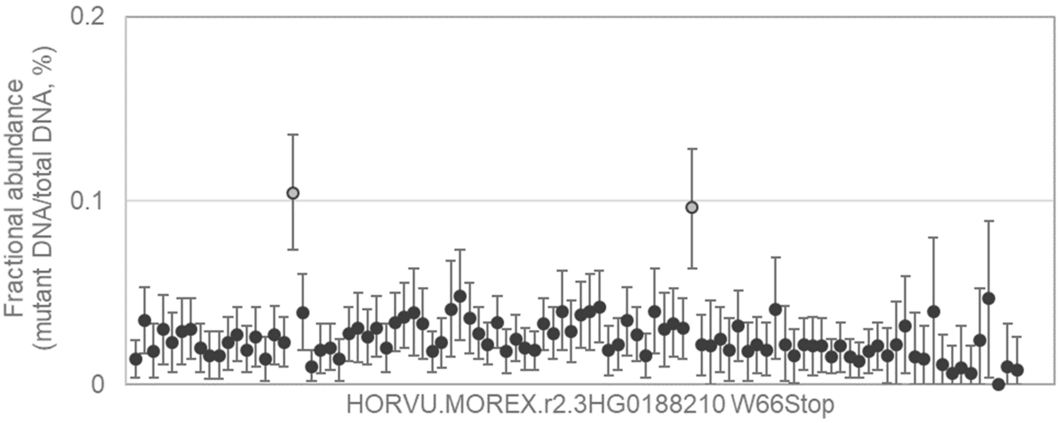


GTTTAACCGGTTCGTCGGGAAGGATATAATGACTGATATTAGGTACCCGGTGAAACAGAGTACTGTGC
CTGCATG[G/A]GCTGTCCCTGTATGTTTCTGAACCTCTGCAATTTCATACCCTGACTCAAGCTTATACTC
ACCCTTCTCTGATA

---

46

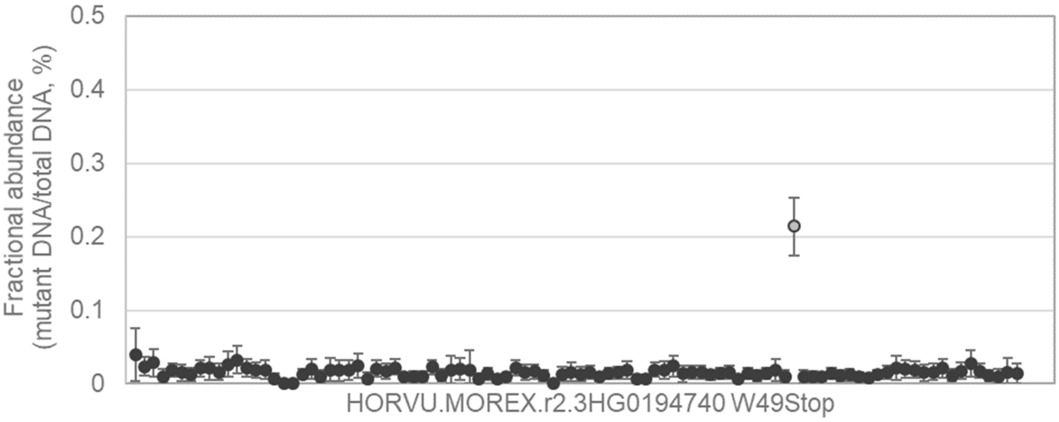


AACAAAAGCAGGTAGGTTACCCTGAAGTTTTACTCAACCTTGTTCTTTATTGTGCAGCCACCACGTTGA
CAAGCTG[G/A]CATCTCTTGGTGACATTCTGTTCGCTCCATGTGGCATTATGCATGAAGCTCTTTGAGC
ACAAACCTTTCGATG

---

47

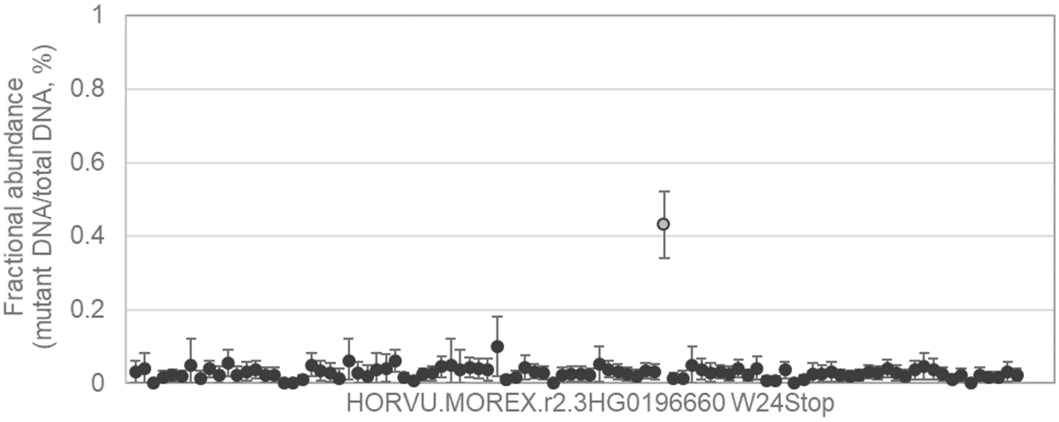


CTAATGGCGATCGTTCACGTGTTACTGGTCGCTCTGATCACCGCGGCGTCTCCTACGGCGCACGGCC
AGACGTG[G/A]CACGGCGACCTCGCGGCGCTCGCCGCGGCAGGCAAGCTCCGCAGCGACCCCAACG
CCACCCTGGCGGC

48

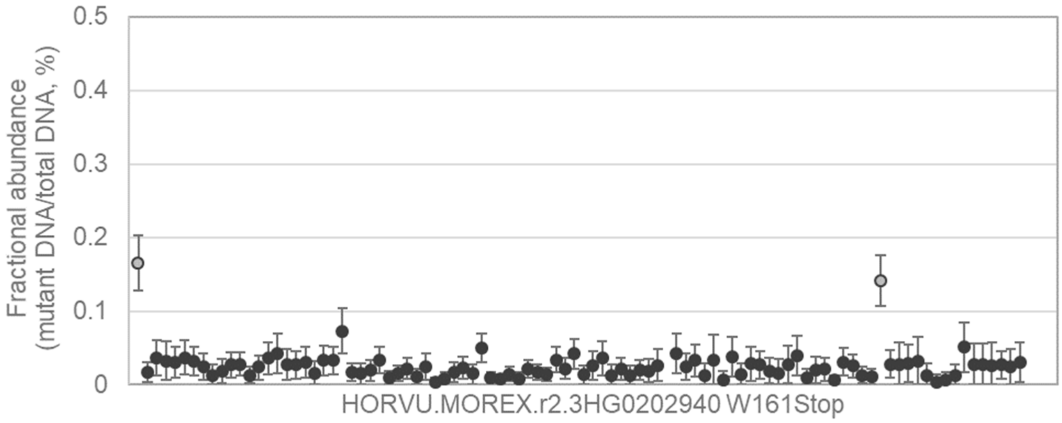


GAAGCTCCACTGCGTCCCGCCTCTCCGTGTCAGTCGAAGGCCAGTGCATAGACGCCGGCGGCGAAC
AGCTGTG[G/A]GTGGATGTGCTGCGCACCGCCCTCGCGCATGGCCTAACGCCGCGTTCCTGGACCGA
CTACCTCCACCTCA

---

49

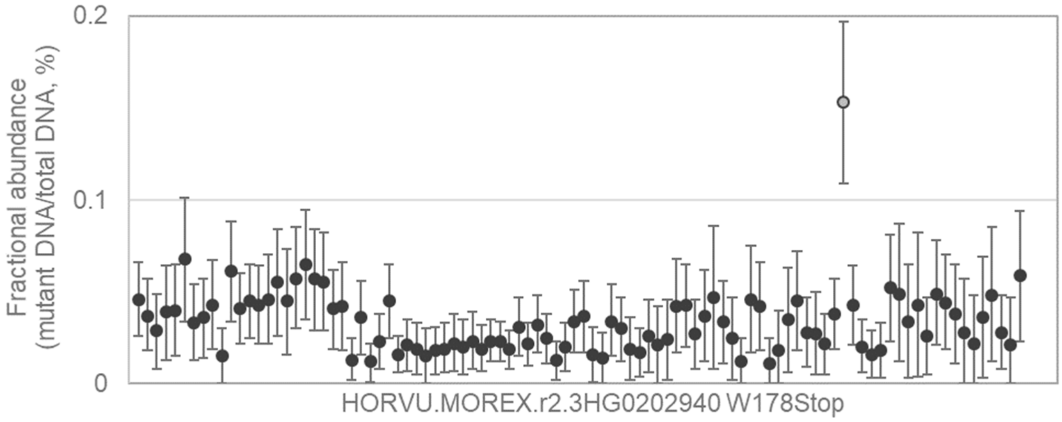


CGCCGGCGGCGAACAGCTGTGGGTGGATGTGCTGCGCACCGCCCTCGCGCATGGCCTAACGCCGC
GTTCCTG[G/A]ACCGACTACCTCCACCTCACCGTCGGCGGCACGCTCTCCAACGCCGGCATCAGCGG
CCAGGCCTTCCGCT

---

50

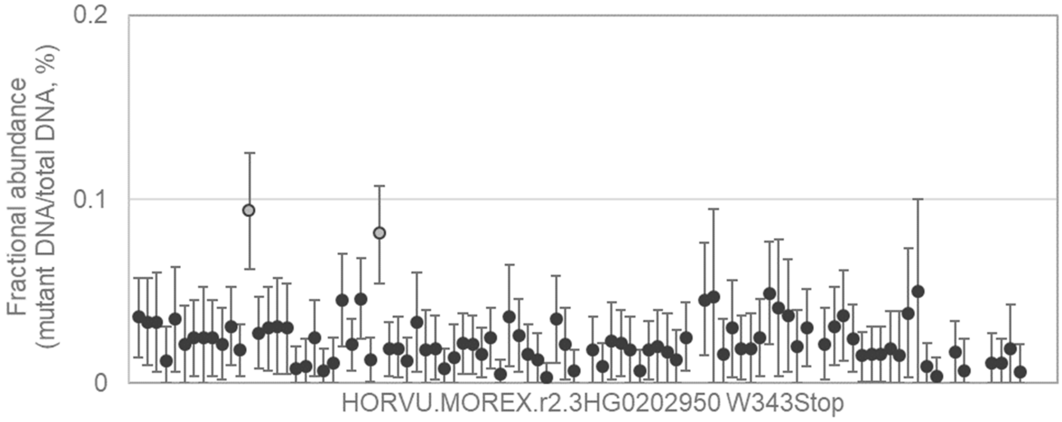


CGCAGTCTCCGGACTCATGGACTACGTCGAGGGCTCGGTGGTGCTCGCAGACCAGGGCCGAGCAGG
GAGCTG[G/A]CGCTCGTCCTTCTTCTCGGACGCCGACGCGGCGCGCATCGCCGCGCTCGCGGAGGA
GGCGGGCGGCATC

51

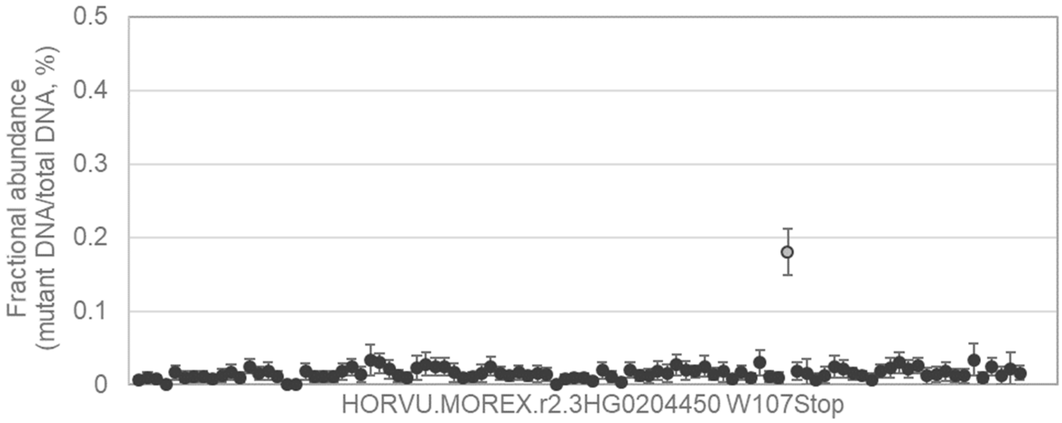


AAGGAGGCCGCCGCCGGCCGGCCGCTCGGGTACGGCAGCAAGCGGATCGGCGTCTCCGGCGACCT
CGGGTG[G/A]ATCGAGTACCTCCTGCTGGGCGTCACCCCCGCCGGCGCGCTGCCCGCCGCCTCCTTC
GCGTCCTGGACAT

---

52

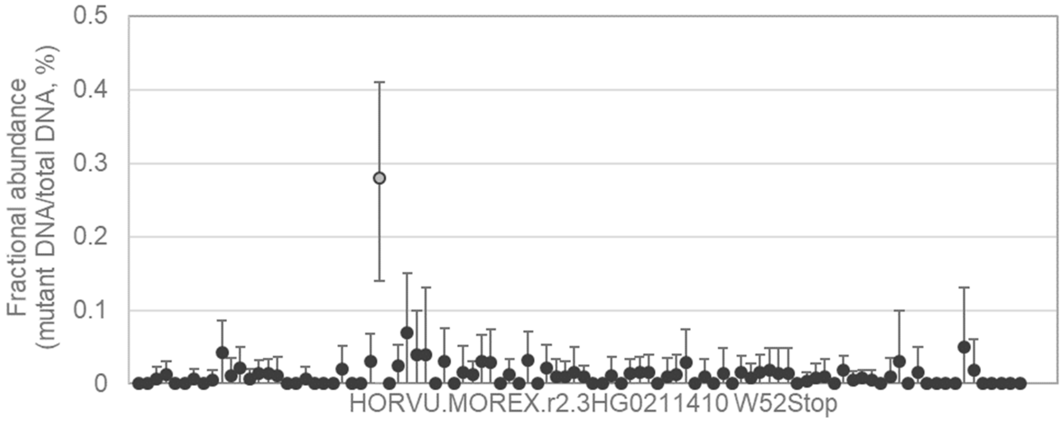


TGAAAAAGAAAATACTCGGTCAGAAAATTGAGTATGACATTATACCAGAGGTGGATATATACAAGCATG
AACCATG[G/A]GATTTACCTGGTAAGTTGAACATGCTTCTATATGGTTGCCTGTGATTATTTTGGTTCAT
AGAAAAATACAGTAT

---

53

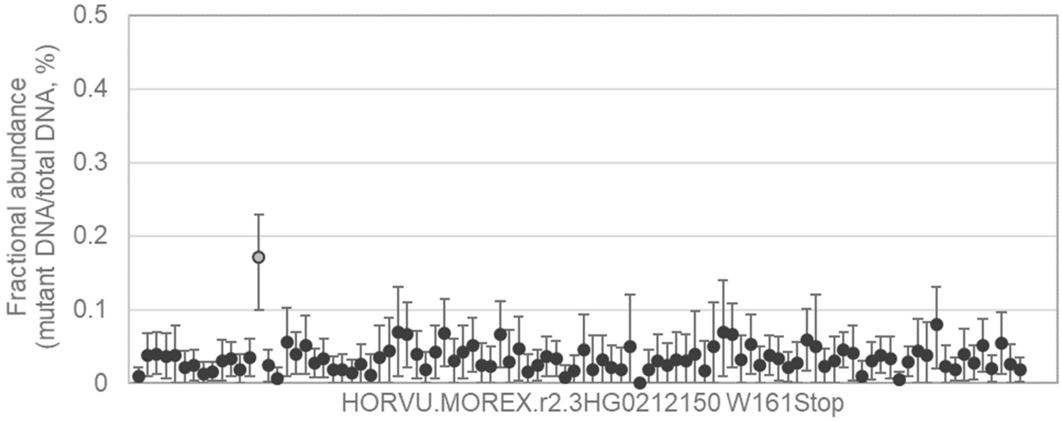


CGAAGCAAGAGCAGTGCTGCAGTGGCACTCAAAGGCTTGCAGTTTGTGACTGCGAAAGTTGGCAACG
ATGGCTG[G/A]ACTGCCGTGGAGAAACGATTCAATCACCTACAGGTTGATGGCATGCTACTCCGTTCAA
GATTTGGGAAATGC

54 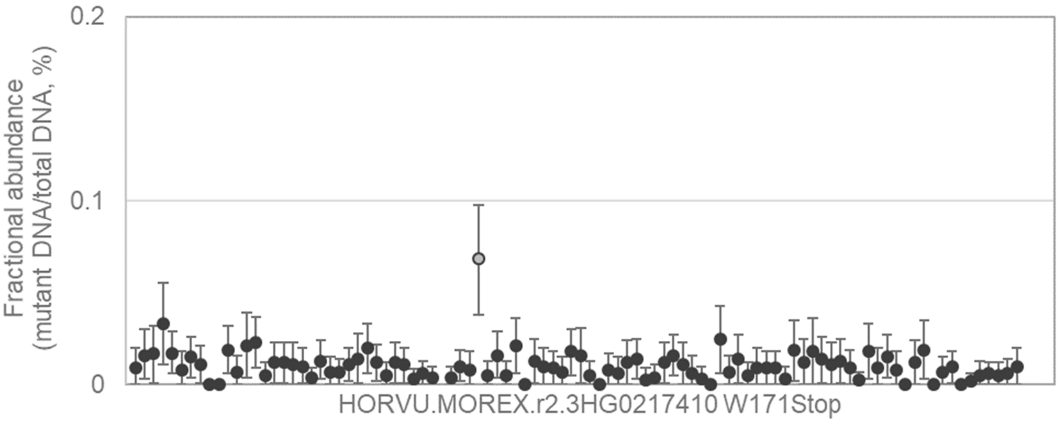


TGCAGGCTGAACAAGGGAGTTCTTTTCTTCCTGGCATCAAGCAACGGTTGCTATCATTTATTTTCCGCA
AGACATG[G/A]AATGAGGAACCTGATCAGGCTTTGGTTAGCTACTTAGCTATCTTACATGCCATTCATTC
ATTTGCTATGTGAT

---

55 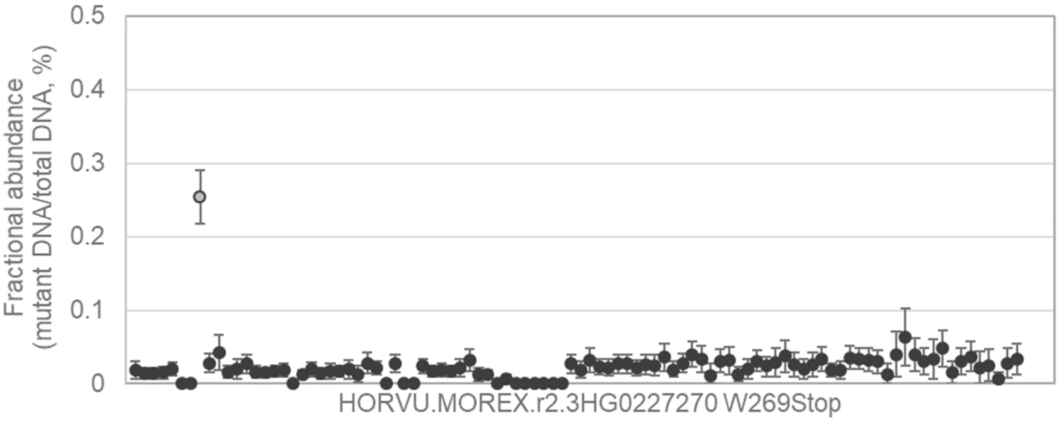


GCACGGGATACAGCTTCGTCAGGTGCAAGCTGACGGGCGTCGGGGCCGGCACCTCCATCCTGGGGC
GTCCATG[G/A]GGCCAGTACTCCCGCGTCGTGTTCGCTCTCACCGACATGTCCGCAGCGGTGAACCCT
CGAGGCTGGGACC

---

56 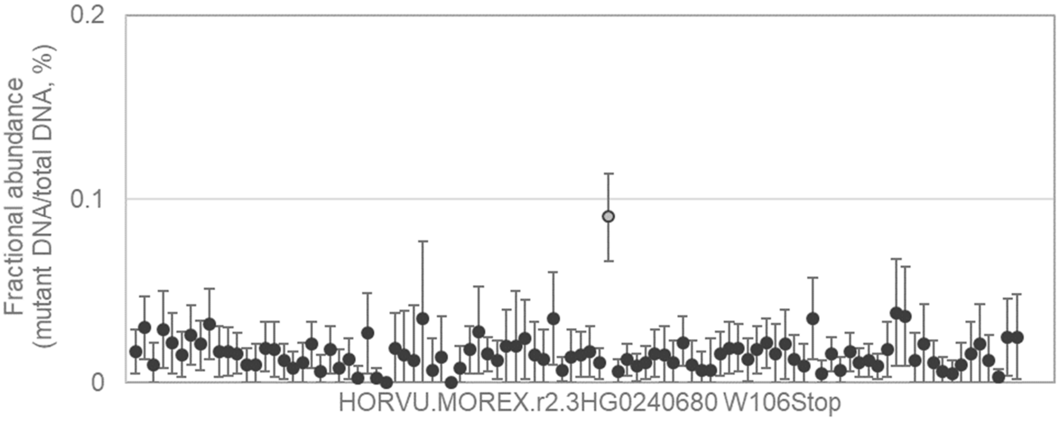


GGACCGCTCCGGCCCGGCCTACCCGTTCGGCTACGGCAGCAAGCGCATCGGGCTCAATGGCGACAT
GGGGTG[G/A]CTCGAGTACCTCCTCCTCGCCGTCGACTCCGCCTCGCTCCCCGCCGCCTCCGCCGTC
CCGTCCTGCGCGC

57

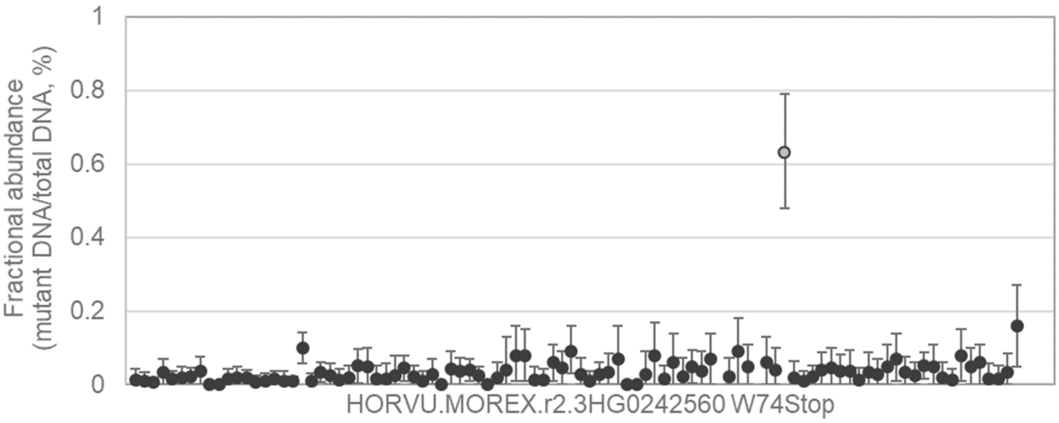


CAGAGAGCGTCGCCTTCGACGCCCAAGGCCACGGCCCGTACAGCGGCGTCTCCGACGGCCGTATTT
TAAAGTG[G/A]AACGGCGACAAGATAGGCTGGACCACTTACGCATATGGCCCCGGCTACGACGAGGA
CATGTGCAGGGCTT

---

58

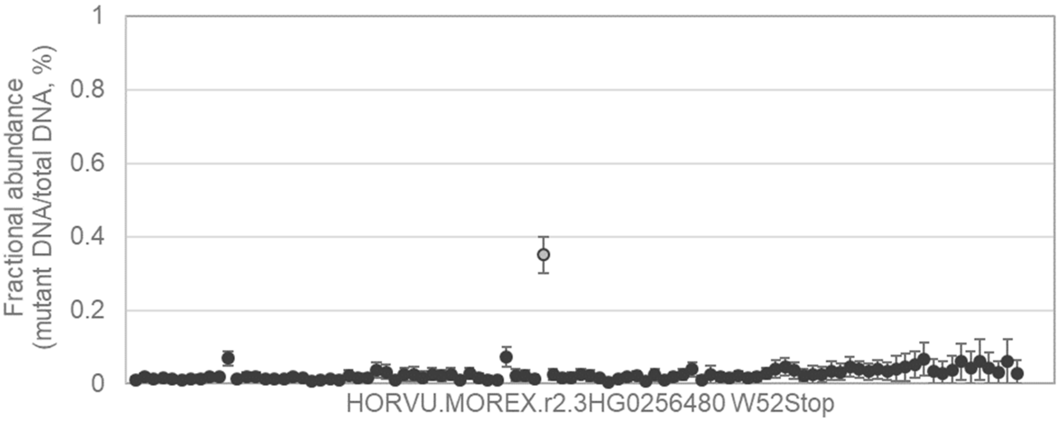


GCCGCCGCTGCGCCGGCGCGCCCATCGCCGTCCCCATCATCACCGAGATCGACCTCTACAAGTTCG
ACCCCTG[G/A]CAGCTCCCAAGTACGTCCTCCATCTCCTCTCTCTCTCTCATCGATCCGTCGTGACTGT
TACCCACACGTTCA

---

59

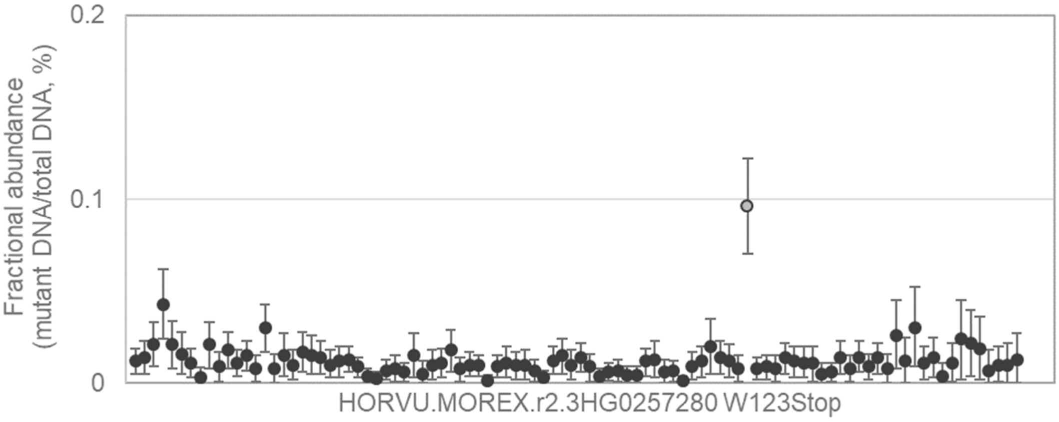


CGAGGTGGAGGAGGACGTGGAGAAGTACGCGGAGCAGCTGCTGGTGCTGGCGCACGCGTACCGTG
TGCCGTG[G/A]CTGAAGCTGTGGTGCCAGGAGGCCATCGGGTCGCGGCTCACCCCGGGCACCGTCG
TCGACGCGCTGCAG

60

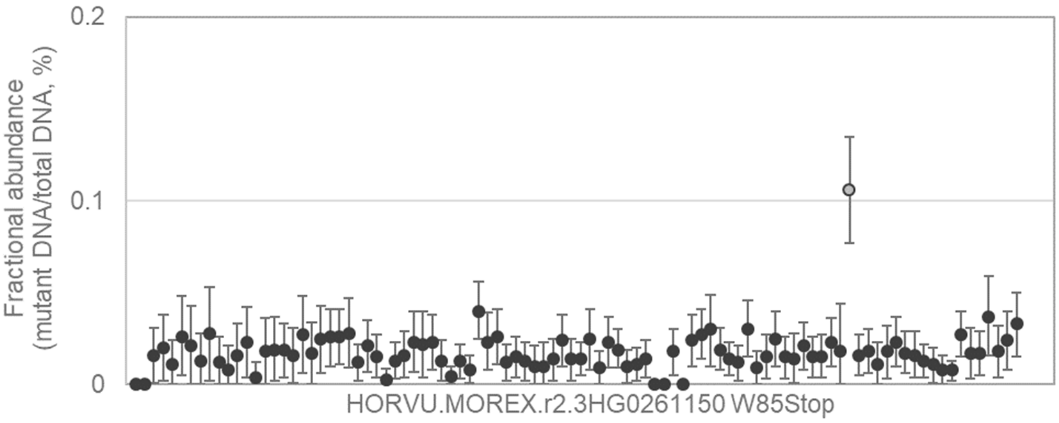


GGCACGGGCGCAACCTGTTCTACTACGTGGTGGAGTCGGAGCGCGACCCGGGCAAGGACCCCGTCG
TGCTCTG[G/A]CTCAACGGCGGCCCCGGCTGCTCCAGCTTCGACGGCTTCGTCTACGAGCACGGTGC
GTCGTTCCTCCTCC

---

61

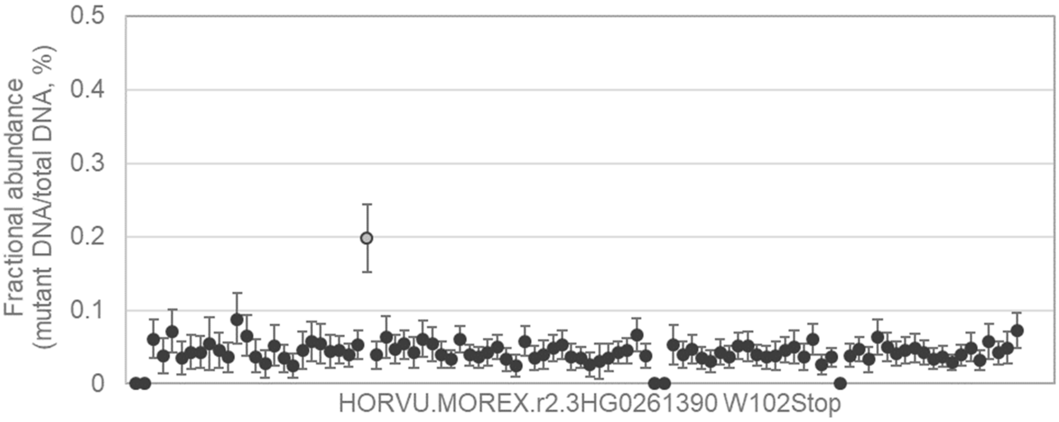


ATGGTCTGATGTTTCAGGATGATGTCAAGCTCATGGTTGACACTCACCTAGAGGCTTACAGATTTTCCA
TCTCCTG[G/A]TCAAGGCTTATACCAGGTATAAGCGCTTCTCGTTACCTCTCCATTGCACTATACTAGAA
AAACACAAAACTTG

---

62

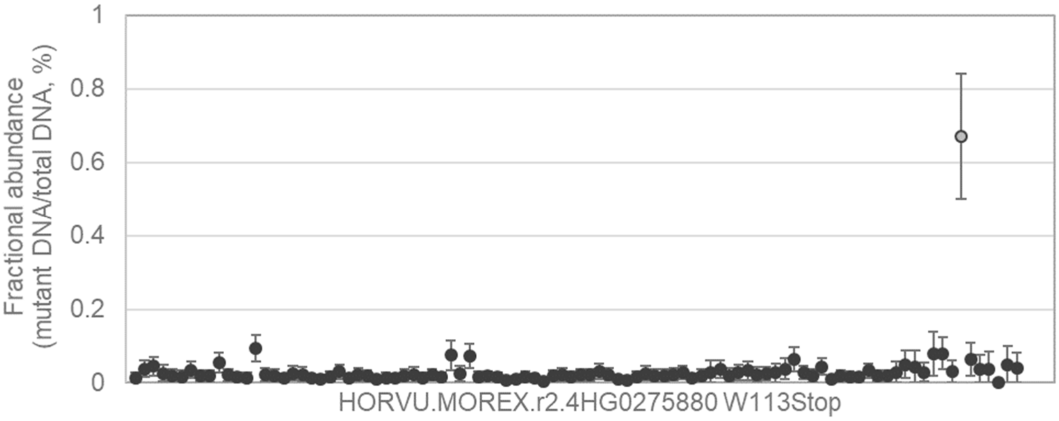


TGGGTGATGCATGGCATGGCATGATGATTAAACAGGTGTACCGTCTTTCGATCGGCGCGGCGTGCGG
TCTGTG[G/A]TGGCCGTCGGACAAGCTTGTGATCCAGGTGCTGGACGACTCGACGGACGCCGGCATC
CGGTCGCTGGTG

63
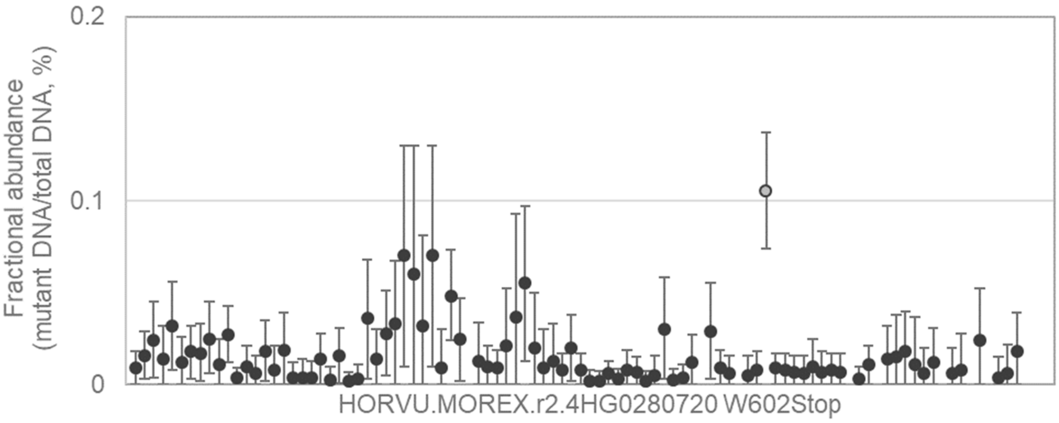


GCTGCTGGCCCTCTTCGCCGGCGGCGACGGGTACAAGGTGGAAGAGAAGGAAGGGTGCCTGACTCT
CGGGTG[G/A]CACACGCGCCCGCTGATCGCCACTTCCGCATGGCGCCTCGCCGCGCCGTGATCGCG
AGTTTTGAACGCTG

---

64
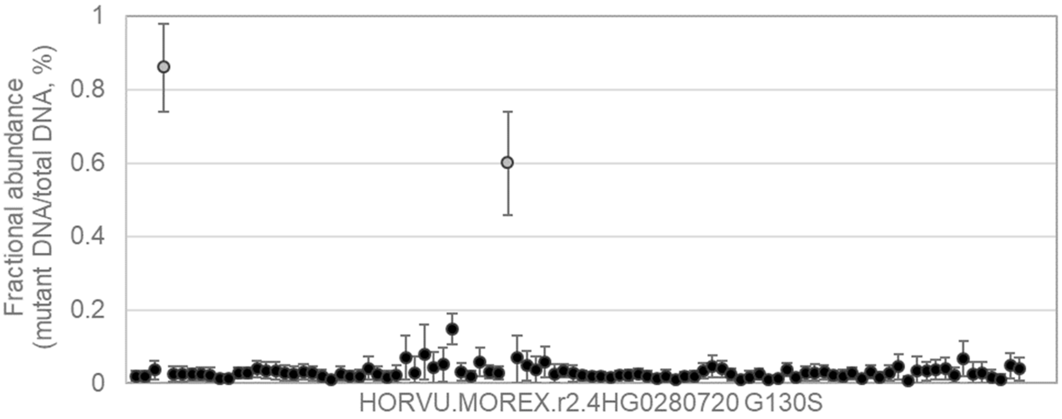


TCAACGCGCCGCCGCCGCCCCTCCCGCCGGCCCCGCCGCAGCTCAACGCCTCCACCTCTTCCACCG
TCACG[G/A]GCGGCGGCGGATACTTCGATCTCCCGCCCTCCGTCGACTCCTCCAGCAGCACCTACGC
CCTGCGCCCG

---

65
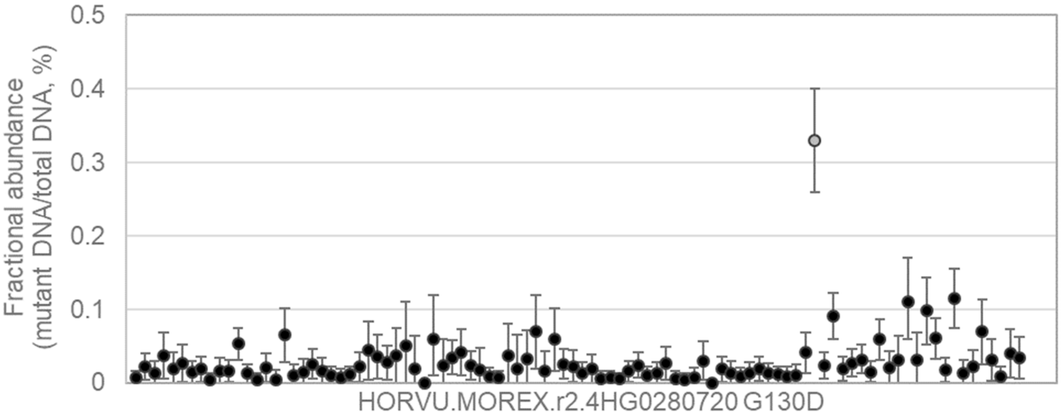


AACGCGCCGCCGCCGCCCCTCCCGCCGGCCCCGCCGCAGCTCAACGCCTCCACCTCTTCCACCGTC
ACGG[G/A]CGGCGGCGGATACTTCGATCTCCCGCCCTCCGTCGACTCCTCCAGCAGCACCTACGCCC
TGCGCCCGA

66 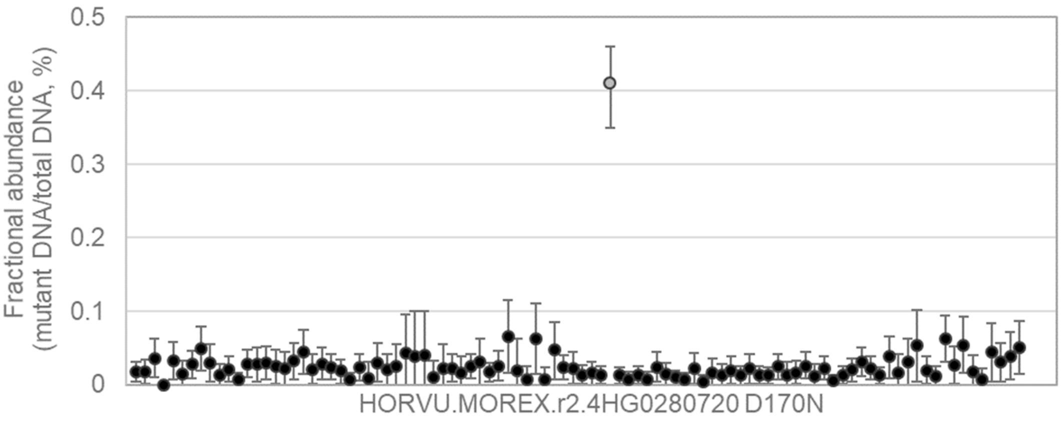


GCACCTACGCCCTGCGCCCGATCATCTCGCCGCCCGTCGCGCCGGCCGACCTCTCCGCTGACTCCG
TCCGG[G/A]ACCCCAAGCGGATGCGCACTGGCGGCAGCAGCACGTCGTCTTCGTCCTCCTCGTCGTC
CTCGCTCGGC

---

67 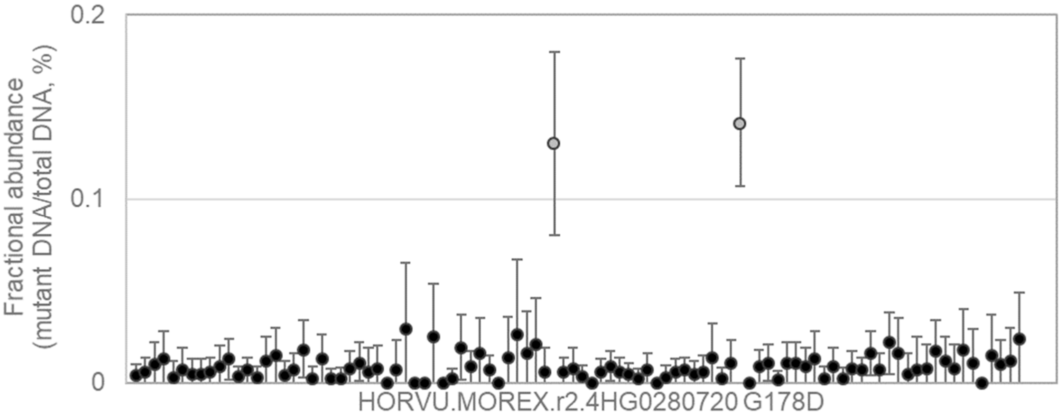


TCGCCGCCCGTCGCGCCGGCCGACCTCTCCGCTGACTCCGTCCGGGACCCCAAGCGGATGCGCACT
GGCG[G/A]CAGCAGCACGTCGTCTTCGTCCTCCTCGTCGTCCTCGCTCGGCGGTGGTGCCGCCAGGA
GCTCTGTGG

---

68 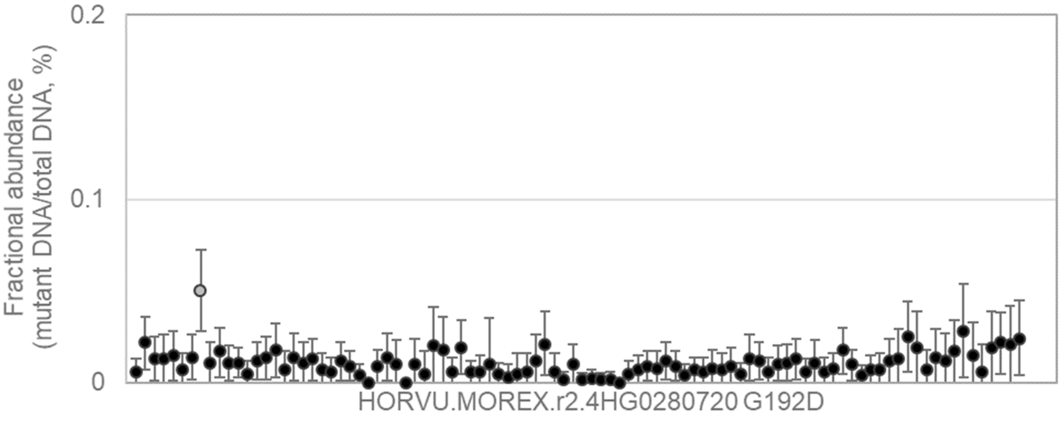


CCGGGACCCCAAGCGGATGCGCACTGGCGGCAGCAGCACGTCGTCTTCGTCCTCCTCGTCGTCCTC
GCTCG[G/A]CGGTGGTGCCGCCAGGAGCTCTGTGGTGGAGGCTGCTCCGCCGGTGGCGGCTGCGGC
TGCTGCGCC

69

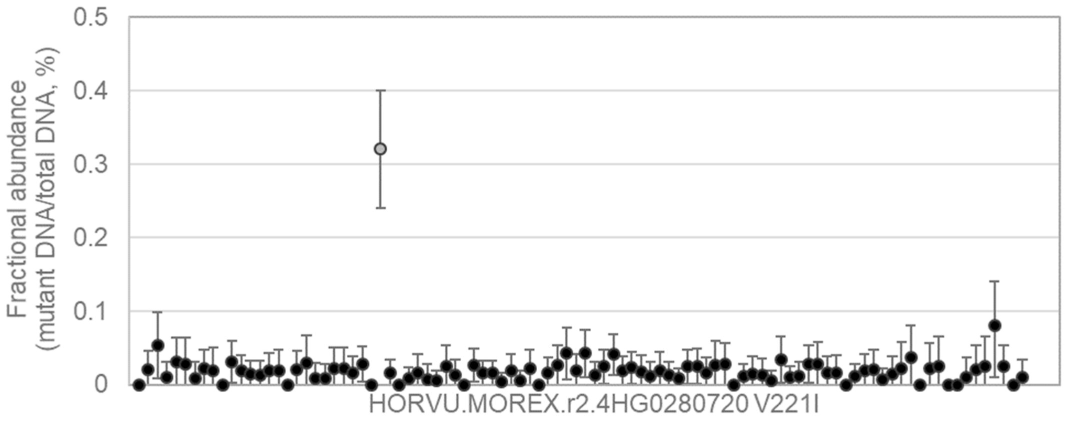


GAGCTCTGTGGTGGAGGCTGCTCCGCCGGTGGCGGCTGCGGCTGCTGCGCCCGCGCTGCCGGTCG
TCGTG[G/A]TCGACACGCAGGAGGCCGGGATTCGGCTGGTGCACGCGCTGCTGGCGTGCGCGGAGG
CCGTGCAGC

---

70

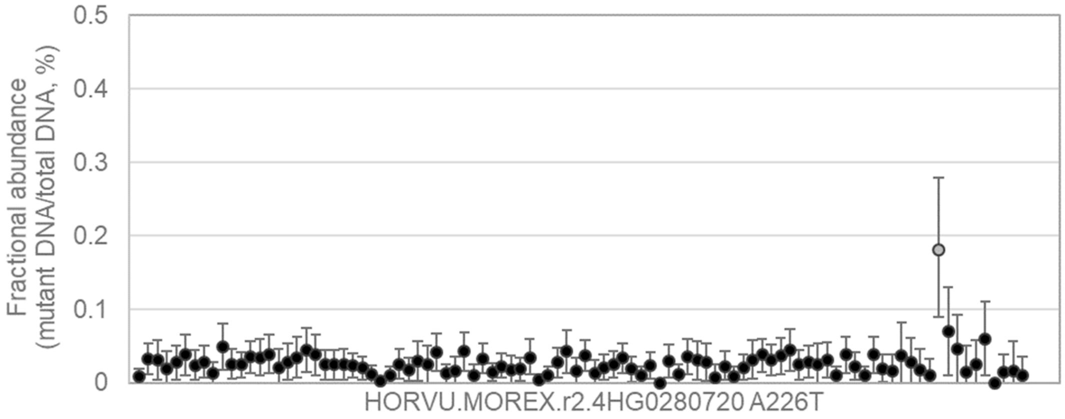


GCTGCTCCGCCGGTGGCGGCTGCGGCTGCTGCGCCCGCGCTGCCGGTCGTCGTGGTCGACACGCA
GGAG[G/A]CCGGGATTCGGCTGGTGCACGCGCTGCTGGCGTGCGCGGAGGCCGTGCAGCAGGAGAA
CCTCTCGGC

---

71

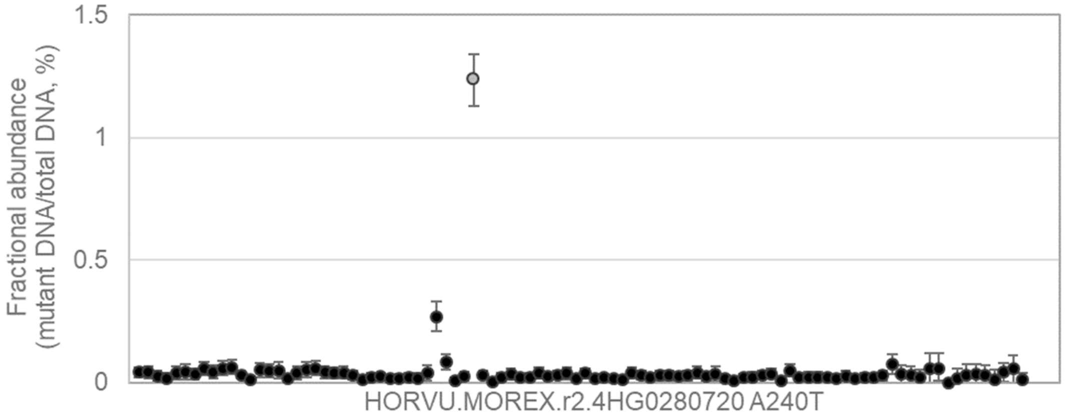


GCCGGTCGTCGTGGTCGACACGCAGGAGGCCGGGATTCGGCTGGTGCACGCGCTGCTGGCGTGCG
CGGAG[G/A]CCGTGCAGCAGGAGAACCTCTCGGCCGCCGAGGCGCTGGTGAAGCAGATACCCTTGCT
GGCAGCGTC

72 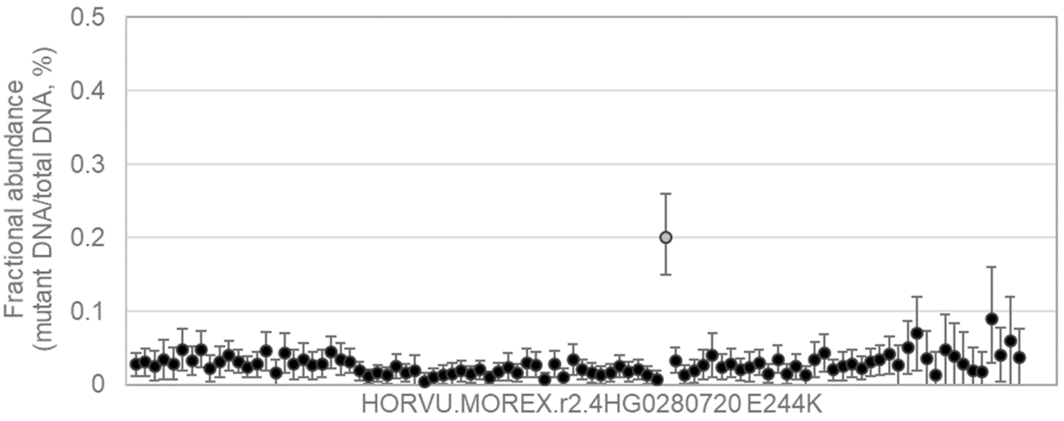


GTCGACACGCAGGAGGCCGGGATTCGGCTGGTGCACGCGCTGCTGGCGTGCGCGGAGGCCGTGCA
GCAG[G/A]AGAACCTCTCGGCCGCCGAGGCGCTGGTGAAGCAGATACCCTTGCTGGCAGCGTCGCAG
GGCGGCGC

73 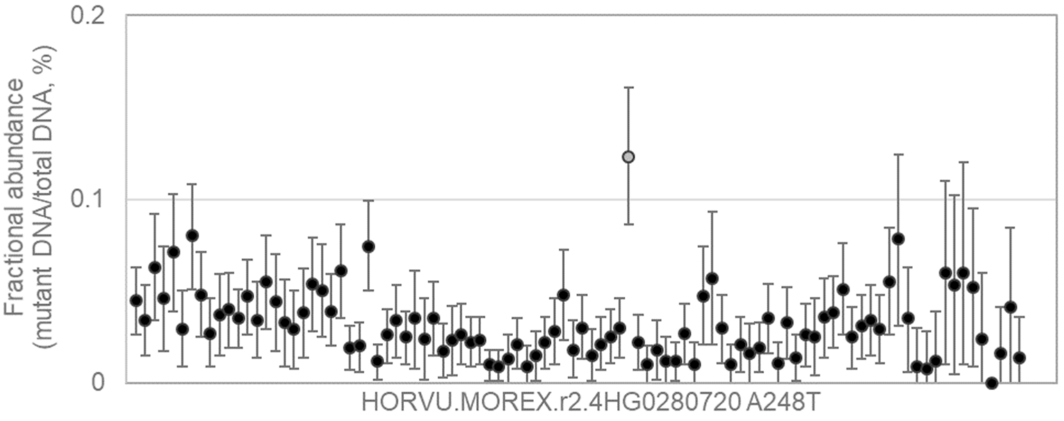


GGAGGCCGGGATTCGGCTGGTGCACGCGCTGCTGGCGTGCGCGGAGGCCGTGCAGCAGGAGAACC
TCTCG[G/A]CCGCCGAGGCGCTGGTGAAGCAGATACCCTTGCTGGCAGCGTCGCAGGGCGGCGCGA
TGCGCAAGGT

74 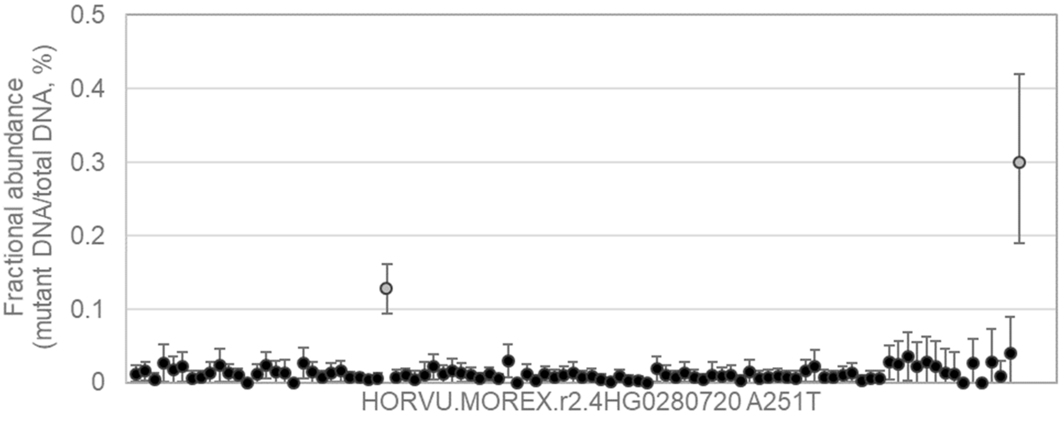


GATTCGGCTGGTGCACGCGCTGCTGGCGTGCGCGGAGGCCGTGCAGCAGGAGAACCTCTCGGCCG
CCGAG[G/A]CGCTGGTGAAGCAGATACCCTTGCTGGCAGCGTCGCAGGGCGGCGCGATGCGCAAGG
TCGCCGCCTA

75

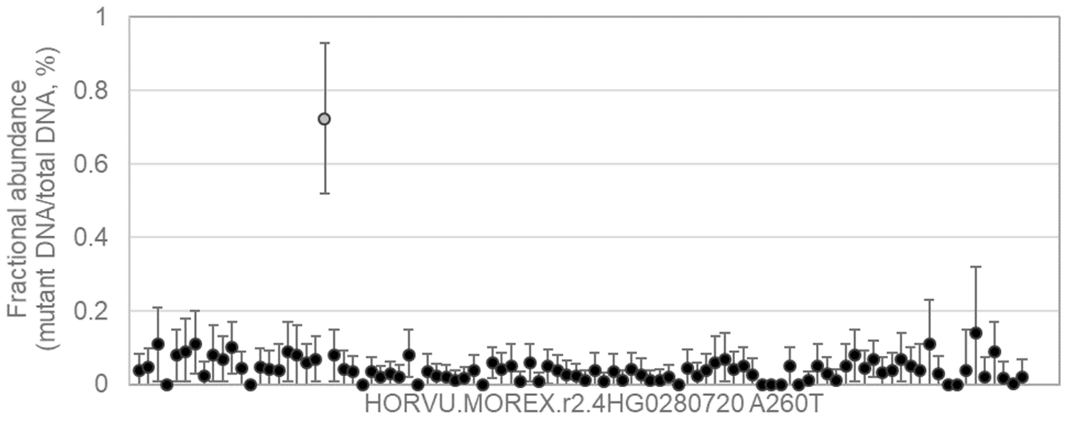


GTGCGCGGAGGCCGTGCAGCAGGAGAACCTCTCGGCCGCCGAGGCGCTGGTGAAGCAGATACCCTT
GCTG[G/A]CAGCGTCGCAGGGCGGCGCGATGCGCAAGGTCGCCGCCTACTTCGGCGAGGCCCTCGC
CCGCCGCG

---

76

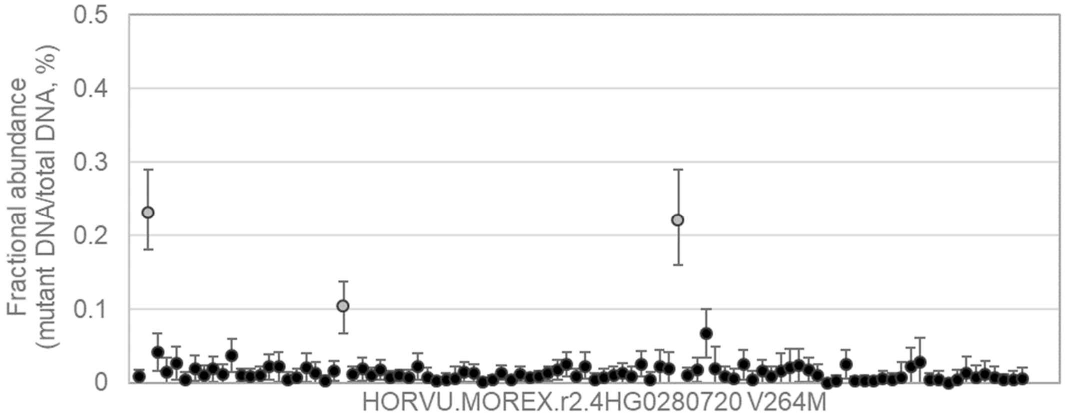


GCGGCTGCGGCTGCTGCGCCCGCGCTGCCGGTCGTCGTGGTCGACACGCAGGAGGCCGGGATTCG
GCTG[G/A]TGCACGCGCTGCTGGCGTGCGCGGAGGCCGTGCAGCAGGAGAACCTCTCGGCCGCCGA
GGCGCTGG

---

77

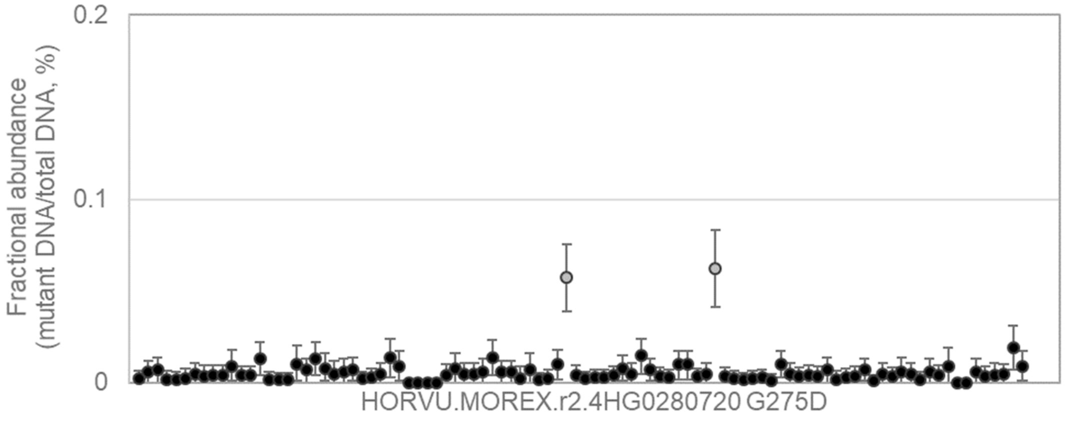


CTGGTGAAGCAGATACCCTTGCTGGCAGCGTCGCAGGGCGGCGCGATGCGCAAGGTCGCCGCCTAC
TTCG[G/A]CGAGGCCCTCGCCCGCCGCGTCTTCCGCTTCCGCCCGCAGCCGGACAGCTCCCTCCTCG
ACGCCGCC

78

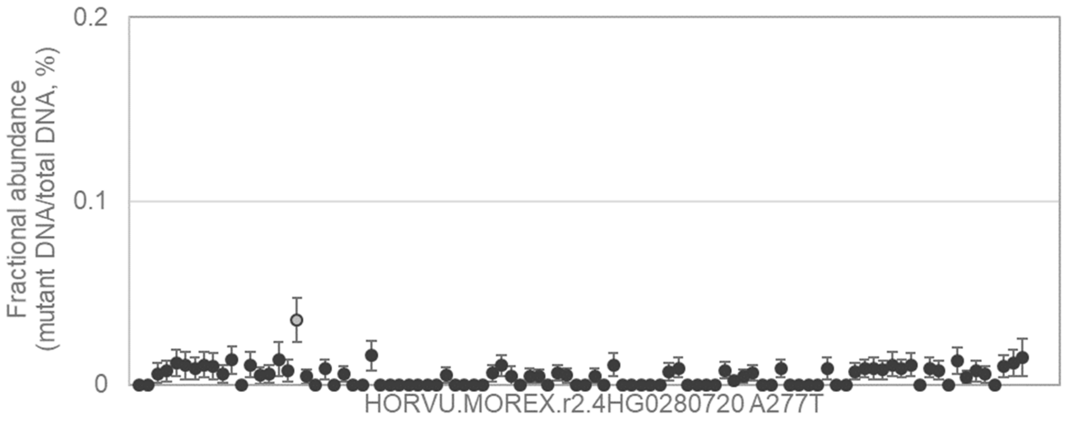


TGAAGCAGATACCCTTGCTGGCAGCGTCGCAGGGCGGCGCGATGCGCAAGGTCGCCGCCTACTTCG
GCG[G/A]CCCTCGCCCGCCGCGTCTTCCGCTTCCGCCCGCAGCCGGACAGCTCCCTCCTCGACGCCG
CCTTCG

---

79

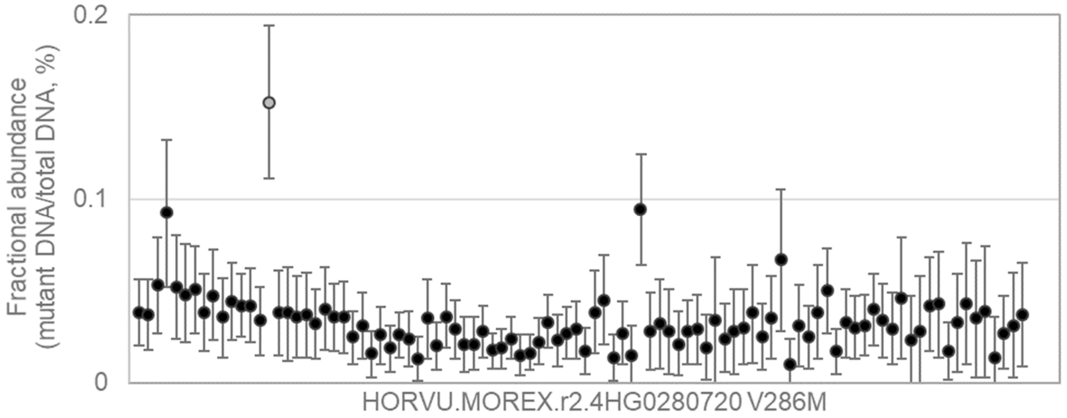


CTGGTGCACGCGCTGCTGGCGTGCGCGGAGGCCGTGCAGCAGGAGAACCTCTCGGCCGCCGAGGC
GCTG[G/A]TGAAGCAGATACCCTTGCTGGCAGCGTCGCAGGGCGGCGCGATGCGCAAGGTCGCCGC
CTACTTCGG

---

80

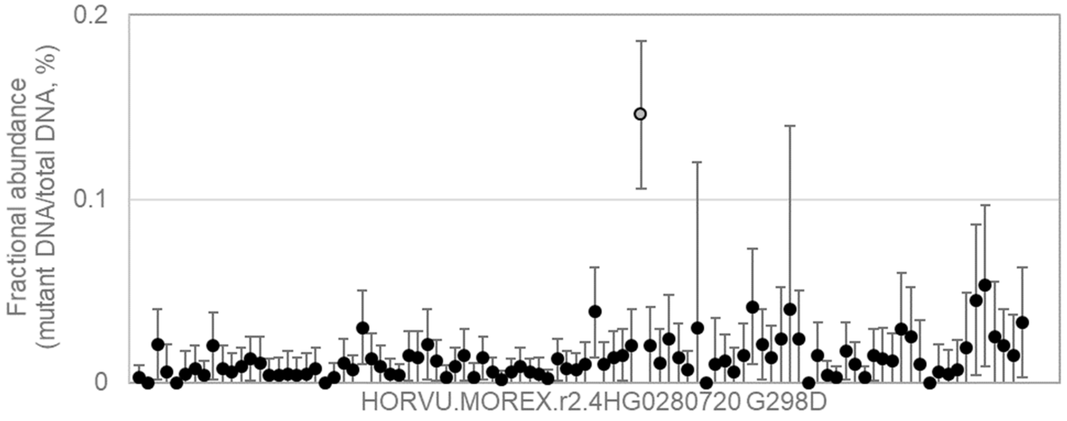


CAGCAGGAGAACCTCTCGGCCGCCGAGGCGCTGGTGAAGCAGATACCCTTGCTGGCAGCGTCGCAG
GGCG[G/A]CGCGATGCGCAAGGTCGCCGCCTACTTCGGCGAGGCCCTCGCCCGCCGCGTCTTCCGC
TTCCGCCCG

81

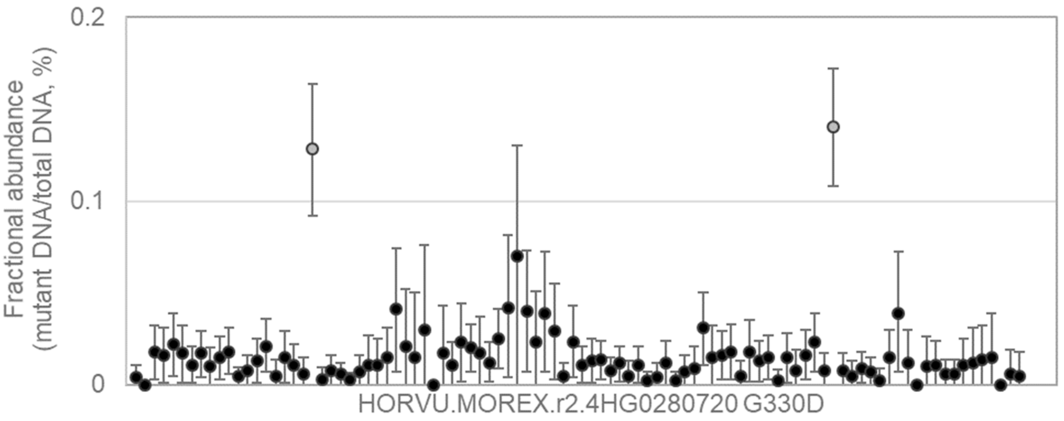


TCTACGAGTCCTGCCCCTACCTCAAGTTCGCCCATTTCACCGCCAACCAGGCCATCCTGGAGGCGTT
CGCCG[G/A]CTGCCGCCGCGTCCACGTCGTCGACTTCGGCATCAAGCAGGGGATGCAGTGGCCGGC
CCTTCTCCAG

---

82

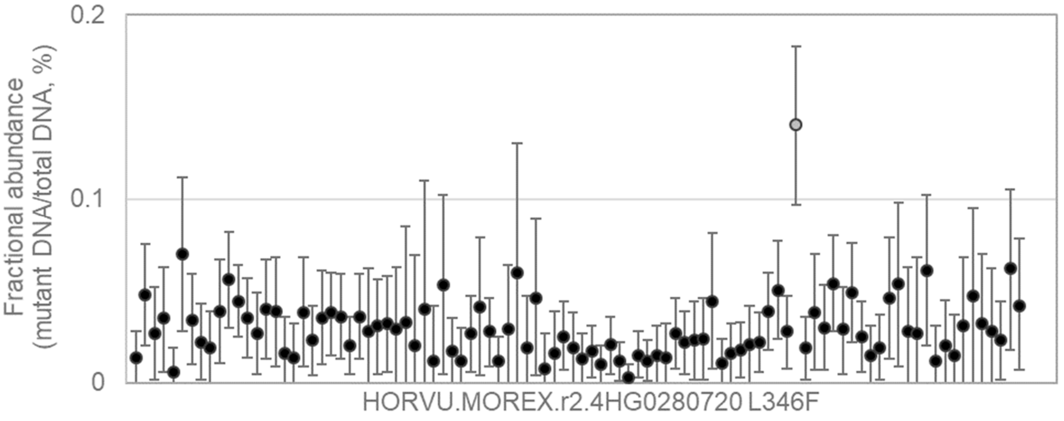


CGGACAGCTCCCTCCTCGACGCCGCCTTCGCCGACCTCCTCCACGCGCACTTCTACGAGTCCTGCCC
CTAC[C/T]TCAAGTTCGCCCATTTCACCGCCAACCAGGCCATCCTGGAGGCGTTCGCCGGCTGCCGCC
GCGTCCAC

---

83

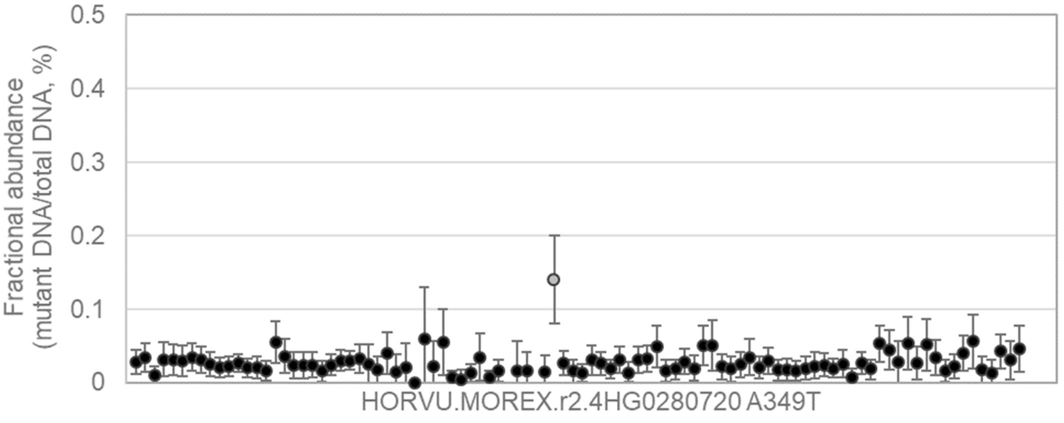


GGAGGCGTTCGCCGGCTGCCGCCGCGTCCACGTCGTCGACTTCGGCATCAAGCAGGGGATGCAGTG
GCCG[G/A]CCCTTCTCCAGGCCCTCGCACTCCGTCCCGGCGGGCCCCCTTCGTTCCGCCTCACCGGC
GTTGGCCCC

84

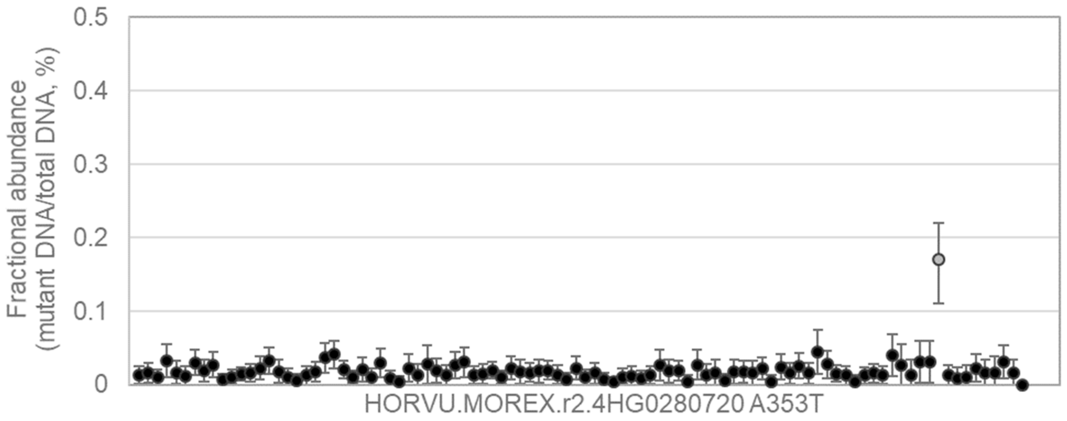


CCGGCTGCCGCCGCGTCCACGTCGTCGACTTCGGCATCAAGCAGGGGATGCAGTGGCCGGCCCTTC
TCCAG[G/A]CCCTCGCACTCCGTCCCGGCGGGCCCCCTTCGTTCCGCCTCACCGGCGTTGGCCCCCC
GCAGCCGGA

---

85

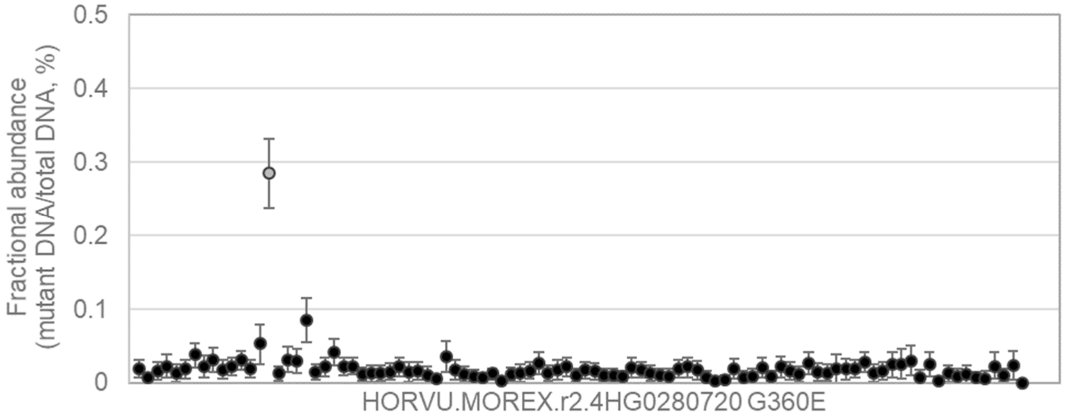


CGTCGACTTCGGCATCAAGCAGGGGATGCAGTGGCCGGCCCTTCTCCAGGCCCTCGCACTCCGTCC
CGGCG[G/A]GCCCCCTTCGTTCCGCCTCACCGGCGTTGGCCCCCCGCAGCCGGACGAGACCGACGC
CCTGCAGCAG

---

86

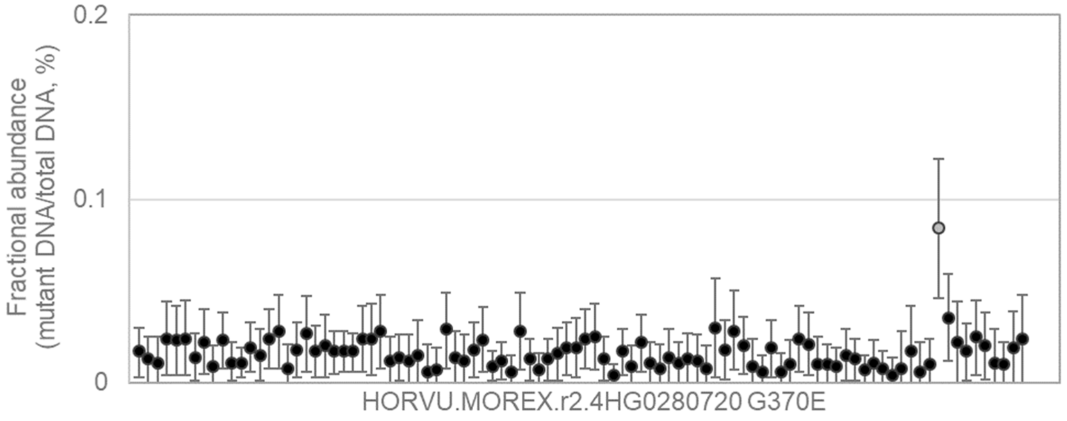


GTGGCCGGCCCTTCTCCAGGCCCTCGCACTCCGTCCCGGCGGGCCCCCTTCGTTCCGCCTCACCGG
CGTTG[G/A]CCCCCCGCAGCCGGACGAGACCGACGCCCTGCAGCAGGTGGGCTGGAAGCTCGCCCA
GTTCGCGCAC

87

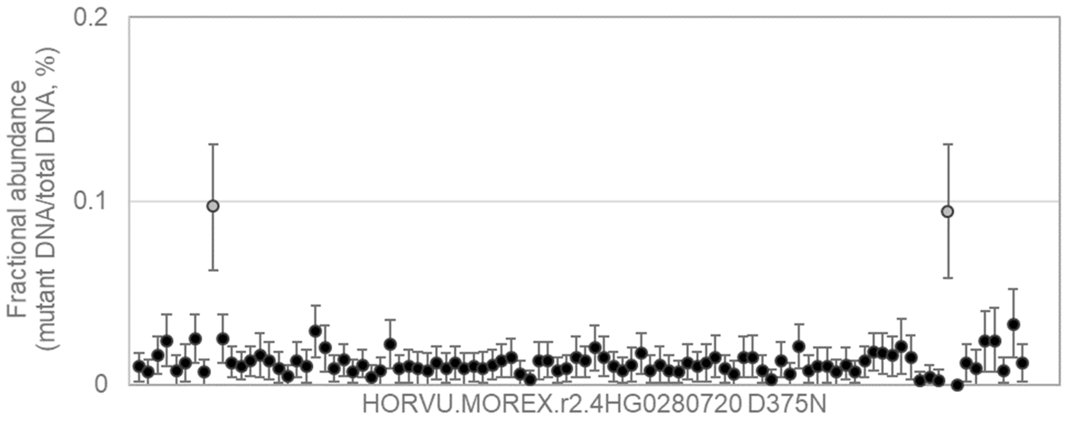


CCAGGCCCTCGCACTCCGTCCCGGCGGGCCCCCTTCGTTCCGCCTCACCGGCGTTGGCCCCCCGCA
GCCG[G/A]ACGAGACCGACGCCCTGCAGCAGGTGGGCTGGAAGCTCGCCCAGTTCGCGCACACCATC
CGCGTCGA

88

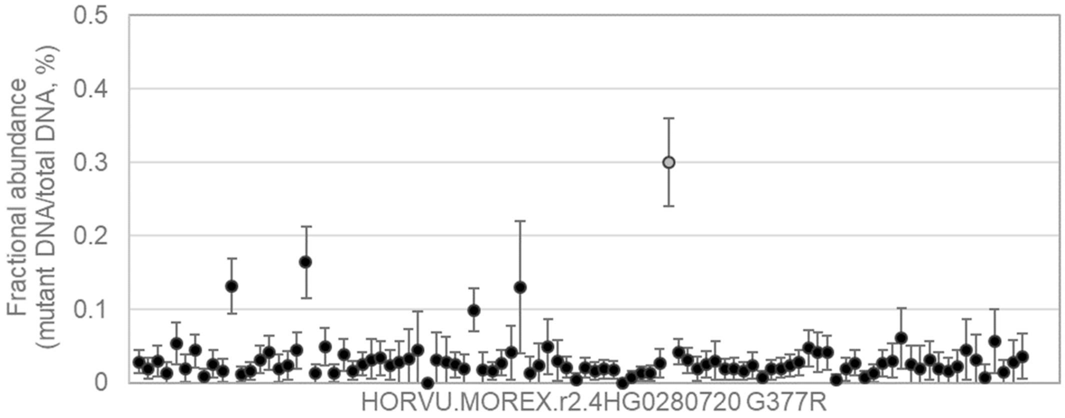


CCAACCAGGCCATCCTGGAGGCGTTCGCCGGCTGCCGCCGCGTCCACGTCGTCGACTTCGGCATCA
AGCAG[G/A]GGATGCAGTGGCCGGCCCTTCTCCAGGCCCTCGCACTCCGTCCCGGCGGGCCCCCTTC
GTTCCGCCT

89

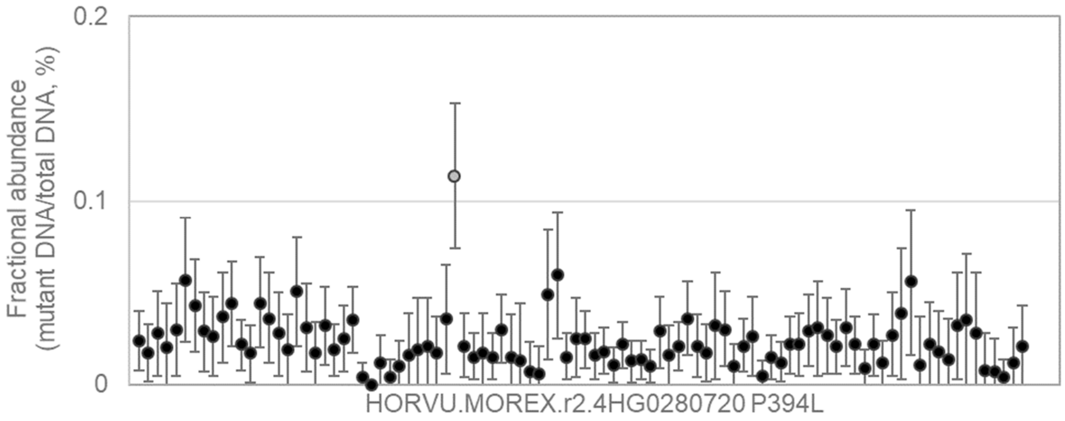


GACTTCGGCATCAAGCAGGGGATGCAGTGGCCGGCCCTTCTCCAGGCCCTCGCACTCCGTCCCGGC
GGGC[C/T]CCCTTCGTTCCGCCTCACCGGCGTTGGCCCCCCGCAGCCGGACGAGACCGACGCCCTG
CAGCAGGTG

90

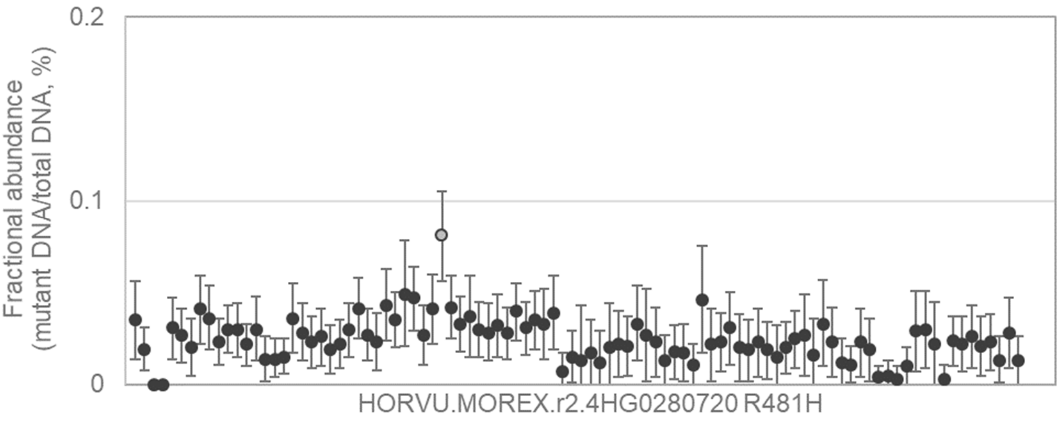


CGCCGTGCGGCCGAGGATCGTCACCGTGGTCGAGCAGGAGGCGAACCACAACTCCGGCTCATTCCT
GGA[G/A]CTTCACCGAGTCCCTGCACTACTACTCCACCATGTTCGATTCTCTCGAGGGCGGCAGCTCC
GGCGGC

---

91

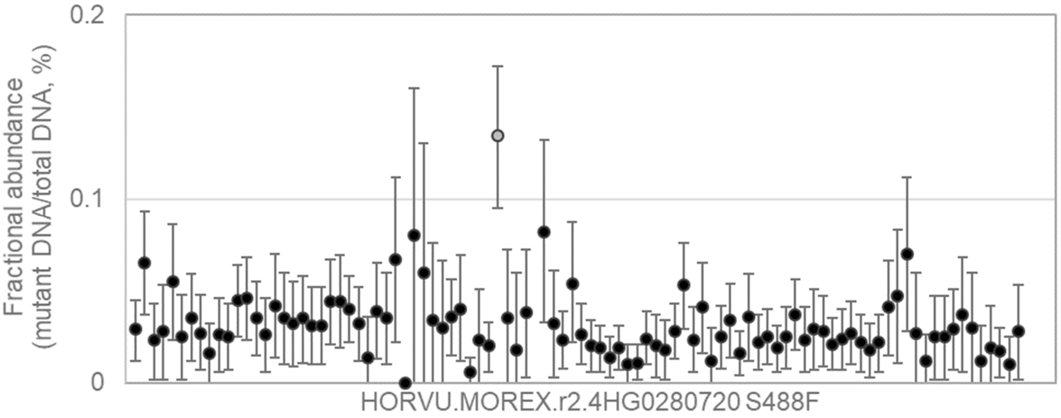


GAGGATCGTCACCGTGGTCGAGCAGGAGGCGAACCACAACTCCGGCTCATTCCTGGACCGCTTCACC
GAGT[C/T]CCTGCACTACTACTCCACCATGTTCGATTCTCTCGAGGGCGGCAGCTCCGGCGGCCCGTC
CGAGGTCT

---

92

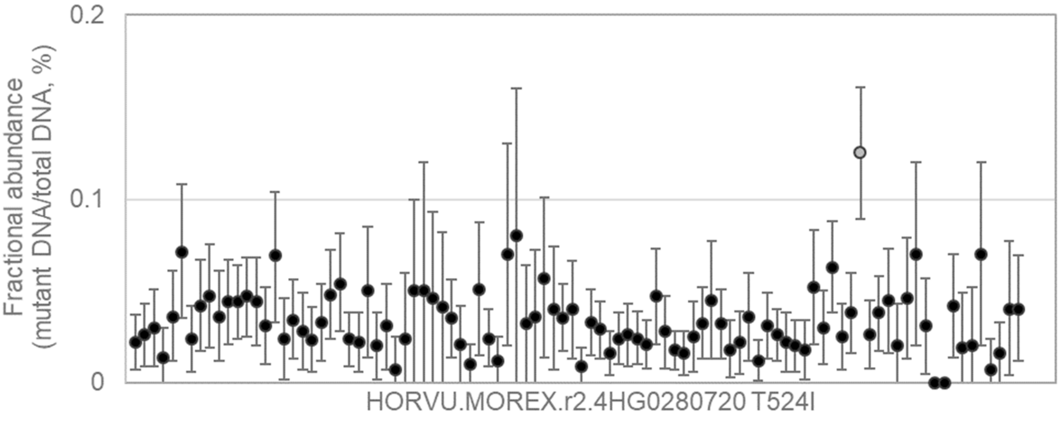


TCGAGCAGGAGGCGAACCACAACTCCGGCTCATTCCTGGACCGCTTCACCGAGTCCCTGCACTACTA
CTCCA[C/T]CATGTTCGATTCTCTCGAGGGCGGCAGCTCCGGCGGCCCGTCCGAGGTCTCATCGGGG
GGTGCCGCT

93 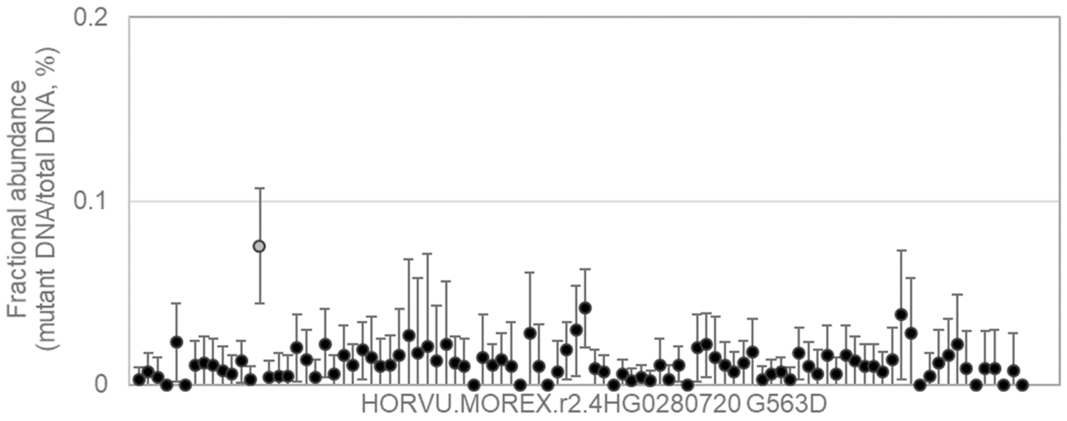


GTCTCATCGGGGGGTGCCGCTCCTGCCGCCGCCGCCGGCACGGACCAGGTCATGTCCGAGGTGTAC
CTCG[G/A]CCGGCAGATCTGCAACGTGGTGGCCTGCGAGGGCACGGAGCGCACAGAGCGGCACGAG
ACACTGGG

94 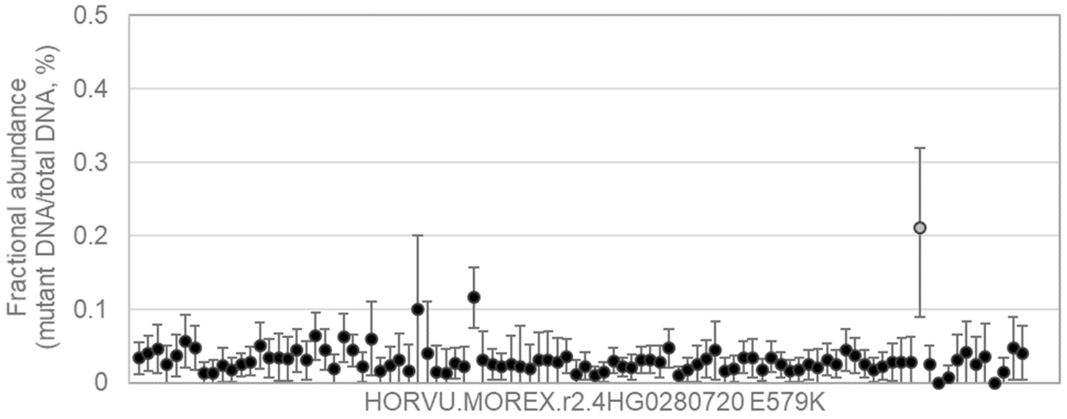


GGTCATGTCCGAGGTGTACCTCGGCCGGCAGATCTGCAACGTGGTGGCCTGCGAGGGCACGGAGCG
CACA[G/A]AGCGGCACGAGACACTGGGGCAGTGGCGGAACCGGCTGGGCAACGCCGGGTTCGAGAC
CGTGCACC

95 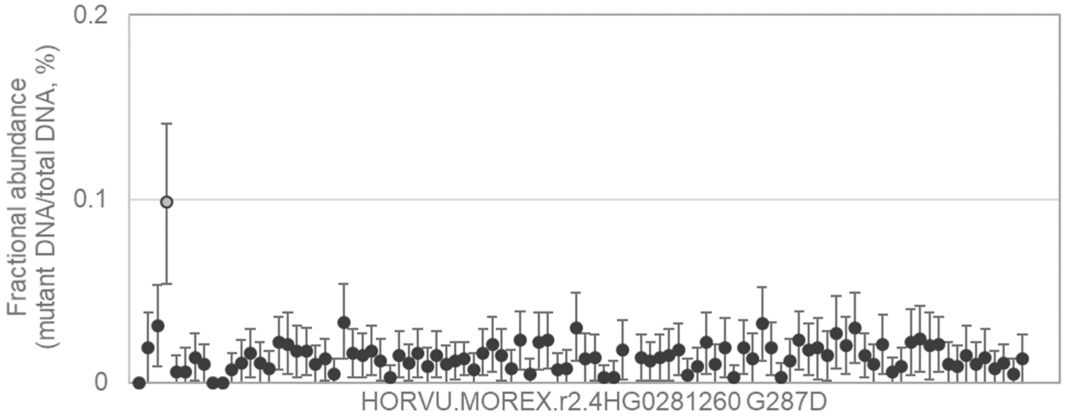


AGGTGATCAAGAGGGCGATCAAGAAGCTCGAGGCGCGGCACACGGAGCACATAGCCGCCTACGGGG
AAG[G/A]CAACGAGCGCCGGCTCACCGGCCGCCACGAGACCGCCGACATCAACACCTTCGTATGGGT
AAGTAAT

96

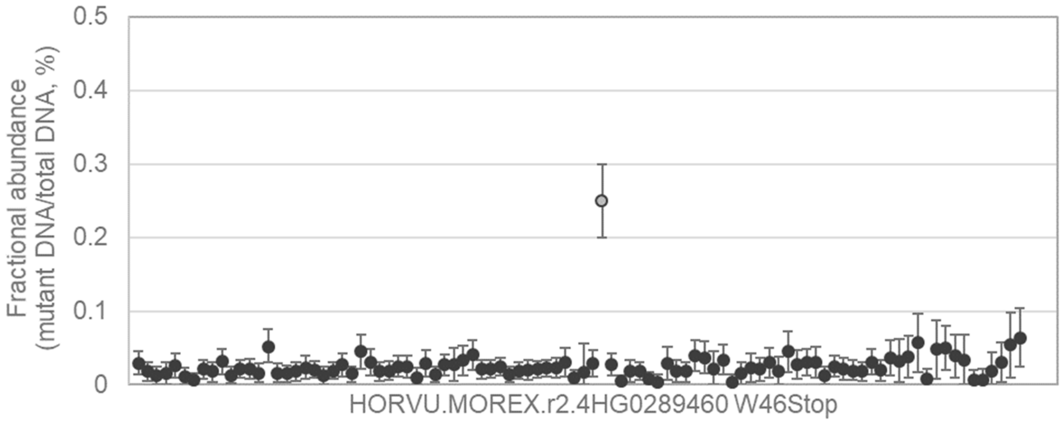


GAGGCCGGGAACGGCCGGGAGTCCGACTGGCTTGACGACGACGGGCGGCCGCGTCGGTCGGGCAC
GGTGTG[G/A]ACGGCGAGCGCCCACATCATCACCGCCGTCATCGGCTCCGGGGTCCTCTCGCTGGCC
TGGGCCATCGCG

---

97

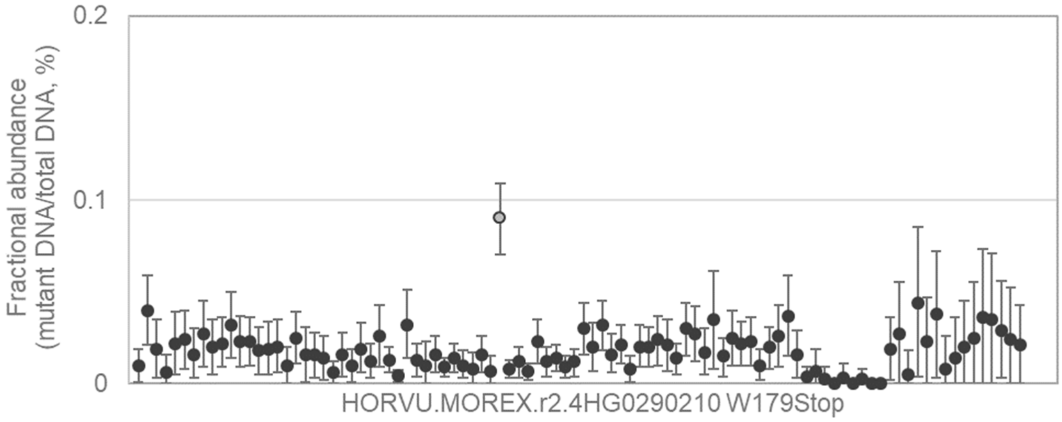


GTCAAATCAGATAAAGATTTGACATATTTCTCTGACAGTGCTGAAGATTAATGAGGTTTACCTGTTTGCA
GAAGTG[G/A]ACAATTGATATGAAAGGGAATCTAAGCCCTTCAATGAGATCAAGCTTGTGGAAGTCCAA
ACGAGTGTTCTTT

---

98

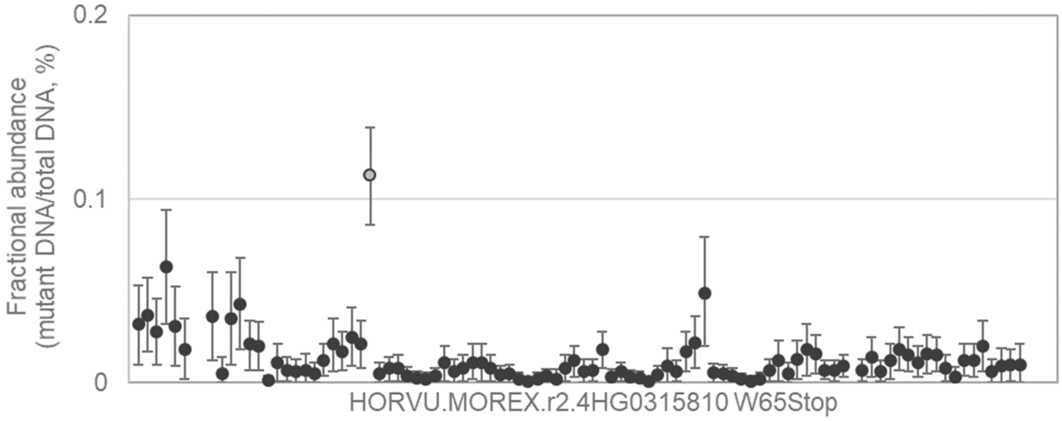


TTAATAATCGTCCGTTATTTTTGTTGGGGCCTGTTAATTGATTGCAGCCAAGGCTAGCTACGGGGACA
GGGAGTG[G/A]TACTTCTTCACGCCGAGGGACCGTAAGTACCCCAACGGTGTCCGGCCGAACCGTGC
TGCGGGTTCCGGCT

99

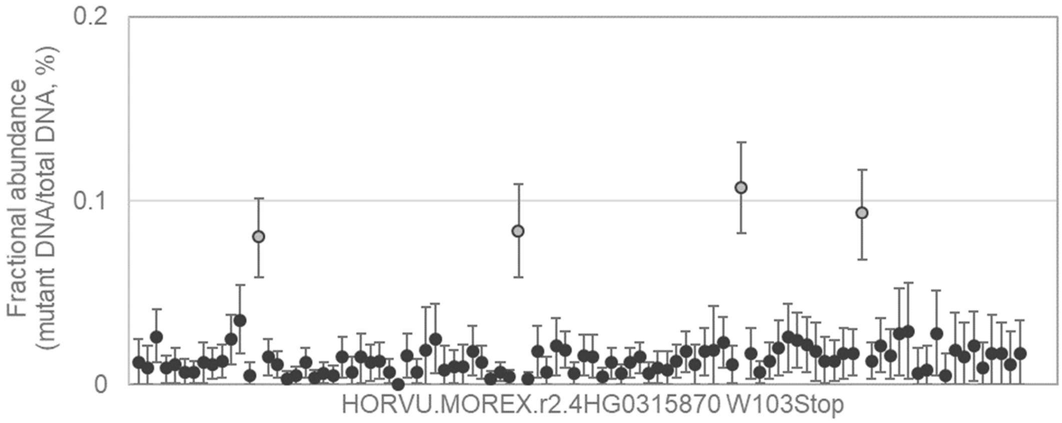


TACTTCTTTTGCCATAAAGACCGCAAGTACCCGACGGGGACGAGGACCAACCGCGCCACCATGAGCG
GCTACTG[G/A]AAGGCCACCGGCAAGGACAAGGAGATCTTCCGGGGGAGGGGCATCCTCGTCGGAAT
GAAGAAGACGCTC

---

100

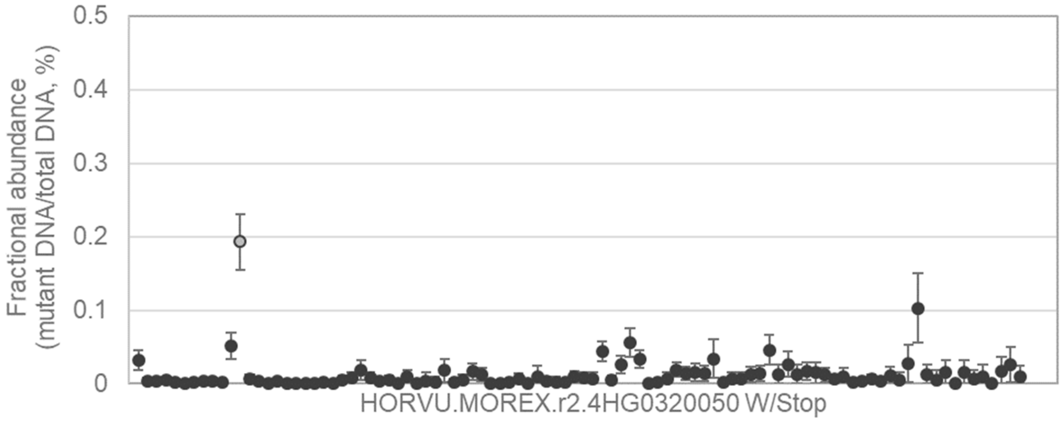


GCGGCGGCGACCGCGGCGCTGCGGACGGCGGTGAGCGTGCCGGAGACGTGCGACCTGTACCGCG
GCAGCTG[G/A]GTGTACGACGAGGTGAACGCGCCGGTGTACAAGGAGACGCAGTGCGAATTCCTGAC
GGAGCAGGTGACCT

---

101

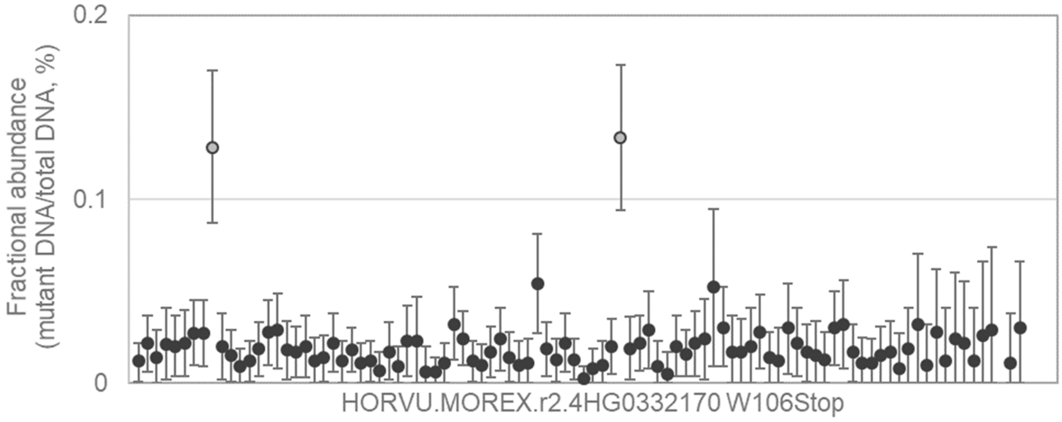


TAAGCTCGAGACGCCGCGCAAGGTGCAGCTCGCCGCGGAGCTGGGCCTCGACACCAAGCAGGTCGC
CGTGTG[G/A]TTCCAGAACCGCCGCGCCCGCTACAAGAGCAAGCTCATCGAGGAGGAGTTCAACAAG
CTCCGGGCCGCCC

102

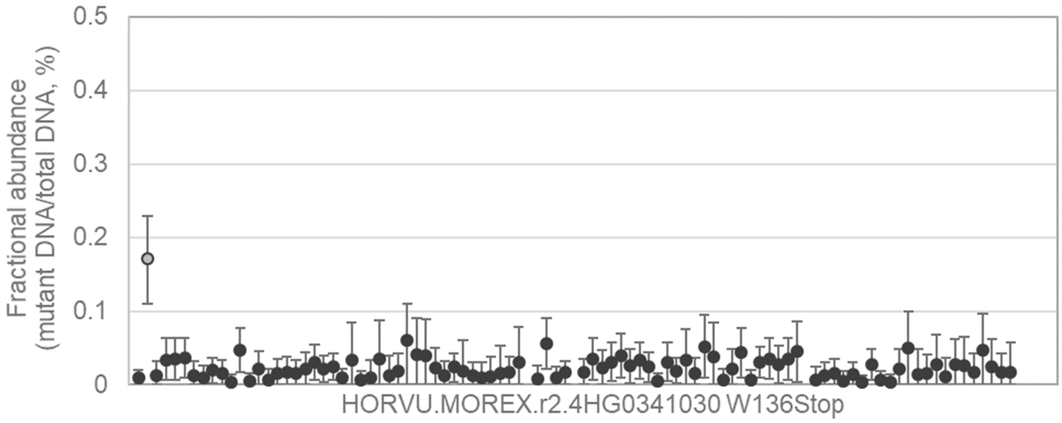


AGCGGGTCATCGGCCTCAAGAAGACGCTCGTCTTCTACCAGGGCCGCGCTCCGCGGGGCACCAAGA
CGGACTG[G/A]GTCATGAACGAGTACCGCCTCCCCGACTCCGGCCCGGCGCCACCCCAGGAGGACAC
GGTGCTGTGCAAG

---

103

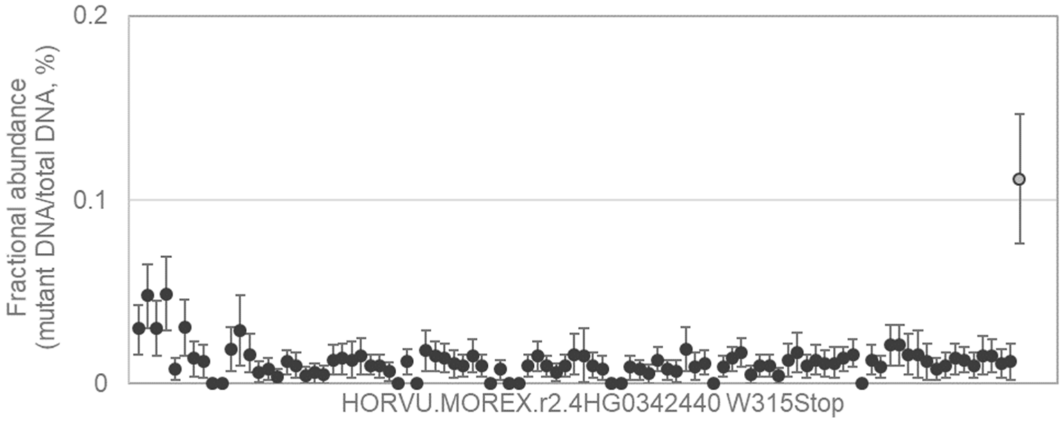


GTATTTGTTACAGGTGCGGGAGATGCCAAATGTTCTAATTCTTCCTCGAAGCGCAGACTATAGTGAAG
AAGTCTG[G/A]ATGGAGATAAGAGAGAAAGCAATCACGATACTACAGTCCTTTTTCTTTGATGGTGTTGT
TCCAAGTAGTTCAA

---

104

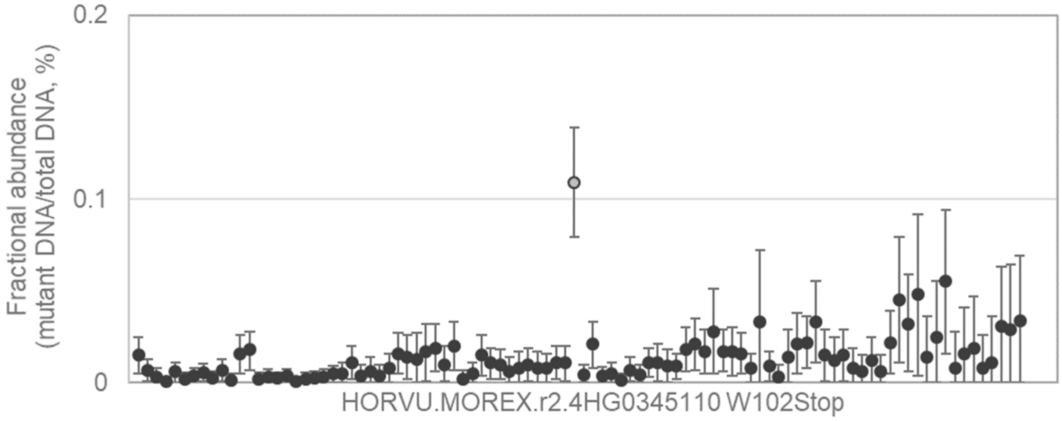


GGAGCAAGAGCACCGCTGCGGTGGCGCTCAAGGGCCTGCAGTTTGTGACTGCCAAAGTTGGCGGCG
ACGGCTG[G/A]GCCGCTGTGGAGAAGCGGTTCAACCACCTGCAGGTTGACGGCGTCCTGCTCCGTTC
CAGATTTGGGAAAT

105 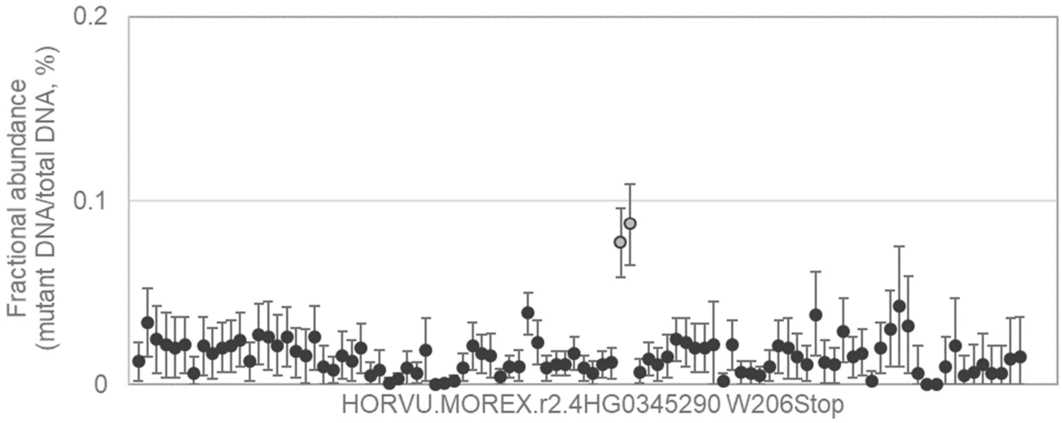


ATTCATAGTCTTGTGCACTGACAGTCCGCTAATTCTGAGAATATTTTTGCAAAATTTCAGCTGGCATATA
TGTTCTG[G/A]ATTCTTGGTTACAATCTTAATGAAGGACATCAACTGCTACAGGTAGTATTAGGAAGTTG
AAAGCTGTTTGTAA

---

106 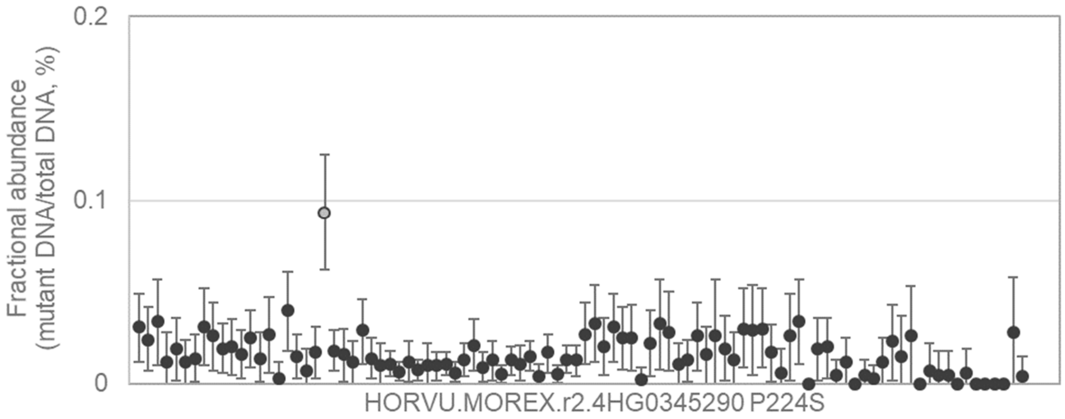


TACCACATGGCTAGTTATCTTGCTTGTGTTTTGTACAGATTTTGAATCCATTTTTAATGTTGTAGAGTAA
AAGG[C/T]CTAGCTTTCCGAAGTTGGAAGCCATCAAATTGGCAACAGCTGACATTGTAAGTAAGACGTA
CGCTGTAT

---

107 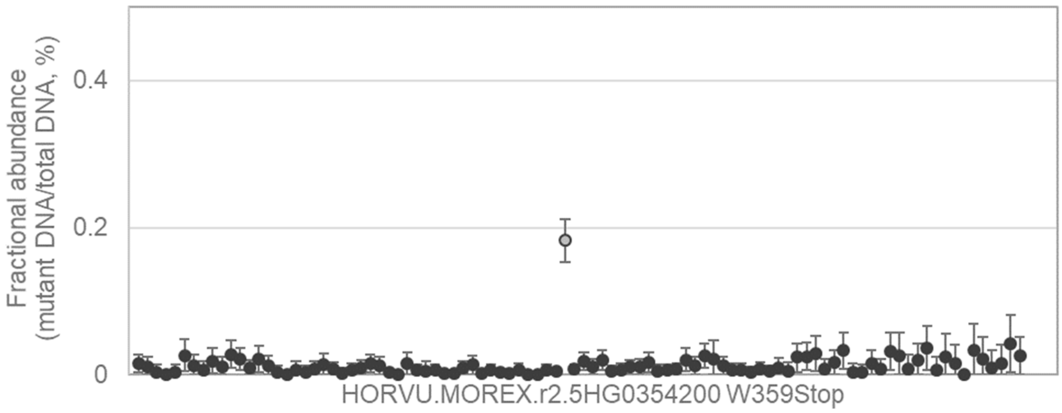


ATGCATTACTACTCCCAATCCTTATTTGTTGCGCCATTCTGTCATCCTTAGTGTTGGACACTGCTGCTG
GAGTTTG[G/A]TGCGACACAAAATCAGTAGTTACAACTCCAAGGACAGGAAGATATAGTGCAGATGCAG
CTGGTGGTGAGGC

108

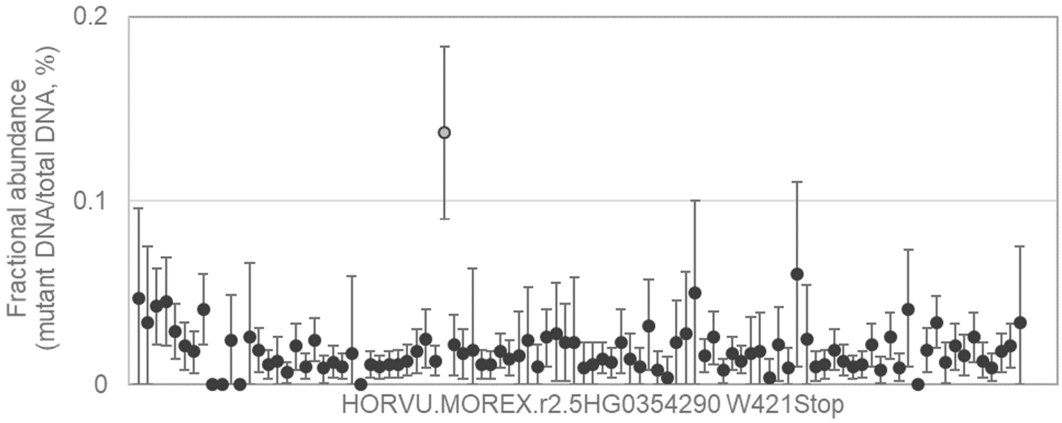


TACGCCAAGGACTCGTCGGAGATGATGTGGGTGGACGTGCCGGACTGCTTCTGTTTCCACCTCTGGA
ACTCGTG[G/A]GAGGAGCCGGAGACGGACGAGGTGGTGGTGATCGGCTCCTGCATGACCCCCGCAG
ACTCCATCTTCAACG

---

109

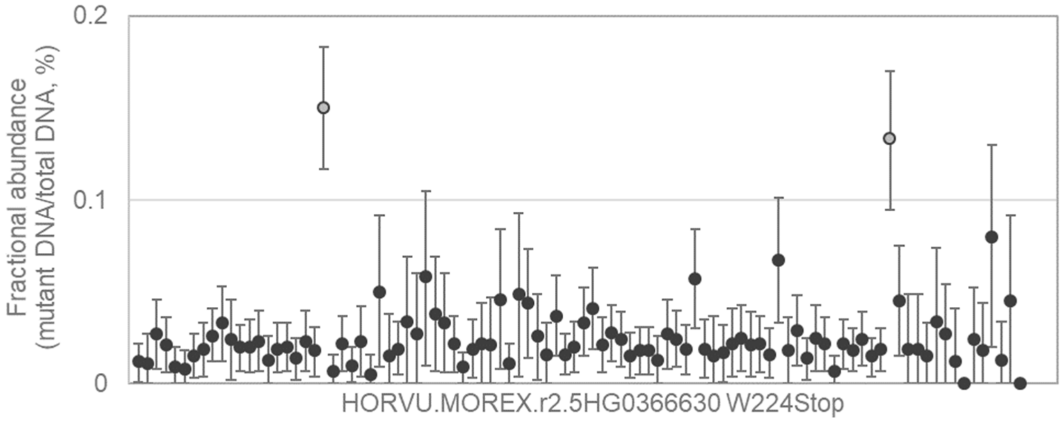


GTTCGCTATCGTGTTGTCATTTGGCCTTTTGCTCACCGCGCTGCGGAGCAGGAAAGCGCGATCCTGG
CGCTATG[G/A]GGCAGGTACATCCACTGTCAATTGTAACTGGTCCTGCAGATGCTATGAGAATTTTATT
GCCTCAAATCTCCTC

---

110

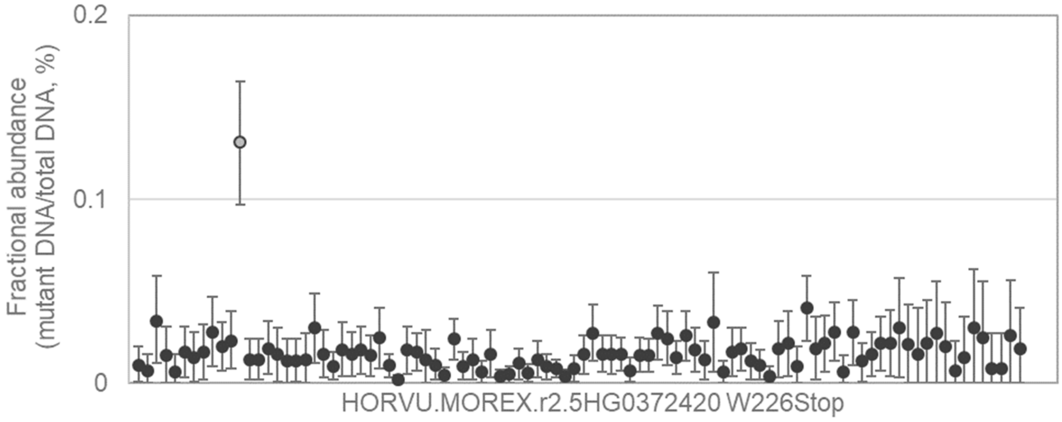


CCACCAAGGTGCTTCTGTATCTCCACAATGTGATAACAGCCTGAGTACACAGAATTATGGTGTGAGGC
CAAGCTG[G/A]AATTCTCGTCGTGGACCACGCGGTAATTTTGTCCCCCCTATCCGAAACAATGGAGGAT
CTACTTGCAATGCA

111

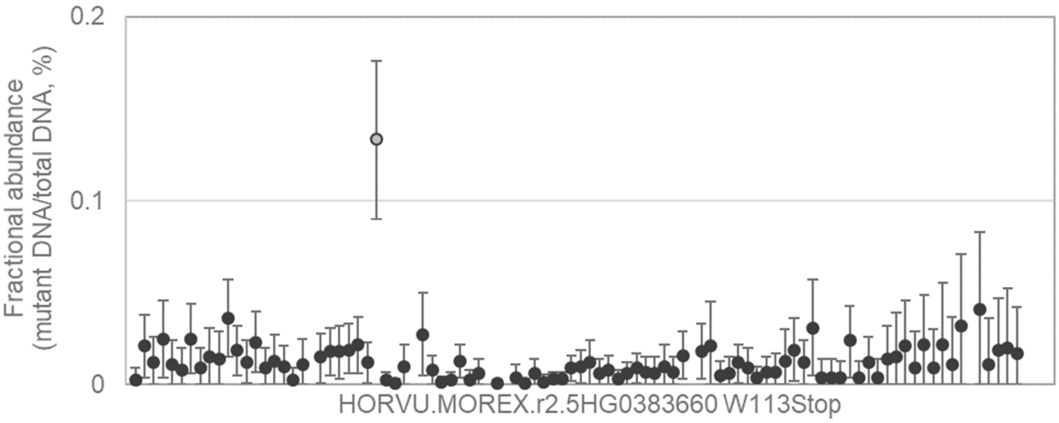


GCAGGCAGGCCGCGTCCATGCCCAGCCCCGTGCCCATCATCGCCGAGGTCAACATCTACAAGTGCA
ACCCATG[G/A]GACCTCCCCGGTATATACATACATAACTAAAAACTAGAGCATCGACGCCATCCTTCTT
GGTTCAACTCTGCTC

---

112

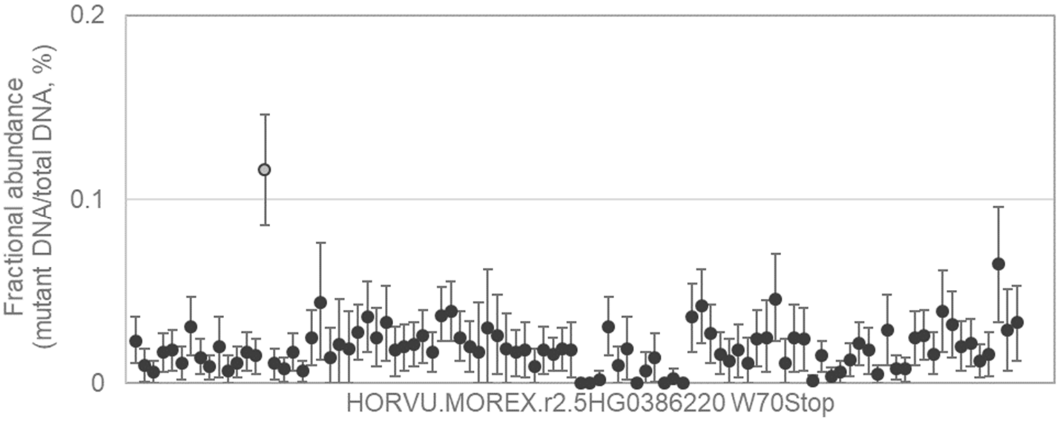


ACCAAGTTCTGCTACTACAACAACTACAGCATGTCCCAGCCGCGCTACTTCTGCAAGGCCTGCCGCC
GCTACTG[G/A]ACCCATGGTGGCTCCCTCCGCAACGTCCCCATCGGTGGTGGTTGCCGCAAGCCCAA
ACGCCCGGGGACCT

---

113

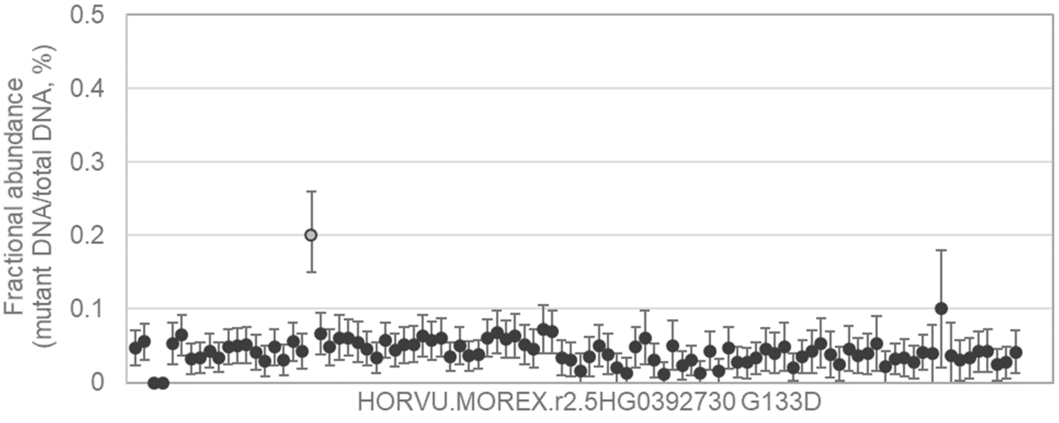


TCAGGGTGAGCTCGGCGGGAGGAGGAGCGGACGCCCCGGCCGACGACGGCCACAGCTGGAGGAAG
TAC[G/A]CCAGAAGGACATCCTCGGAGCCCATCACCCAAGGTTGGTCTCGGCGCATTTCATCCGTTCC
AAAGTTC

114

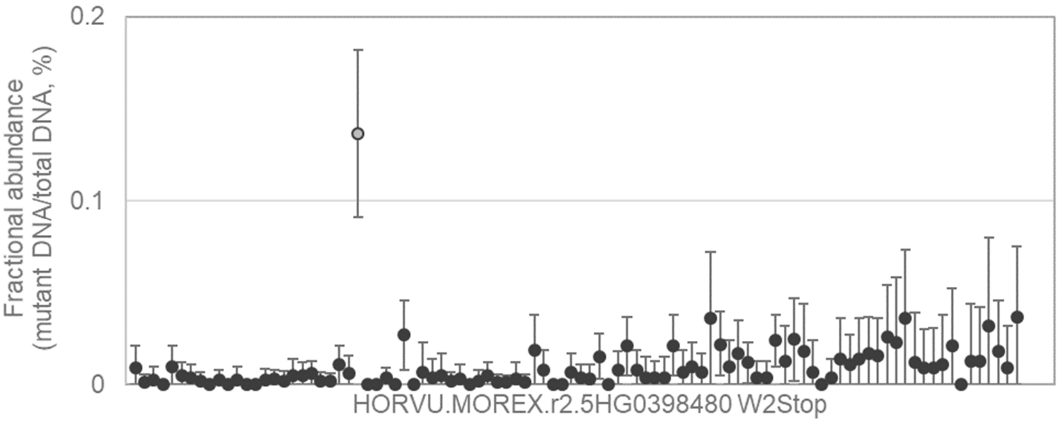


TATTCTTCTTCTTCCTCCTCCTCCTCCCTGCTTGCTGCTCTCGCGGGCGGGGGCGGGGGCGCGG
CCATGTG[G/A]ACGCCGTCGCGCGGGAGCAACGCGCGGCGGGCCGGCCACCGGCGCATCGCGGAG
GGCCTCCCGGACGA

---

115

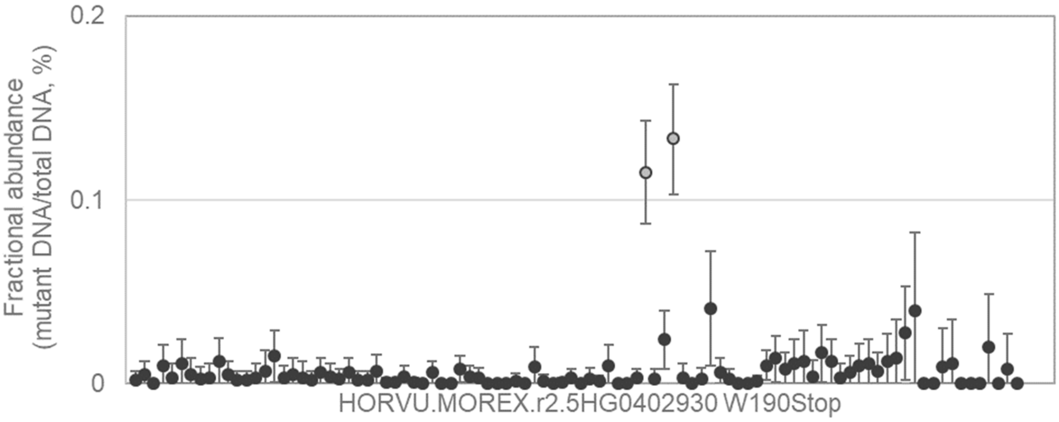


GCTCGGCCCCGAGGCCCTGCGCAAGCTCGTGCCGGACATCGAGGCCGCCGTTCGCTCCACGCTCGC
CGCCTG[G/A]GCGGACGGCGACGTCGCCAGCACTTTCCACGCCATGAAGAGGGTAAGCTATAGCAGC
TAGCTGCGCTGCG

---

116

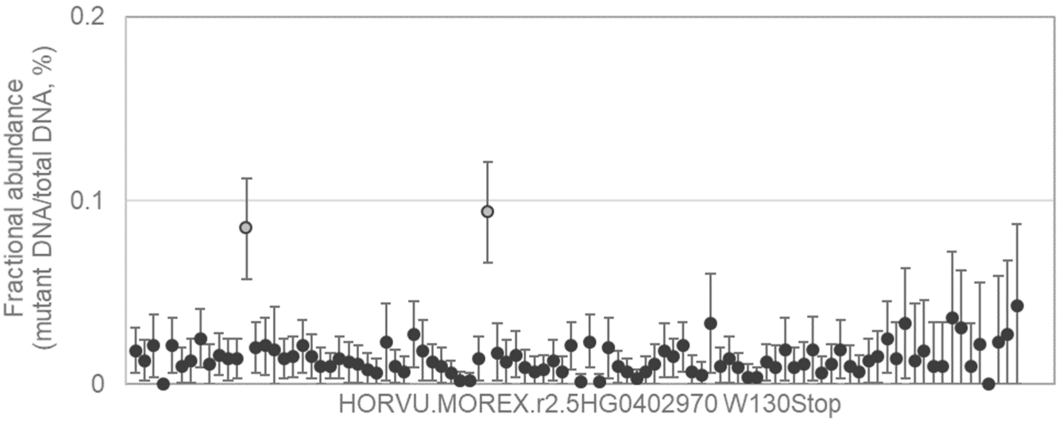


GGCGTGCGGCAGCGGCCGTGGGGGAAGTGGGCCGCGGAGATCCGCGACCCGCGCCGCGCGGTGC
GCAAGTG[G/A]CTCGGGACGTTCGACACCGCCGAGGAGGCCGCGCGGGCGTACGACCTCGCCGCGC
TCGAGTTCCGCGGC

117 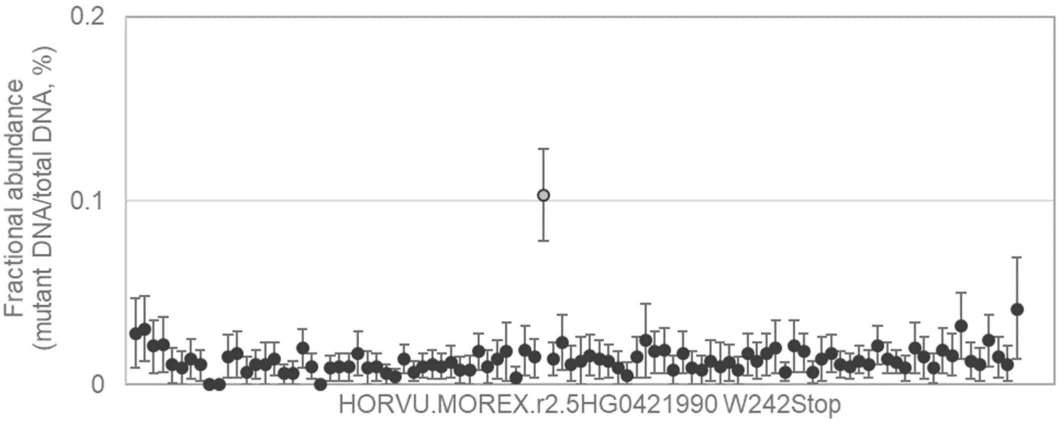


GCGCCGCCGACGCGTCGGAGATGGTGTGGGTGGACGTGCCGGACTGCTTCTGCTTCCACCTCTGGA
ACGCGTG[G/A]GAGGAGGAGGAGACCGACGAGGTGGTGGTGATCGGCTCCTGCATGACCCCCGCCG
ACTCCATCTTCAACG

---

118 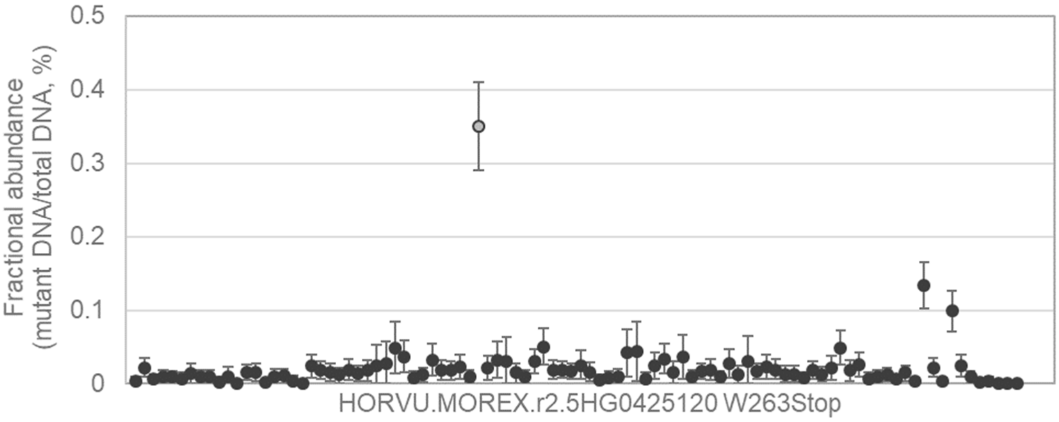


TTGTAAGAAATGATGTCTCTTATCCTGTGAAATTCTATGGCAAAGTGGTTGAAGGCACTGATGGGAGAA
AACACTG[G/A]ATTGGAGGAGAGAATATCAAGGCTGTGGCACATGACGTTCCTATTCCTGGCTACAAGA
CTAAAACTACTAAC

---

119 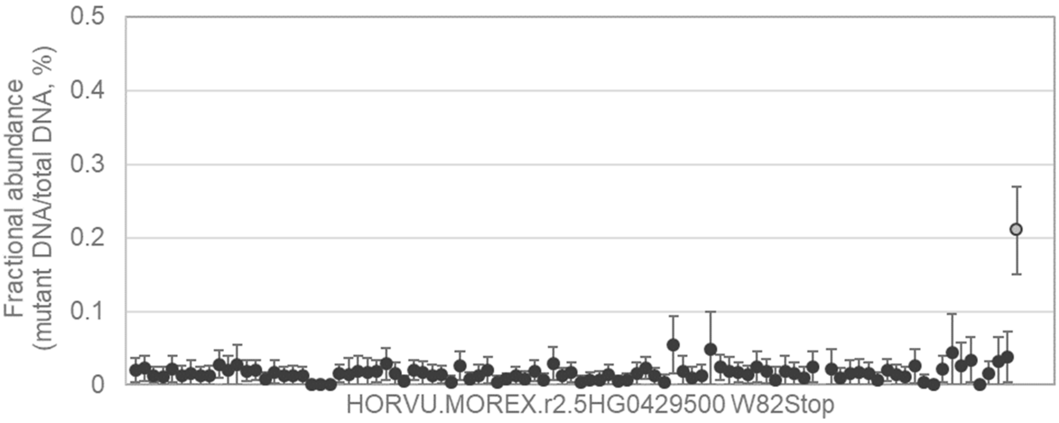


ACAAGAAGTTCTTGAAGTCGCAGGCGCTCGGCGGCCACCAGAACGCGCACAAGAAGGAGCGCAGCA
TCGGCTG[G/A]AACCCCTACTTCTACATGCCGCCCGCCTCCTCCGCGCACCTCCACGCCAATGCCGCG
CCGCCCGCCGCCG

120 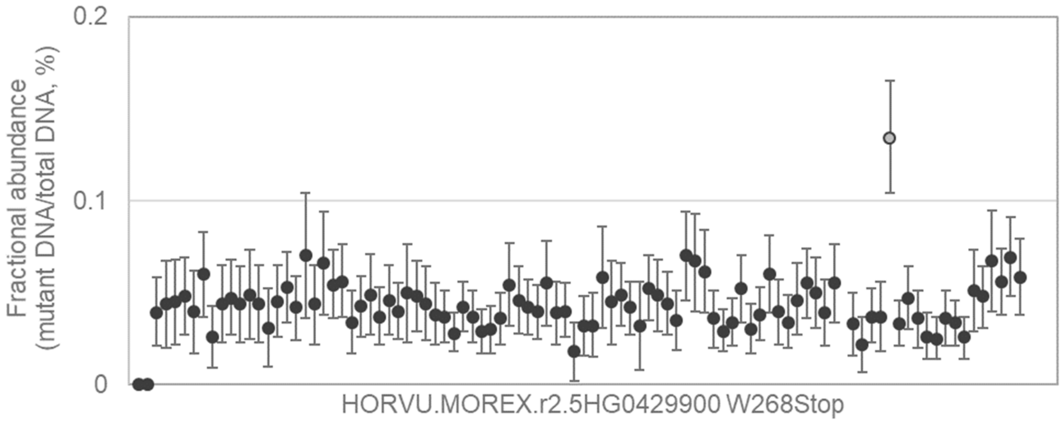


GCAATCGCCGGCGAAGACGGCTGGTACCATCCCAGAAGCATCCAGGAGTTGCATACCCTCTTCGACT
CCAACTG[G/A]TTCGACGGAAATTCAGTCAAGATCGTGGCGTCAAACACGGGCGCCGGAGTGTACAAG
GATCAAGACCTGTA

---

121 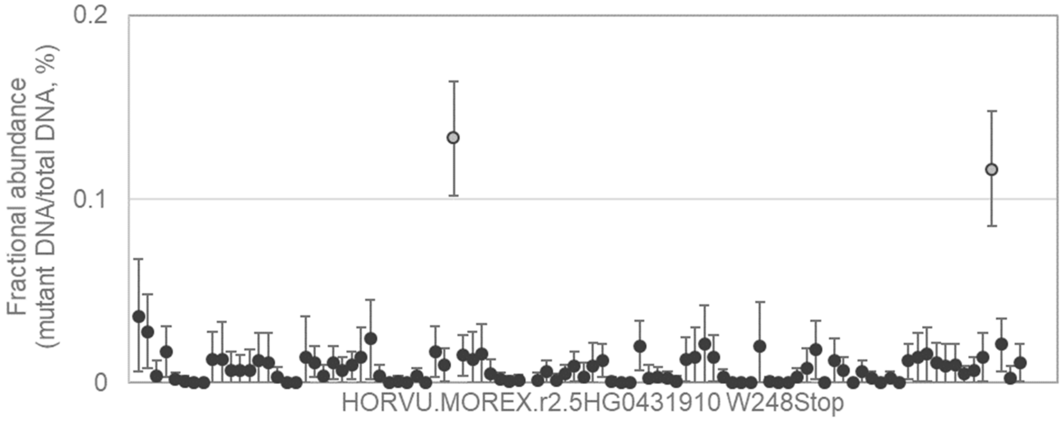


ATCCAGAAGCCGGCCTTCCTCGACTTCTTCACCAACCACTTGGACTGGGCCGTCTTCGAGAACGAGC
TCAAGTG[G/A]TACCACACGGAGGTGCAGCAGGGCCAGCTCAACTACGCCGACGCCGACGCGCTGCT
CGCGTTCTGCGACC

---

122 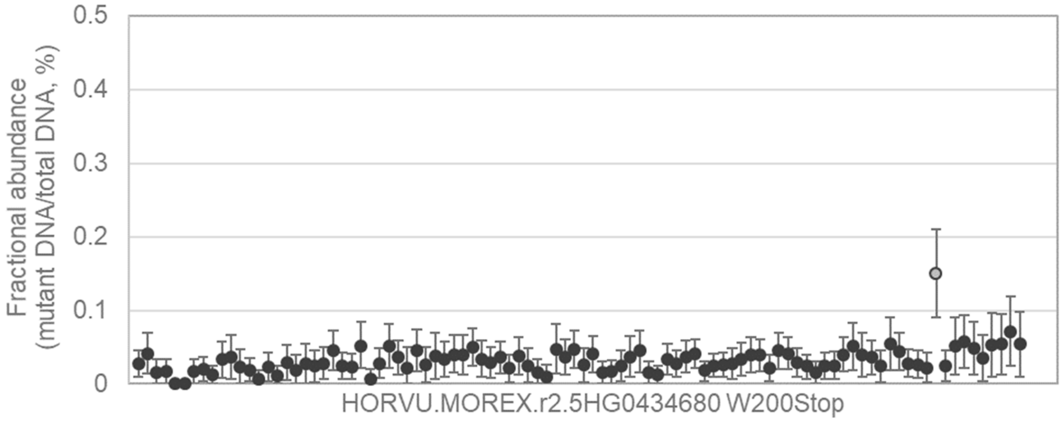


TTGTTGCCTCATGCTTCTGAATGTTTACAGCAAATCCATCAAGGGAGTTCTCTGGTAGCCTTGGCAATG
TTGCGTG[G/A]AAAGAGAGAGTTGATGGCTGGAAAATGAAAGACAAGGGTGCAATTCCCATGACTAAT
GGAACAAGCATTGCT

123

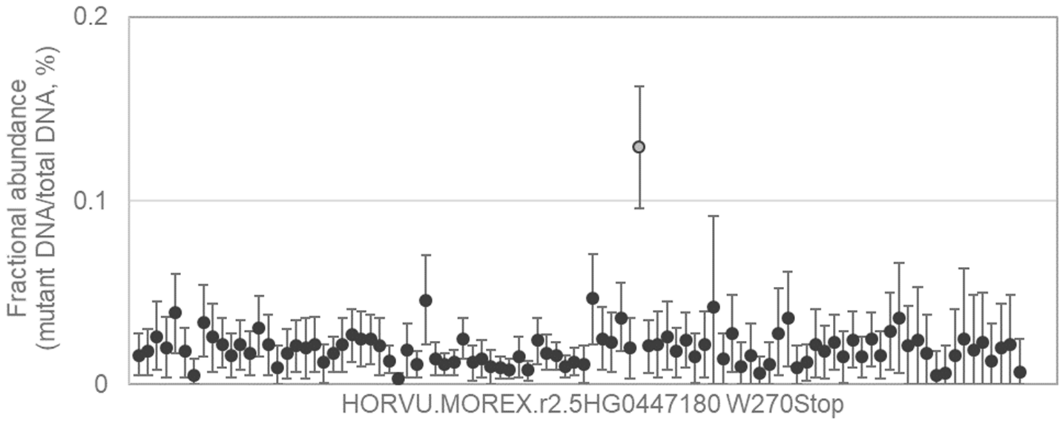


CCTTTGTAGGCACCGTGACATATATGTCACCTGAGAGAATTCGTAATGAGAACTACTCTTATGCTGCTG
ATATTTG[G/A]AGTCTTGGACTAACGATATTGGAGTGTGCTACTGGTAAATTTCCATATAATGTCAATGA
AGGCCCAGCCAAT

124

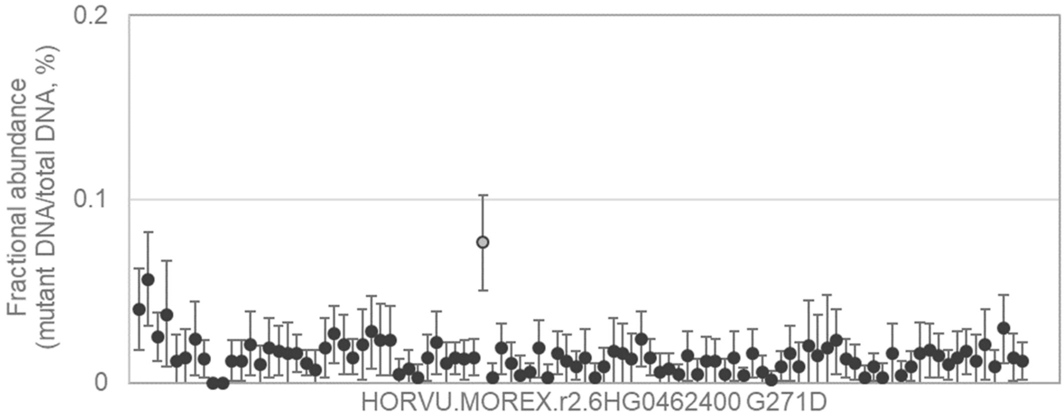


CACTATTTTTTTATTAATTATCATTTTAAACATCCGAGATGCTCAGGTTAAAATTGAAGGTGACATCAATC
AAG[G/A]TGTGGCATATTACAAGAAAGCTCTATTCTATAATTGGCACTATGCTGATGCAATGTATAATCT
TGGTGTT

125

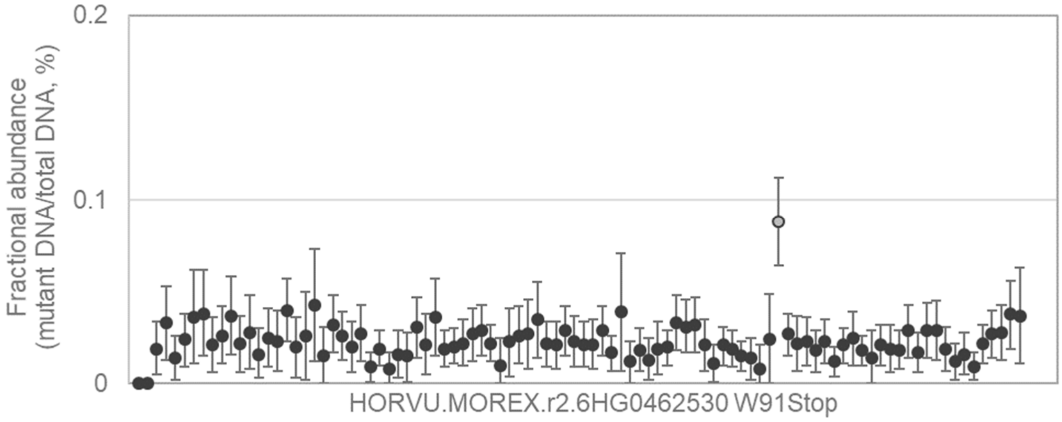


TCGTCGTGCTTGTTCATGCGCTCGGGCGGGATCATCATCCATCAGAGAAGGCGACCTTCGGGGAGCA
CGAGTG[G/A]TACTTCTTCAGCCCGCGCGACCGCAAGTACGCCAACGGCGCGCGGCCGAACCGGGC
GGCGACGTCGGGC

126

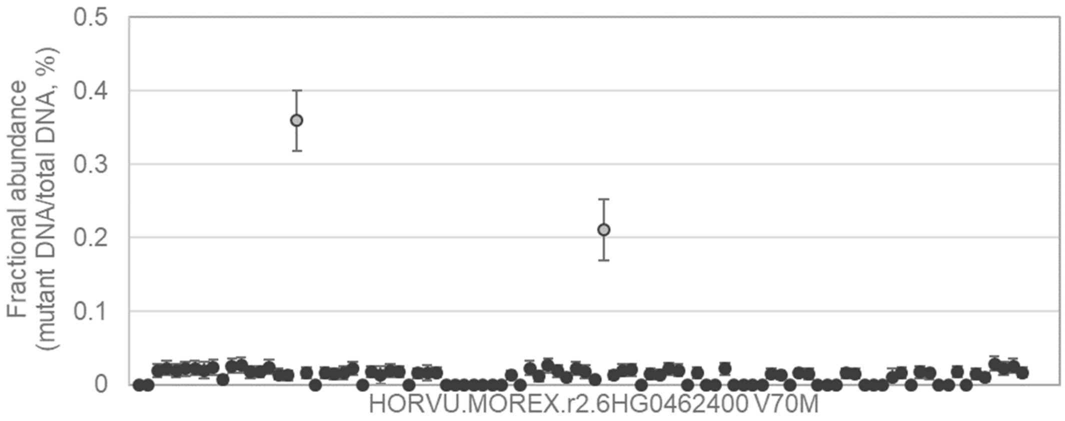


GGAGCTGGTCGTGCACTACCTCAAGAAGAAGGCCGCCAAGGCGCCGCTCCCCGTCACCATCATCGC
CGAG[G/A]TGGACCTCTACAAGTTCGACCCATGGGAGCTCCCCGGTATGTACTACTAGTTAGTACTATG
TCTATCCC

---

127

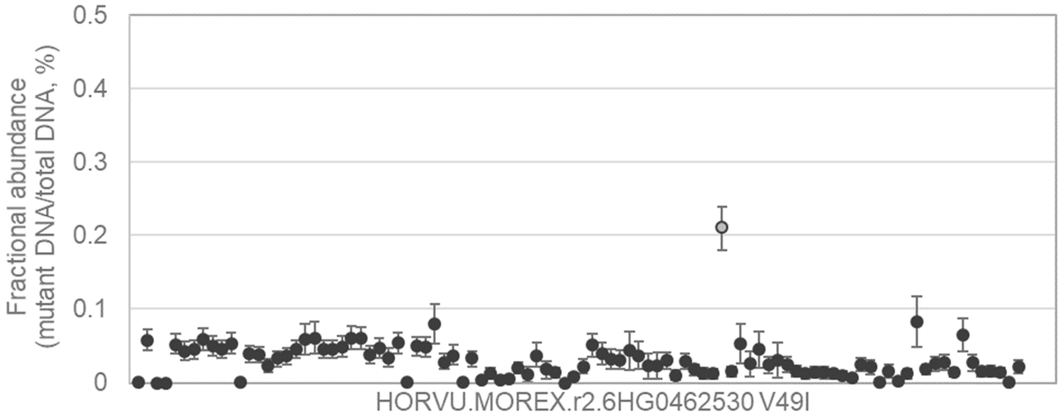


CCGCGGCGGCAGGGCTCGGCGCCGGAGCTCCCTCCCGGCTTCCGGTTCCACCCGACGGACGAGGA
GCTG[G/A]TCGTGCACTACCTCAAGAAGAAGGCCGCCAAGGCGCCGCTCCCCGTCACCATCATCGCC
GAGGTGGA

---

128

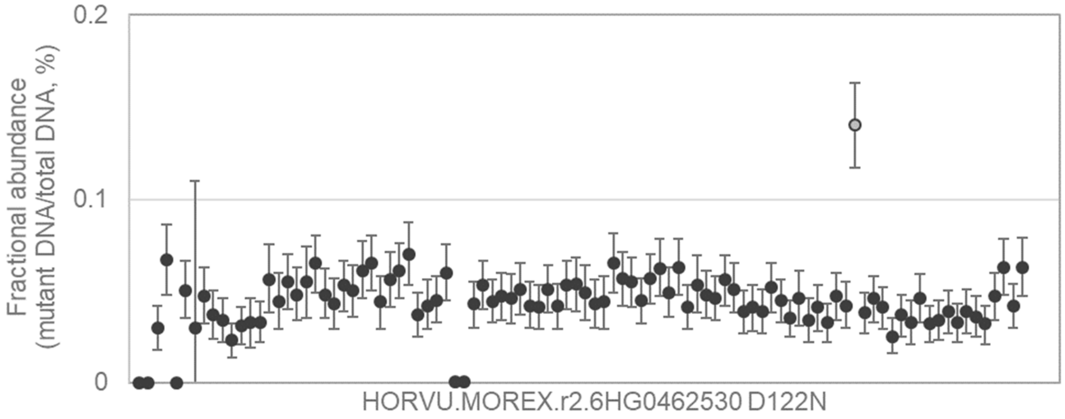


GCAAGTACGCCAACGGCGCGCGGCCGAACCGGGCGGCGACGTCGGGCTACTGGAAGGCCACCGGC
ACG[G/A]ACAAGCCTATCCTGGCCTCGGCCACCGGGTGCGGCCGGGAGAAGGTCGGCGTCAAGAAG
GCGCTCG

129

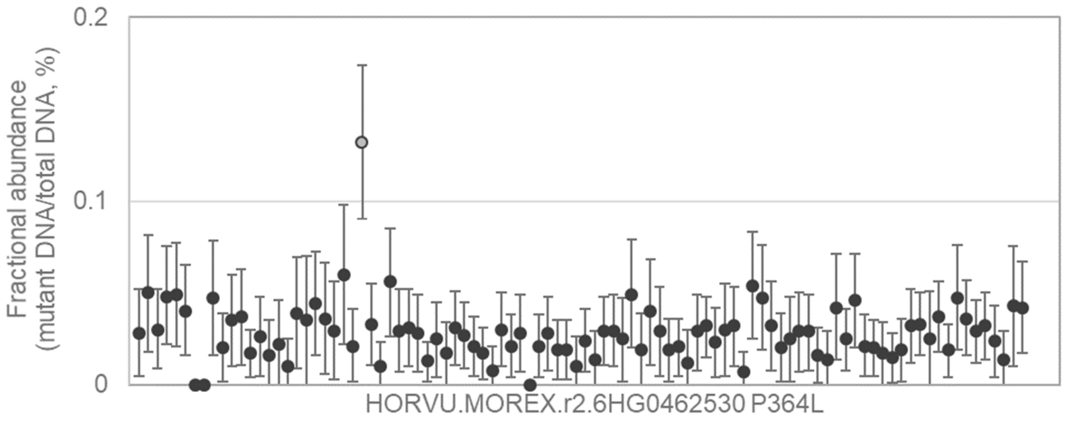


CGGCCGACGACATGGCTGGCGCGGTGGACGTCAGCGACGGAGGCAATGCGGTGAACGCCATGTATC
TCC[C/T]CGTGCAAGACGGGACCTACCATCAGCATGTCATCCTCGGAGCTCCGCTGGCGCCAGAGGC
CATCGCG

---

130

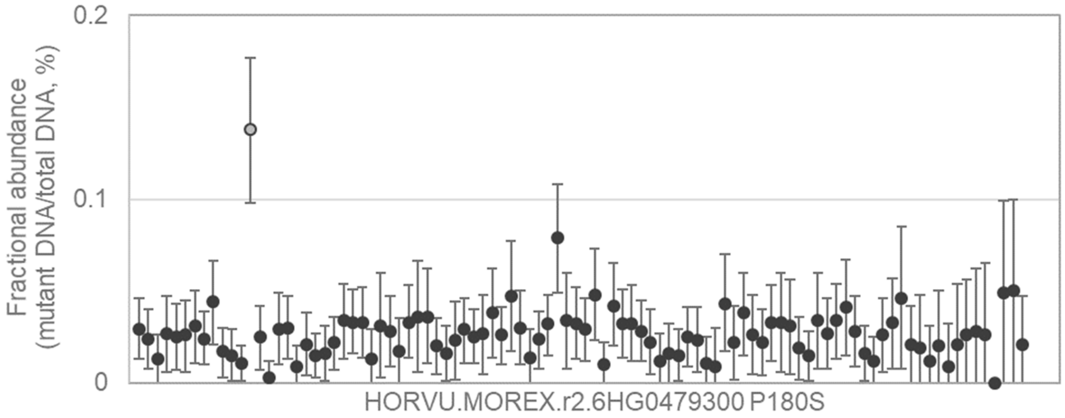


TGGATCATGCGGACGGAGGGGTGGCCCGGCCTCTTCCGCGGCAACGCCGTCAACGTCCTCCGCGTC
GCG[C/T]CGAGCAAGGCCATCGAGGCAAGCCAGCCACCTCACGTGTTTTCCTTTGAAATTTCCTATAGT
AGACGC

---

131

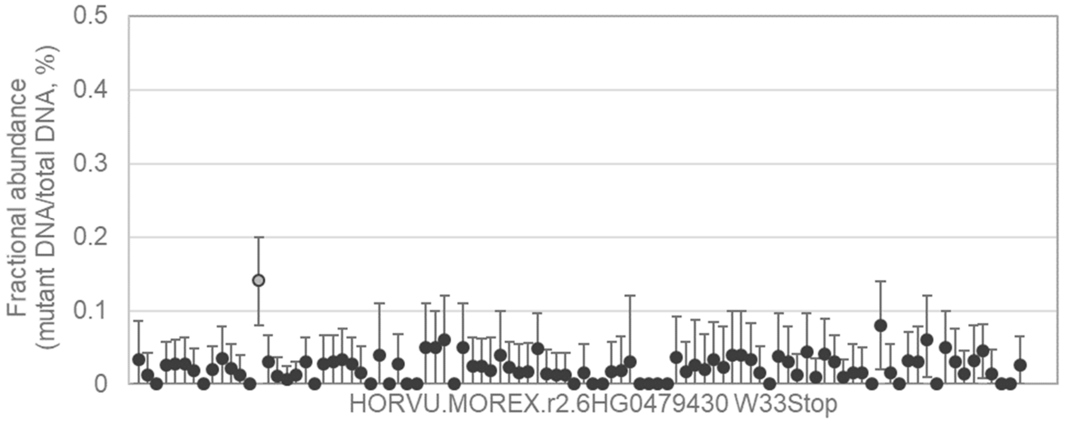


TAGTGGTTTCTTCGCCGGTTTACAAGATTCAAACTAACAGCATGCGTGTCTTTTGTGAAGTAAGAGGAG
GGTGTG[G/A]CTTGGTACCTTTGAAACCGCTGAAGAGGCCGCACGGGCCTATGACGAGGCTGCCGTT
CTGATGAGCGGAC

132
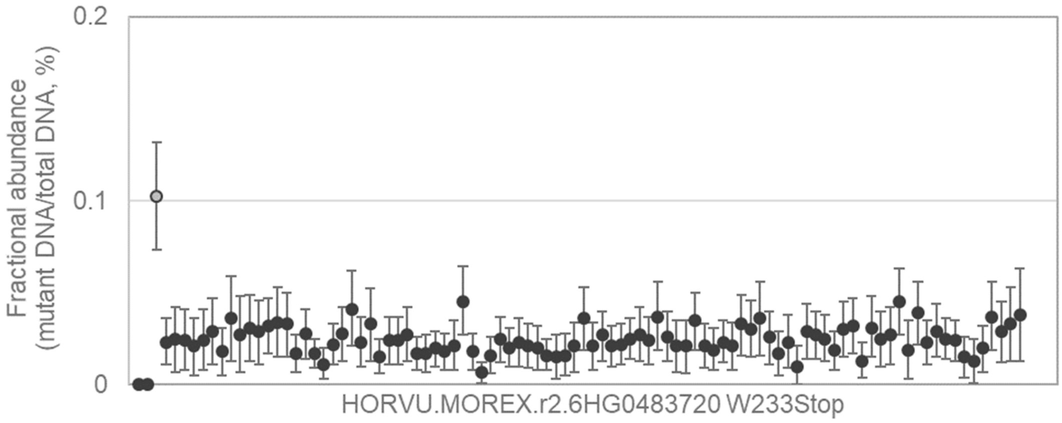


GTAAATTGTGTTTTGCTTACAGTGATGGTAACATAGACATGAATATAGAGGACATGATGGTTATGGAAG
CGATTTG[G/A]CGTTCAATTCAGGTAAAGTAAAAGGTTCTTATATGCCCCTTGAATGAACAGTATCTGGT
TAGTGCAAGATGTA

---

133
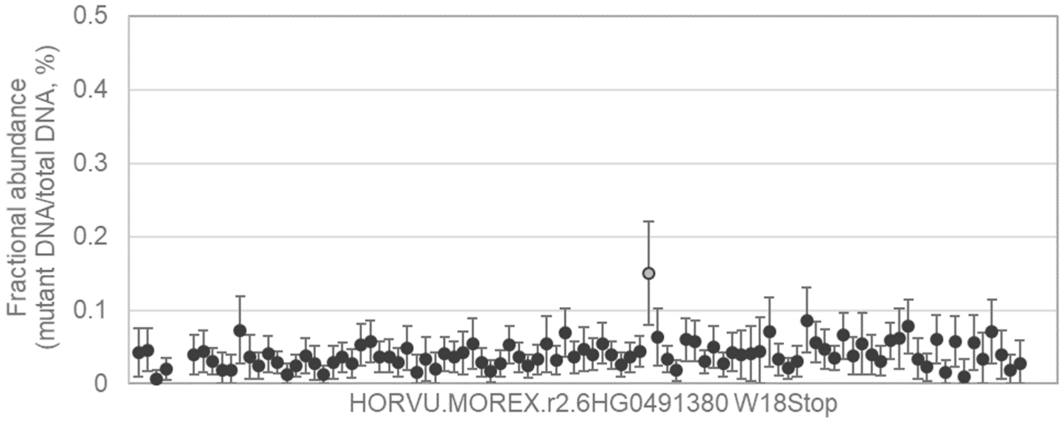


CGTGTGAGGTTCCCGTGGGAATGGCAGAGTCGGACGGCGCCGAGTACCGCTGCTTCGTCGGCAGCC
TGTCCTG[G/A]AACACCGACGACCGCGGCCTCGAGGCTGCTTTTAGCTCCTTCGGAGAGATCCTCGAC
GCCAAGGTCCGCC

---

134
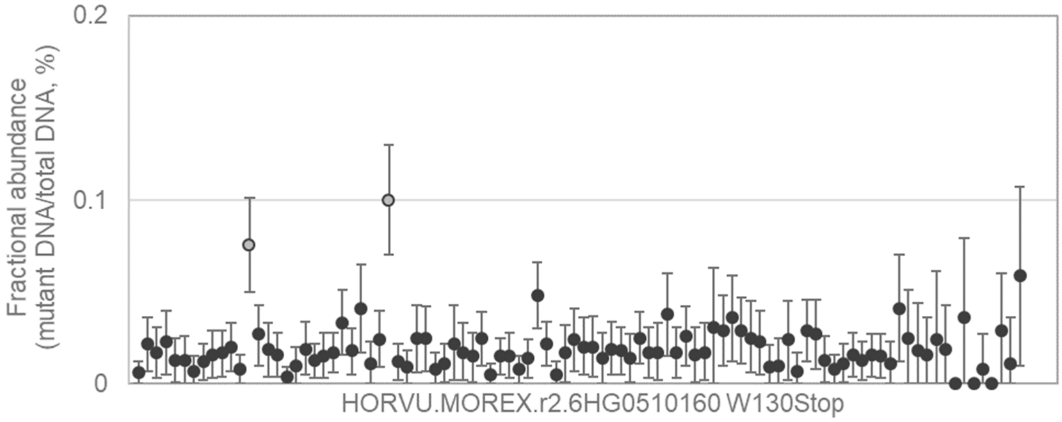


ATATTCACGAACCGATTCGATGTGCATATGTTCTGATGGATGATTGTGCTTGCATGCGCAGGTCCCGT
TCTACTG[G/A]GCGGCGCAGTTCACGGGCGCCATCTGCGCGTCCTTCGTGCTCAAGGCGGTGCTCCA
CCCCATCACCGTGA

135

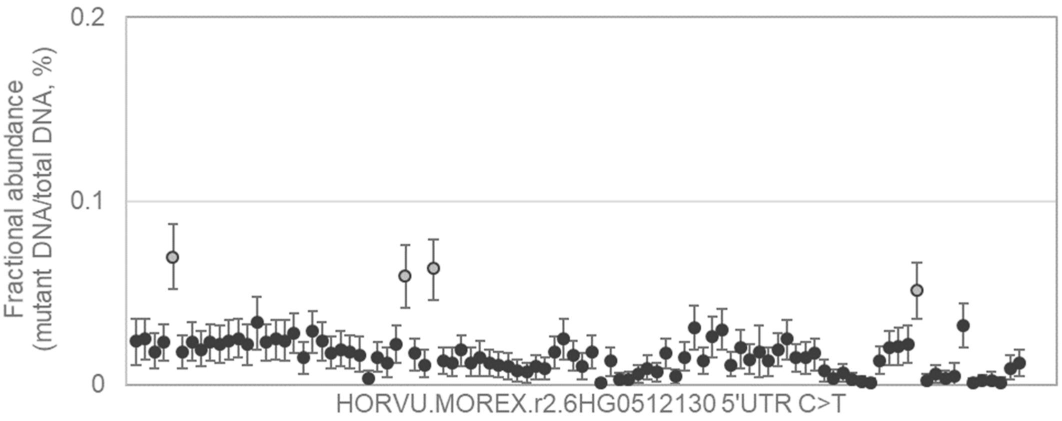


ATTCGATGAAAAATGCTCGAATGTTCTGTCCTCAGAAAAACAGAGGTTGAGGATAACTGACGGTCGTA
TTGA[C/T]CGGTGCCTTCTTATGGAAGGCGAAGGCTGCCTCCATCTACATCACTTGGGCATTGAATCGC
CTTTTGA

---

136

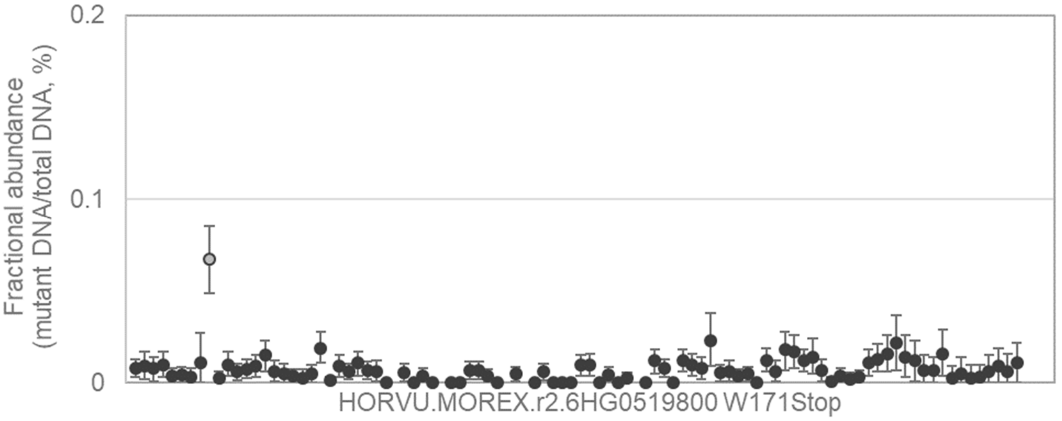


AATTAAATTATTTGGTCTTGACGTTCTTTGTTTGACACATTTTGCTACAGGAACTATCCTCTCCTGATGG
CTGTATG[G/A]AAGGTAGCCCCTGCTCTGGCAGCTGGTTGTACAGCTGTACTGAAACCATCTGAATTAG
CTTCCGTGTAAGTT

---

137

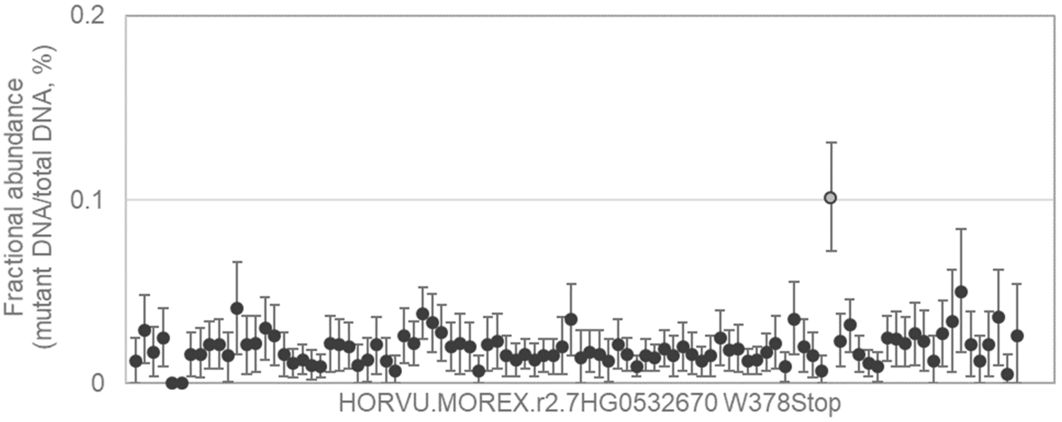


GAGTTGGATCATCAGGTGAAACCAGAGGGCAACCATAAGCTGACGACGAAGGAGAACGCAATGAACC
AGCCTTG[G/A]TCCAGACTACCTCCAGCACCAGTGAGGAACAACAGTGAGTGCCGTACTTGGAATGGC
TTCCGTTTCATTCAA

138

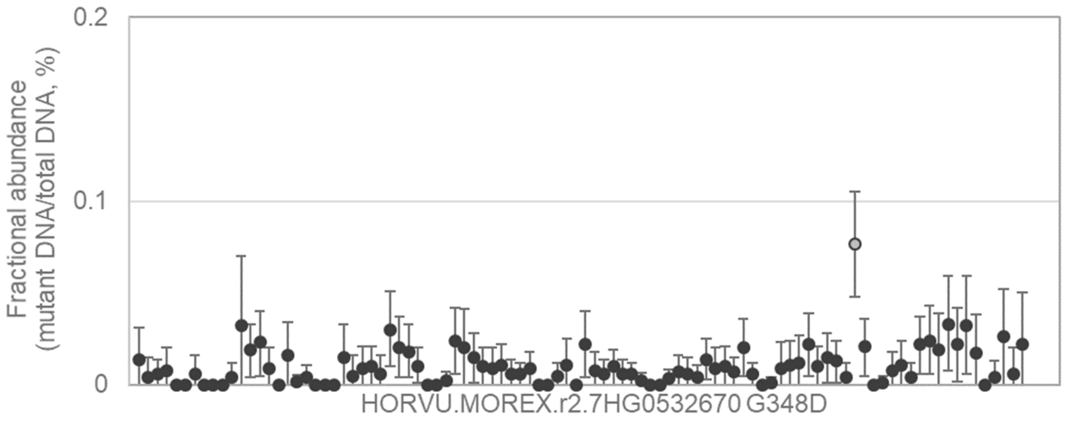


GACTTTATCCAATGGGAGGCCGGCAGTTAATAACTCCTACCTTAGCACCAACGCCCCAGCAAGAGCA
GCTG[G/A]CGTGCAAAGGAAGCTAGAGTTGGATCATCAGGTGAAACCAGAGGGCAACCATAAGCTGAC
GACGAAG

---

139

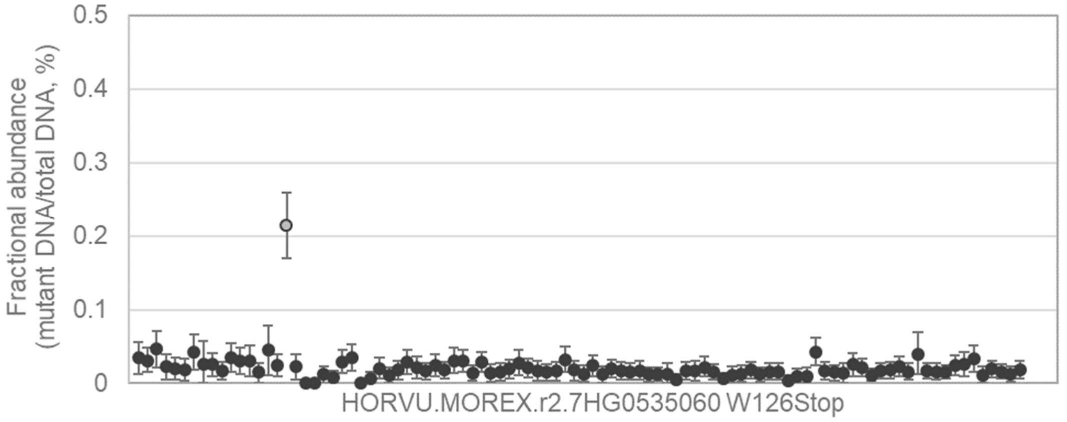


GCGTGTCTGTTCAGGCCAACGGTCACCGGGTGATGGTCGTCTCCCCGCGCTACGATCAGTACAAGGA
CGCCTG[G/A]GACACCAGCGTCATCTCCGAGGTAGGTATCCGCCATATGAATGATCACGCTTCACATG
CTCCTGCACATTTC

---

140

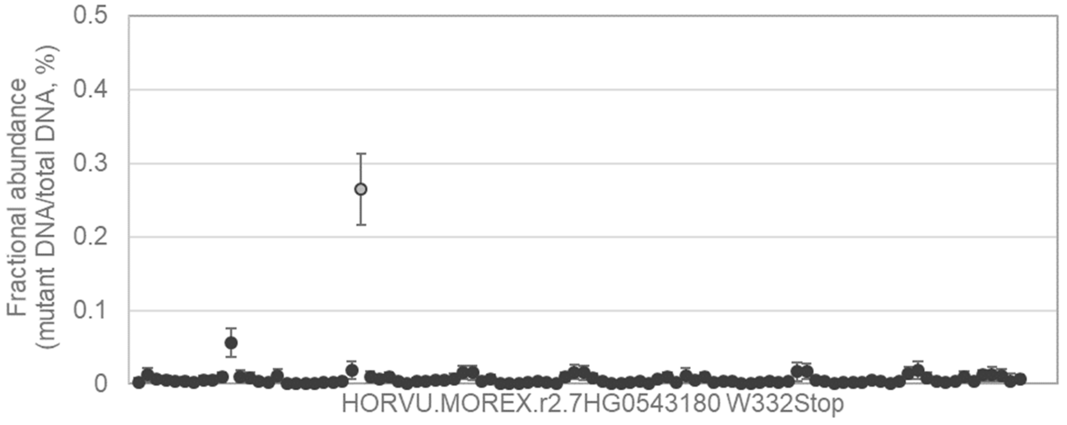


AATGTATGTTATTTCAATGTCAATAGGGTGTGGAGCCTGCAAGTACATATCCTGATCTGGGATTGCCAC
CTGAATG[G/A]TATGGAGCTTTAGAATGGGTATTTCCAGAATGGGCAAGAAGGCATGCCCTTGACAAG
GGTGAGGCGGTTAA

141

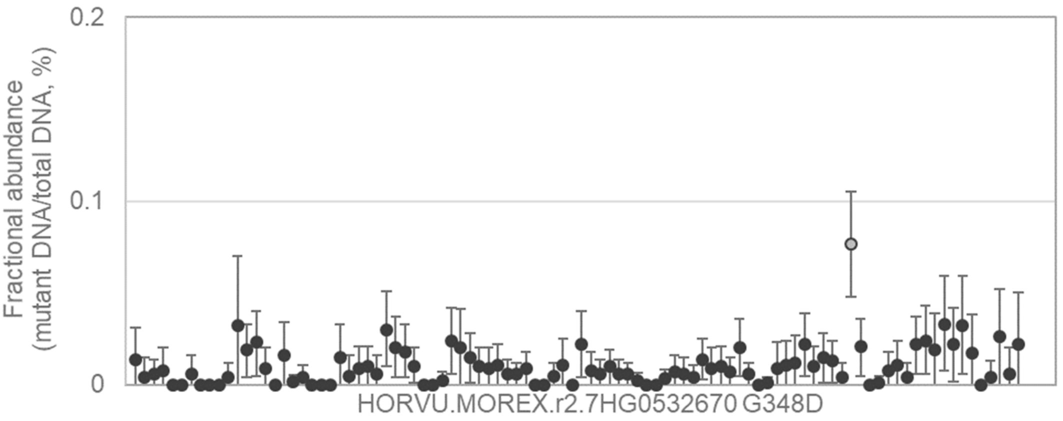


TGGCCATTCCAGATCTCATGAGGGAAGACGTACAGTTTGTCATGCTTGGATCGGGGGATCCAGTTTTT
GAAG[G/A]CTGGATGAGATCTACCGAGTCGAGCTACAAGGATAAATTCCGTGGATGGGTTGGATTTAG
TGTTCCAG

---

142

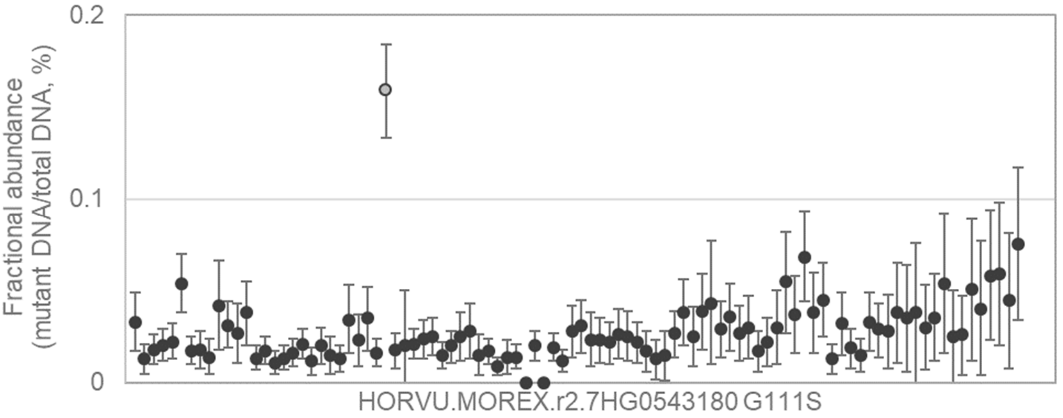


CAGGATTCTGAGATCATGGATGCCAACGACCAACCTCTAGCTAAAGTTACACGTAGCATCGTGTTTGT
GACT[G/A]GTGAAGCTGCTCCTTATGCAAAGTCAGGGGGGCTGGGAGATGTTTGTGGTTCGTTGCCAA
TTGCTCTT

---

143

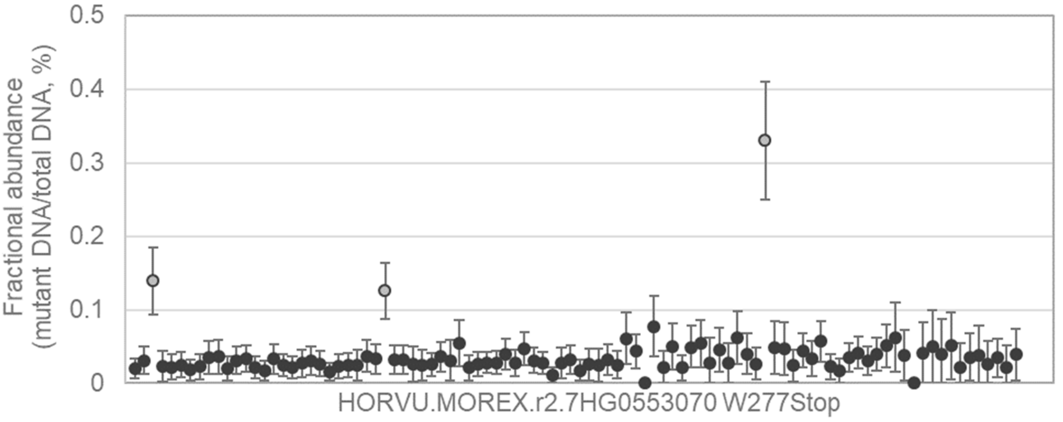


GCTGATTGTCAAAGAAGCTCCAAAACCAAAGGCTCTTTCGGCCCCTGCAGCCCCCGCTGTACAAGAA
GACCTTTG[G/A]GATTTCAAGAAATACATTGGTTTCGAGGAGCCCGTGGAGGCCAAGGATGATGGCTC
GGCTGTTGCAGATGA

144

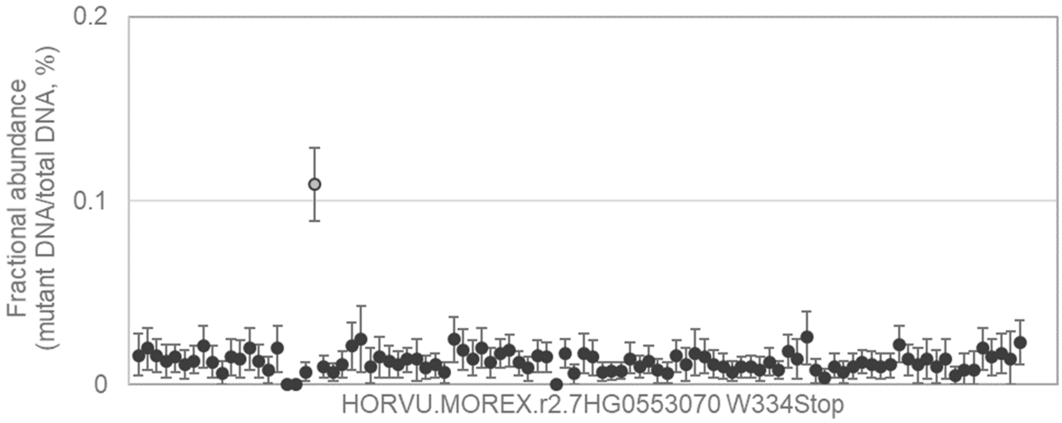


AATCATGATTCCGGACCTTTGGCAGGGGAGAACGTCATGAACGTGGTCGTCGTTGCTGCTGAATGTTC
TCCCTG[G/A]TGCAAAACAGGTATGGGCATTGCCTTTTCAGTCTCTCTCTCTCTCTCTTTCTGTTGTT
CATCAAACTTTGCT

---

145

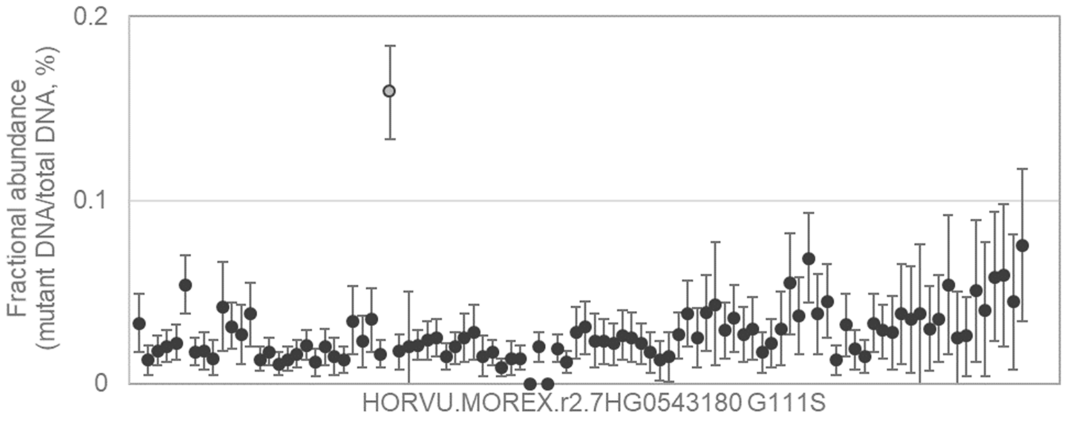


GCAAGGAGGCCCTGCAGCGCGAGCTGGGGCTGCAGGTCCGCGGCGACGTGCCGCTGCTCGGGTTC
ATCG[G/A]GCGGCTGGACGGGCAGAAGGGCGTGGAGATCATCGCGGACGCGATGCCCTGGATCGTG
AGCCAGGA

---

146

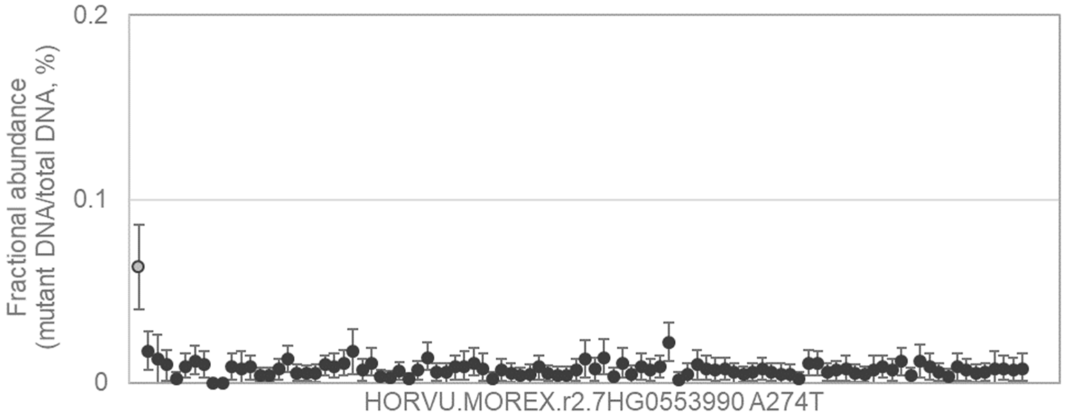


CTCCATGGTTCTCATAGAAATCCAGGACACGGTGAAGGCGCCGCCGGCCGAGAACAAGACAATGAGG
AAG[G/A]CAAACCTACTGGGGGTCTCCACCACCACGGCCTTCTACATGCTCTGCGGCTGCCTCGGCTA
CGCTGC

147
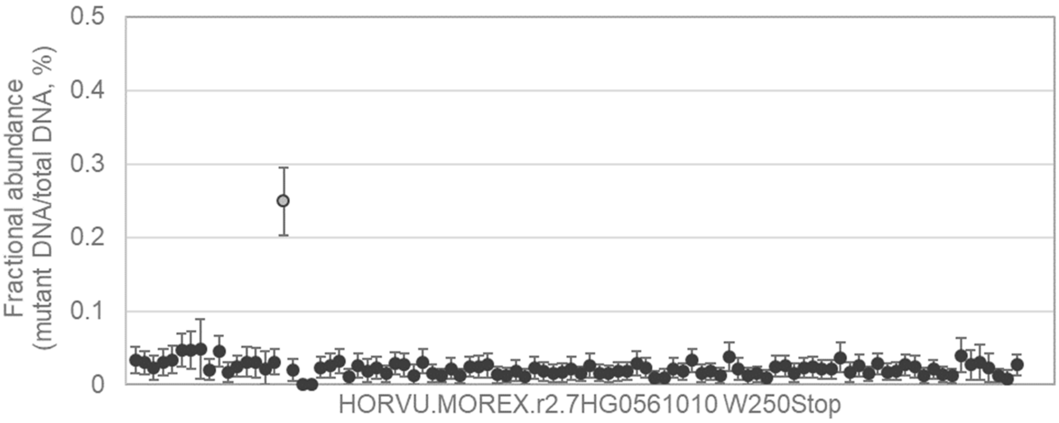


GCCGCGACGCCTTCCGCGCCTCCAAGATCATGCAGGCCCGGGCTCTCTCCAACCCCAAAATCCAGGT
CGTGTG[G/A]GACTCCGAGGTCGTTGAGGCCTACGGCGGCTCGGACGGCGGCCCCTTGGCTGGCGT
CAAGGTAAAAAACC

---

148
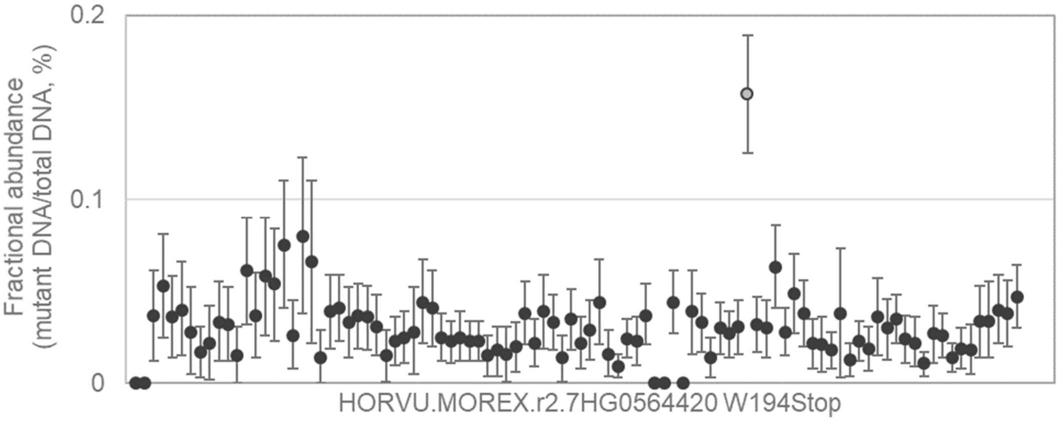


GAACCCGGCAGGTTCAGAAGCTACCTGCTGGATTTCTCCTACCCAAGGGTTCCAAGCAGCGTGCGGG
ATGCATG[G/A]CCGGGGGCTCGGCCAGGCTACCGGATGCCCGGTGAAATCCAGTGGCAAGGGAACCT
AGACCTGCGTCCT

---

149
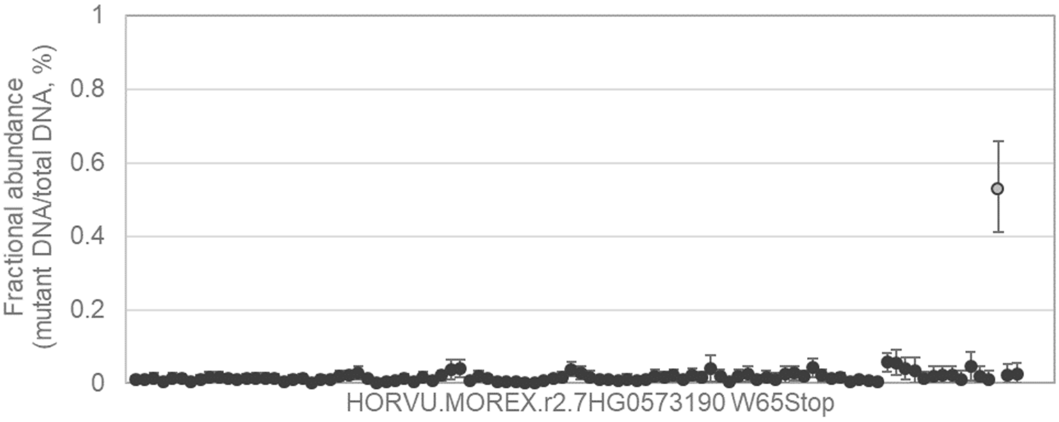


GAATCTTCCAGAAGGCTGGGCCTGGTCCTCAAAACGGGGCACAGTATGGAGCACCCTTCGTCGAGGA
GGAGTG[G/A]GAGGCTGATGAGGATGATGATTTTGGTCCTCTGCCTGTGGAGAGGAACCTCGTTGTTG
AGCATGAGGCTCC

150

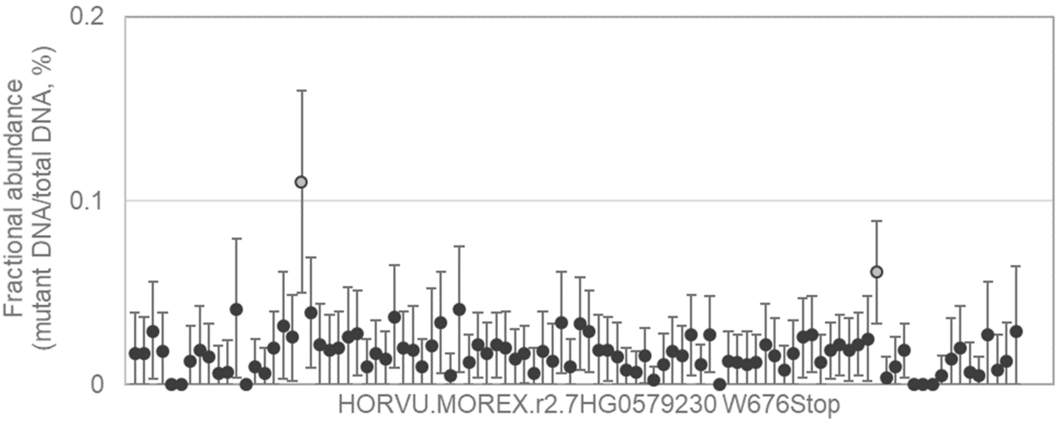


TCCATCTACCCACACGCCTTCATCGGCACCGCCCCCATCAACCTCACGGAGAGGCTCTTCCAGGTGC
TCCGCTG[G/A]TCCACGGGATCCCTCGAGATCTTCTTCTCCAAGAACAACCCGCTCTTCGGCAGCACA
TACCTCCACCCGCT

151

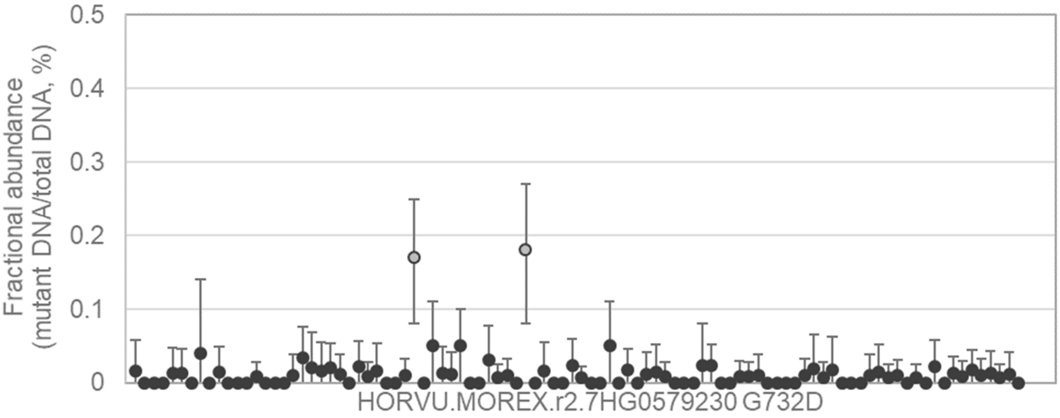


TCACCACTTACCCCTTCACCGCCATCTTCCTCATCTTCTACACCACCGTGCCGGCGCTATCCTTCGTC
ACCG[G/A]CCACTTCATCGTGCAGCGCCCGACCACCATGTTCTACGTCTACCTGGGCATCGTGCTATC
CACGCTGC

152

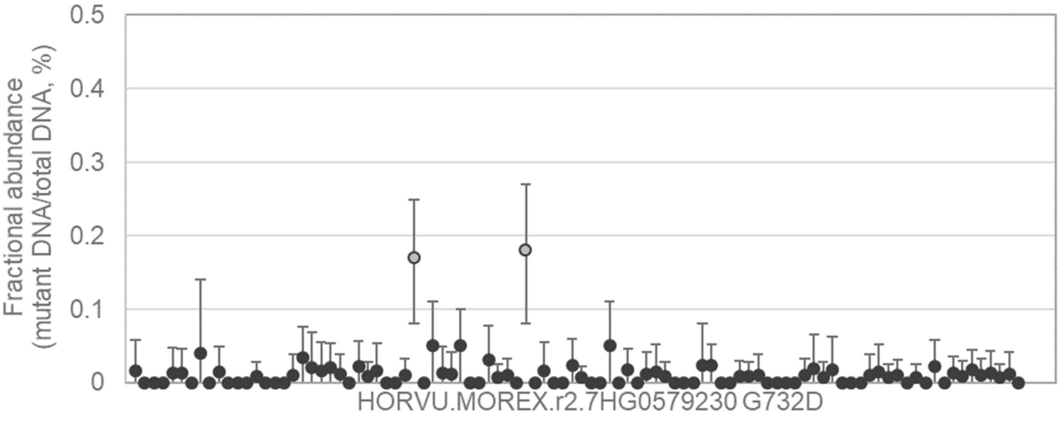


GCCGGCGCTATCCTTCGTCACCGGCCACTTCATCGTGCAGCGCCCGACCACCATGTTCTACGTCTAC
CTGG[G/A]CATCGTGCTATCCACGCTGCTCGTCATCGCCGTGCTGGAGGTCAAGTGGGCCGGGGTCA
CAGTCTTC

153

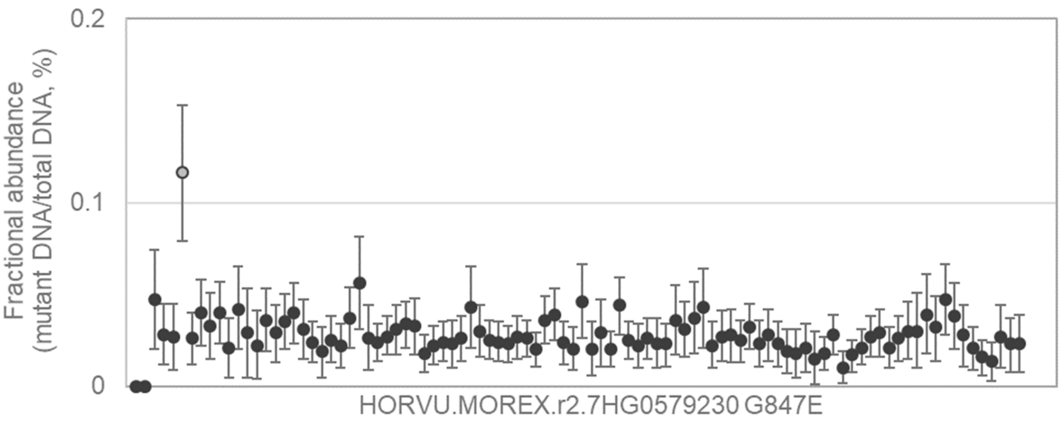


CCGACCTCTACGTGGTGCGCTGGACGCCGCTCATGATTACACCCATCATCATCATCTTCGTCAACATC
ATCG[G/A]ATCCGCCGTGGCCTTCGCCAAGGTTCTCGACGGCGAGTGGACGCACTGGCTCAAGGTCG
CCGGCGG

154

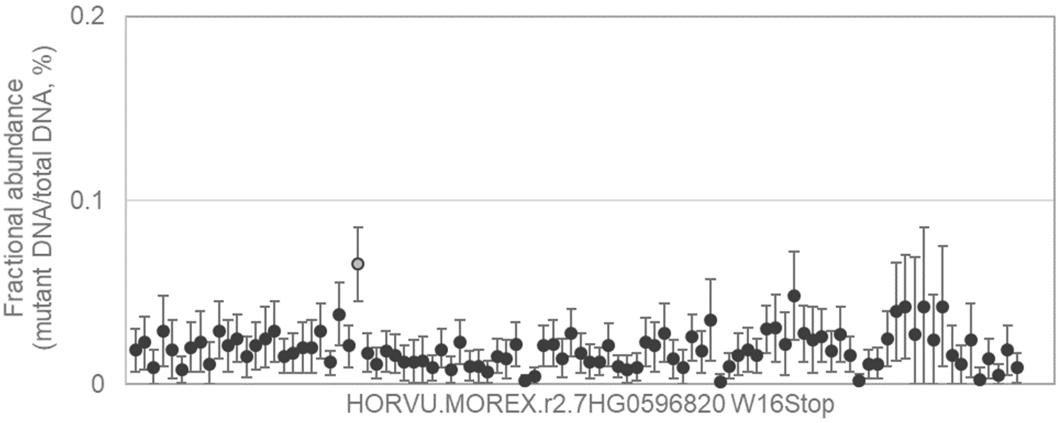


TCGCTCTGCATTCCGCAGAGAAACATCATGGTACAGTCCAAGAAGAAGTTTCGCGGCGTCAGGCAGC
GCCACTG[G/A]GGCTCCTGGGTCTCCGAGATCAGGCATCCTCTCCTGTAAGCCTCTCGCTAGCTCTCT
CTCTCTCTATAGCTG

155

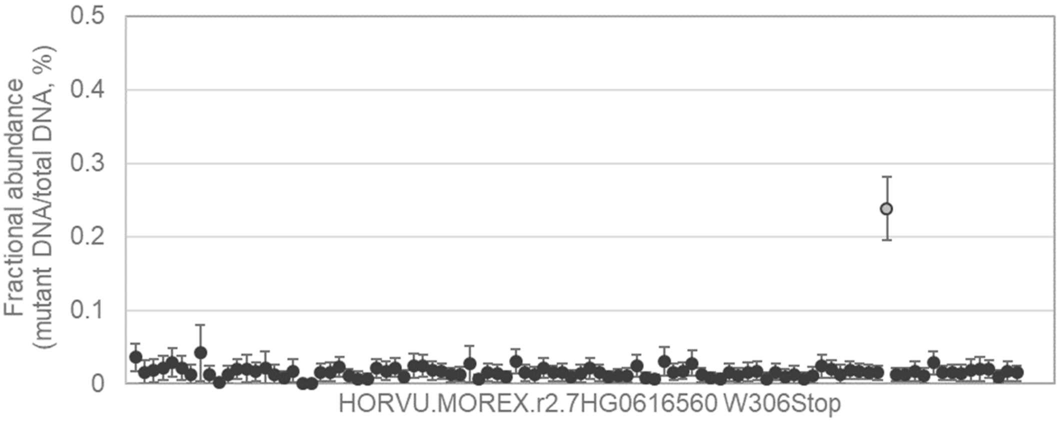


TAAAGGGATACAAGGATCAGGATCTATATAACAAGTATATTGACATCGGTGAGATTCCAGAGCTTTCAG
CTATTTG[G/A]AAGAAAGACAGTGGCATTGAAATTGGAGCAGCTACACCAATTTCTAAGACCATCGAGA
TACTTGAGCAAGAA

**Supplementary Notes 2, Section 2: Beyond barley. Fractional abundance values of variant DNA to total DNA following ddPCR in Phase 1 of the FIND-IT screening process in wheat (*Triticum aestivum*), yeast (*Saccharomyces cerevisiae*) and bacteria (*Lactobacillus pasteurii* (DSM 23907)).** Dots represent individual pools on a library plate, with the positive pool(s) identified for further downstream screening in FIND-IT phase 2 marked in gray open circles. The gene sequences and the targeted SNP are shown under the graphs.

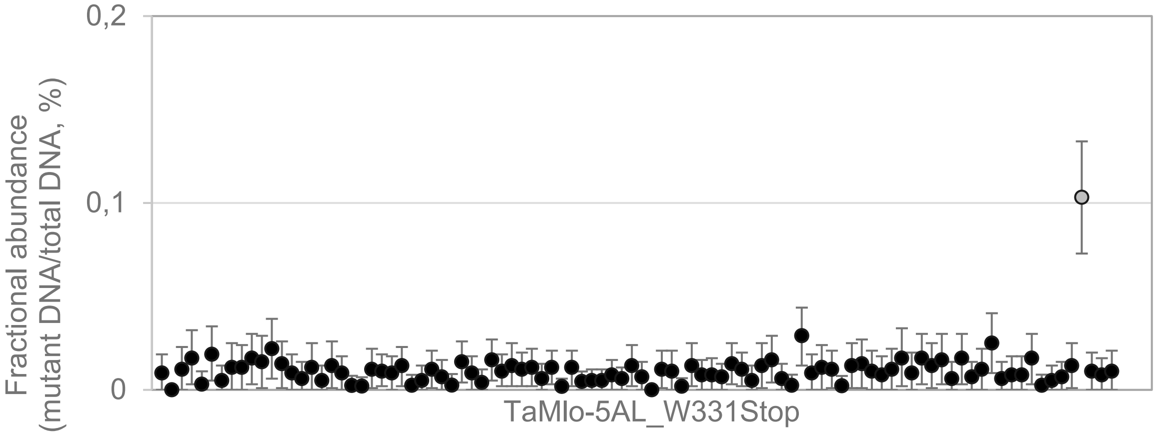


CAGATCCTCTTGTGTGTTGGAACCAAGCTAGAGATGATCATCATGGAGATGGCCCTGGAGATCCAGGACC
GGTCGAGCGTCATCAAGGGGGCACCCGTGGTCGAGCCCAGCAACAAGTTCTTCTG[G/A]TTCCACCGCC
CCGACTGGGTCCTCTTCTTCATACACCTGACGCTGTTCCAGAACGCGTTTCAGATGGCACATTTCG

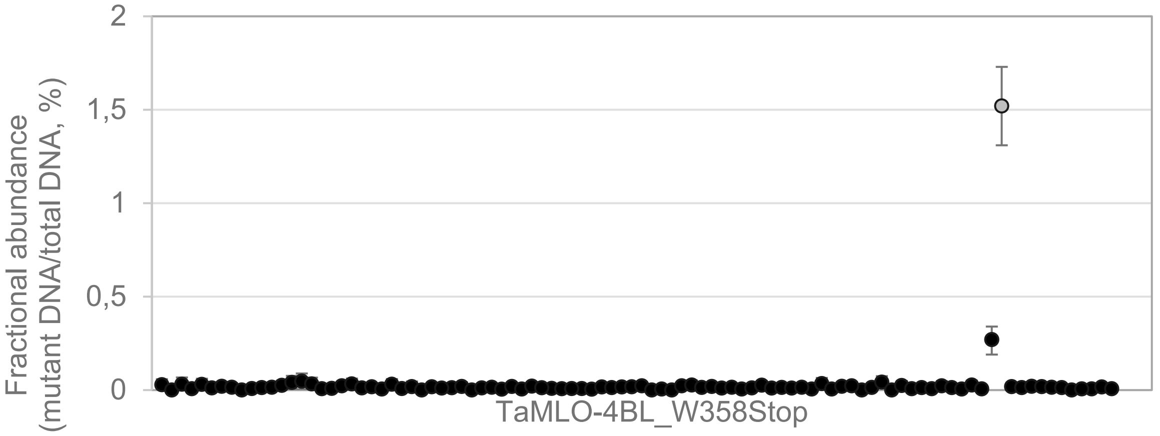


CCGCCCCGACTGGGTCCTCTTCTTCATACACCTGACGCTATTCCAGAACGCGTTTCAGATGGCACATTTC
GTGTG[G/A]ACAGTGGTATGCACCAGTAATTGGCAGTTCAGTTAGGGATGCAAGGCACCAAGTAGTGCTG
ATGAACTGCACTGACGGAGATTTACTTGTTCGTAGGCCACGCCCGGCTTGAAGAA

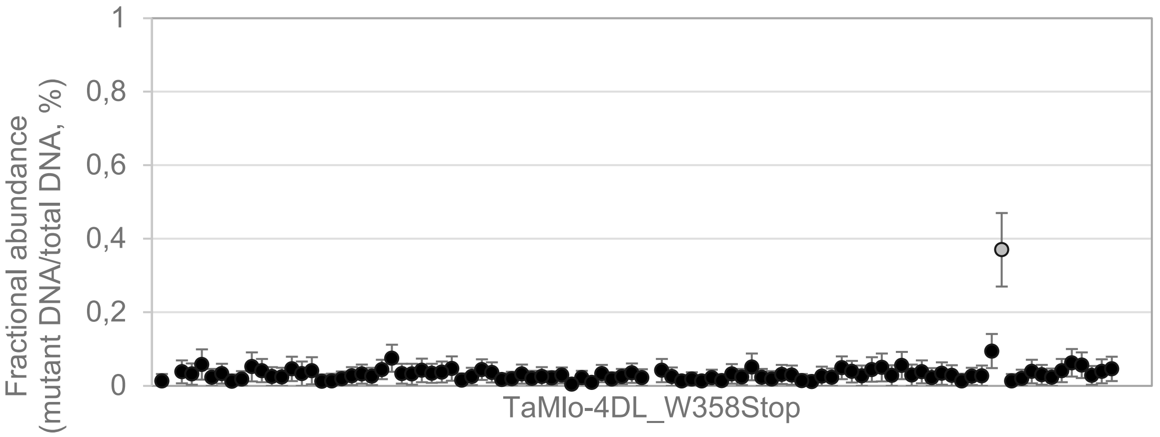


CAAGGGGGCGCCCGTGGTTGAGCCCAGCAACAAGTTCTTCTGGTTCCACCGCCCCGACTGGGTCCTCTT
CTTCATACACCTGACGCTGTTCCAGAATGCGTTTCAGATGGCACATTTCGTCTG[G/A]ACAGTGGTATGTA
CCAGTAATTGGCAGTTCGGTTAGGGATGCAAGGCACCACGTGCTGCTGCTGATGAACTG

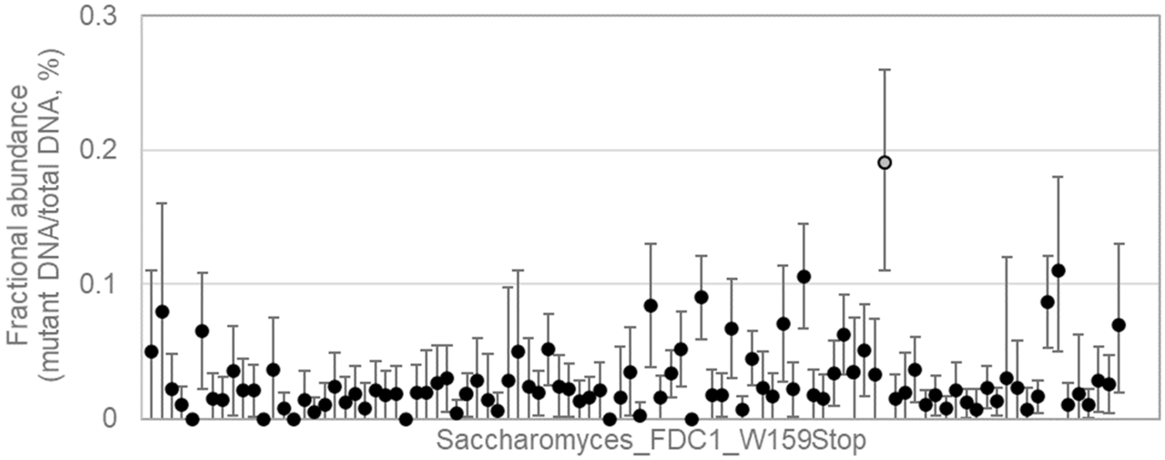


GCCTGCCAACACCATATCTACATGTTTCAGACGGTGGCAAGTACTTACAAACGTACGGAATGT[G/A]GATT
CTTCAAACTCCAGATAAAAAATGGACTAATTGGTCAATTGCTAGAGGTATGGTTGTAGATGACAAGCATAT
C

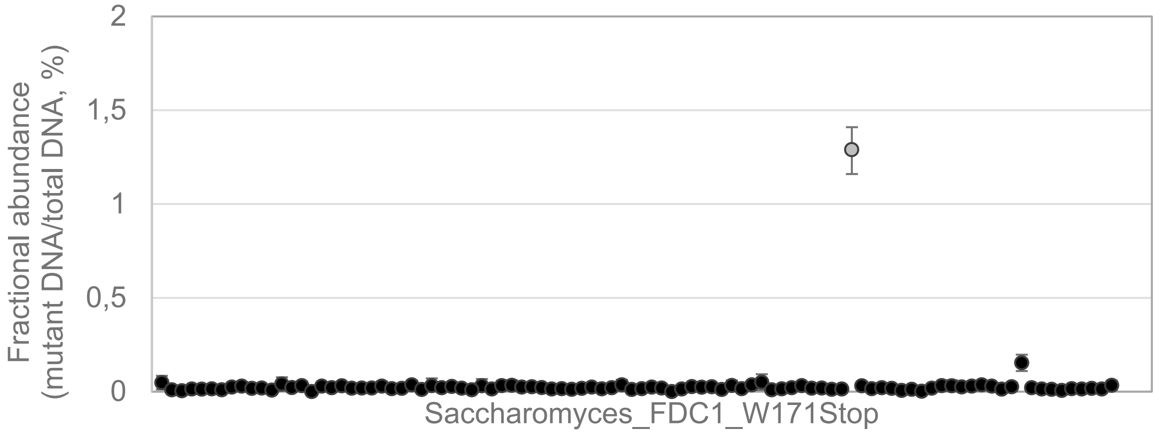


GACGGTGGCAAGTACTTACAAACGTACGGAATGTGGATTCTTCAAACTCCAGATAAAAAATGGACTAATTG
[G/A]TCAATTGCTAGAGGTATGGTTGTAGATGACAAGCATATCACTGGTCTGGTAATTAAACCACAACATAT
TAGACAAATTGCTGACTCTTGGGCAGCAATTGGAAAAGCAAA

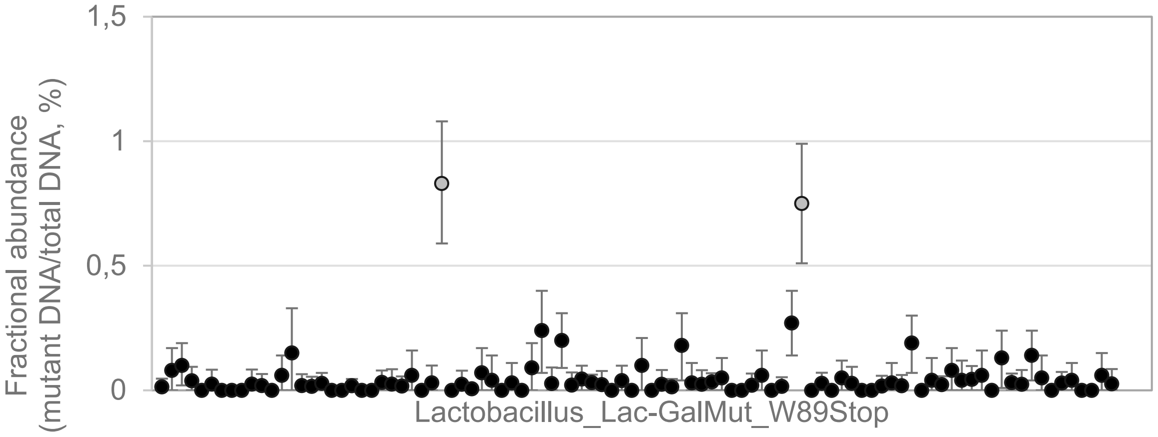


TAGTTGATGATACTACTTCTAAAGAAATTTTAAATGATTTATTGCACGATTTAGCTACGCCAATCCCTAGTG
AGATAAATCGTACTTTTT[G/A]GGAAAAGATGGGGACATCTCCCAAAGAAGCCACTGATTGGTTTTACAACT
TATGTATAAAGAATAATTACATAAAGAAAAAAGCGATTGATAAAAATATTGTTTATGAGGGAACCAGTAGCA

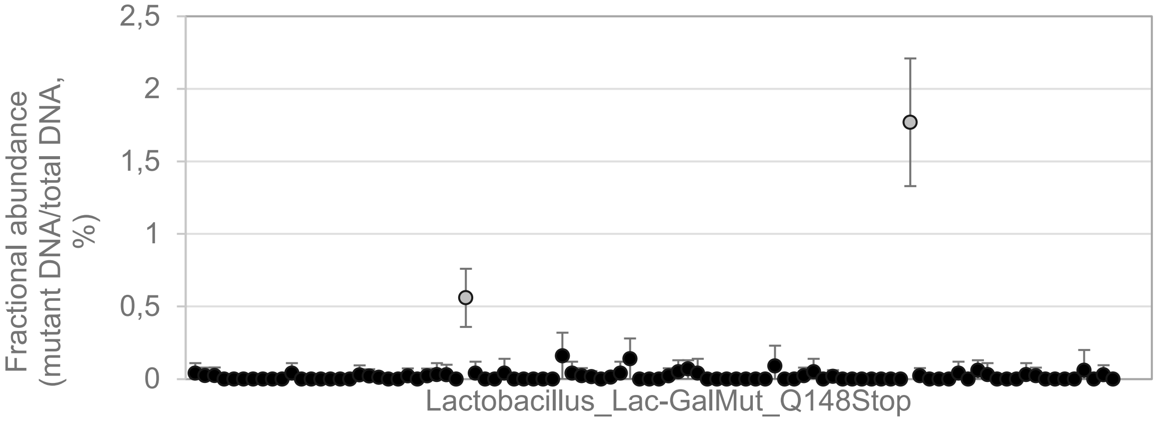


AACTTATGTATAAAGAATAATTACATAAAGAAAAAAGCGATTGATAAAAATATTGTTTATGAGGGAACCAGT
AGCAAGGGGCATGAATTAGAAATTACTATTAACTTGTCAAAGCCAGAAAAAGATCCA[C/T]AATCGATAGCT
AAAGCTGCTCTTATTAAAAATAATGATTATCCAAAATGTGCGCTTTGTCTGGAGAATGAGGGATATCTTG
