## Supplemental notes 3 for "FIND-IT: Ultrafast mining of genome diversity"

### Supplementary Notes 3

**Contents**

Supplementary Notes 3, Section 1. FIND-IT method verification and demonstration of possibilities: modulating plant- and grain morphology by targeting barley knockout variants. 3

Example *GBSS1:* 3

Fig. Supplementary Notes 3.2. Identification and isolation of a novel mutation in the gene *GBSS1* (*gbss1*$^{W126*}$). 5

Fig. Supplementary Notes 3.3. Principal coordinate analysis of isolated FIND-IT variants and respective library parents genotyped using a 50K SNP chip. 6

Fig. Supplementary Notes 3.4. Phenotypic characterization and agronomic evaluation of *gbss1*$^{W126*}$. 7

Example *JMJ706/VRS3*: 7

Fig. Supplementary Notes 3.5. Gene model of barley *JMJ706*. 7

Fig. Supplementary Notes 3.6. Phenotypic characterization and agronomic evaluation of *jmj706*$^{W313*}$ 8

Example *Nud:* 9

Fig. Supplementary Notes 3.7. Gene model of barley *Nud*. 9

Fig. Supplementary Notes 3.8. Phenotypic characterization and agronomic evaluation of *nud*$^{W16*}$ 10

Example *GW2:* 10

Fig. Supplementary Notes 3.9. Natural variation of *GW2* orthologues of barley and rice in barley exome diversity collections and the rice PAN genome. 12

Fig. Supplementary Notes 3.10. Gene model of barley *GW2*. 12

Fig. Supplementary Notes 3.11. Phenotypic characterization and agronomic evaluation of a barley variant carrying the *gw2*$^{W233*}$ allele. 13

Supplementary Notes 3, Section 2. Targeting distinct non-synonymous nucleotide variants as well as variants that modulate transcript abundance demonstrate the strength of FIND-IT for precision breeding. 13

Establishing a mutant framework for improved nitrogen use efficiency and sustainable agriculture of barley green revolution varieties. 13

SLN1/DELLA 14

Fig. Supplementary Notes 3.12. SLN1/DELLA protein in barley with novel variants present in FIND-IT libraries 15

Fig. Supplementary Notes 3.13. Phenotypic characterization and agronomic evaluation of *sln1*$^{A277T}$ 15

GRF4 15

Fig. Supplementary Notes 3.14. Gene model of barley *GRF4*. 16

**Supplementary Notes 3, Section 1. FIND-IT method verification and demonstration of possibilities: modulating plant- and grain morphology by targeting barley knockout variants.**

**Example *GBSS1:***

The biosynthesis of amylose is associated with the activity of granule bound starch synthases. In barley it has been described that variants carrying a loss of function allele of *GBSS1* (*Granule-Bound Starch Synthase I*; HORVU.MOREX.r2.7HG0535060) (Fig. Supplementary Notes 3.1) produce waxy endosperm starch while low expression of this gene results in waxy-like starch phenotypes, due to the absence or reduced content of amylose in the endosperm starch, respectively[1–3].

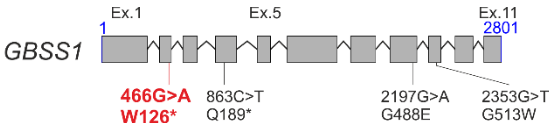

**Fig. Supplementary Notes 3.1. Gene model of barley *GBSS1*.** Position of described mutations causing a waxy phenotype and a novel mutation identified in this study are shown in black and red, respectively.

Using the FIND-IT technology (Online Methods), we targeted and isolated a variant of *GBSS1*, resulting in a premature stop W126* (Fig. Supplementary Notes 3.1) in a library of spring barley cultivar Planet M3 grains that initially were mutagenized with 1.67 mM $NaN_3$. Here we exemplify the isolation procedure in full detail: the genomic DNA library pool identified in phase 1 of the FIND-IT procedure (Fig. Supplementary Notes 3.2a) that contained the mutated *GBSS1* grains originated from a grain pool weighing 280 g. This amounts to approximately 5600 grains assuming an average thousand grain weight (TGW) of 50 g. The identified pool was split into 25% (70 g or approximately 1400 grains) for genomic DNA extraction and screening (phase 1), and 75% (210 g or approximately 4200 grains) of whole grain for variant grain identification (phase 2 and phase 3). From ddPCR analysis a fractional abundance is estimated, which denotes the proportion of the total DNA in a sample that represents the rare allele. A fractional abundance of 0.21% of grain carrying the *gbss1*$^{W126*}$ variant thus corresponds to approximately 8.8 homozygous grains out of 4200 grains in the 75% grain pool kept for variant grain identification.

In phase 2, genomic DNA from approximately 1000 grains from the 75% grain pool (of the corresponding genomic DNA pool identified in phase 1) was sampled non-destructively by drilling a small hole in the endosperm and collecting the flour. Genomic DNA was extracted from the flour sample and screened in pools of 10. The corresponding drilled grains were stored at 4°C. Three wells were identified with a variant fractional abundance of 4.1%, 8.2% and 11.5% (Fig. Supplementary Notes 3.2b). The well with a fractional abundance of 11.5% was chosen for further analysis as this corresponds to the presence of one homozygous grain carrying the *gbss1*$^{W126*}$ variant in the pool of 10 grains.

In phase 3, the 10 grains identified in phase 2 were germinated and genomic DNA was extracted from the first emerging leaf and used for screening. One plant was identified as a homozygous variant as the variant fractional abundance was ~94% (Fig. Supplementary Notes 3.2c). For further validation, the *gbss1*$^{W126*}$ variant was confirmed by Sanger sequencing (Fig. Supplementary Notes 3.2d).

a

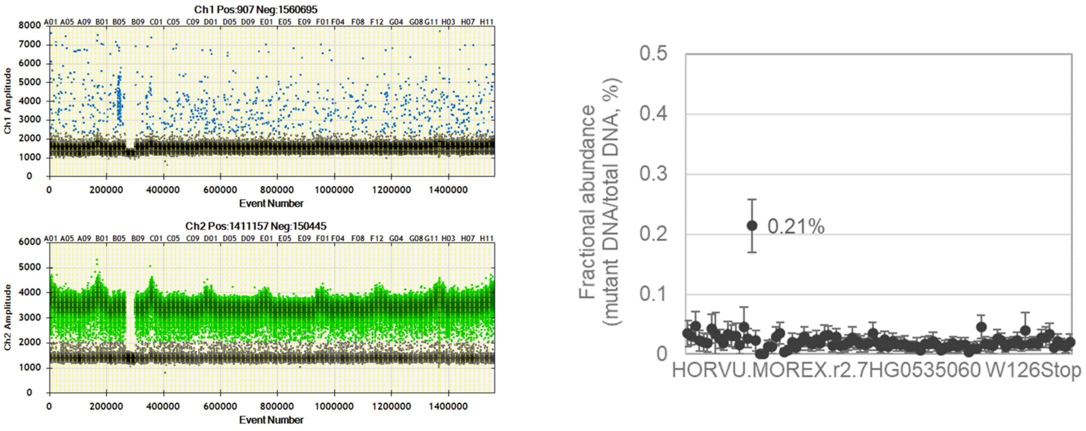

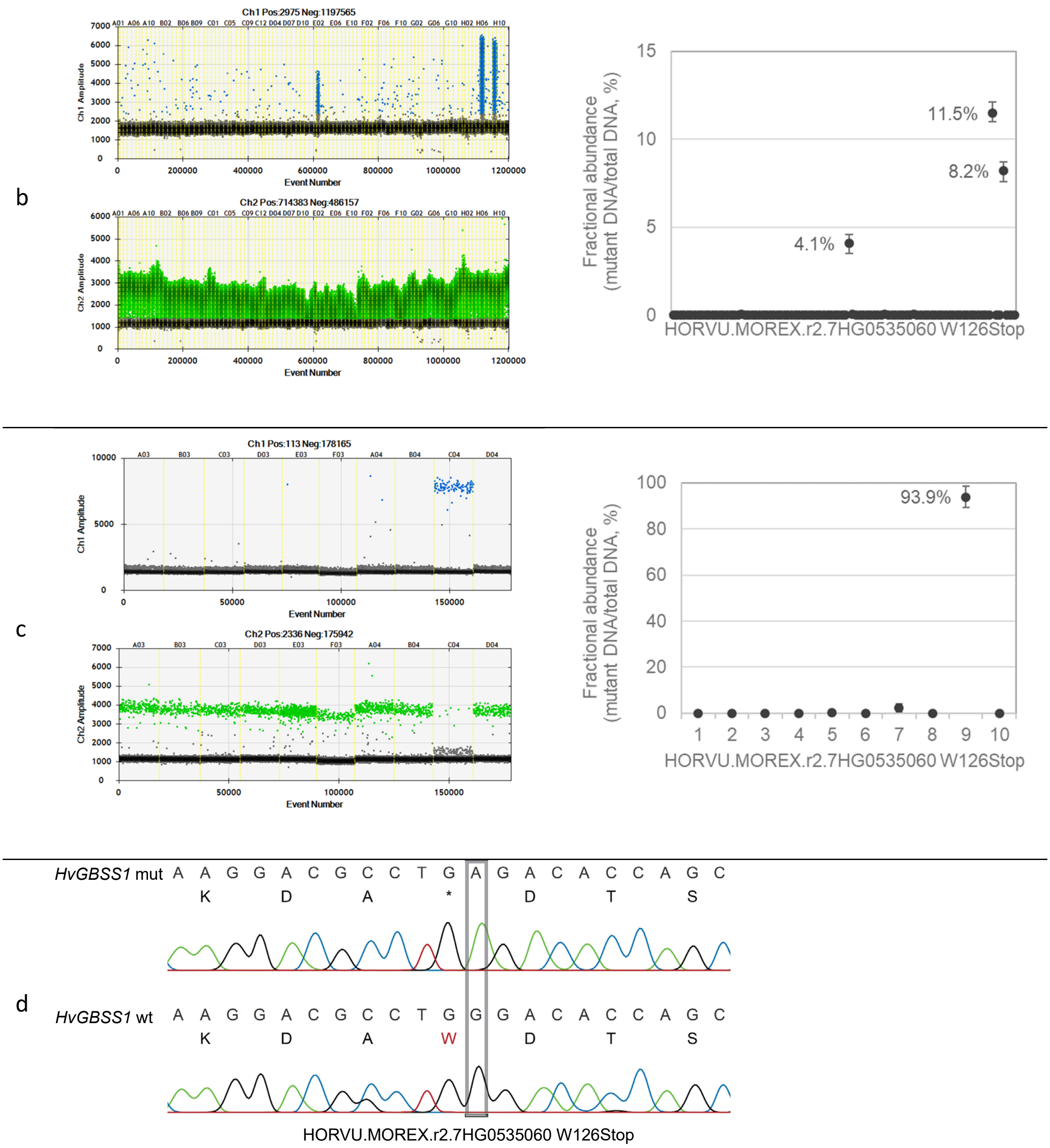

**Fig. Supplementary Notes 3.2. Identification and isolation of a novel mutation in the gene *GBSS1* (*gbss1*$^{W126*}$).** Phase 1- 3 of the FIND-IT procedure. **a**, Phase 1 - Initial library screening. **b**, Phase 2 – Sub-pool screening. **c**, Phase 3 - Confirmation of homozygous variant. **d**, Sequence validation of homozygous variant. Fluorescence amplitude in **a-c**, of variant (blue) and wildtype (green) signals (left) and corresponding fractional abundance of variant DNA copies per total DNA copies (right) in the different phases of FIND-IT screening.

Genetic heritage of isolated homozygous plants was validated to ensure the integrity of variant characterisation before agronomic evaluation. Genotyping original variants using a 50K SNP chip genotyping platform verified that variants isolated from Quench, Paustian or Planet libraries have the respective genetic background (Fig. Supplementary Notes 3.3; Online Methods).

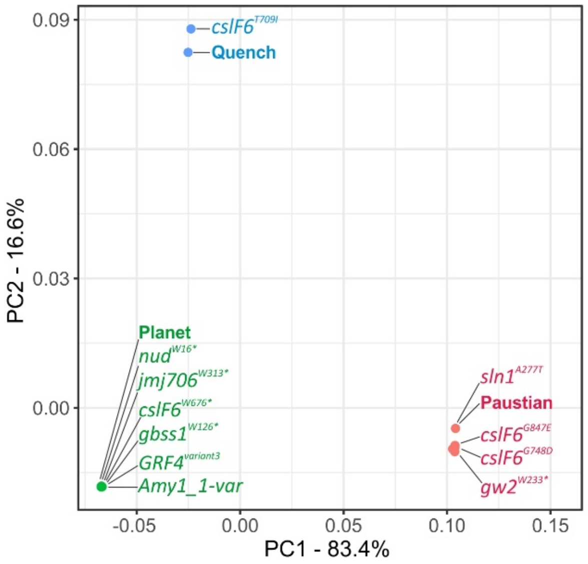

**Fig. Supplementary Notes 3.3. Principal coordinate analysis of isolated FIND-IT variants and respective library parents genotyped using a 50K SNP chip**.

Confirmed homozygous *gbss1*$^{W126*}$ variant plants were grown to maturity in a greenhouse, harvested and then further propagated in field row plots followed by field trial plots for agronomic evaluation (Online Methods). The isolated Planet variant carrying the novel loss of function allele *gbss1*$^{W126*}$ features the waxy barley grain phenotype described by Patron et al. (2002)[1] with the characteristic reduction of amylose in the grain starch (Fig. Supplementary Notes 3.4a,b). When isolated in an elite spring barley background and evaluated on the field, overall TGW is slightly increased (Fig. Supplementary Notes 3.4c) while no large change in grain size was measured (Fig. Supplementary Notes 3.4d,e). Overall grain yield was slightly reduced (Fig. Supplementary Notes 3.4f). Waxy starches are of high value in many processed foods as the amylose content directly influences the texture of cooked starches[4]. Using the FIND-IT technology, waxy variants of common cereals used in the food industries can be directly isolated from elite, high yielding varieties and evaluated on the field for their commercial value and market potential.

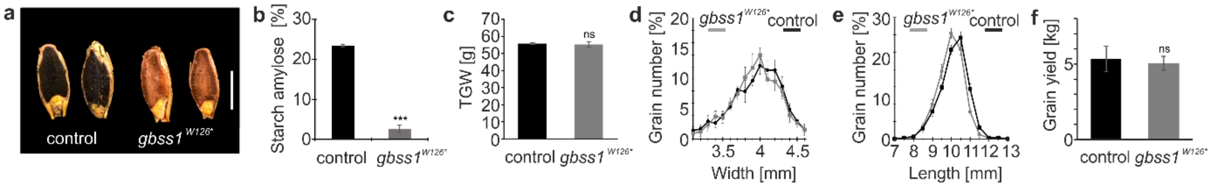

**Fig. Supplementary Notes 3.4. Phenotypic characterization and agronomic evaluation of *gbss1*$^{W126*}$.** **a**, Starch Lugol-staining of half grains. **b**, Endosperm starch amylose content. **c-e**, Thousand grain weight (TGW), grain width distribution and grain length distribution of field propagated lines. **f**, Overall grain yield (7.5 sqm field blots). Control, cv. Planet. Scale bar in a, 5 mm. Error bars are standard diviation, two-tailed t-test was performed to obtain P values (*P<0.05, **P<0.01, ***P<0.001, ns not statistically significant). See Supplementary Table 3 for statistical test details and number of replicates.

##### Example *JMJ706/VRS3*:

The gene *JMJ706/VRS3* (HORVU.MOREX.r2.1HG0041980) is a main controller of the row-type trait in barley inflorescences (spikes)[5]. Bull et al. (2017)[5] identified the barley *VRS3* (*SIX-ROWED SPIKE3*) locus to encode a putative Jumonji C-type H3K9me2/me3 demethylase, involved in the suppression of barley lateral spikelet fertility. Loss of function mutations (Fig. Supplementary Notes 3.5) result in a leaky six-rowed spike phenotype characterized by a gradual filling of lateral spikelets from the bottom to the top end of the spike[5].

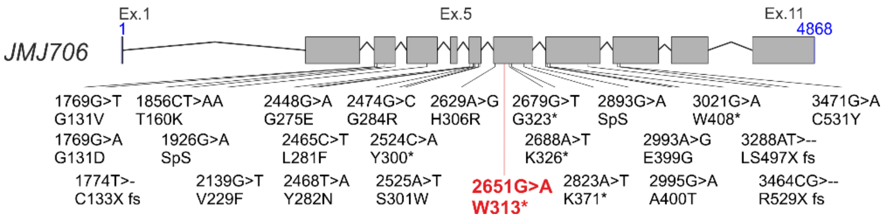

**Fig. Supplementary Notes 3.5. Gene model of barley *JMJ706*.** Position of described mutations and a novel mutation identified in this study are shown in black and red, respectively.

Using the FIND-IT technology (Online Methods), we targeted and isolated a novel allele of *JMJ706/VRS3*, resulting in a premature stop W313* (Fig. Supplementary Notes 3.5) in a library of spring barley cultivar Planet M3 grains that initially were mutagenized with 1.67 mM $NaN_3$. Confirmed homozygous *jmj706*$^{W313*}$ variant plants were grown to maturity in a greenhouse, harvested and then further propagated in field row plots followed by field trial plots for agronomic evaluation (Online Methods). The isolated Planet variant carrying the novel loss of function allele *jmj706*$^{W313*}$ features the barley spikelet phenotype with the characteristic partly six-rowed spike with a gradual increase of the spikelet triplet fertility from the bottom towards the top section of the spike (Fig. Supplementary Notes 3.6a). When used in a modern two-rowed barley background, overall TGW is slightly reduced (Fig. Supplementary Notes 3.6b) while grain size and grain width are similar to the Planet control (Fig. Supplementary Notes 3.6c,d) after seed size sorting above 2.5 mm diameter (here, the generally smaller grains derived from the lateral spikelets, as seen in Fig. Supplementary Notes 3.6a, were sorted out as a standard procedure before agronomic evaluation). As reported in Bull et al., (2017)[5], loss of function *jmj706* alleles show their full agronomic potential in six-rowed barley background, resulting in desired overall yield increase and seed size uniformity. Using FIND-IT, novel alleles of row-type genes can be directly isolated from two- and six-rowed barley libraries to finetune inflorescence architecture.

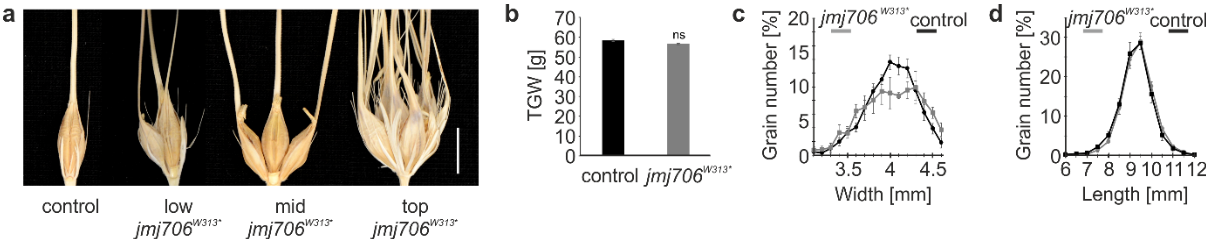

**Fig. Supplementary Notes 3.6. Phenotypic characterization and agronomic evaluation of *jmj706*$^{W313*}$. a**, Spikelet phenotype with a partly six-rowed spike phenotype; lateral spikelets in lower spike section are infertile or partly fertile with increasing fertility towards the top spike section; top triplets can have additional fertile spikelets[5]. **b-d**, Thousand grain weight (TGW), grain width distribution and grain length distribution of field propagated lines. Control, cv. Planet. Scale bar in a, 5 mm. Error bars are standard diviation, two-tailed t-test was performed to obtain P values (*P<0.05, **P<0.01, ns not statistically significant). See Supplementary Table 3 for statistical test details and number of replicates.

**Example *Nud:***

The *Nud* gene controls the covered/naked caryopsis trait in barley. *Nud* is an ethylene response factor (ERF) family transcription factor (HORVU.MOREX.r2.7HG0596820) with homology to the Arabidopsis WIN1/SHN1 transcription factor gene[6]. *Nud* regulates a lipid biosynthesis pathway which at grain maturity forms a lipid layer on the pericarp epidermis that is mandatory for hull adhesion. Taketa et al. (2008)[6] and Yu et al. (2016)[7] described natural and induced loss-of-function alleles of *Nud* (Fig. Supplementary Notes 3.7, shown in black font), including a naturally selected deletion variant *nud1.a*, that are missing the respective lipid layer to adhere the hull. Thus, threshing of mature grain results in a naked/hulless barley grain phenotype.

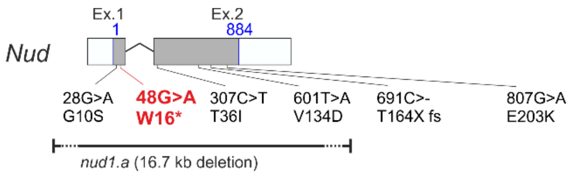

**Fig. Supplementary Notes 3.7. Gene model of barley *Nud*.** Position of described mutations and a novel mutation identified in this study are shown in black and red, respectively. In the naturally occurring variant *nud1.a*, the *Nud* gene is deleted.

Using the FIND-IT technology (Online Methods), we targeted and isolated a novel variant of *Nud*, carrying a premature stop W16* (Fig. Supplementary Notes 3.7), in a library of spring barley cultivar Planet M3 grains that initially were mutagenized with 1.67 mM $NaN_3$. Confirmed homozygous $nud^{W16*}$ variant plants were grown to maturity in a greenhouse, harvested and further propagated in field row plots followed by field trial plots for agronomic evaluation (Online Methods). The isolated Planet variant carrying the novel loss of function allele $nud^{W16*}$ features the characteristic hulless barley grain phenotype (Fig. Supplementary Notes 3.8a). As a result, both grain length and width and thus TGW and overall yield are reduced (Fig. Supplementary Notes 3.8b-e).

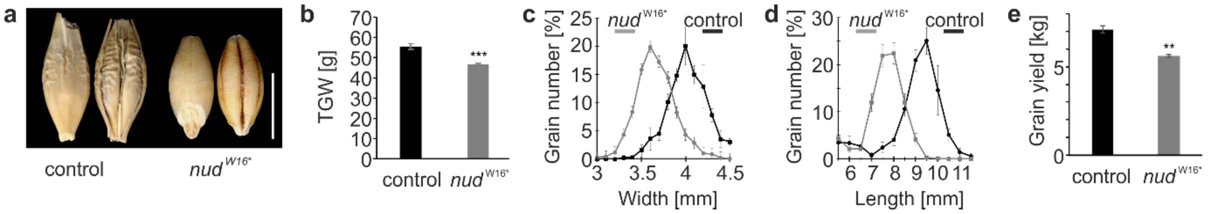

**Fig. Supplementary Notes 3.8. Phenotypic characterization and agronomic evaluation of *nud*$^{W16*}$.** **a**, Mature barley grain after threshing with attached (control) and de-attached hull (*nud*$^{W16*}$). **b**, Thousand grain weight (TGW), **c**, grain width distribution and **d**, grain length distribution of field propagated lines. **e**, Overall grain yield (7.5 sqm field blots). Control, cv. Planet. Scale bar in a, 5 mm. Error bars are standard diviation, two-tailed t-test was performed to obtain P values (*P<0.05, **P<0.01, ***P<0.001, ns not statistically significant). See Supplementary Table 3 for statistical test details and number of replicates.

This known domestication trait, here isolated in a modern elite barley, makes barley suitable for human consumption. Our agronomic evaluation shows the obvious yield reduction for farmers (as hulls are separated from the grain in the harvest process), that needs to be considered for trait market value evaluation in comparison to elite hulled barley varieties.

**Example *GW2:***

Song et al. (2007)[8] reported modifications in a *RING-type E3 ubiquitin ligase* gene to be causative for the quantitative trait locus (QTL) *Grain Width 2* (*OsGW2*) modulating grain width and weight in rice. Loss of *GW2* function in the varieties WY3 and Oochikara (via a 1-bp deletion in the *GW2* gene causing a premature stop codon) was associated with increased grain width and weight. The barley reference genome harbors a *RING-type E3 ubiquitin ligase*-like gene (HORVU.MOREX.r2.6HG0483720) in the peri-centromeric region on chromosome 6H which likely represents the barley *GW2* orthologue.

When screening available barley exome datasets of diverse collections of wild barley, landraces and modern cultivars for variation in *GW2* (267 and 371 diverse barley accessions, respectively;[9,10]), a minor set of six natural alleles was identified (average of 9.4 alleles per 1,000 accessions). These alleles represent five amino acid exchanges and one deletion of the residue Pro268 which is not found in the rice peptide sequence, and thus most likely of minor importance for protein function. However, no premature stop codons or larger deletions likely representing loss of function alleles as known in rice were present (Fig. Supplementary Notes 3.9). When screening the rice PAN genome, containing 3010 diverse accessions[11], a larger set of 42 natural *GW2* alleles was found for rice (average of 13.95 alleles per 1,000 accessions), including a point mutation causing the change of Trp234 to a premature stop (W234*; Fig. Supplementary Notes 3.9).

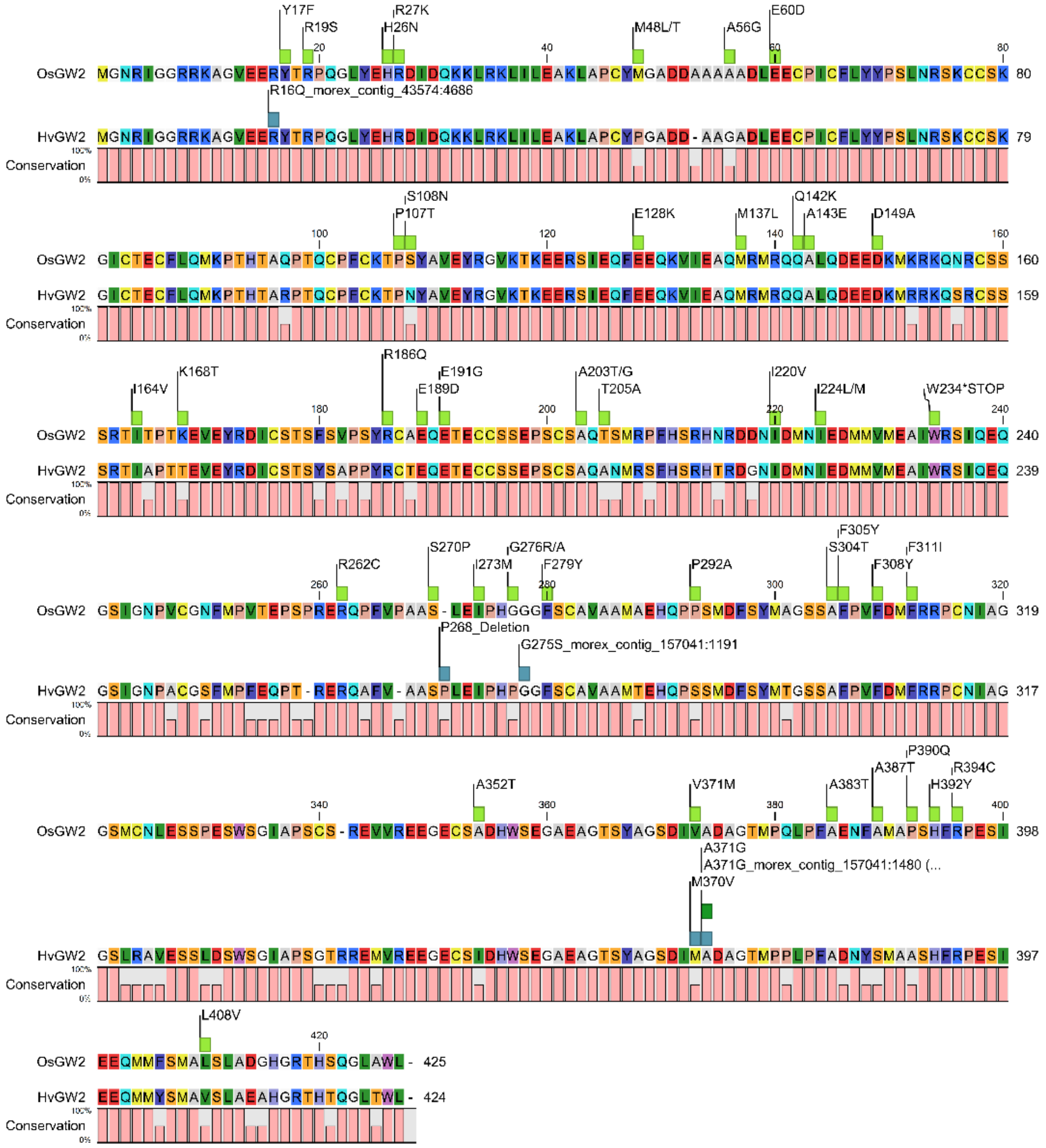

**Fig. Supplementary Notes 3.9. Natural variation of *GW2* orthologues of barley and rice in barley exome diversity collections and the rice PAN genome.**

A recent barley GWAS (genome-wide association scan) study of grain yield traits, identified the chromosome 6H peri-centromeric region, harboring *GW2*, as a yield-associated QTL hotspot but suggested a genome editing or TILLING approach to verify the actual contribution of barley *GW2* to the control of grain width, grain length, TGW or grain yield per area[12]. Using the FIND-IT technology (Online Methods), we mimicked the rice natural loss of function allele $gw2^{W234*}$ by targeting the corresponding loss of function allele $gw2^{W233*}$ in an elite malting barley (Fig. Supplementary Notes 3.10). This novel allele of *GW2* was identified and isolated in a library of spring barley cultivar Paustian M2 grains that initially were mutagenized with 0.3 mM $NaN_3$. Confirmed homozygous $gw2^{W233*}$ variant plants were grown to maturity in a greenhouse, harvested and then further propagated in field row plots followed by field trial plots for agronomic evaluation (Online Methods).

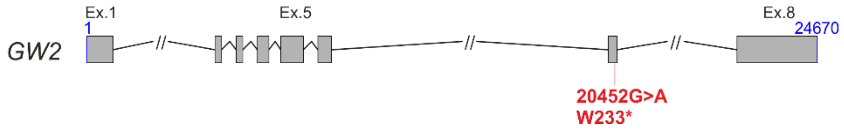

**Fig. Supplementary Notes 3.10. Gene model of barley *GW2*.** Position of a novel FIND-IT identified and isolated induced variation is shown in red.

Increasing yield per area is a main cereal breeding target. The rice QTL *Grain Width 2* (*OsGW2*), encoding a *RING-type E3 ubiquitin ligase* gene, and respective orthologues were recently published as promising gene targets to increase cereal yield[8,13]. While natural and induced loss of function alleles of the *GW2* locus in rice and wheat were identified and selected by breeders, no allele was reported for barley yet. Using FIND-IT, we isolated the first *GW2* loss of function allele in barley by mimicking a natural variant identified in a rice landrace. Here, we show in field yield trials that a loss of GW2 protein function in elite European spring malting barley, like in rice, results in increased thousand grain weight due to wider grain size (Fig. Supplementary Notes 3.11a-d). However, when tested in Danish fields in season 2020, yield per area was not increased (Fig. Supplementary Notes 3.11e). Song et al. (2007) reported that loss of GW2 function caused accelerated grain milk filling rate. This might result in a yield advantage in suboptimal growth conditions during grain filling. For instance, heat stress shortens grain filling resulting in yield losses through decreased starch deposition per grain[14]. Variants with loss of function alleles might partly compensate this through accelerated grain filling. The novel barley GW2 loss of function allele isolated here, can be tested in different climatic environments to improve grain yield per area.

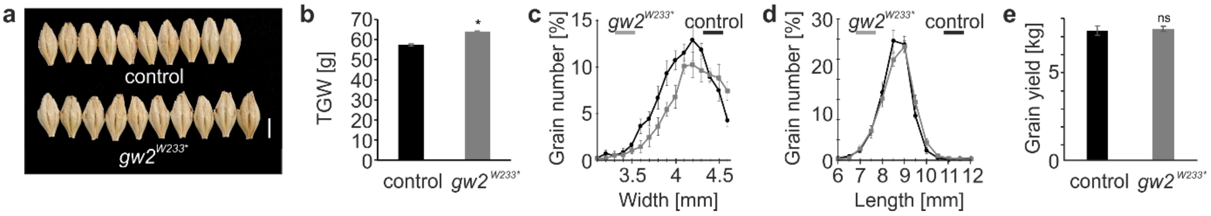

**Fig. Supplementary Notes 3.11. Phenotypic characterization and agronomic evaluation of a barley variant carrying the *gw2*$^{W233*}$ allele. a**, Size of 10 randomly picked mature barley grain after threshing. **b**, Thousand grain weight, **c**, grain width distribution and **d**, grain length distribution of field propagated lines. **e**, Overall grain yield (7.5 sqm field blots). Control, cv. Paustian. Scale bar in a, 5 mm. Error bars are standard diviation, two-tailed t-test was performed to obtain P values (*P<0.05, **P<0.01, ns not statistically significant). See Supplementary Table 3 for statistical test details and number of replicates.

#### Supplementary Notes 3, Section 2. Targeting distinct non-synonymous nucleotide variants as well as variants that modulate transcript abundance demonstrate the strength of FIND-IT for precision breeding.

##### Establishing a mutant framework for improved nitrogen use efficiency and sustainable agriculture of barley green revolution varieties.

Advances in agricultural practices during the green revolution in the 1960s boosted crop plant growth and cereal productivity through fertilizer application paired with plant height control via the introduction of natural and induced variants of semi-dwarfing genes[15]. In wheat, variants of the *Reduced height-1* (*Rht-1*) locus, encoding the gibberellic acid (GA) signaling repressor DELLA, reduce the sensitivity to the GA growth promoting signal[16]. In rice and barley, variants of the GA biosynthesis gene *GA20ox2* lower GA hormone abundance and increase DELLA accumulation[17,18].

These short-strawed variants form the genetic basis of green revolution varieties of wheat, rice and barley used today[19]. Both strategies cause excess DELLA repression that inhibits plant growth but also impair plant nitrogen uptake, thereby necessitating increased use of nitrogen fertilizers[19].

This excess application of nitrogen fertilizers negatively impacts the environment, increase farming costs, and reduces the utilization of green revolution varieties in developing countries. A recent study in rice identified variants of *GROWTH-REGULATING-FACTOR 4* (*GRF4*) that increase nitrogen uptake in cereal green revolution varieties to overcome the undesired inhibitory effects of DELLA on this parameter[19], thus constituting a promising breeding target for sustainably grown cereal crops.

In barley, green revolution genes *GA20ox2* and *DELLA* are encoded by loci *SEMIDWARF1* (*Sdw1*; HORVU.MOREX.r2.3HG0256590) and *SLENDER1* (*SLN1;* HORVU.MOREX.r2.4HG0280720), respectively [18,20]. An orthologue of rice *GRF4* is present in the barley reference genome (HORVU.MOREX.r2.2HG0160430). Sequencing identified the *GA20ox2* loss-of-function allele in barley cvs. Paustian and Planet (*sdw1.d;*[18]) verifying these elite malting lines as green revolution varieties and as suitable backgrounds. Accordingly, we screened our Paustian and Planet FIND-IT libraries for their potential to harness diverse *DELLA* and *GRF4* variants suitable for breeding strategies to balance growth suppression and nitrogen uptake. We targeted *i*) *SLN1/DELLA* for non-synonymous variants (to lower DELLA repression) and *ii*) *GRF4* for variants with increased transcript abundance via *miR396*-regulation interference.

**SLN1/DELLA:** Targeted screening of the libraries of the spring barley cultivars Planet (M3 grains, initially mutagenized with 1.67 mM $NaN_3$) or Paustian (M2 grains, initially mutagenized with 0.3 mM $NaN_3$) identified 31 variants of SLN1/DELLA with specific amino acid exchanges (Fig. Supplementary Notes 3.12). A single variant, $sln1^{A277T}$, was isolated from the Paustian library, grown to maturity and further propagated to test and evaluate the effects of DELLA fine-regulation in elite green revolution varieties of barley (Online Methods).

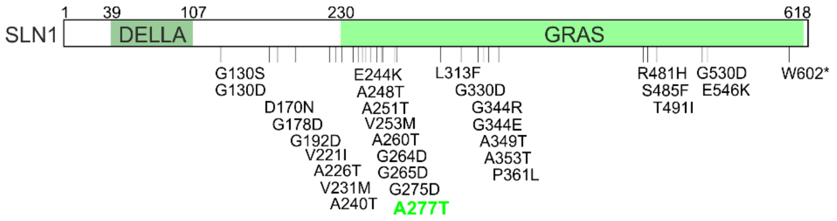

**Fig. Supplementary Notes 3.12. SLN1/DELLA protein in barley with novel variants present in FIND-IT libraries.** Functional DELLA and GRAS domains with positions of identified amino acid exchange variants are indicated.

Interestingly, the *sln1*$^{A277T}$ variant showed no significant change in plant height (Fig. Supplementary Notes 3.13a), combined with increased TGW likely due to elongated grains, while yield per area was constant and not decreased (Fig. Supplementary Notes 3.13b-e). Our results indicate that fine-modulating DELLA abundance in green revolution varieties does not reset the agronomic performance to pre-green revolution times and presents a possible way forward to test and improve nitrogen use efficiency of modern malting barley. FIND-IT libraries contain multiple variants of *DELLA* that can be tested alone or in combination with *GRF4* variants in breeding programs.

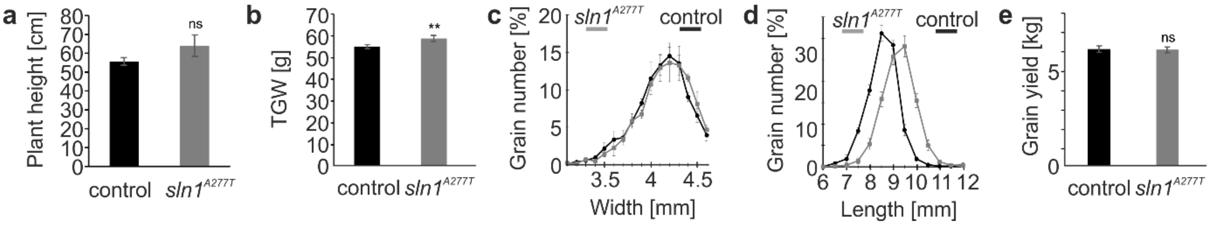

**Fig. Supplementary Notes 3.13. Phenotypic characterization and agronomic evaluation of *sln1*$^{A277T}$. a**, Plant height, **b**, thousand grain weight, **c**, grain width distribution and **d**, grain length distribution of field propagated lines. **e**, Overall grain yield (7.5 sqm field blots). Control, cv. Paustian. Error bars are standard diviation, two-tailed t-test was performed to obtain P values (*P<0.05, **P<0.01, ns not statistically significant). See Supplementary Table 3 for statistical test details and number of replicates.

**GRF4:** To enable future breeding strategies to improve nitrogen uptake in green revolution varieties, we targeted the 21-bp *miR396*-binding site of barley *GRF4.* In rice *GRF4*, mutations in this binding site (haplotype *OsGRF4*$^{ngr2}$, Fig. Supplementary Notes 3.14;[19]) interfere with the miR396-mediated cleavage of the *GRF4* transcript, causing increased GRF4 protein levels and resulting in the desired improved nitrogen use efficiency. We identified three novel nucleotide substitutions within the 21-bp *miR396*-binding site of barley *GRF4* in our Planet library (M3 grains, initially mutagenized with 1.67 mM $NaN_3$) (Online Methods) (Fig. Supplementary Notes 3.14). Seeds of these variants can be isolated for propagation and assessment of their potential in improving nitrogen uptake and agronomic performance of elite green revolution varieties. Best candidates can additionally be used in crosses with novel DELLA variants or elite non-GA green revolution varieties (e.g. *DEP1* semi-dwarf varieties[21]) to breed a new type of green revolution varieties with improved nitrogen metabolism. This shows that FIND-IT provides an efficient and targeted approach for breeders and scientists to modulate protein function.

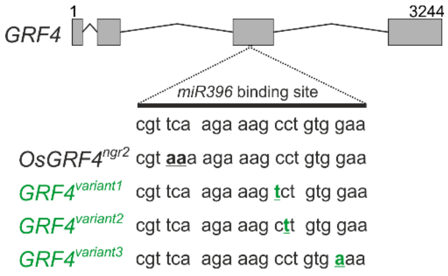

**Fig. Supplementary Notes 3.14. Gene model of barley *GRF4*.** The 21-bp *miR396* binding site in exon 3 is shown including a known double substitution in rice (*OsGRF4*$^{ngr2}$) and three novel barley variants (in green).

#### Targeting a promoter variant of barley α-amylase to enhance gene expression and seed germination vigour.

Starch is the major storage polysaccharide in cereal grains and its degradation is central to early seedling vigour, and hence to crop productivity. An efficient starch degradation is also a major determinant of cereal malt quality and optimization of the respective amylolytic enzyme levels is a key breeding target for cereal crop improvement[22].

The (1,4)-α-glucan endohydrolase, α-amylase, is a main starch degrading enzyme which hydrolyses the internal (1,4)-α-glucosyl linkages. α-Amylase enzymes are encoded by 12 genes in the Morex barley reference genome[23], and their expression, after initiation of seed germination, is tissue specific[24], and tightly regulated by transcriptional repressors and activators[25]. Several cis-acting elements are known in the promoters of α-amylase genes, including the W-box/O2S binding domains for the WRKY38 repressor (Fig. Supplementary Notes 3.15a).

The WRKY38 transcription factor was shown to interfere with the binding of GAMYB, the transcriptional activator of α-amylase expression[25]. Transcript levels of WRKY38 are high in aleurone tissues of mature barley grain but quickly decrease upon the initiation of grain germination in a time- and tissue-dependent manner, while at the same time, the expression of α-amylase genes increases. Here, transcripts of the single *Amy1_2* gene increase to the highest levels, followed by transcripts of the *Amy1_1* and *Amy2* gene cluster members[26].

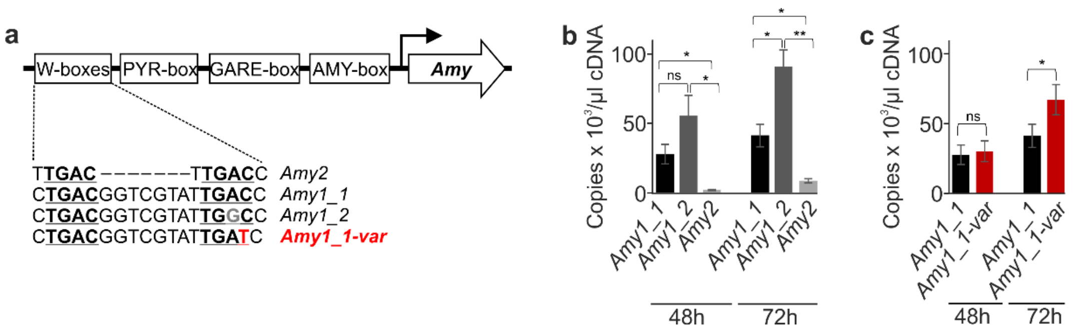

**Fig. Supplementary Notes 3.15. Known cis-acting elements in α-amylase promoters with natural and induced W-box sequence variation. a**, TGAC core motifs in W-boxes are underlined and in bold, natural variation in the *Amy1_2* gene in grey and a novel, induced variation in an *Amy1_1* gene in red. **b**, Expression of α-amylase genes in Planet grain 48h and 72h after initiation of germination. **c**, Expression of *Amy1_1* genes in Planet grain (black) and Planet variant *Amy1_1-var* (red) 48h and 72h after initiation of germination. Error bars are standard diviation, two-tailed t-test was performed to obtain P values (*P<0.05, **P<0.01, ns not statistically significant). See Supplementary Table 3 for statistical test details and number of replicates.

Genes in the *Amy1_1* and *Amy2* clusters have a tandem repeat W-box with two TGAC core motifs in common (Fig. Supplementary Notes 3.15a, shown underlined and bold; [23,25]) which in *Amy1* genes are separated by a short linker sequence that do not affect WRKY38 binding activity[27]. The single

*Amy1_2* gene however, with the strongest α-amylase expression, has a modified tandem repeat W-box with one core motif changed from TGAC to TGGC (Fig. Supplementary Notes 3.15a, shown in grey), which might decrease the WRKY38 binding affinity to the *Amy1_2* promoter and thus decrease the repression of transcription.

To test our hypothesis that variations in the TGAC WRKY38 repressor binding motif could contribute to superior α-amylase expression, we identified and isolated a novel allele of a single *Amy1_1* gene with a single nucleotide substitution in the tandem repeat W-box changing one of the TGAC core motif to TGAT (Fig. Supplementary Notes 3.15a, shown in red) in a library of spring barley cultivar Planet M3 grains that initially were mutagenized with 1.67 mM $NaN_3$. Confirmed homozygous variant plants harboring the *Amy1_1-var* gene variant, were grown to maturity in a greenhouse, harvested and then further propagated in field row plots followed by field trial plots for further characterization (Online Methods).

For testing α-amylase expression, grains of control Planet and the novel barley variant *Amy1_1-var* were germinated as described in Betts et al. (2017) and transcription levels were captured using quantitative ddPCR with probes specific for the *Amy1_2* gene, for genes in *Amy1_1* cluster or for genes in the *Amy2* cluster (Online Methods). Here, and as described in Betts et al. (2020), expression of the *Amy1_2* gene in the Planet control exceeded the expression of genes in the *Amy1_1* or *Amy2* clusters showing the integrity of the test system (Fig. Supplementary Notes 3.15b). We then determined that the overall *Amy1_1* transcript level in the novel Planet variant *Amy1_1-var* was significantly increased 72h after initiation of germination (Fig. Supplementary Notes 3.15c). This result for the first time confirms the function of the W-box in α-amylase promoters *in vivo* and highlights the possibilities of modifying gene transcript levels by targeting promoter domains using FIND-IT for gene function analyses and cereal breeding purposes.

**Targeting variants of *CslF6* for modification of cereal grain quality parameters.**

Low (1,3;1,4)-β-glucan is a preferred grain trait for the distilling and brewing industries as high (1,3;1,4)-β-glucan levels in the wort can cause filtration difficulties and undesirable hazes in the final product[28]. The *cellulose synthase like* gene *CslF6* (HORVU.MOREX.r2.7HG0579230) is a key determinant controlling the biosynthesis and structure of (1,3;1,4)-β-glucan in barley grains[29]. Knockout of *CslF6* identified by either screening for low (1,3;1,4)-β-glucan or by utilising CRISPR-Cas9, showed detrimental effect on grain yield and TGW in greenhouse grown variants [29,30]. A knockdown mutant of *CslF6*, leading to 70% reduction in (1,3;1,4)-β-glucan levels in the grain, showed no morphological difference to wild-type. However, here grain hardness was lowered leading to increased rates of broken grains after threshing [31], lowering grain and malt quality. No yield data was presented by the authors[31].

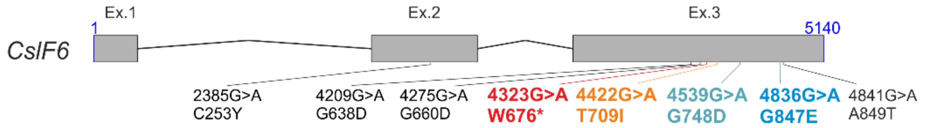

**Fig. Supplementary Notes 3.16. Gene model of barley *CslF6*.** Position of described mutations as well as novel mutations identified in this study are shown in black and colour, respectively.

To test the hypothesis that FIND-IT could be used to obtain applicable low (1,3;1,4)-β-glucan lines via *CslF6* gene variation, we targeted *CslF6* for a premature stop codon as proof of concept, and additionally for more delicate SNPs to identify and isolate knockdown versions.

**Knocking out *CslF6* results in grain with ultralow (1,3;1,4)-β-glucan content and unacceptably low yield potential in modern malting barley.**

As proof of concept we targeted and isolated a novel knockout allele of *CslF6*, causing a premature stop W676* (Fig. Supplementary Notes 3.16), in our library of spring barley cultivar Planet M2 grains that initially was mutagenized with 0.3 mM $NaN_3$ (Online Methods). Confirmed homozygous *cslF6*$^{W676*}$ variant plants were grown to maturity in a greenhouse, harvested and then further propagated in field row plots followed by field trial plots for agronomic evaluation (Online Methods).

The isolated Planet variant carrying the novel loss of function *cslF6*$^{W676*}$ allele had very low (1,3;1,4)-β-glucan content as shown by (1,3;1,4)-β-glucan Calcofluor staining and quantification by HPAEC-PAD (Fig. Supplementary Notes 3.17a,b). Field-testing for agronomic performance showed that *cslF6*$^{W676*}$ TGW was severely reduced (Fig. Supplementary Notes 3.17c) mainly due to a narrow grain morphology (Fig. Supplementary Notes 3.17d,e), which confirms published greenhouse experiments[30]. Most importantly, the grain yield at harvest was reduced by 35% (Fig. Supplementary Notes 3.17e), detrimental to any applicability of the line.

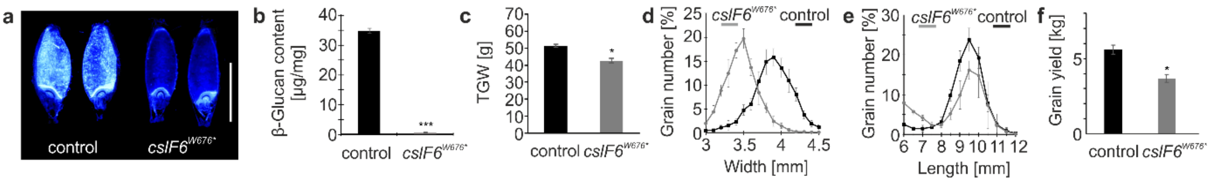

**Fig. Supplementary Notes 3.17. Phenotypic characterization and agronomic evaluation of allele *cslF6*$^{W676*}$. a**, (1,3;1,4)-β-Glucan Calcofluor staining of mature grain. **b**, (1,3;1,4)-β-Glucan content of mature grain. **c**, Thousand grain weight, **d**, Grain width distribution and **e**, Grain length distribution of field propagated lines. **f**, Overall grain yield (7.5 sqm field blots). Control, cv. Planet. Scale bar in a, 5 mm. Error bars are standard diviation, two-tailed t-test was performed to obtain P values (*P<0.05, **P<0.01, ns not statistically significant). See Supplementary Table 3 for statistical test details and number of replicates.

##### Specific amino acid exchanges in CslF6 result in barley grain with lowered (1,3;1,4)-β-glucan content, unchanged yield in elite malting barley.

Using the FIND-IT technology (Online Methods), we aimed to identify amino acid exchange variants of CslF6, featuring reduced (1,3;1,4)-β-glucan grain content without any detrimental effect on agronomical performance. We targeted and isolated three *CslF6* variants (Fig. Supplementary Notes 3.16) that carried amino acid exchanges in either close proximity to the TED motif that controls the (1,3;1,4)-β-glucan fine structure[32] (Fig. Supplementary Notes 3.18; variant T709I) or within CslF6 transmembrane helices (TMH) (Fig. Supplementary Notes 3.18; variant G748D in TMH4 and variant G847E in TMH6).

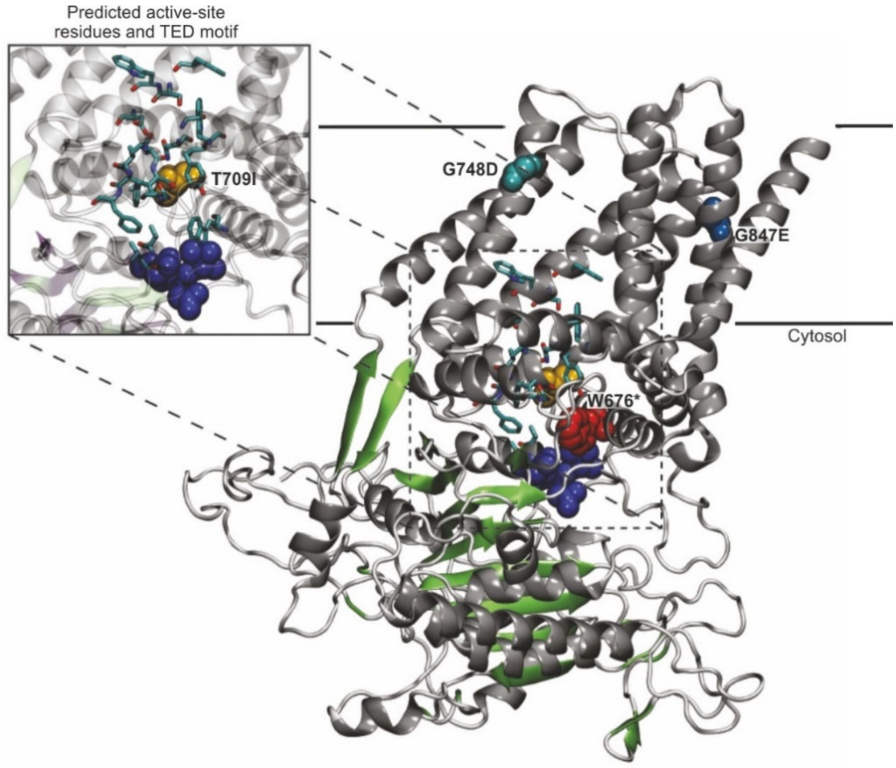

**Fig. Supplementary Notes 3.18. Protein homology model of CslF6 based on poplar cellulose synthase isoform 8 (PDB# 6wlb.1.A).** Amino acid positions at residue W676, T709, G748 and G847 are shown as spheres (red, yellow, turquoise, blue, respectively). Insert shows glucose binding sites - predicted by alignment of CslF6 and poplar cellulose synthase isoform 8 - as light blue sticks and the TED motif is shown as spheres (dark blue).

Intermediate amounts of (1,3;1,4)-β-glucan were detected in all three variants (Fig. Supplementary Notes 3.19a). In contrast to the knockout variant, TGW was not or little affected in *cslF6*$^{G748D}$, *cslF6*$^{G847E}$ and *cslF6*$^{T709I}$ (Fig. Supplementary Notes 3.19b). All three variants showed slightly narrower grain morphology (Fig. Supplementary Notes 3.19c,d). Stability of the mature grain at harvest and grain breakage in hard mechanical threshing was changed in variants *cslF6*$^{G847E}$ and *cslF6*$^{T709I}$. Overall grain yields were, as TGW, unaltered for *cslF6*$^{G748D}$,*cslF6*$^{G847E}$ and *cslF6*$^{T709I}$ (Fig. Supplementary Notes 3.19f).

Whether the slightly bigger decrease in (1,3;1,4)-β-glucan levels in *cslF6*$^{T709I}$, compared to the other two variants, induced the TGW penalty observed cannot be concluded from the present data.

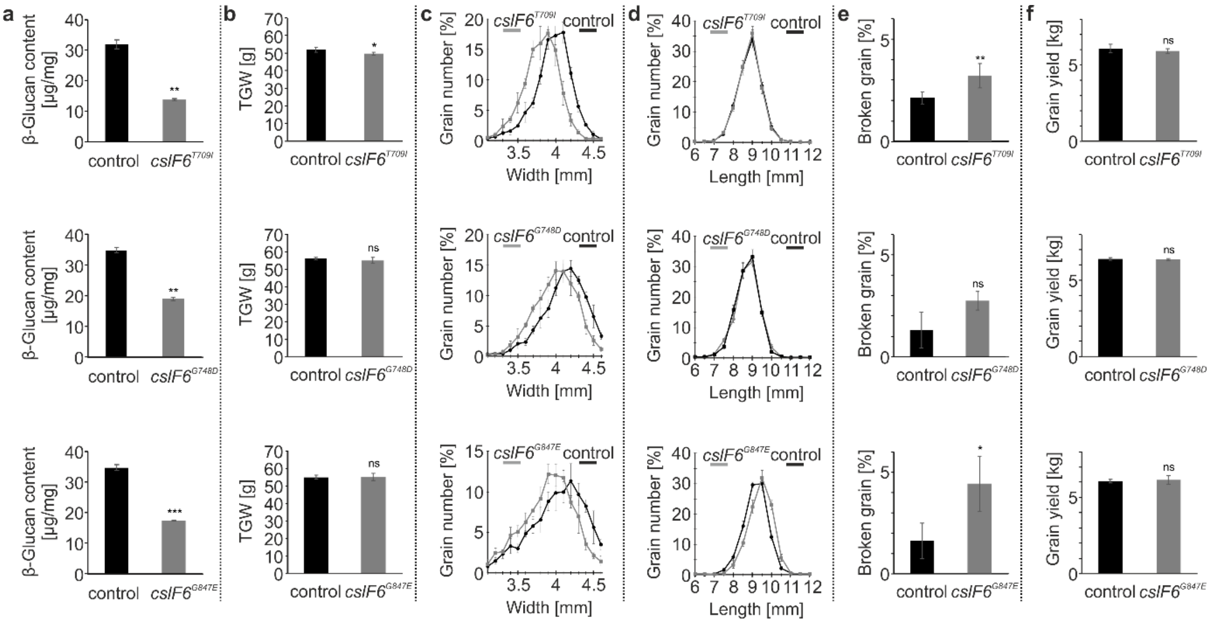

**Fig. Supplementary Notes 3.19. Grain characterization and agronomic evaluation of variants *cslF6*$^{T709I}$, *cslF6*$^{G748D}$ or *cslF6*$^{G847E}$. a**, (1,3;1,4)-β-Glucan content of mature grain. **b**, Thousand grain weight (TGW), **c**, Grain width distribution and **d**, Grain length distribution of field propagated lines. **e**, Percentage of broken grain after threshing. **f**, Overall grain yield (7.5 sqm field blots). Control *cslF6*$^{G748D}$ or *cslF6*$^{G847E}$, cv. Paustian; Control *cslF6*$^{T709I}$, cv. Quench. Error bars are standard diviation, two-tailed t-test was performed to obtain P values (*P<0.05, **P<0.01, ***P<0.001, ns not statistically significant). See Supplementary Table 3 for statistical test details and number of replicates.

Using the FIND-IT technology, we generated and identified four novel variants with non-synonymous base changes in the barley *CslF6*. While the *CslF6* knockout line as expected will be of limited usefulness for malting and brewing applications because of the yield penalty, *cslF6*$^{T709I}$, *cslF6*$^{G748D}$ and *cslF6*$^{G847E}$ showed promising results. Especially *cslF6*$^{G748D}$ was unaltered in yield, in TGW and in grain breakage in hard mechanical threshing but had a 50% reduction in (1,3;1,4)-β-glucan levels, making it a prime candidate gene variation to be included in elite malting lines.

**Supplementary Notes 3, Section 3. Beyond barley, using FIND-IT to identify loss-of-function variants in wheat, yeast, and bacterial libraries.**

**Wheat:**

In wheat, we screened a winter wheat library (supplementary Notes 1, Section 1) and targeted specific stop codons in the three sub-genome copies of the *TaMLO* gene to introduce broad spectrum resistance to the fungal pathogen, *Erisiphe graminis* f. sp. hordei, the causal agent of powdery mildew[33]. We identified TaMlo-5AL$^{W331Stop}$, TaMlo-4BL$^{W358Stop}$ and TaMlo-4DL$^{W358Stop}$ (Supplementary Notes 2, Section 2). The three TaMLO variant lines can now be isolated, crossed and directly evaluated on the field for yield and disease performance.

**Yeast:**

To demonstrate the applicability of FIND-IT for microbial libraries we screened a *Saccharomyces cerevisiae* library (Supplementary Notes 1, Section 1) for specific loss-of-function variants of the gene *ferulic acid decarboxylase* (*FDC1*). The FDC1 enzyme converts aromatic carboxylic acids such as ferulic acid from malt to their corresponding vinyl derivatives, which often impart undesirable clove-like off-flavours in beer[34]. We identified and isolated two strain-variants with premature stop codons FDC1$^{W159*}$ and FDC1$^{W171*}$ (Supplementary Notes 2, Section 2) that showed the expected growth inhibition phenotype when exposed to cinnamic acid (Fig. Supplementary Notes 3.20).

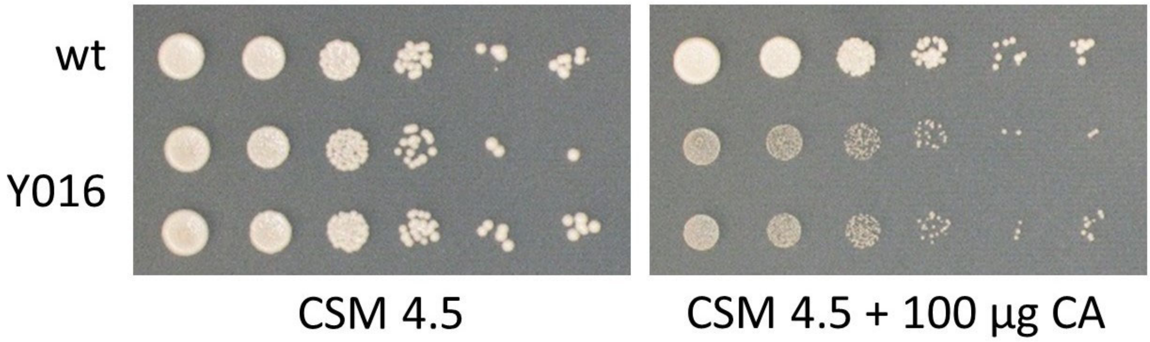

**Fig. Supplementary Notes 3.20. Targeted *fdc1* mutants in *Saccharomyces cerevisiae* are sensitive to cinnamic acid.** Two single cell isolates for the *FDC1* TaqMan sample Y016 (W159Stop) and the parental wild-type strain were grown for 16 h in yeast extract-peptone-dextrose medium at 30 °C. Cultures were adjusted to an $OD_{600}$ of 1 and another five 10-fold serial dilutions were performed. Each dilution (5 µl) was spotted onto complex synthetic medium pH 4.5 (CSM 4.5) and the same medium supplemented with 100 µg/ml cinnamic acid (CA). Plates were subsequently incubated at 30 °C for three to four days.

**Bacteria:**

In a *Lactobacillus pasteurii* (DSM 23707) library totalling approximately 1 million lines (Supplementary. Notes 1, Section 1; Supplementary Notes 2, Section 2) we targeted and identified two premature stop variants of the Galactose-1-phosphate uridylyltransferase gene, galT (*GalT*$^{W89*}$ and *GalT*$^{Q148*)}$ which mediates the second step of the Leloir pathway in which D-galactose is metabolised[35,36].

Together, these examples show the applicability of FIND-IT for targeting genetic variants in diverse genes, genetic element and across species.
