## Supplemental Table 1 for "FIND-IT: Ultrafast mining of genome diversity"

**Supplementary Table 1. Novel barley gene variants identified by FIND-IT.**

Identification of 155 barley variants with novel nucleotide substitutions causing premature stops, amino acid substitutions or changes of regulatory regions in target genes of interest.

| | Barley Gene ID | Genome location | Identified Variant |
|---|---|---|---|
| 1 | HORVU.MOREX.r2.1HG0005650 | chr1H:14296429-14298092 | W152* |
| 2 | HORVU.MOREX.r2.1HG0005660 | chr1H:14386006-14387653 | W58* |
| 3 | HORVU.MOREX.r2.1HG0005970 | chr1H:15161134-15162759 | W143* |
| 4 | HORVU.MOREX.r2.1HG0005990 | chr1H:15661663-15663425 | W137* |
| 5 | HORVU.MOREX.r2.1HG0006010 | chr1H:15671377-15673010 | W281* |
| 6 | HORVU.MOREX.r2.1HG0006010 | chr1H:15671377-15673010 | G387D |
| 7 | HORVU.MOREX.r2.1HG0013190 | chr1H:49443436-49446567 | W319* |
| 8 | HORVU.MOREX.r2.1HG0017860 | chr1H:82048286-82057404 | W111* |
| 9 | HORVU.MOREX.r2.1HG0019190 | chr1H:93118163-93121783 | W88* |
| 10 | HORVU.MOREX.r2.1HG0019190 | chr1H:93118163-93121783 | W301* |
| 11 | HORVU.MOREX.r2.1HG0038860 | chr1H:332135908-332140252 | W53* |
| 12 | HORVU.MOREX.r2.1HG0038870 | chr1H:332140981-332145711 | W346* |
| 13 | HORVU.MOREX.r2.1HG0039600 | chr1H:338188726-338191284 | W230* |
| 14 | HORVU.MOREX.r2.1HG0041980 | chr1H:359066146-359071013 | W313* |
| 15 | HORVU.MOREX.r2.1HG0047170 | chr1H:397790780-397794463 | W46* |
| 16 | HORVU.MOREX.r2.1HG0048100 | chr1H:405077941-405081372 | W210* |
| 17 | HORVU.MOREX.r2.1HG0051960 | chr1H:430403307-430404841 | W62* |
| 18 | HORVU.MOREX.r2.1HG0062960 | chr1H:483432737-483433878 | W113* |
| 19 | HORVU.MOREX.r2.1HG0066860 | chr1H:493915229-493921495 | W237* |
| 20 | HORVU.MOREX.r2.1HG0067690 | chr1H:495456953-495464334 | W15* |
| 21 | HORVU.MOREX.r2.2HG0079410 | chr2H:1727295-1739674 | W73* |
| 22 | HORVU.MOREX.r2.2HG0086340 | chr2H:18205047-18209880 | W131* |
| 23 | HORVU.MOREX.r2.2HG0086340 | chr2H:18205047-18209880 | G265D |
| 24 | HORVU.MOREX.r2.2HG0088780 | chr2H:25092220-25095704 | W178* |
| 25 | HORVU.MOREX.r2.2HG0091480 | chr2H:34645995-34647649 | W96* |
| 26 | HORVU.MOREX.r2.2HG0094840 | chr2H:49673203-49674327 | G110D |
| 27 | HORVU.MOREX.r2.2HG0096950 | chr2H:59028646-59034216 | W175* |
| 28 | HORVU.MOREX.r2.2HG0107310 | chr2H:128182530-128190128 | W170* |
| 29 | HORVU.MOREX.r2.2HG0107310 | chr2H:128182530-128190128 | W371* |
| 30 | HORVU.MOREX.r2.2HG0107310 | chr2H:128182530-128190128 | W431* |
| 31 | HORVU.MOREX.r2.2HG0107310 | chr2H:128182530-128190128 | W431* |
| 32 | HORVU.MOREX.r2.2HG0113880 | chr2H:196367510-196368424 | W42* |
| 33 | HORVU.MOREX.r2.2HG0115270 | chr2H:207285072-207286124 | W49* |
| 34 | HORVU.MOREX.r2.2HG0136670 | chr2H:465287027-465299202 | W218* |
| 35 | HORVU.MOREX.r2.2HG0138830 | chr2H:483868696-483871821 | W69* |
| 36 | HORVU.MOREX.r2.2HG0139520 | chr2H:489015186-489023414 | W86* |
| 37 | HORVU.MOREX.r2.2HG0151680 | chr2H:575913255-575913869 | W119* |
| 38 | HORVU.MOREX.r2.2HG0152930 | chr2H:581365914-581367006 | W169* |
| 39 | HORVU.MOREX.r2.2HG0160430 | chr2H:616161133-616164376 | P165S |
| 40 | HORVU.MOREX.r2.2HG0160430 | chr2H:616161133-616164376 | P165L |
| 41 | HORVU.MOREX.r2.2HG0160430 | chr2H:616161133-616164376 | E167K |
| 42 | HORVU.MOREX.r2.2HG0164090 | chr2H:628491325-628498606 | W393* |
| 43 | HORVU.MOREX.r2.2HG0164340 | chr2H:628985934-628987326 | W54* |
| 44 | HORVU.MOREX.r2.2HG0167990 | chr2H:638302138-638303184 | W232* |
| 45 | HORVU.MOREX.r2.3HG0188210 | chr3H:15398805-15400487 | W66* |

| | | | |
|---|---|---|---|
| 46 | HORVU.MOREX.r2.3HG0194740 | chr3H:37009034-37011619 | W49* |
| 47 | HORVU.MOREX.r2.3HG0196660 | chr3H:48708368-48710489 | W24* |
| 48 | HORVU.MOREX.r2.3HG0202940 | chr3H:101372717-101375861 | W161* |
| 49 | HORVU.MOREX.r2.3HG0202940 | chr3H:101372717-101375861 | W178* |
| 50 | HORVU.MOREX.r2.3HG0202950 | chr3H:102162183-102166377 | W343* |
| 51 | HORVU.MOREX.r2.3HG0204450 | chr3H:115820435-115822873 | W107* |
| 52 | HORVU.MOREX.r2.3HG0211410 | chr3H:185377055-185380367 | W52* |
| 53 | HORVU.MOREX.r2.3HG0212150 | chr3H:193583368-193588172 | W161* |
| 54 | HORVU.MOREX.r2.3HG0217410 | chr3H:261884890-261931076 | W171* |
| 55 | HORVU.MOREX.r2.3HG0227270 | chr3H:390955444-390956888 | W269* |
| 56 | HORVU.MOREX.r2.3HG0240680 | chr3H:498497030-498498813 | W106* |
| 57 | HORVU.MOREX.r2.3HG0242560 | chr3H:507973316-507974068 | W74* |
| 58 | HORVU.MOREX.r2.3HG0256480 | chr3H:572032746-572034295 | W52* |
| 59 | HORVU.MOREX.r2.3HG0257280 | chr3H:574702874-574706606 | W123* |
| 60 | HORVU.MOREX.r2.3HG0261150 | chr3H:589709313-589713500 | W85* |
| 61 | HORVU.MOREX.r2.3HG0261390 | chr3H:590169963-590173530 | W102* |
| 62 | HORVU.MOREX.r2.4HG0275880 | chr4H:173882-176015 | W113* |
| 63 | HORVU.MOREX.r2.4HG0280720 | chr4H:16668801-16671743 | W602* |
| 64 | HORVU.MOREX.r2.4HG0280720 | chr4H:16668801-16671743 | G130S |
| 65 | HORVU.MOREX.r2.4HG0280720 | chr4H:16668801-16671743 | G130D |
| 66 | HORVU.MOREX.r2.4HG0280720 | chr4H:16668801-16671743 | D170N |
| 67 | HORVU.MOREX.r2.4HG0280720 | chr4H:16668801-16671743 | G178D |
| 68 | HORVU.MOREX.r2.4HG0280720 | chr4H:16668801-16671743 | G192D |
| 69 | HORVU.MOREX.r2.4HG0280720 | chr4H:16668801-16671743 | V221I |
| 70 | HORVU.MOREX.r2.4HG0280720 | chr4H:16668801-16671743 | A226T |
| 71 | HORVU.MOREX.r2.4HG0280720 | chr4H:16668801-16671743 | A240T |
| 72 | HORVU.MOREX.r2.4HG0280720 | chr4H:16668801-16671743 | E244K |
| 73 | HORVU.MOREX.r2.4HG0280720 | chr4H:16668801-16671743 | A248T |
| 74 | HORVU.MOREX.r2.4HG0280720 | chr4H:16668801-16671743 | A251T |
| 75 | HORVU.MOREX.r2.4HG0280720 | chr4H:16668801-16671743 | A260T |
| 76 | HORVU.MOREX.r2.4HG0280720 | chr4H:16668801-16671743 | V264M |
| 77 | HORVU.MOREX.r2.4HG0280720 | chr4H:16668801-16671743 | G275D |
| 78 | HORVU.MOREX.r2.4HG0280720 | chr4H:15883636-15885492 | A277T |
| 79 | HORVU.MOREX.r2.4HG0280720 | chr4H:16668801-16671743 | V286M |
| 80 | HORVU.MOREX.r2.4HG0280720 | chr4H:16668801-16671743 | G298D |
| 81 | HORVU.MOREX.r2.4HG0280720 | chr4H:16668801-16671743 | G330D |
| 82 | HORVU.MOREX.r2.4HG0280720 | chr4H:16668801-16671743 | L346F |
| 83 | HORVU.MOREX.r2.4HG0280720 | chr4H:16668801-16671743 | A349T |
| 84 | HORVU.MOREX.r2.4HG0280720 | chr4H:16668801-16671743 | A353T |
| 85 | HORVU.MOREX.r2.4HG0280720 | chr4H:16668801-16671743 | G360E |
| 86 | HORVU.MOREX.r2.4HG0280720 | chr4H:16668801-16671743 | G370D |
| 87 | HORVU.MOREX.r2.4HG0280720 | chr4H:16668801-16671743 | D375N |
| 88 | HORVU.MOREX.r2.4HG0280720 | chr4H:16668801-16671743 | G377R |
| 89 | HORVU.MOREX.r2.4HG0280720 | chr4H:16668801-16671743 | P394L |
| 90 | HORVU.MOREX.r2.4HG0280720 | chr4H:15883636-15885492 | R481H |
| 91 | HORVU.MOREX.r2.4HG0280720 | chr4H:16668801-16671743 | S488F |
| 92 | HORVU.MOREX.r2.4HG0280720 | chr4H:16668801-16671743 | T524I |
| 93 | HORVU.MOREX.r2.4HG0280720 | chr4H:16668801-16671743 | G563D |
| 94 | HORVU.MOREX.r2.4HG0280720 | chr4H:16668801-16671743 | E579K |
| 95 | HORVU.MOREX.r2.4HG0281260 | chr4H:18575867-18578536 | G287D |

| | | | |
|---|---|---|---|
| 96 | HORVU.MOREX.r2.4HG0289460 | chr4H:73680496-73682091 | W46* |
| 97 | HORVU.MOREX.r2.4HG0290210 | chr4H:82552859-82562826 | W179* |
| 98 | HORVU.MOREX.r2.4HG0315810 | chr4H:413053908-413055642 | W65* |
| 99 | HORVU.MOREX.r2.4HG0315870 | chr4H:414475706-414476933 | W103* |
| 100 | HORVU.MOREX.r2.4HG0320050 | chr4H:461499644-461502515 | W218* |
| 101 | HORVU.MOREX.r2.4HG0332170 | chr4H:560241643-560242657 | W106* |
| 102 | HORVU.MOREX.r2.4HG0341030 | chr4H:599987252-599988397 | W136* |
| 103 | HORVU.MOREX.r2.4HG0342440 | chr4H:604389161-604392593 | W315* |
| 104 | HORVU.MOREX.r2.4HG0345110 | chr4H:613186867-613190782 | W102* |
| 105 | HORVU.MOREX.r2.4HG0345290 | chr4H:613477070-613481667 | W206* |
| 106 | HORVU.MOREX.r2.4HG0345290 | chr4H:613477070-613481667 | P224S |
| 107 | HORVU.MOREX.r2.5HG0354200 | chr5H:14656004-14664743 | W359* |
| 108 | HORVU.MOREX.r2.5HG0354290 | chr5H:15562611-15564434 | W421* |
| 109 | HORVU.MOREX.r2.5HG0366630 | chr5H:99679211-99683078 | W224* |
| 110 | HORVU.MOREX.r2.5HG0372420 | chr5H:165230529-165253645 | W226* |
| 111 | HORVU.MOREX.r2.5HG0383660 | chr5H:318783743-318785113 | W113* |
| 112 | HORVU.MOREX.r2.5HG0386220 | chr5H:340792835-340793836 | W70* |
| 113 | HORVU.MOREX.r2.5HG0392730 | chr5H:399761478-399763004 | G133D |
| 114 | HORVU.MOREX.r2.5HG0398480 | chr5H:439858216-439867112 | W2* |
| 115 | HORVU.MOREX.r2.5HG0402930 | chr5H:467079543-467082282 | W116* |
| 116 | HORVU.MOREX.r2.5HG0402970 | chr5H:467395763-467396491 | W130* |
| 117 | HORVU.MOREX.r2.5HG0421990 | chr5H:529801624-529803402 | W242* |
| 118 | HORVU.MOREX.r2.5HG0425120 | chr5H:538916691-538923734 | W263* |
| 119 | HORVU.MOREX.r2.5HG0429500 | chr5H:551480093-551480959 | W82* |
| 120 | HORVU.MOREX.r2.5HG0429900 | chr5H:553107429-553116126 | W268* |
| 121 | HORVU.MOREX.r2.5HG0431910 | chr5H:558457293-558459137 | W248* |
| 122 | HORVU.MOREX.r2.5HG0434680 | chr5H:565291916-565296808 | W200* |
| 123 | HORVU.MOREX.r2.5HG0447180 | chr5H:596729695-596737128 | W270* |
| 124 | HORVU.MOREX.r2.6HG0462400 | chr6H:50083893-50090159 | G271D |
| 125 | HORVU.MOREX.r2.6HG0462530 | chr6H:51261970-51263563 | W91* |
| 126 | HORVU.MOREX.r2.6HG0462530 | chr6H:51261970-51263563 | V70M |
| 127 | HORVU.MOREX.r2.6HG0462530 | chr6H:51261970-51263563 | V49I |
| 128 | HORVU.MOREX.r2.6HG0462530 | chr6H:51261970-51263563 | D122N |
| 129 | HORVU.MOREX.r2.6HG0462530 | chr6H:51261970-51263563 | P364L |
| 130 | HORVU.MOREX.r2.6HG0479300 | chr6H:190184822-190186307 | P180S |
| 131 | HORVU.MOREX.r2.6HG0479430 | chr6H:191265639-191266365 | W33* |
| 132 | HORVU.MOREX.r2.6HG0483720 | chr6H:246535403-246560072 | W233* |
| 133 | HORVU.MOREX.r2.6HG0491380 | chr6H:351537513-351538087 | W18* |
| 134 | HORVU.MOREX.r2.6HG0510160 | chr6H:515913031-515916013 | W130* |
| 135 | HORVU.MOREX.r2.6HG0512130 | chr6H:526695485-526696955 | TTGA[C/T]C |
| 136 | HORVU.MOREX.r2.6HG0519800 | chr6H:555348843-555354517 | W171* |
| 137 | HORVU.MOREX.r2.7HG0532670 | chr7H:10836462-10840039 | W378* |
| 138 | HORVU.MOREX.r2.7HG0532670 | chr7H:10836462-10840039 | G348D |
| 139 | HORVU.MOREX.r2.7HG0535060 | chr7H:15977348-15980148 | W126* |
| 140 | HORVU.MOREX.r2.7HG0543180 | chr7H:40203741-40210284 | W332* |
| 141 | HORVU.MOREX.r2.7HG0543180 | chr7H:40203741-40210284 | G463D |
| 142 | HORVU.MOREX.r2.7HG0543180 | chr7H:40203741-40210284 | G111S |
| 143 | HORVU.MOREX.r2.7HG0553070 | chr7H:91928500-91938490 | W277* |
| 144 | HORVU.MOREX.r2.7HG0553070 | chr7H:91928500-91938490 | W334* |
| 145 | HORVU.MOREX.r2.7HG0553070 | chr7H:91928500-91938490 | G623E |

| | | | |
|---|---|---|---|
| 146 | HORVU.MOREX.r2.7HG0553990 | chr7H:98808452-98810950 | A274T |
| 147 | HORVU.MOREX.r2.7HG0561010 | chr7H:147466779-147471410 | W250* |
| 148 | HORVU.MOREX.r2.7HG0564420 | chr7H:175704026-175708788 | W194* |
| 149 | HORVU.MOREX.r2.7HG0573190 | chr7H:265532875-265539857 | W65* |
| 150 | HORVU.MOREX.r2.7HG0579230 | chr7H:367614501-367619640 | W676* |
| 151 | HORVU.MOREX.r2.7HG0579230 | chr7H:367614501-367619640 | G847E |
| 152 | HORVU.MOREX.r2.7HG0579230 | chr7H:367614501-367619640 | G748D |
| 153 | HORVU.MOREX.r2.7HG0579230 | chr7H:367614501-367619640 | G732D |
| 154 | HORVU.MOREX.r2.7HG0596820 | chr7H:529060345-529061228 | W16* |
| 155 | HORVU.MOREX.r2.7HG0616560 | chr7H:617743377-617758724 | W306* |
