## Supplemental Table 2 for "FIND-IT: Ultrafast mining of genome diversity"

**Supplementary Table 2. Sequencing primers for validation of FIND-IT variants.**

| | | | |
|---|---|---|---|
| HORVU.MOREX.r2.7HG0596820 (*nud* $^{W16*}$ ) | | | |
| Fwd | CATTCCTTCCCTCCGCTCTG | Rev | TCTCAGGACACGGTGGGATA |
| HORVU.MOREX.r2.7HG0579230 (*cslF6* $^{W676*}$ ) | | | |
| Fwd | CAAAGAGATCGGCTGGGTGT | Rev | TGAGGAAGATGGCGGTGAAG |
| HORVU.MOREX.r2.1HG0041980 (*jmj706* $^{W313*}$ ) | | | |
| Fwd | TCAATTATCATCATTGTGGGGCA | Rev | GCATCCTCACCATCACCAGT |
| HORVU.MOREX.r2.7HG0535060 (*gbss1* $^{W126*}$ ) | | | |
| Fwd | CCATTACAACTGACGGCGTG | Rev | TCCAGCCAGTCGGTAGAATC |
| HORVU.MOREX.r2.6HG0483720 (*gw2* $^{W233*}$ ) | | | |
| Fwd | TCATGCTGTAGGTGTGTGACA | Rev | ACAAGTCGCATTCATAACATTATTCA |
| HORVU.MOREX.r2.4HG0280720 (*sln1* $^{A277T}$ ) | | | |
| Fwd | GTGGTCGACACGCAGGAG | Rev | CTTGATGCCGAAGTCGACGA |
| HORVU.MOREX.r2.7HG0579230 (*cslF6* $^{G847E}$ ) | | | |
| Fwd | CAAGTTGCTCCGCCTACCTC | Rev | ATGACGAAGGTGAATGCCCA |
| HORVU.MOREX.r2.7HG0579230 (*cslF6* $^{G748D}$ ) | | | |
| Fwd | CCCATCAACCTCACGGAGAG | Rev | ATCACCTTGGTCAGCACCTG |
| HORVU.MOREX.r2.7HG0579230 (*cslF6* $^{T709I}$ ) | | | |
| Fwd | CCCATCAACCTCACGGAGAG | Rev | ATCACCTTGGTCAGCACCTG |
| HORVU.MOREX.r2.7HG0579230 (*cslF6* $^{G732D}$ ) | | | |
| Fwd | CCCATCAACCTCACGGAGAG | Rev | ATCACCTTGGTCAGCACCTG |
| HORVU.MOREX.r2.2HG0160430 (*GRF4* $^{variant3}$ ) | | | |
| Fwd | AGCATTCTTGACGTGGGTGT | Rev | ATCTTTGTTTCCGGCAGCAC |
| (*Amy1_1-var* ) | | | |
| Fwd | TGCTCGAATGTTCTGTCCTCA | Rev | AGTACGATGGAGAACTGATGCA |
