## Supplemental Table 3 for "FIND-IT: Ultrafast mining of genome diversity"

**Supplementary Table 3. Statistical summary of barley variant agronomic evalutation.**

**Paired two-sided t-tests were performed to obtain P-values**

| Gene (variant) | Seed Harvest Year | Evaluation | *n* | Type[a] | Mean | SD | df | T-statistic | P-Value | Figure reference |
|---|---|---|---|---|---|---|---|---|---|---|
| ***Nud* (*W16**)** | DK 2019 | TKW [g] | *4* | Var | 46,70 | 0,644 | 3 | 15,717 | *** P = 5,6 *10$^{-4}$ | Fig. 2d, Supp. 3.8b |
| | | | | c | 55,64 | 1,469 | | | | |
| | | Grain yield [kg] | *3* | Var | 5,62 | 0,076 | 2 | 11,048 | ** P = 8,09 *10$^{-3}$ | Fig. 2e, Supp. 3.8e |
| | | | | c | 7,12 | 0,197 | | | | |
| ***JMJ706* (*W313**)** | DK 2020 | TKW [g] | *2* | Var | 56,59 | 0,057 | 1 | 4,743 | ns P = 1,32 *10$^{-1}$ | Fig. 2d, Supp. 3.6b |
| | | | | c | 58,07 | 0,368 | | | | |
| | | Grain yield [kg] | *2* | Var | 6,27 | 0,014 | 1 | 15,800 | * P = 4,02 *10$^{-2}$ | Fig. 2e |
| | | | | c | 8,44 | 0,209 | | | | |
| ***GW2* (*W233**)** | DK 2020 | TKW [g] | *2* | Var | 64,24 | 0,204 | 1 | -12,769 | * P = 4,98 x 10$^{-2}$ | Fig. 2d, Supp. 3.11b |
| | | | | c | 57,38 | 0,556 | | | | |
| | | Grain yield [kg] | *2* | Var | 7,44 | 0,145 | 1 | -1,303 | ns P = 4,17 x 10$^{-1}$ | Fig. 2e, Supp. 3.11e |
| | | | | c | 7,33 | 0,262 | | | | |
| ***GBSS1* (*W126**)** | NZ 2018 | TKW [g] | *3* | Var | 55,32 | 1,526 | 2 | 0,531 | ns P = 6,48 x 10$^{-1}$ | Fig. 2d, Supp. 3.4c |
| | | | | c | 55,87 | 0,387 | | | | |
| | | Grain yield [kg] | *3* | Var | 5,06 | 0,459 | 2 | 1,278 | ns P = 3,3 x 10$^{-1}$ | Fig. 2e, Supp. 3.4f |
| | | | | c | 5,34 | 0,842 | | | | |
| | | Starch amylose [%] | *3* | Var | 2,64 | 0,737 | 2 | 39,185 | *** P = 6,51 x 10$^{-4}$ | Fig. 2g, Supp. 3.4b |
| | | | | c | 23,41 | 0,074 | | | | |
| ***SLN1* (*A277T*)** | DK 2019 | TKW [g] | *4* | Var | 58,74 | 1,454 | 3 | -12,896 | ** P = 1,01 x 10$^{-3}$ | Fig. 3d, Supp. 3.13b |
| | | | | c | 55,11 | 0,942 | | | | |
| | | Grain yield [kg] | *4* | Var | 6,12 | 0,149 | 3 | 0,255 | ns P = 8,15 x 10$^{-1}$ | Supp. 3.13e |
| | | | | c | 6,15 | 0,178 | | | | |
| | | Plant height [cm] | *3* | Var | 64,00 | 5,560 | 2 | -1,674 | ns P = 2,36 x 10$^{-1}$ | Fig. 3b, Supp. 3.13a |
| | | | | c | 55,67 | 2,082 | | | | |
| ***CslF6* (*W676**)** | NZ 2018 | TKW [g] | *3* | Var | 42,65 | 1,329 | 2 | 7,138 | * P = 1,91 x 10$^{-2}$ | Fig. 2d, Supp. 3.17c |
| | | | | c | 51,33 | 0,995 | | | | |
| | | Grain yield [kg] | *3* | Var | 3,67 | 0,260 | 2 | 6,530 | * P = 2,27 x 10$^{-2}$ | Fig. 2e, Supp. 3.17f |
| | | | | c | 5,60 | 0,296 | | | | |
| | | β-Glucan content [μg/mg] | *3* | Var | 0,57 | 0,021 | 2 | -58,478 | *** P = 2,92 x 10$^{-4}$ | Fig. 2f, Supp. 3.17b |
| | | | | c | 34,79 | 0,791 | | | | |
| ***CslF6* (*T709I*)** | DK 2017 | TKW [g] | *4* | Var | 49,79 | 0,790 | 3 | 3,627 | 5* P = 3,61 x 10$^{-2}$ | Supp. 3.19b |
| | | | | c | 51,87 | 1,331 | | | | |
| | | Grain yield [kg] | *5* | Var | 5,92 | 0,161 | 4 | -1,542 | ns P = 1,98 x 10$^{-1}$ | Fig. 3l, Supp. 3.19f |
| | | | | c | 6,07 | 0,278 | | | | |
| | | Broken grain [%] | *4* | Var | 3,22 | 0,597 | 3 | -6,239 | ** P = 8,31 x 10$^{-3}$ | Fig. 3k, Supp. 3.19e |
| | | | | c | 2,13 | 0,294 | | | | |
| | | β-Glucan content [μg/mg] | *3* | Var | 13,93 | 0,318 | 2 | 17,990 | ** P = 3,08 x 10$^{-3}$ | Fig. 3j, Supp. 3.19a |
| | | | | c | 31,91 | 1,419 | | | | |
| ***CslF6* (*T748D*)** | DK 2017 | TKW [g] | *3* | Var | 55,05 | 1,588 | 2 | 0,882 | ns P = 4,71 x 10$^{-1}$ | Supp. 3.19b |
| | | | | c | 56,10 | 0,555 | | | | |
| | | Grain yield [kg] | *3* | Var | 6,35 | 0,080 | 2 | 0,559 | ns P = 6,32 x 10$^{-1}$ | Fig. 3l, Supp. 3.19f |
| | | | | c | 6,39 | 0,083 | | | | |
| | | Broken grain [%] | *4* | Var | 2,73 | 0,455 | 3 | -2,931 | ns P = 6,09 x 10$^{-2}$ | Fig. 3k, Supp. 3.19e |
| | | | | c | 1,30 | 0,558 | | | | |
| | | β-Glucan content [μg/mg] | *3* | Var | 18,92 | 0,547 | 2 | 19,144 | ** P = 2,72 x 10$^{-3}$ | Fig. 3j, Supp. 3.19a |
| | | | | c | 34,79 | 0,890 | | | | |
| ***CslF6* (*T847E*)** | DK 2017 | TKW [g] | *5* | Var | 55,03 | 2,034 | 4 | -0,132 | ns P = 9,01 x 10$^{-1}$ | Supp. 3.19b |
| | | | | c | 54,84 | 1,180 | | | | |
| | | Grain yield [kg] | *5* | Var | 6,16 | 0,288 | 4 | -0,646 | ns P = 5,53 x 10$^{-1}$ | Fig. 3l, Supp. 3.19f |
| | | | | c | 6,08 | 0,116 | | | | |
| | | Broken grain [%] | *5* | Var | 4,44 | 1,532 | 4 | -3,524 | * P = 2,44 x 10$^{-2}$ | Fig. 3k, Supp. 3.19e |
| | | | | c | 1,63 | 0,698 | | | | |
| | | β-Glucan content [μg/mg] | *3* | Var | 17,47 | 0,043 | 2 | 35,323 | *** P = 8,01 x 10$^{-4}$ | Fig. 3j, Supp. 3.19a |
| | | | | c | 34,79 | 0,890 | | | | |
| **cv. Planet** | NZ 2019 | Expression of Amy genes 48 h [Copies x 10^3/μl cDNA] | *3* | Amy1_1 | 27,72 | 7,145 | 2 | -3,301 | * P = 8,08 x 10$^{-2}$ | Fig. 3g, Supp. 3.15b |
| | | | | Amy1_2 | 55,26 | 14,678 | | | | |
| | | | | Amy1_1 | 27,72 | 7,145 | 2 | 6,147 | * P = 2,55 x 10$^{-2}$ | |
| | | | | Amy2 | 2,072 | 0,360 | | | | |
| | | | | Amy1_2 | 55,26 | 14,678 | 2 | 6,416 | * P = 2,34 x 10$^{-2}$ | |
| | | | | Amy2 | 2,072 | 0,360 | | | | |
| | | Expression of Amy genes 72 h [Copies x 10^3/μl cDNA] | *3* | Amy1_1 | 41,27 | 8,197 | 2 | -6,590 | * P = 2,23 x 10$^{-2}$ | Fig. 3g, Supp. 3.15b |
| | | | | Amy1_2 | 90,61 | 12,284 | | | | |
| | | | | Amy1_1 | 41,27 | 8,197 | 2 | 7,616 | * P = 1,68 x 10$^{-2}$ | |
| | | | | Amy2 | 8,536 | 1,486 | | | | |
| | | | | Amy1_2 | 90,61 | 12,284 | 2 | 10,689 | ** P = 8,64 x 10$^{-3}$ | |
| | | | | Amy2 | 8,536 | 1,486 | | | | |
| **cv. Planet & *Amy1_1-variant*** | NZ 2019 | Expression of Amy genes 48 h [Copies x 10^3/μl cDNA] | *3* | Amy1_1-Var | 30,27 | 7,477 | 2 | -1,515 | ns P = 2,69 x 10$^{-1}$ | Fig. 3h, Supp. 3.15c |
| | | | | Amy1_1 | 27,72 | 7,145 | | | | |
| | | Expression of Amy genes 48 h [Copies x 10^3/μl cDNA] | *3* | Amy1_1-Var | 67,14 | 10,896 | 2 | -4,808 | * P = 4,06 x 10$^{-2}$ | Fig. 3h, Supp. 3.15c |
| | | | | Amy1_1 | 41,27 | 8,197 | | | | |

ns **Statistical not significant**

* **Statistically significant at the 5 % level**

* * **Statistically significant at the 1 % level**

*** **Statistically significant at the 0.1 % level**

[a] Var = Genetic variant, c = control
